# A Cell Model for the Detection of a Planar Surface Using Motion Stereo in Area MST of the Visual Cortex

**DOI:** 10.1101/2024.01.16.575918

**Authors:** Susumu Kawakami

**Affiliations:** Tohoku University, Research Institute of Electrical Communication, 2-1-1 Katahira, Aoba-ku, Sendai 980-8577, Japan

## Abstract

I propose a series of modeled cells for detecting three plane parameters (i.e. a time-to-contact to a plane, its orientation, and its shortest distance) with motion stereo. This series is composed of lateral geniculate nucleus cells, nondirectionally selective simple cells, directionally selective (DS) simple cells, DS complex cells, motion-detection cells, and planar-surface-detection cells. These cell types perform the time delay, Hough transform, spatio-temporal correlation, accumulation, inverse Hough transform, and a combinations of the cross-ratio and polar transforms (or a small-circle transform) to detect these plane parameters, respectively. Each connection of the neural network connecting this series of cell types is modeled mathematically one by one using these transforms. The selective responses of SDCs to planar parameters are consistent with those of physiological experiments.

## 1. Introduction

The purpose of this paper is to model the cells, using motion stereo, that perceive the plane parameters of Figure 1 (i.e. the time-to-contact T to a plane, its three-dimensional (3D) orientation n, and its shortest distance D): the cells are thought to exist in areas such as the medial superior temporal (MST) area of the visual cortex. The pilot of Figure 1 recognizes this orientation n of a runway (i.e. a plane) to control the attitude of the airplane. Then, when this time T becomes appropriate, the pilot raises the nose of the airplane to land while recognizing the distance D.

**Figure 1.**
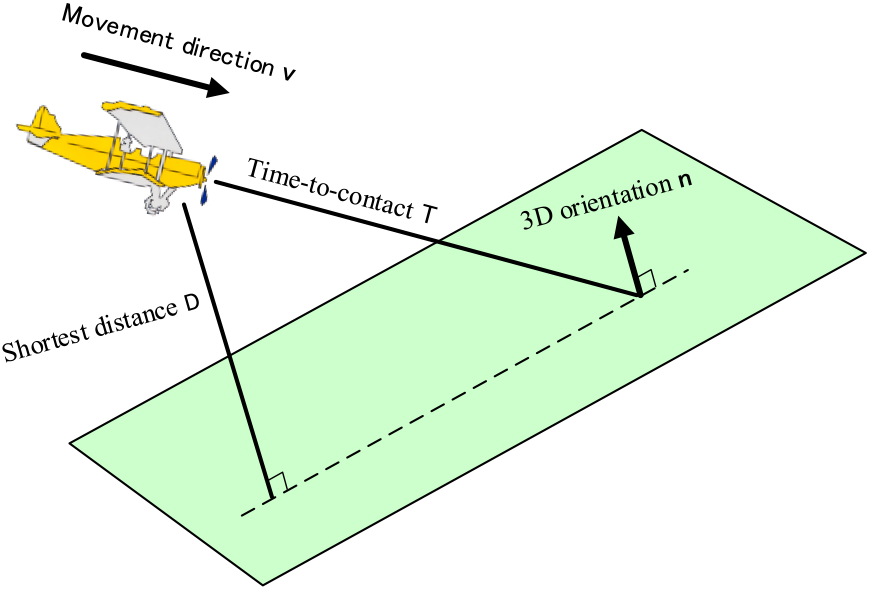
Recognition of three plane parameters. A pilot recognizes the plane parameters (i.e. the time-to-contact T to a plane, its 3D orientation n, and the shortest distance D to it) by motion stereo to land on a runway (i.e. a plane). That is, he recognizes the 3D direction n of the runway to control the attitude of the airplane. Then, when this time T becomes appropriate, he raises the nose of the airplane to land while recognizing the shortest distance D.

Although we previously reported this purpose (Kawakami et al., 2003, 2000), we were unable to fully explain it due to page limit. Therefore, this paper enabels this purpose to be explained in detail from a different point of view. Further, this paper allows Figure 19 in Kawakami et al. (2020) to be described in detail: this figure proposed for the first time, as far as we knew, a series of cell models that unified the form recognition and space recognition; the motion and binocular stereos in Figures 19(B) and 20(B) belong to the space recognition.

First, how these plane parameters have been thought of is explained. Gibson (1950) discovered that a plane is recognized by its optical flow pattern (OFP) that is generated on the retina: Figure 2 shows the OFP proposed by Gibson. This OFP is composed of local motions (i.e. motion vectors), each of which is caused within a receptive fields (RF) by self-locomotion. The magnitude of each local motion increases as the airplane approaches the runway (i.e. the plane), and thus its magnitude corresponds to the time-to-contact T to the plane. Also, when the plane tilts (or when the airplane’s tilt relative to the plane changes), the shape of this OFP changes and thus the shape corresponds to the 3D orientation n of the plane. However, the shortest distance D has not been mentioned by Gibson.

**Figure 2.**
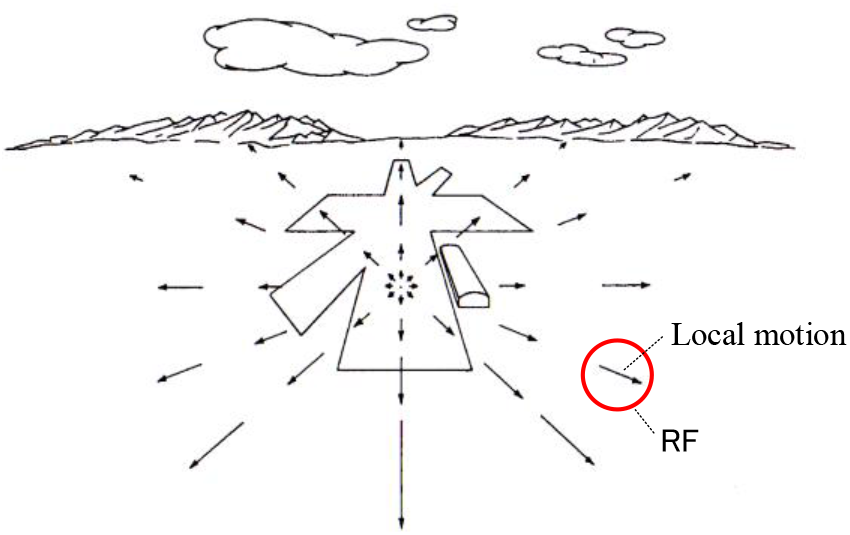
Optical flow pattern (OFP). The pilot’s retina in flight causes an OFP. This OFP consists of local motions, each of which occurs in a small region (i.e. a receptive field (RF)) on his retina. This figure was quoted by altering Gibson (1950).

A new perspective, different from this Gibson’s discovery, is described as follows. I tried to think psychologically about how I perceives the time-to-contact to the plane and its 3D direction. As shown in Figure 3, consider a scene in which a person walks toward a wall (i.e. a plane) in a movement direction v. I discovered that the wall always looks as a plane, but appears to degenerate into a straight line instead of the plane only at the moment he crosses the plane (shown in red). Based on this discovery, I realized that I can recognize the time-to-contact to the plane and its 3D orientation. That is, if the time to cross this plane can be predicted, this time is the time-to-contact T. Also, the normal vector n of the red plane at the moment he crossed it is the 3D orientation of the plane to be detected. Therefore, knowing the moment of crossing the plane enables the time-to-contact and 3D orientation to be detected. Note that the shortest distance D to the plane will be described in Section **3** later.

**Figure 3.**
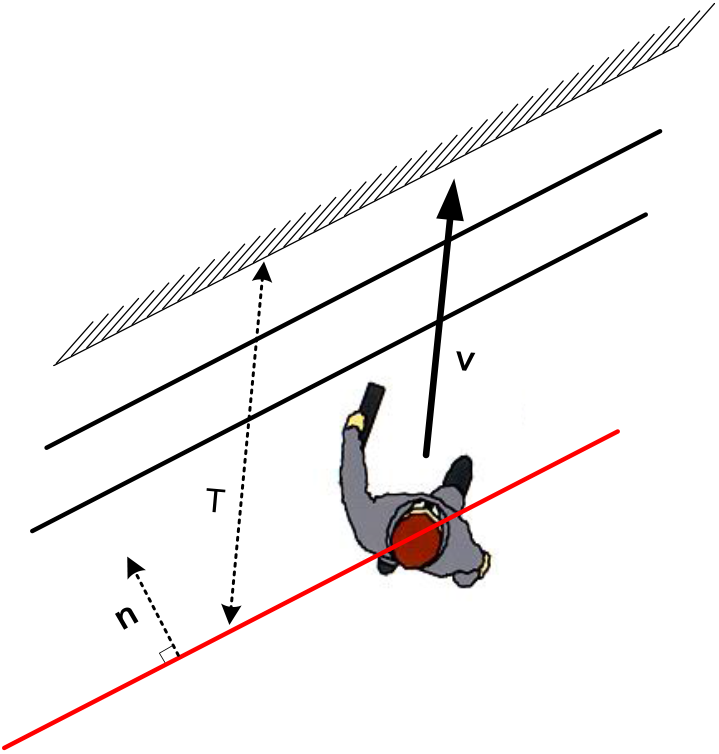
Focus on the moment when a person crosses a planar surface. Consider a scene in which a person walks toward a wall (i.e. a plane) in a direction v. I discovered that the wall always looks as a plane, but appears to degenerate into a straight line instead of the plane only at the moment he crosses the plane (shown in red). Based on this discovery, I realized that the time-to-contact to a plane and its 3D orientation are able to be recognized. That is, if the time to cross this plane can be predicted, this time is the time-to-contact T to the plane. Also, the normal vector n of the red plane at the moment he crossed it is the 3D orientation of the plane to be detected. Therefore, knowing the moment of crossing the plane enables the time-to-contact and 3D orientation to be detected.

I had two experiences related to this moment of crossing the plane. The first experience is as follows. Transmission towers are often arranged in a row to form a plane (Figure 4(A)). I have had many times the experience of being startled (or surprised) on a high speed train to see a straight line instead of the plane when I cross the plane, and to think that I have broken through this plane. This made me psychologically convinced that there was a mechanism in the cerebrum to detect that moment, and that it was possible to detect the time-to-contact to the plane and its 3D direction. The second experience is as follows. Similarly, utility poles are often arranged in a row to form a plane (Figure 4(B)). The moment I crossed that plane in a car, I had many times the experience of being startled (or surprised) to see a straight line instead of the plane, and to think that I have broken through this plane. This experience reinforced the above convictions.

**Figure 4.**
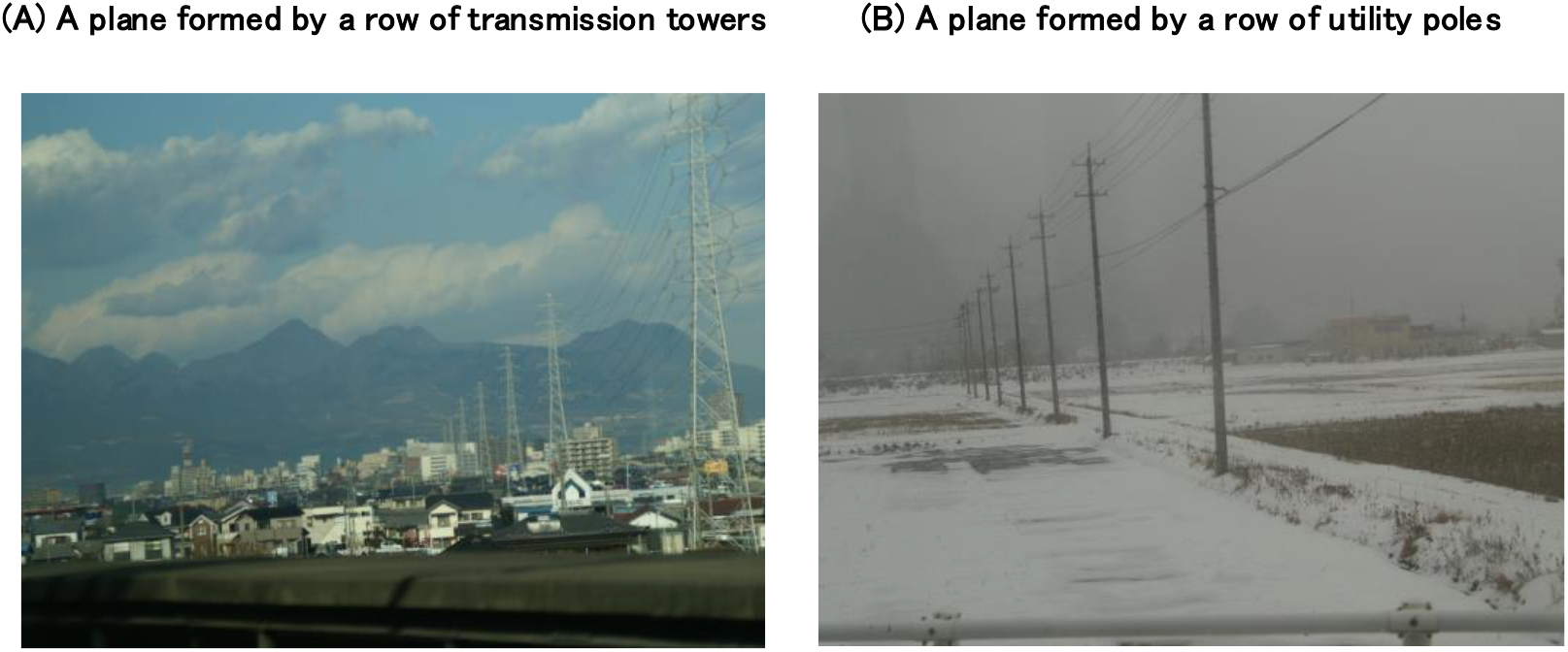
Experiences of the moment when crossing a plane. I had two experiences related to the moment of crossing the plane in Figure 3. The (A) shows the first experience. Transmission towers are often arranged in a row to form a plane. The moment I crossed that plane on a high-speed train, I had a startling (or surprising) experience that it looked like a straight line instead of the plane, and that I had broken through this plane. The (B) shows another experience. Utility poles are often arranged in a row to form a plane. The moment I crossed that plane in a car, I had a startling (or surprising) experience that it looked like a straight line instead of the plane, and that I had broken through this plane. Based on these, I was psychologically convinced that there is a mechanism in my cerebrum that detects that moment, and that this mechanism enables the time-to-contact to the plane and its 3D orientation to be detected.

These startled moments were, exactly, not the moment of crossing, but just a moment before that. That is, the moments were about 1 second before the crossing and were fairly constant regardless of movement speeds. This time (i.e. this moment) is almost the same as the time it takes for a gannet to fold its wings just before it plunges into the sea (Lee & Reddish, 1981). Also, Section **4.2.2.2** will show that there are cells, in area MST of monkeys, that detect this time of about 1 second. Section **2.2** will enable a series of cells that detects this moment to be modeled mathematically, and Section **4.2.1** will show that this model can explain the wing-folding time above.

Next, let us think about how this moment of Figure 3 that appears to be the straight line instead of the plane is reflected on the eyeball (Figure 5). At the current time T_0_ and the next time T_1_, triangles on the plane also appear as triangles on the eyeball. However, only at the moment T_C_ when the plane crosses the center of the eyeball, it appears to degenerate into a pink great circle instead of a triangle on the eyeball: this circle is the largest circle on the eyeball, which corresponds to a straight line on the plane. This degeneracy occurs with any shape on the plane, not just triangles. Extending this degeneracy causes every plane in space to degenerate into a great circle on the eyeball.

**Figure 5.**
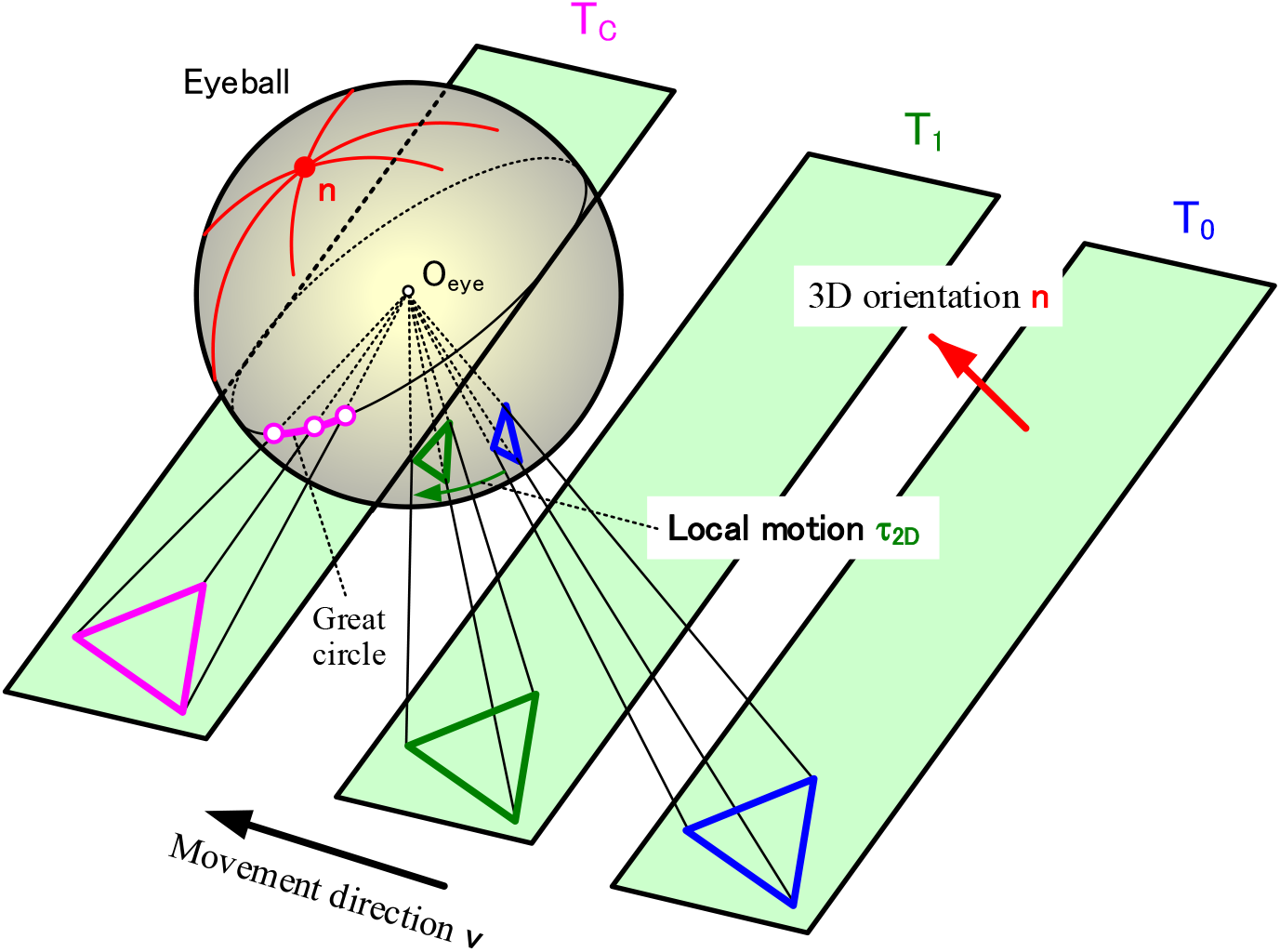
The moment when a triangle on the eyeball degenerates into a great circle. Let us consider how the moment of Figure 3 that appears to be the straight line instead of the plane is reflected on the eyeball. At the current time T_0_ and the next time T_1_, triangles on the plane also appear as triangles on the eyeball, respectively. However, only at the moment T_C_ when the plane crosses the center O_eye_ of the eyeball, it appears to degenerate into a pink great circle instead of the triangle on the eyeball: this circle is the largest circle on the eyeball, and corresponds to a straight line on the plane. This corresponds to the fact in Figures 3 and 4 that the plane appeared to be a straight line. Therefore, at this moment, the plane is seen as a pink great-circle on the eyeball. Detecting the moment T_C_ when this great circle appears enabls the time-to-contact T to be known as T_C_-T_0_. In addition, the 3D orientation n of the plane can be known as follows. Polar transformation (see Figure 6) of any three pink points on the great circle, into which the plane was degenerated at the time T_C_, results in three red great-circles: this transformation is such a transformation that converts the north pole to the equator, assuming that the eyeball is the earth. These red great-circles intersect at one point n. Thus, the 3D orientation n of the plane can be known as a direction from the eyeball center O_eye_ to this intersection.

If you can detect the moment T_C_ when this great circle appears, you can know the time-to-contact T as T_C_-T_0_. In addition, you can know the 3D orientation n as follows (Figure 5). Polar transformation of 3 points on this pink great circle at the time T_C_ results in 3 red great circles: this polar transform (Gurevic, 1962) corresponds to drawing an equator, assuming each of these points as a north pole (see Figure 6 and Section **4.5.3** for details). These 3 great circles intersect at one point n. The orientation connecting this intersection and the eyeball center O_eye_ is the 3D orientation n of the plane. In this way, drawing the red great circles enables the 3D orientation to be detected as the intersection n of these great circles.

**Figure 6.**
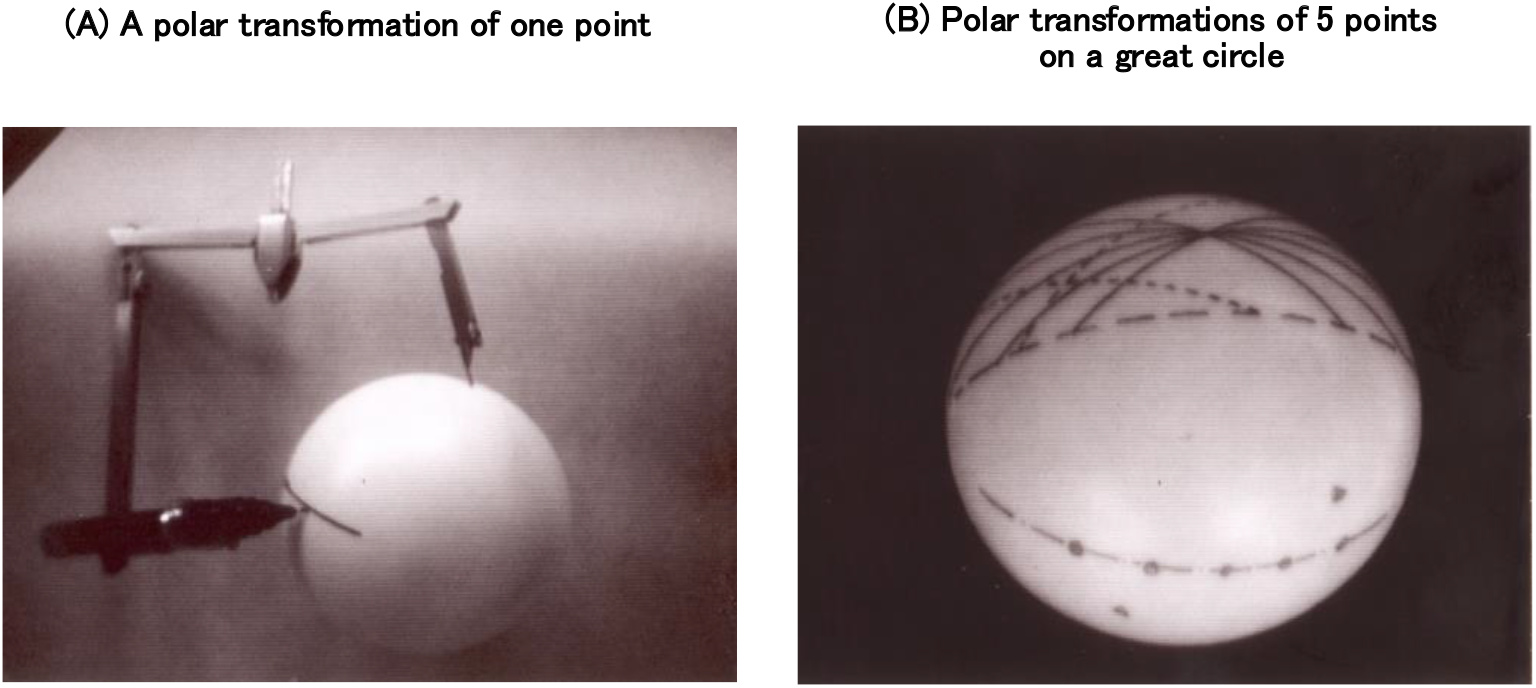
Drawing of polar transformations. The polar transformation described in Figure 5 was drawn on a plastic ball using a compass. In the (A), a great circle was drawn by placing the fulcrum of the compass at any position of the ball. This allowed me to draw the polar transformation on the ball. Assuming that this ball is the earth, it is equivalent to drawing the equator with the north pole as a fulcrum. Then, in the (B), five great circles were drawn by polar transformations with fulcrums at five points on the great circle that is shown in the bottom of the (B). These great circles intersected at one point as shown at the top of the (B). In this way, drawing with a compass has enabled the polar transformations and the intersection of great circles to be confirmed. Note that great circles (shown as dotted lines), which were drawn by a polar transformation of two points (shown as small triangles) outside this great circle above, does not pass through this intersection.

This intersection was drawn on the plastic ball using a compass, as follows (Figure 6). First, the polar transformation of a point was drawn. As shown in the (A), by placing the fulcrum of the compass at an arbitrary position on the ball and drawing a largest circle whose radius corresponded to 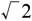 times that of the ball, a great circle could be drawn. This shows the drawing of the polar transformation. Next, by the polar transformations with five points on a great circle of the (B) as fulcrums, five great circles were drawn which intersected at one point. In this way, drawing with a compass enabled the polar transformations and the intersection of the great circles of to be confirmed. Note that the rough dotted line in the upper part of Figure 6(B) indicates the boundary of each hemisphere where points or great circles are drawn.

Each straight line (i.e. each side of the triangle) on the plane in Figure 5 is projected as a great circle on the eyeball. This projection was confirmed by taking a photograph (Figure 7). This photograph was taken from a height of about 10 cm on a tiled floor using a fisheye lens (see Section **2.1.3**) that can capture an image equivalent to the eyeball. It seems that someone is riding a ball, but the author’s shoes were photographed with the fisheye lens. The straight lines of the tiled floor appear as great circles, and the floor (i.e. the plane) looks like a sphere.

**Figure 7.**
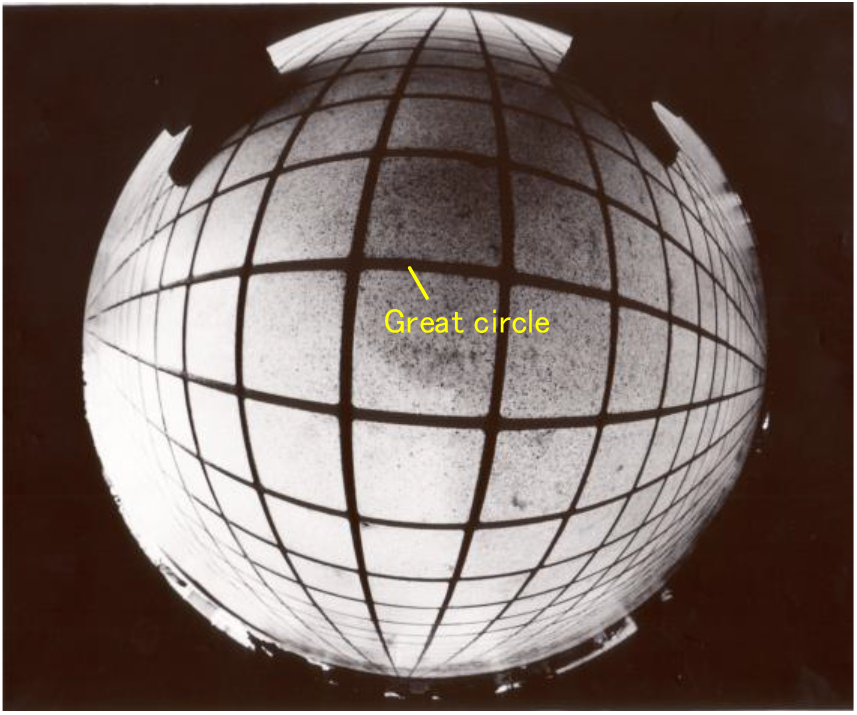
Photographing a floor with a fisheye lens. A tiled floor was photographed from a height of about 10 cm using a fish-eye lens that is capable of capturing an image equivalent to the eyeball. Straight lines of the tiled floor appear as great circles, making the floor (i.e. the plane) look like a sphere. In this way, it has been confirmed that straight lines in space appears as great circles on the eyeball.

In this way, I confirmed that each straight line in space appears as a great circle on the eyeball.

To summarize the above, if the moment T_C_ in which the plane in space degenerates into a pink great circle on the eyeball can be known (see Figure 5), this moment is the time when it crosses the plane, and thus the time-to-contact T to the plane can be detected as T_C_-T_0_. In addition, the 3D orientation n of the plane can be detected as the intersection of the red great circles, each of which is obtained by the polar transformation of a pink point on the degenerated great-circle.

However, there are still issues to be solved. In Figure 5, the moment T_C_ in which the plane in space is degenerated into a great circle on the eyeball is not known because it is a future time. All we know is the position on the eyeball at a current time T_0_ (shown as a blue triangle) and the movement τ _2D_ from time T_0_ to time T_1_ (shown as a green arrow and called local motion). The cell model that detects this local motion was reported (Kawakami & Okamoto, 1996a, 1995a; Kawakami, 1996b): this will be described in Section **2.1.2**.

Therefore, based on these two known parameters on the eyeball (i.e. the position P_0_ at a current time T_0_ and the local motion τ _2D_ from P_0_ to P_1_ in Figure 8(A)), let us predict the position P_T_ at the future arbitrary time T: P_1_ is the position at the next time T_1_. This prediction is explained below based on previous reports (Kawakami et al., 2003, 2000).

**Figure 8.**
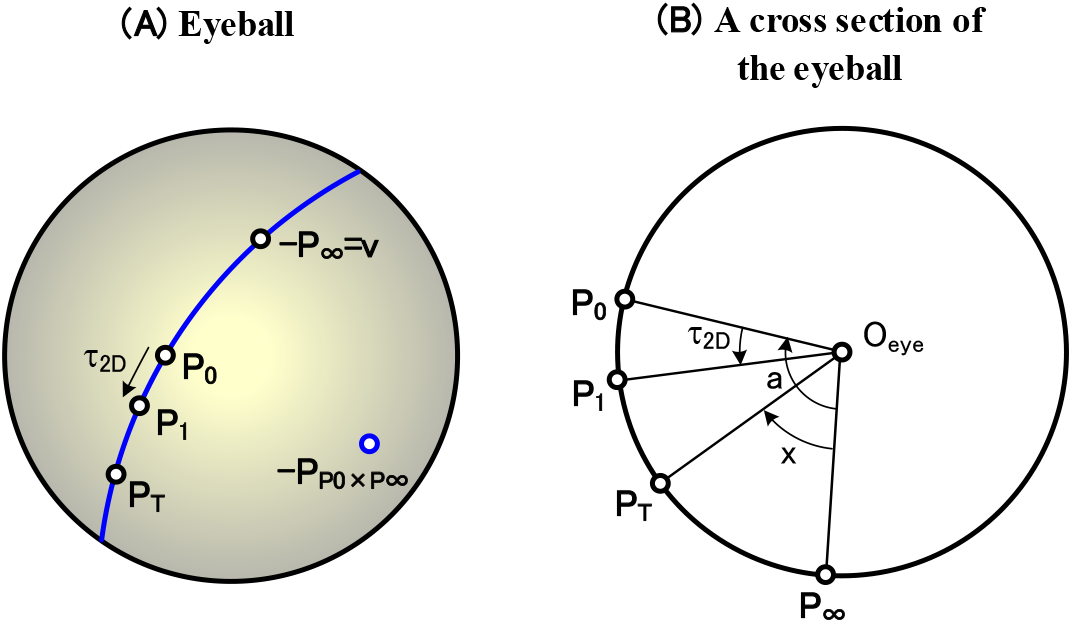
Various parameters on the eyeball and the cross-ratio transform. (A) Various parameters on the eyeball are shown. A position P_0_ of a current time T_0_ moves as follows. That is, P_0_ moves to P_1_ after time t_d_, and reaches position P_∞_ opposite to the movement direction v after infinite time: the movement from P_0_ to P_1_ is the local motion τ_2D_, and P_p0×p∞_ is the pole of the blue movement trajectory above. Since P_0_ and τ_2D_ are known, the position P_T_ at any time T can be predicted using the cross-ratio transform of equation(1.0-4). (B) A cross section of the eyeball passing both the blue movement trajectory ((A)) and the eyeball center O_eye_ is shown. Using these central angles (τ_2D_, a, and x), the cross-ratio transform of equation(1.0-4) is expressed analytically as described in equation(1.0-6).

Before this explanation, note the following. For ease of description, it is assumed below that, in principle, the opponent (or object) moves, not the self (or eyeball). Since this movement change is a relative movement, there is no problem. To make it our own move, just invert the sign of the move direction.

In Figure 8(A), assuming that the opponent’s movement direction v is known, the position P_0_ reaches a position P_∞_ opposite to v after infinite time. Thus, the position P_∞_ is expressed by equation(1.0-1). Note that since the position P _∞_ is not visible on the back side of the eyeball, the inversion -P_∞_ is shown.

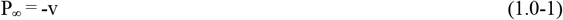

A normalized time-to-contact t and unit time t_d_ are defined by the following equation.

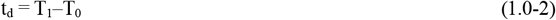

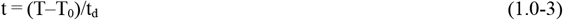

As proved in Appendix **1**, the following relationship holds between these four positions (P_0_, P_1_, P_T_, and P_∞_).

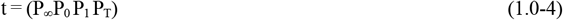

where (P_∞_P_0_P_1_P_T_) is a cross ratio (also called the double ratio or anharmonic ratio) defined in projective geometry (Izumi et al., 1987; Iwata, 1981). Based on the definition, describing this cross ratio using directed segments (i.e. P_∞_P_1_, P_0_P_1_, and P_∞_P_T_ etc.) on a spherical surface yields the following equation.

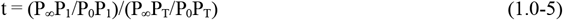

In order to express this equation analytically, Figure 8(B) is used as follows: this figure shows a cross section of the eyeball passing both the blue movement trajectory in Figure 8(A) and the eyeball center O_eye_. The directed segments P_∞_P_1_, P_0_P_1_, P_∞_P_T_, and P_0_P_T_ on the eyball correspond to the central angles (a+τ_2D_), τ_2D_, x, and (x-a) in Figure 8(B), respectively. Thus, rewriting equation(1.0-5) using these angles yields the following equation: see Appendix **2** for the proof of equation (1.0-6) and why it is expressed in the sin function.

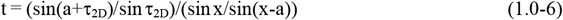

Here, these angles (i.e. a, x, and τ_2D_) and 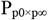 used are calculated using the positions (i.e. P_0_, P_1_, P_T_, and P_∞_) by the following equation (see Appendix **4**).

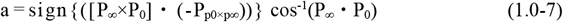

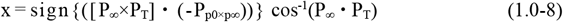

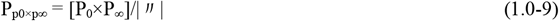

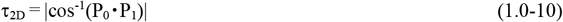

[P_∞_×P_0_] etc. are vector products, and (P_∞_•P_0_) etc. are scalar products. The 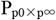 is the pole of the blue movement trajectory (i.e. the great circle) in Figure 8(A): this pole corresponds to the north pole when the trajectory is the equator (Gurevic, 1962). The sign{x} is a function that determines the sign, and if x is positive, it is 1, if it is negative, it is -1. The|″| is the absolute value of [P_∞_×P_0_], and is used to make 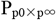 the unit vector.

Solving equation(1.0-6) for x yields the following equation.

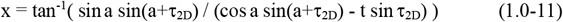

Since the current time position a and the local motion τ_2D_ are known as described above, substituting them into equation (1.0-11) enables a position x at arbitrary time t (i.e. the P_T_ of Figure 8(B)) to be predicted. Since this method using the cross ratio enables the position P_T_ of the arbitrary time T to be predicted, we have named equation (1.0-11) as the cross-ratio transform, including the underlying equations(1.0-4 and 6) (Kawakami et al., 2003, 2000). If two of the three parameters (t, P_1_, P_T_) in equation(1.0-4) are known, this transformation allows us to calculate one of the remaining. In equation(1.0-11), P_T_ was calculated by knowing t and P_1_ (i.e. knowing t and τ_2D_).

In this way, using the two known parameters (i.e. the position P_0_ on the eyeball at the current time T_0_ and the local motion τ_2D_ from P_0_ to P_1_), we have been able to predict the position P_T_ of a future arbitrary time T. Therefore, this prediction has enabled the aforementioned problem to be solved. Based on this solution, the moment T_C_ when the plane crosses the eyeball and its 3D orientation n can be determined by the following steps.

[Step 1] Give any two parameters (i.e. the position P_0_ and local motion τ_2D_). Then, assuming that the movement direction v is known, calculate P_∞_ by equation(1.0-1).

[Step 2] Set a time T arbitrarily and calculate the normalized time-to-contact t by equation(1.0-3).

[Step 3] Predict a position P_T_ on the eyeball at the time T (Figure 8(A)) using the cross-ratio transform of equation(1.0-11): the variable a used in this transform is calculated by equation(1.0-7).

[Step 4] Plot this P_T_ on the eyeball (shown as a pink point in Figure 5), and then draw a red great circle that is obtained by the polar transformation of the P_T_. There is an important property that the 3D orientation n of a plane to be determined lies on this great circle.

[Step 5] Draw such great circle for all positions P_0_ at every time including the T_0_ and T_1_. These great circles do not intersect at a general time T, but intersect at a red point n only at a certain time T_C_ in Figure 5. This time intersected is the moment when a plane crosses the eyeball. Thus, this moment is determined as the time T_C_, and the time-to-contact to the plane is detected as T_C_–T_0_. In addition, the 3D orientation n of the plane is determined as an orientation from the eyeball center O_eye_ to this intersection.

In this way, knowing the P_0_ and τ_2D_ enables the moment T_C_, at which the plane crosses the eyeball, and the 3D orientation n of the plane to be determined using the cross-ratio transform: the time-to-contact T is obtained as T_C_-T_0_.

Section **2.2** will show that these steps are accurately expressed by mathematical equations, and Section **2.3** will show that the T_C_ and n of the plane are correctly determined using computer simulations: the validity of this concept proposed in Figure 5 will be shown in Section **2.3.1**. In addition, Section **3** will show a method for determining the remaining parameter (i.e. the shortest distance D) of Figure 1. Furthermore, Supplementary materials will show the computational procedure for simulating the neural network connecting cells that are modeled in Sections **2** and **3**.

Because these steps are represented by complex mathematical equations (such as equation(1.0-11)), you may think that it is difficult for cells in the cerebrum to perform them. However, it will be indicated in Section **2.2**(12) that this is not a problem. The main points are as follows. These equations represent the neural networks after learning is complete. Therefore, they are not necessary for the cell operations that are performed after learning, and thus there is no problem.

The above algorithm for recognizing the plane parameters has been realized by the following four types of my attention that continue temporally. In 1977, I started to study this plane recognition. My first attention was on the moment when the plane crosses (Figure 3), and based on that, I got the idea of Figure 5 in 1982. As the second attention, in 1990, I discovered a cross-ratio transform that is performed in equation(1.0-4): this transform predicts the position P_T_ at an arbitrary time T based on two known parameters (i.e. the position P_0_ at a current time and a local motion τ_2D_). As the third attention, in 1992, I modeled a series of cells for detecting the local motion τ_2D_, and we later reported the modeled series (Kawakami & Okamoto, 1996a, 1995a). As the final attention, in 1997, I realized the idea of Figure 5 based on these attentions, and modeled a series of cells (shown in Figure 9 later) using mathematical equations to detect the time-to-contact T_C_ to the plane and its 3D orientation n: we later reported the modeled series (Kawakami et al., 2003, 2000). At the same time, in 1997, I also modeled a series of cells (shown in Figure 17 later) using mathematical equations to detect the shortest distance D to the plane and its 3D orientation n: we later reported the modeled series (Kawakami et al., 2003, 2000).

**Figure 9.**
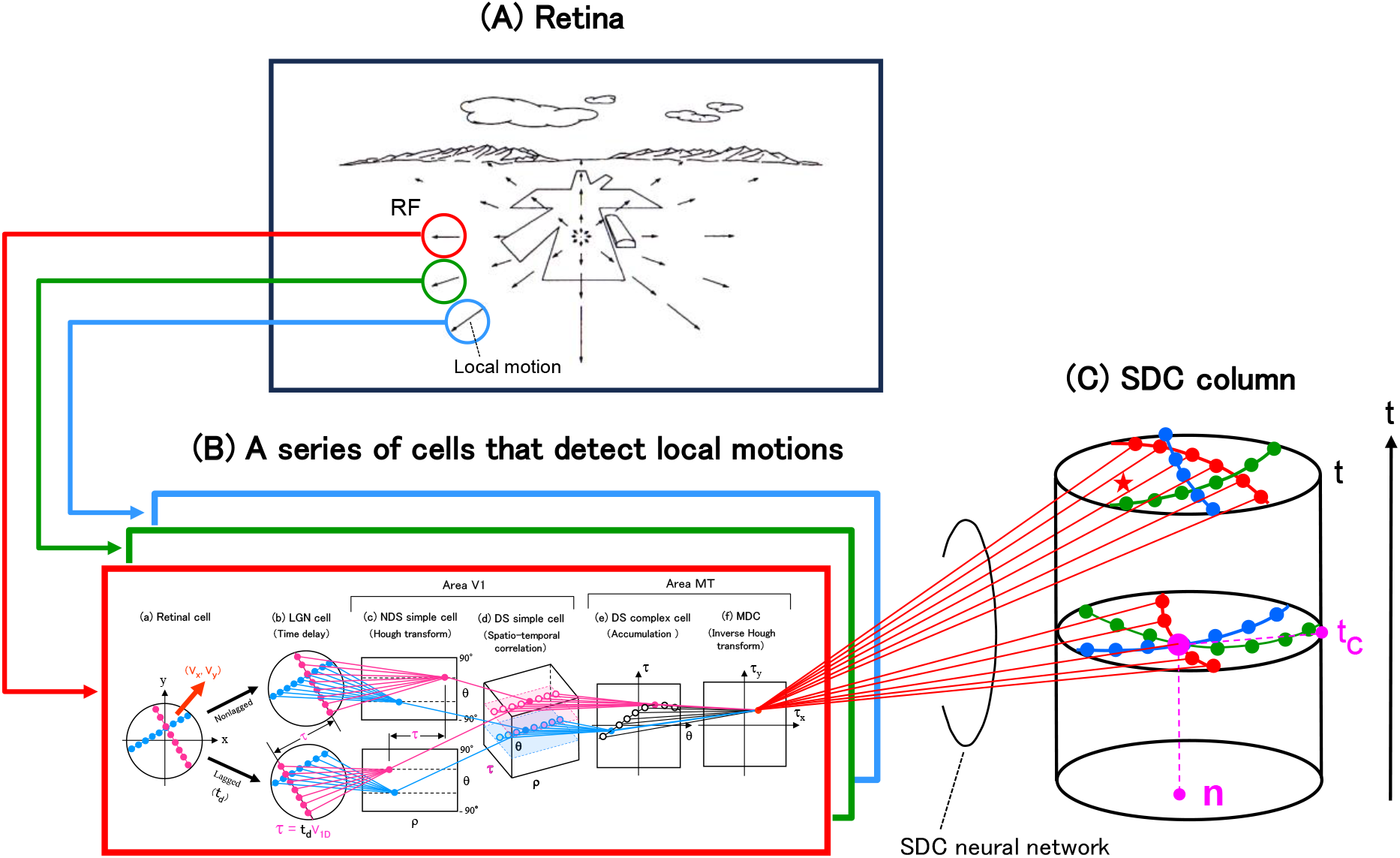
A cell model for detecting the time-to-contact to a plane and its 3D orientation. A series of modeled cells that detect two parameters in Figure 1 (i.e. the normalized time-to-contact t to a plane and its 3D orientation n) is shown. (A) An OFP in the retina consists of local motions, each of which occurs in a small region (i.e. a RF). (B) Each local motion is detected by a series of cells (i.e. LGN cells, NDS simple cells, DS simple cells, DS complex cells, and MDCs). In this series, a series of transforms (i.e. the time delay, Hough transform, spatio-temporal correlation, accumulation, and inverse Hough transform) are performed, respectively: in LGN cells, the 2D convolution with a DOG filter is also performed at the same time to emphasize the contours of figures. The pink and blue straight lines represent the neural networks connecting each cell array, and perform the above transforms. (C) A column composed of SDCs is shown. Each SDC detects the normalized time-to-contact t to a plane as the height coordinate, and detects the 3D orientation n as the polar coordinate of the cross section at this height. The red straight lines represent the neural network connecting each MDC in the (B)(f) and the SDC column in the (C), and perform a combination of the cross-ratio and polar transforms.

In fact, the origin of these four types of attention mentioned above stems from developing a stereoscopic vision function for robots. In other words, the origin of this work was to develop this function for robots that was scheduled to be used in nuclear power plants: this development was done by focusing psychologically on a motion stereo that birds perform during flight (Kawakami, 1987; Morita et al., 1989).

## 2. A cell model for detecting the time-to-contact to a plane and its 3D orientation

### 2.1. Cell model

Based on the five steps above, a series of cells in Figure 9 has been modeled, and are explained as follows. This figure consists of the (A), (B), and (C). In the retina of the (A), an OFP is generated by the movement of an opponent (or an object), and consists of local motions each of which occurs in a small region (i.e. a receptive field (RF)). These local motions are detected by a series of cells in the (B). The (C) is a cylindrical cell array named an SDC column, where SDC is an abbreviation for planar-surface-detection cell. Each SDC in the column integrates every local motion (τ_X_,τ_Y_) detected in the (B)(f) to extract the OFP in the (A), and then detects the time-to-contact to a plane and its 3D orientation. This series of modeled cells are described below.

#### 2.1.1. Retina

Although the OFP occurs on the eyeball, the retina of the (A) is drawn on a plane for easy understanding. Each local motion (i.e. each velocity vector) that constitutes this OFP is generated within a RF by the motion of the the opponent (or object).

In the example of Figure 2, the magnitude of this local motion increases as the airplane approaches the runway (i.e. the plane), and thus its magnitude corresponds to the time-to-contact to the plane. Also, when the plane tilts (i.e. when the airplane’s tilt relative to the plane changes), the shape of this OFP changes, and thus the shape corresponds to the 3D orientation of the plane. These time and orientation is detected by the corresponding SDC: this detection will be described in Section **2.1.4**.

#### 2.1.2. A series of cells that detect local motions

A local motion within each RF in the (A) is detected by a block (i.e. a rectangle) in the (B) that corresponds to this RF. That is, a local motion in the red RF is detected within the red rectangle, that in the green RF is detected within the green rectangle, and that in the blue RF is detected within the blue rectangle. A series of modeled cells for detecting this local motion within each rectangle is the same, and has been reported so far (Kawakami & Okamoto, 1996a, 1995a, 1992d; Kawakami, 1996b; Kawakami et al., 1992b).

The responses of these modeled cells to various light stimuli were consistent with actual physiological experiments (Okamoto et al., 1999; Kawakami & Okamoto, 1996a; Kawakami, 1996b). Also, the NDS simple and DS simple cells exist in the primary visual cortex area (i.e. in area V1), and the DS complex cells and MDCs exist in both area V1 and the middle temporal area (MT) (Kawakami and Okamoto, 1996a; Kawakami, 1996b; Okamoto et al., 1999).

About 1,000 cells are arranged in a circle or rectangle of the (B). About 20,000 cells are arranged in the cuboid of the (B)(d). In this series of cells (i.e. lateral geniculate nucleus (LGN) cells, nondirectionally selective (NDS) simple cells, directionally selective (DS) simple cells, DS complex cells, and motion detection cells (MDCs)), a series of transforms (i.e. the time delay, Hough transform, spatio-temporal correlation, accumulation, and inverse Hough transform) is performed, respectively. The neural network connecting this series of cell types is shown as pink or blue straight lines.

An outline of the local-motion detection, which is performed in the red rectangle of the (B), will be described in the subsections (1) to (6) below, respectively (see Kawakami & Okamoto (1996a) and Kawakami (1996b) for details): an enlarged view of the (B) is shown in Figure S2-1 of Supplementary materials.

1. Retinal cells in the (a) The (a) shows the red RF of the (A), in which a cross figure moves in a local motion (i.e. a velocity vector) of (V_X_,V_Y_). Each dot indicates a retinal cell that is firing. This local motion is detected by the following series of cell types.
2. LGN cells in the (b) that perform the time delay This retinal cell array in the (a) branches into two LGN cell arrays in the (b): that is, one is an array without time delay (i.e. nonlagged) in the upper part, and the other is an array with time delay (i.e. lagged) in the lower part. This time delay t_d_ enables the local motion related to both time and space to be transformed into a spatial-only parameter τ (shown in pink) that cell arrays are good at: physiologically, this t_d_ was reported to be about 50 ms (Mastronarde, 1987a, 1987b; Kawakami & Okamoto, 1996a; Kawakami, 1996b). The above is described in detail as follows. The velocity of each straight line that constitutes the cross figure in the (a) is called the one-dimensional (1D) velocity V_1D_. The pink straight line in the (a) is converted to two pink straight lines in the upper and lower LGN arrays of the (b), and the distance between them is τ shown in pink. That is, the 1D velocity V_1D_ is converted to the spatial interval τ (= t_d_ V_1D_) by the above time delay t_d_. Thus, the role of this time delay is to convert the velocity V_1D_ into the spatial interval τ. Note that in these LGN cells, two-dimensional (2D) convolution by the DOG filter is performed at the same time to emphasize figure contours (Kawakami & Okamoto, 1996a; Kawakami, 1996b): this convolution is done by equation(2) in Kawakami & Okamoto (1996a).
3. NDS simple cells in the (c) that perform the Hough transform The Hough transform, which is performed by each NDS simple cells in the (c), enables the 2D interval τ in the (b) to be detected one-dimensionally, as follows. The Hough transform detects a pink straight line in each array in the (b), and the corresponding pink cell in the (c) fires: this cell detects the orientation θ and position ρ of each line as its coordinate (ρ,θ) (see Figure 33(D) later; Kawakami & Okamoto, 1996a). Since these two pink lines in the (b) are parallel, two pink cells with the same θ coordinate in the (c) fire. In this way, the role of the Hough transform is to detect the pink 2D interval τ in the (b) as a pink 1D interval τ in the (c) (i.e. as a difference τ between two ρ coordinates). Note that this Hough transform is done by equation(4) in Kawakami & Okamoto (1996a), and was explained geometrically in Figure A1(i) in Kawakami & Okamoto (1996a) (see also Section **4.5.1** later). In addition, note that this Hough transform will be a special case of the polar transform in Figure 6, as shown in Section **4.5.2** later.
4. DS simple cells in the (d) that perform the spatio-temporal correlation The detection of the above difference τ between two ρ coordinates is described as follows. Performing a spatio-temporal correlation between these two (ρ,θ) arrays in the (c) causes a new coordinate τ to be added, and thus the 2D array (ρ,θ) in the (c) is converted to a 3D array (ρ,θ, τ) in the (d): this conversion is called a spatio-temporal correlation because it is spatially correlated between the lagged and nonlagged (ρ,θ) arrays. In the (ρ,θ, τ) array of the (d), a pink cell fires, and thus the difference τ of the ρ coordinates in the (c) is detected as the τ coordinate in the (d): in detail, this difference τ between the pink (or blue) NDS simple cells with the same θ coordinate in the (c) is detected as the pink (or blue) τ coordinate in the (d). Thus, the 1D velocity V_1D_ of the pink straight line in the (a) has been detected as the pink coordinate τ of this 3D array (ρ,θ,τ). Note that this spatio-temporal correlation is done by equation(5) in Kawakami & Okamoto (1996a). Here, the following caveats are pointed out regarding this correlation. In the correlation above, multiplication was used (see equation(5) in Kawakami & Okamoto (1996a); Kawakami & Okamoto, 1995a; Kawakami, 1996b). However, there are two physiological problems with this multiplication. The first is that this operation easily overflows beyond the cell’s dynamic range, whose upper limit seems to be about 100 (page 92 in Kawakami (1996b)): for example, 20×30(= 600) greatly exceeds the upper limit of 100. The other is that a physiological mechanism that performs the sign operation of multiplication (e.g. negative x positive is negative) is unknown. To address these problems, a multiplication-like operation that is neurophysiologically feasible (i.e. feasible by combining the presynaptic inhibition and the postsynaptic excitation and inhibition) was proposed (Kawakami, 1996b; Kawakami & Okamoto, 1995b). In this operation, absolute values are added in the same sign operation as multiplication, so the above problems do not occur. That is, for the first problem, 20+30(= 50) does not exceed the upper limit 100 described above. For another problem, the sign operation that is the same as multiplication are performed by a neurophysiologically viable synaptic-circuit. Thus, this multiplication-like operation can solve the above physiological problems of multiplication using this synaptic circuit: the synaptic circuit for the multiplication-like operation has been described in Kawakami (1996b) and Kawakami & Okamoto (1995b). A computer simulation confirmed that the spatio-temporal correlation in the (d), into which this multiplication-like operation was applied, was performed correctly (page 101 in Kawakami (1996b); Kawakami & Okamoto, 1995b). For this reason, the multiplication-like operation should be used for the correlation in the (d), but multiplication, which is functionally almost equivalent to the multiplication-like operation, has been used here. This is because a description of the synaptic circuit for this operation takes a considerable number of pages.
5. DS complex cells in the (e) that perform the accumulation Each DS complex cell (θ, τ) in the (e) accumulates all DS simple cells ((d)) that have the same θ and τ: this accumulation is done by equation(12) in Kawakami & Okamoto (1996a). Thus, the ρ coordinates are erased, and the array (ρ,θ,τ) of the (d) is converted to the array (θ,τ) of the (e). Since the position coordinate ρ of each straight line has been erased by this accumulation, a DS complex cell can detect the orientation and 1D velocity of the line as the cell’s coordinate (θ,τ) no matter where the line is within the RF. Therefore, the orientation θ and 1D velocity τ of each straight line, that constitutes the cross figure of the (a), can be detected by a DS complex cell wherever the figure is in the RF. So far, the cell types (i.e. LGN cells, NDS simple cells, DS simple cells, and DS complex cells) relevant to detecting 1D velocities of straight lines has been described. Here, regarding NDS simple cells in the (c), DS simple cells in the (d), and DS complex cells in the (e), note the following. It was discovered that even if the RF size changed about 30 times between the fovea and periphery of the visual field, the orientation resolution Δθ of the simple and complex cells remained almost constant at about 10 degrees (Hubel & Wiesel, 1984; Hubel et al, 1978). This discovery (i.e. the invariance of Δθ) will be explained in Section **S1.2.6.1**(6) of Supplementary materials.
6. MDCs in the (f) that perform the inverse Hough transform Finally, each MDC of the (f) detects a local motion (V_X_,V_Y_) of the cross figure in the (a) as follows, where this motion is also called the 2D velocity. Pink and blue cells in the (e) fire on a sinusoidal pattern. The amplitude of this pattern is the magnitude τ_2D_ of the local motion (Figure 8(A)) and its phase is the direction of the motion: see Kawakami & Okamoto (1996a) and Kawakami (1996b) for why the sinusoidal pattern is generated. A MDC in the (f) extracts this sinusoidal pattern by the inverse Hough transform to detect the local motion. That is, the red MDC at the position (τ_X_, τ_Y_) fires and detects the local motion (V_X_,V_Y_) by the following equation.

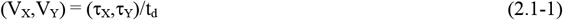 This local motion (τ_X_, τ_Y_) is detected wherever the cross figure in the (a)is in the RF: this is because, as described in subsection (5) above, a DS complex cell can detect each line, which makes up this figure, wherever the line is in the RF. Note that this inverse Hough transform is done by equation(14) in Kawakami & Okamoto (1996a), and is explained geometrically in Figure A1(ii) in Kawakami & Okamoto (1996a) (see also Section **4.5.1** later). In addition, note that this inverse Hough transform will be a special case of the inverse polar transform, as shown in Sections **4.5.2** and **4.5.3** later. Summarizing the above, the local motion τ_2D_ is detected by the series of cells (i.e. the series of transforms) of the (B). Each connection of the neural network connecting these cell types in the (B) is accurately expressed one by one using these transforms (Kawakami & Okamoto, 1996a, 1995a): the pink and blue straight lines in the (B) represent this network. The computational procedure for this neural network is shown in Sections **S2.1, S2.2**, and **S3.1** of Supplementary materials.
7. Things to note Although this local-motion detection in the (6) might be thought to be limited to the cross figure, complex shapes containing curves and single dots are more robustly detected than the cross figure. The reason is as follows. Since a more continuous sine wave, in the DS complex cell array of the (e), is generated for these complex shapes than for the cross figure, the sine waves for these shapes are extracted robustly by the inverse Hough transform and thus their local motions τ_2D_ are robustly detected by the MDC. The reason why this continuous sine wave is generated for these curve and single dot is described as follows. First, since the single dot is equivalent to all straight lines that pass through it, a continuous sine wave that is obtained by detecting the orientations θ and 1D velocities τ of these lines is generated (see Figures 33(A and C); Figure 6(B) in Kawakami & Okamoto (1996a)). Next, since the curve is equivalent to all straight lines that touch it, a continuous sine wave that is obtained by detecting the orientations θ and 1D velocities τ of these lines is generated (see Figure 7 in Kawakami & Okamoto (1996a)). Conversely, since the cross figure in the (a) has only two straight lines, two points on the sine wave is generated rather than a continuous sine wave (see Figure 6(A) in Kawakami & Okamoto (1996a)). Thus, this figure is the least suitable shape for the sinusoidal extraction.

This local-motion detection is very robust to noise. An example is shown in Figure 7(C) of Kawakami & Okamoto (1996a). A sine wave appears in the DS complex cell array of Figure 7(C)(iv) even when random-dot noises are superimposed on the ring within the RF of the retina. This wave is extracted by the inverse Hough transform, and thus a MDC of Figure 7(C)(v) fires in isolation. This robustness that is not affected by noise is due to the DOG filter of LGN cells described in the (2) above. The reason is shown as follows. Since this filter produces blue negative-responses (or inhibitions) in the LGN cell array in Figure 7(C)(i), these inhibitions cancel the noises and thus a sinusoidal pattern appears in the complex cell array of Figure 7(C)(iv). In this way, the DOG filter of LGN cells in the (2) above plays an important role in the noise suppression in the complex cell array.

The above series of modeled cells in Figure 9(B) is represented by the complex equations (such as the Hough and inverse Hough transforms), and you may think that it is difficult for cerebral cells to execute them. However, that worry is not a problem for the same reasons as in Section **2.2**(12) later.

Since I have shown that the cells in Figure 9(B) are arranged in 2D (or 3D) arrays, you may think that it is impossible for the cerebrum to form such an orderly array. However, that worry is not a problem for the same reasons as in Section **2.2**(12) later. Note the following. Even if the arrays of NDS simple cells of (c), DS complex cells of (e), and MDCs of (f) in Figure 9(B) are highly distorted, a method has been proposed in Section **6.2.3** of Kawakami & Okamoto (1996a) that can physiologically validate the neural network of each array using the optical imaging.

I point out that a VLSI implementation research (Akima et al., 2017) was reported to realize the series of cells from LGN cells to MDCs (Figure 9(B)) using electric circuit and to detect a local motion in each RF.

#### 2.1.3. SDC column for detecting planes

As described in Section **2.1.2**, a MDC (τ_X_, τ_Y_) of Figure 9(B)(f) has detected the local motion (V_X_,V_Y_) of the (B)(a). A SDC neural network connecting this MDC and the SDC column of the (C) is described as follows.

The cross section of this SDC column at each height t corresponds to a disk to which the eyeball of Figure 5 is converted. Stacking this disk in the height (i.e. in the normalized time-to-contact) direction yields the SDC column. This conversion of the eyeball to the disk is carried out as follows (Figure 10): the conversion is the same as that for a cell model that detects curvatures constituting a figure (see Figure 4(C) in Kawakami et al. (2020)).

**Figure 10.**
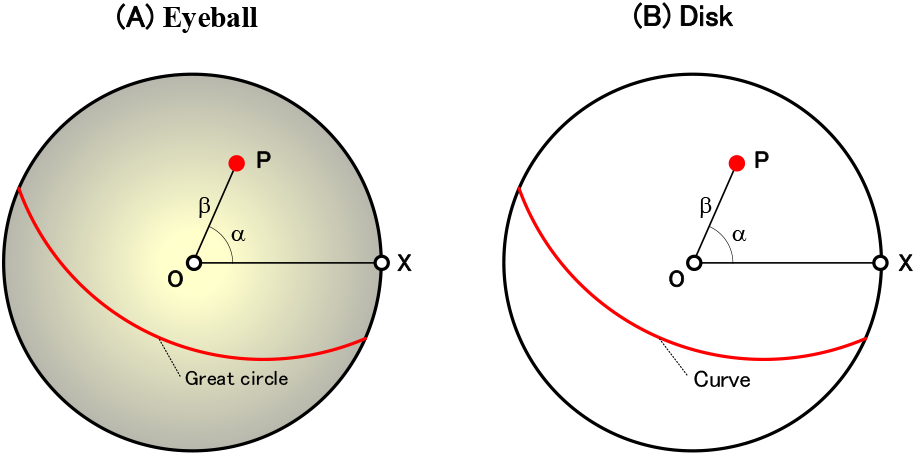
Equidistant projection from the eyeball to a disk. The equidistant projection transforms the eyeball into a disk, as follows. (A) First, express an arbitrary point P on the eyeball as (α,β) in a polar coordinate. (B) Next, express a point P on the disk in the polar coordinate (α,β) that has the same value as on the eyeball. This is the equidistant projection from the eyeball to the disk. The backward projection (i.e. the projection from the disk to the eyeball) is performed in a similar manner. A great circle on the eyeball ((A)) is converted into a curve on the disk. Here, O is the center of the visual field and X is the basic vector (1,0,0).

Express any point P on the eyeball in Figure 10(A) as (α,β) in the polar coordinate: O is the center of the visual field and X is the basic vector (1,0,0); assuming the O as the north pole of the earth gives the α and β to correspond to the longitude and the 90 deg - latitude, respectively. Also, express a point P on the disk in the polar coordinate that has the same values (α,β) as on the eyeball (Figure 10(B)). This is a conversion from the eyeball of the (A) to the disk of the (B). This conversion is called the equidistant projection, which is a type of projection method for fisheye lenses: Figure 7 showed a photograph taken with this fisheye lens.

Here, if the surface of the eyeball in Figure 10(A) is made of rubber, let us consider how the surface is stretched and converted into the disk of Figure 10(B). The larger the value of β, the greater the elongation in the circumferential direction, and finally the maximum elongation occurs at β=π/2. Thus, the surface of the eyeball is converted to the disk of Figure 10(B) by extending π/2 times in the circumferential direction. It is such a conversion.

Stacking this disk of Figure 10(B) in the height direction t yields a cylinder (α,β,t). Since the 3D orientation n of a plane is expressed as (α_n_,β_n_) in the polar coordinate, this cylinder is expressed as (n,t) and represents the SDC column of Figure 9(C). About 10,000 SDCs are arranged in this cylinder (n,t). Each SDC detects a normalized time-to-contact t as its height coordinate, and detects a 3D orientation n as its polar coordinate of the cross section.

A red great-circle on the eyeball in Figure 10(A) is converted into a red curve on the disk in Figure 10(B): the curve is obtained by converting this circle using the equidistant projection. These great circle and curve can be expressed by mathematical equations. That is, both the great circle and curve are expressed by the same equation(A3-1) in Appendix **3**. Therefore, this curve on the disk will be called a great circle in the same way as that on the eyeball.

#### 2.1.4. Neural network connecting MDCs and the SDC column

Based on the five steps described in Section **1**, let us determine a neural network connecting each MDC of Figure 9(B)(f) and the SDC column of Figure 9(C).

[Step 1] Give two parameters of Figure 8(A) (i.e. the position P_0_ on the eyeball at a current time and a local motion τ_2D_ from P_0_ to P_1_), as follows. First, give the position P_0_ as the RF center O_IJ_ based on equation(A4-3) in Appendix **4**. Next, give the local motion τ_2D_ by the following equation, where (τ_X_,τ_Y_) is the coordinate of the MDC in Figure 9(B)(f).

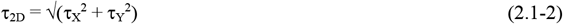

Here, how a local motion τ_2D_ on the eyeball is detected by this MDC array (τ_X_, τ_Y_) is described in Section **A4.1** of Appendix **4**. Further, assuming that the movement direction v is known, calculate P_∞_ by equation(1.0-1).

[Step 2] Set a time T arbitrarily and calculate the normalized time-to-contac t by equation(1.0-3).

[Step 3] Predict a position P_T_ on the eyeball at the time T (i.e. at the normalized time-to-contact t) using the cross-ratio transform of equation(1.0-11): the variable a used in this transform is calculated by equation(1.0-7).

[Step 4] Plot this P_T_ on the cross section at a height t in Figure 9(C): this plotted P_T_ is shown as a red mark ★. Then, draw a red great circle, on the cross section, which is obtained by the polar transformation of the P_T_. There is an important property that the 3D orientation n of a plane to be determined exists on this great circle.

Finally, connect all SDCs (indicated as red dots) on this great circle to the MDC (τ_X_, τ_Y_) of Figure 9(B)(f). This connection shown as red straight lines represents a SDC neural network in Figure 9(C). The above is for the red RF in Figure 9(A), and for the green and blue RFs, the green and blue great circles and all SDCs on each great circle are drawn: the neural networks are omitted for ease of viewing. These great circles do not intersect at a single point.

[Step 5] When such a great-circle drawing is performed for all heights t, the great circles intersect at one point only at some height t_C_ (see Figure 9(C) for t_C_). The normalized time-to-contact t_C_ to the plane is detected as the height coordinate of the pink intersected SDC, and its 3D orientation n is detected as the polar coordinate of this SDC.

In this way, a neural network connecting each MDC of Figure 9(B)(f) and the SDC column of Figure 9(C) has been modeled. This network detects the normalized time-to-contact t to the plane and its 3D orientation n. Although the essential points of the network have been explained with an emphasis on comprehensibility, Section **2.2** will show that the network is expressed accurately using mathematical equations. Section **2.3** will also show that the validity of this network is confirmed by computer simulations.

### 2.2. Representation of the SDC neural network by mathematical equations

Let us express the neural network, described in Section **2.1.4**, by mathematical equations using the form of a flowchart, as follows (see Figure 11). Here, you need to note that Appendix **4** shows important properties such as how to detect each local motion τ_2D_ on the eyeball with a MDC array (τ_X_, τ_Y_) of Figure 9(B)(f) and the relationship of various parameters.

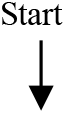

**Figure 11.**
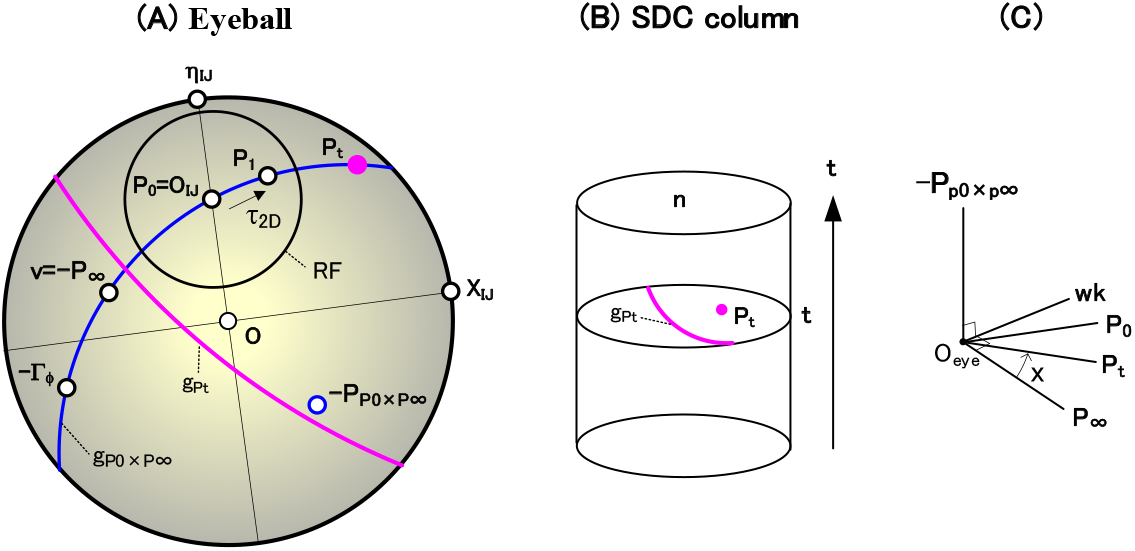
Detection of the normalized time-to-contact t and 3D orientation n using the cross-ratio and polar transforms. (A) The parameters on the eyeball are shown. The position P_0_ at a current time T_0_ moves as follows. That is, P_0_ moves to P_1_ after time t_d_ and reaches position P_∞_ in the opposite direction of movement v after infinite time: the movement from P_0_ to P_1_ is a local motion τ_2D_, and P_p0×p∞_ is the pole of the blue movement trajectory. Since P_0_ and τ_2D_ are known, the pink position P_t_ at any time T can be predicted using the cross-ratio transform of equations(1.0-11 and 2.2-8). The polar transformation of this position P_t_ yields a pink great circle g_Pt_. There is an important property that the 3D orientation n of a plane to be determined lies on this great circle. Here, X_IJ_ and η_IJ_ belong to each RF coordinate system (see equations(A4-4 and 10) in Appendix **4**) and P_0_=O_IJ_ (see equation(A4-3) in Appendix **4**). Γ_ϕ_ is the direction of movement from P_0_ to P_1_ (see equation(A4-7) in Appendix **4**). (B) The SDC column of Figure 9(C) is shown. The equidistant projection in Figure 10 converts each position P_t_ and great circle g_Pt_ in the (A) into a pink dot P_t_ and curve g_Pt_ in the (B), respectively. When such curve g_Pt_ is drawn for all positions P_t_ and all heights, these curves intersect only at a certain height t_C_ (see Figure 9(C) for t_C_). The normalized time-to-contact t_C_ to the plane is detected as this height coordinate, and its 3D orientation n is detected as the polar coordinate of this intersection. The SDC neural network that performs this detection is expressed by mathematical equations in Section **2.2**. (C) The variables x and each vector required to calculate equation(2.2-8) are shown.

1. Give a movement direction v (see Figure 11(A)) With respect to the center of the visual field O, if the movement direction v is (O•v)>0, it is an approaching movement, if (O•v)<0, it is a separating movement, and if (O •v)=0, it is a horizontal movement. Calculate P_∞_ using the following equation (see equation(1.0-1)).

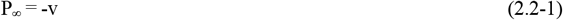
2. Sweep a RF center O_IJ_ 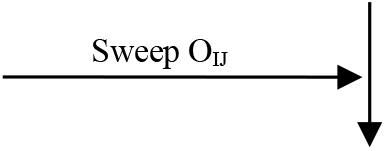 Calculate P_0_ using the following equation (see equation(A4-3) in Appendix **4**).

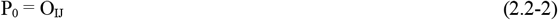
3. Sweep a normalized time-to-contact t 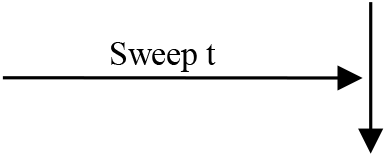
4. Sweep a MDC array (τ_X_,τ_Y_) 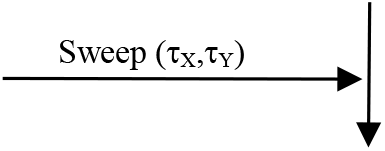 Convert this (τ_X_,τ_Y_) to its polar coordinate (τ_2D_,ϕ) by the following equation (see Figure A4(D) in Appendix **4**).

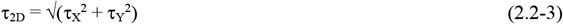

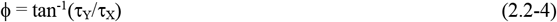
5. Calculate x(=cos^-1^(P_∞_ • P_t_)) with the following equation (see equation(1.0-11))

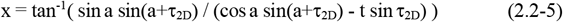 Here, the variable a used above is calculated with the following equations (see equations(1.0-7 and 9)).

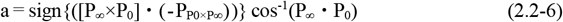

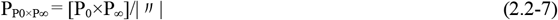
6. Calculate the position P_t_ at the time t using the following equation (See Figure 11(C))

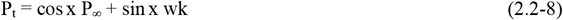 Here, the unit vector wk used above is calculated by the following equation (see Figure 11(C)).

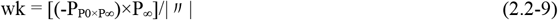
7. Calculate a pink great circle g_Pt_ whose pole is the position P_t_ (see Figure 11(A)) First, P_t_ expressed by equation(2.2-8) is represented in an orthogonal coordinate system by the following equation.

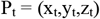 Convert the P_t_ to polar coordinate (α_t_,β_t_) by the following equation.

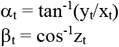 Next, when an arbitrary point on this circle g_Pt_ is represented as a polar coordinate (α,β), there is a following relationship (see equation(A3-1) in Appendix **3**).

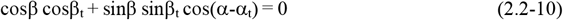 This is the mathematical expression of the great circle g_Pt_, which is obtained by the polar transformation of the position P_t_. There is an important property that the 3D orientation n of a plane to be determined lies on this great circle.
8. Draw this great circle g_Pt_ in the SDC column Using the equidistant projection of Figure 10, convert this great circle g_Pt_ on the eyeball into a pink great-circle on the cross section at the height t of the SDC column in Figure 11(B). Then, draw this converted great circle on the cross section. Note that the great circle on the cross section is represented by the same equation(2.2-10) as that on the eyeball (see Section **A3.2** in Appendix **3**).
9. Connect all SDCs on this great circle with the MDC (τ_X_,τ_Y_) Connect all SDCs (indicated as red dots in Figure 9(C)) on this great circle at height t to the MDC (τ_X_,τ_Y_) of Figure 9(B)(f). This connection shown as red straight lines represents an SDC neural network in Figure 9(C). 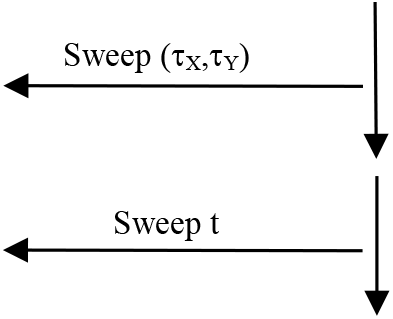
10. Up to this point, the red neural network in Figure 9(C) have been created. That is, the neural network for the red RF in Figure 9(A) has been completed. 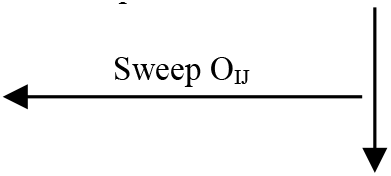
11. A neural network for all RFs in Figure 9(A), including red, green, and blue, has been completed. Each connection of the neural network (Figure 9(C)) connecting the MDCs and the SDC column has been accurately expressed one by one using mathematical equations. 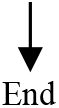 In this way, the SDC neural network of Figure 9(C) has been accurately expressed by mathematical equations at the level of the neural connection of which cell is connected to which cell. This neural connection is shown in detail as the forward connection table in Section **S2.3.2** of Supplementary materials.
12. Things to note The above is represented by the complex equations such as equation(2.2-5), and you may think that it is difficult for cerebral cells to execute them. However, it is not a problem for the following reasons. These mathematical equations are only needed to determine to which SDCs (n,t) each MDC (τ_X_, τ_Y_) is connected. Once the neural network composed of these neural connections has been determined, no calculation is required for the operation of the SDC, and thus there is no need to worry about the above. That is, responses of SDCs, which are performed in the neural network determined above, do not require these equations, and do not suffer from the above problems. Note the following two things. First, I think that these equations represent the neural networks after learning is complete. Thus, they are not necessary for the cell operations that are performed by the networks after learning. Next, in Supplementary materials describing simulation procedures for the neural network, these equations are used only to calculate the neural network (see Sections **S2.3.1** and **S2.3.2** of the materials) and not to calculate cell responses (see Section **S3.2** of the materials). In this way, these equations are only used to calculate the neural networks and not to calculate the cell responses.

The neural network that detects local motions in Figure 9(B) is also represented by complex mathematical-equations such as the Hough and inverse Hough transforms, but there is no problem for the same reasons as described above.

Since I have shown that the cells in Figure 9 are arranged in 2D (or 3D) arrays, you may think that it is difficult for the cerebrum to form such an orderly array. However, it is not a problem for the following reasons. That is, no matter how deformed the array of each cell types in Figure 9 is, if the neural network is correct, the time-to-contact to a plane and its 3D orientation can be detected. This is explained as follows. In Figure 9, the various types of cells constitute the arrays: that is, the (B) is constituted with circular, rectangular, and cuboidal arrays, and the (C) is constituted with a cylindrical array. No matter how deformed these arrays are, as long as the neural connections of which cells are connected to which cells are maintained, the function of each cell is performed correctly, and finally, in the (C), the time-to-contact to the plane and its 3D orientation can be detected: as an extreme case of this deformation, it is also conceivable that the cuboidal array above is deformed two-dimensionally. In short, no matter how deformed the arrays are, if the neural network in Figure 9 is correct, each cell will behave correctly.

It is surprising that the neural network (Figure 9) that detects the time-to-contact to a plane and its 3D orientation is performed in the linear processings (i.e. in the processings by addition and subtraction), except for the spatio-temporal correlation of Figure 9(B)(d) (see Section **2.1.2**(4)). These linear processings can be performed with the postsynaptic excitation and inhibition by which the network is made up. Similarly, the neural network (i.e. Figure 17 described later) for detecting the shortest distance to a plane and its 3D orientation is represented by a neural network that performs the linear processings.

### 2.3. Simulation results

#### 2.3.1. Case where MDCs are arranged continuously

Consider the case where the MDCs in Figure 9(B)(f) are continuously arranged. That is, the array (τ_X_, τ_Y_) of MDCs has a continuous address of the real number.

Using the mathematical expression in Section **2.2**, a computer simulation was performed to detect the time-to-contact to a plane and its 3D orientation (Kawakami et al., 2003, 2000).

The result is shown in Figure 12 (Kawakami et al., 2003, 2000). A case approaching a perpendicular plane (which has eight points on it) 3 m away was calculated: this approaching speed is 1 m/s, and n (i.e. the 3D orientation) = v (i.e. the movement direction) = O (i.e. the center of the visual field). Based on physiological data (Mastronarde, 1987a, 1987b; Kawakami & Okamoto, 1996a), 50 ms was used as the unit time t_d_ in equation(1.0-2). Therefore, the normalized time-to-contact t_C_ to reach the plane is calculated as (3m/(1m/s))/50ms, i.e. 60.

**Figure 12.**
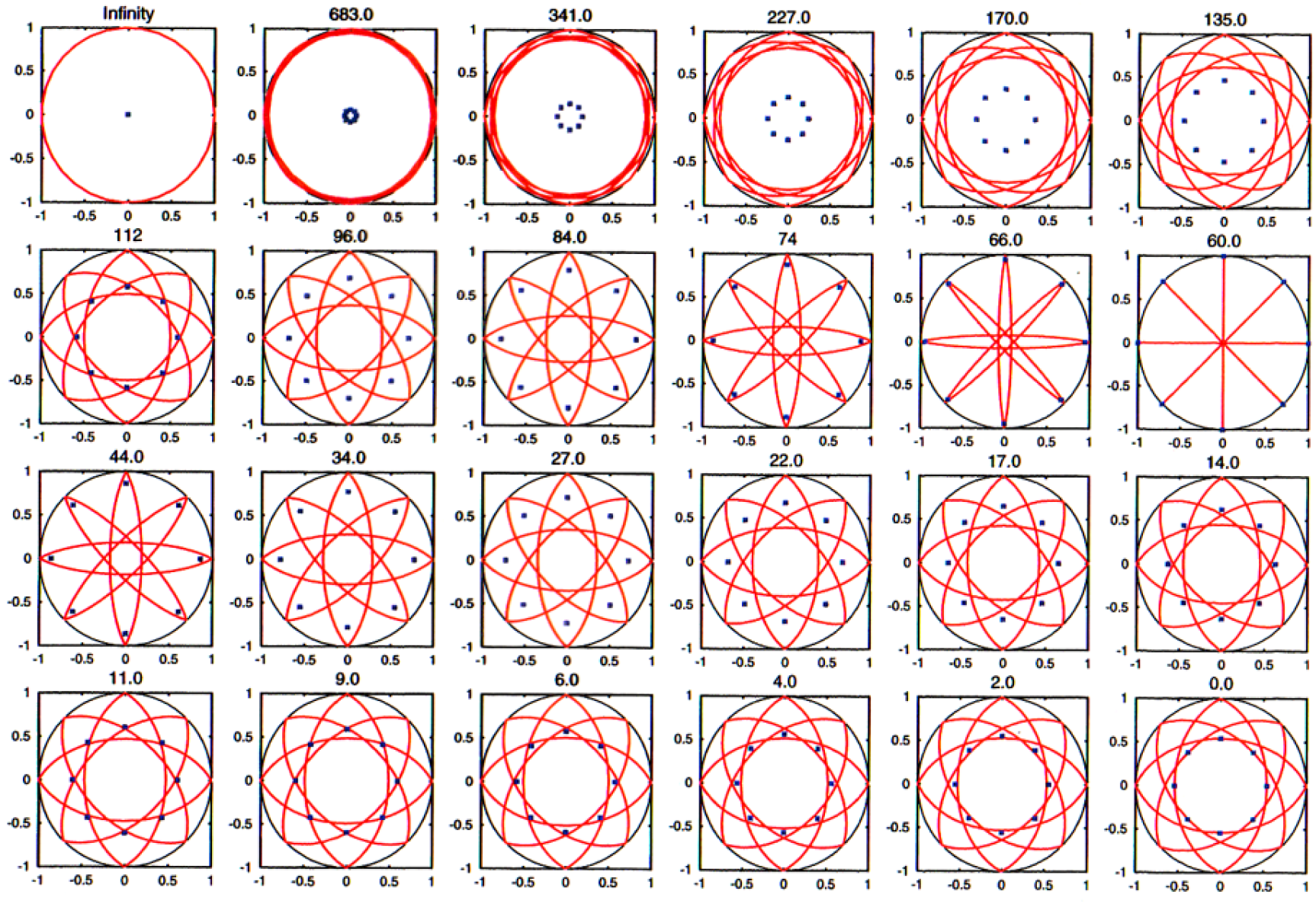
Simulation results detecting the normalized time-to-contact t and 3D orientation n when MDCs are arranged continuously. The simulation results when the MDCs of Figure 9(B)(f) are continuously arranged are shown. Cross sections of all heights t in the SDC column of Figure 9(C) were arranged. In a cross section of each height t, the blue small-square shows the position P_t_, which corresponds to the pink dot in Figure 11(B), predicted by equation(2.2-8). Also, the red curve shows a great circle, which corresponds to the pink great circle in Figure 11(B), obtained by polar transformation of the P_t_. As the height t increased, a circle enveloped by these great circles became smaller, and became zero at t=60. That is, the great circles intersected at one point with the given normalized-time-to-contact t=60. In addition, this intersection is at the visual field center O and coincides with the given 3D orientation n of the plane. Therefore, the given normalized time-to-contact t and 3D orientation n were detected as the coordinates of the intersection at t=60 of the SDC column. When t exceeds 60, the circle enveloped by these great circles becomes larger.

The cross sections of all heights t of the SDC column in Figure 9(C) are arranged in Figure 12. In a cross section of each height t, blue small-squares show the positions P_t_ predicted by equation(2.2-8) (see Figure 11(A)). Also, red curves show great circles obtained by polar transformation of these positions P_t_ using equation(2.2-10) (see Figure 11(A)). In the cross section of t = 0, the eight points given are shown because P_t_ is equal to the position P_0_ at the current time.

As the height t increased, a circle enveloped by these great circles became smaller, and became zero at t = 60. That is, these great circles intersected at one point in the given time (t = 60). In addition, their intersection is at the center of the visual field O, which coincides with the 3D orientation n of the given plane. Therefore, the given time-to-contac t_C_ and 3D orientation n were detected correctly as the coordinates of this intersection at t = 60 of the SDC column. When t exceeds 60, the circle enveloped by the great circles becomes larger.

In this way, it has been confirmed that the normalized time-to-contact t_C_ to the plane and its 3D orientation n are accurately detected using the mathematical expressions in Section **2.2**.

Let us compare the above results with the idea of Figure 5 which enabled these parameters n and t to be detected (see Section **1**). In conclusion, an arrangement of the eyeballs (Figure 5) for all times corresponds to the arrangement of the cross sections in Figure 12. In addition, the moment when the plane crosses on the eyeball center O_eye_ (Figure 5) corresponds to the cross section of t = 60 in Figure 12. Thus, the validity of the idea in Figure 5 has been confirmed by this simulation.

Let us explain why this confirmation above is reached. Based on the equidistant projection of Figure 10, a cross section of the column at each time (shown in Figure 12) is equivalent to the eyeball at the corresponding time (shown in Figure 5). In addition, points and great circles on the cross section for each time (shown in Figure 12) are equivalent to points and great circles on the eyeball for the corresponding time (shown in Figure 5), respectively. Therefore, these equivalencies have caused the confirmation above to be reached.

#### 2.3.2. Case where MDCs are arranged discretely

Consider the case where the MDCs in Figure 9(B)(f) are discretely arranged (see Section **4.3.3** later for details). That is, the array (τ_X_, τ_Y_) of MDCs has a discrete address of the integer: this discrete array will be shown in Figure 25(A)(i) later. The procedure for calculating the responses of discretely arranged cells (Figure 9) is shown in Supplementary materials. Its overview is described as follows.

First, the procedures for calculating the neural networks (i.e. for calculating the connection tables representing these networks) of the NDS simple cells (Figure 9(B)(c)) and MDCs (Figure 9(B)(f)) are shown in Sections **S2.1** and **S2.2** of Supplementary materials, respectively. Next, using these tables, the procedures for calculating the series of cell responses in Figure 9(B) are shown in Section **S3.1** of Supplementary materials. These procedures were also used in previous simulations (Kawakami & Okamoto, 1996a, 1995a, 1992d; Kawakami, 1996b; Kawakami et al., 2003, 2000, 1992b).

Next, the procedure for calculating the neural network (i.e. for calculating the connection table representing this network), which connects each MDC in Figure 9(B)(f) and the SDC column in Figure 9(C), is shown in Section **S2.3** of Supplementary materials. Using this table, the procedure for calculating the cell responses of the SDC column (Figure 9(C)) is shown in Section **S3.2** of Supplementary materials. These procedures were also used in previous simulations (Kawakami et al., 2003, 2000; Akima et al., 2017).

All these procedures were installed on a computer to simulate the responses of all cells in Figures 9(B and C). Figure 13 shows the results taken from previous reports (Kawakami et al., 2003, 2000). It is shown below that the normalized time-to-contact to a plane and its 3D orientation can be detected correctly. Figures 9(B and C) are consisted of the about 16 million cells and about 460 million neural connections between them.

**Figure 13.**
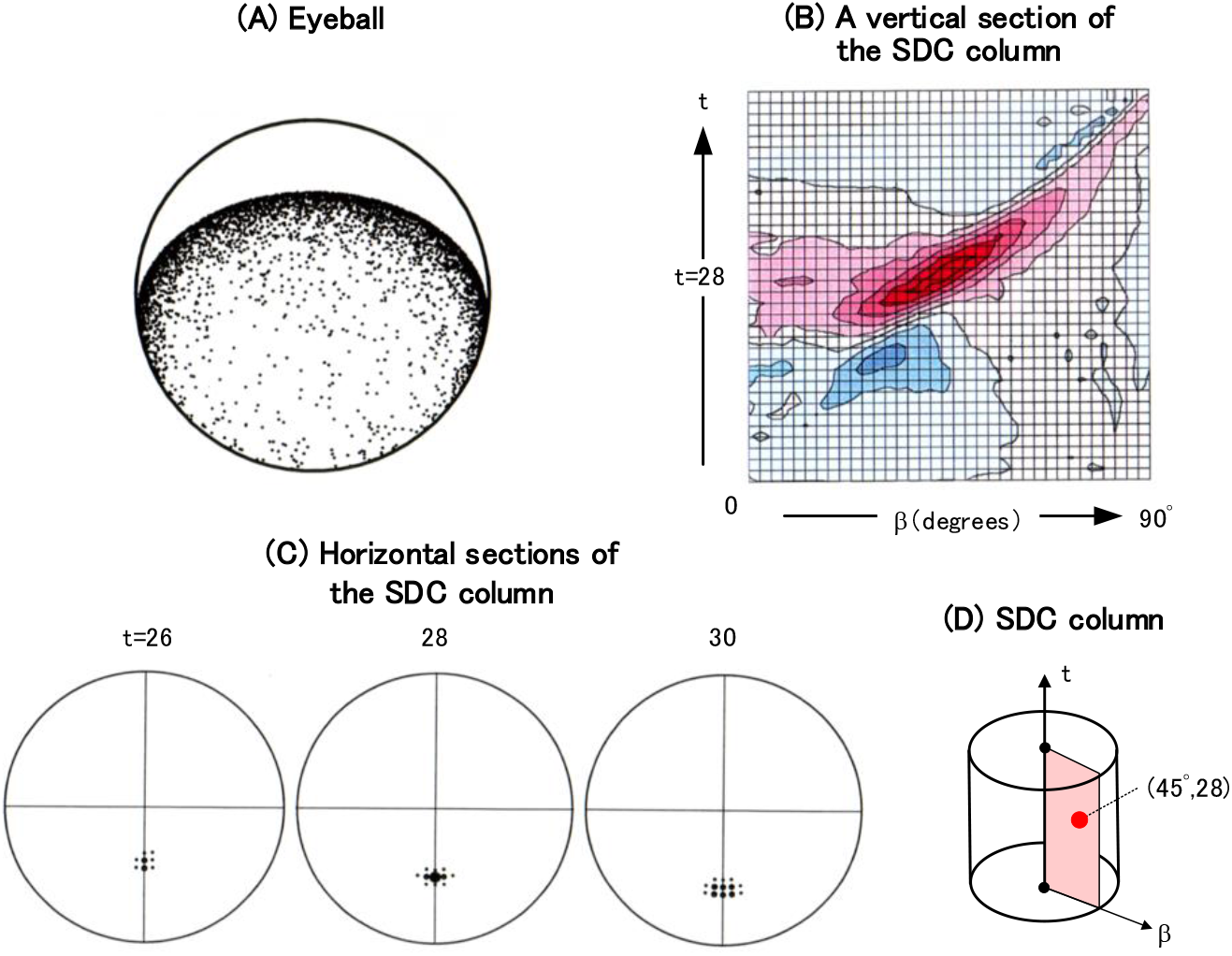
Simulation results detecting the normalized time-to-contact t and 3D orientation n when MDCs are arranged discretely. The simulation results when the MDCs of Figure 9(B)(f) are discretely arranged are shown. (A) The image of a plane inclined by 45 degrees on which random dots are drawn is shown on the eyeball. (B) A vertical cross-section (see (D)) including the maximum-firing SDC and the column-central axis t is shown in a contour map. Red represents a positive response, blue represents a negative response, the horizontal axis is the radial direction β of the column, and the vertical axis is the column height t. (C) Cross sections at three heights (centered at height t=28) of the SDC column are shown: a response of each SDC is displayed as the dot’s size. In a cross section of t=28, the cell fired maximally at 45 degrees from the center, correctly detecting the plane with a given normalized-time-to-contact of 28 and a 3D orientation of 45 degrees.

A case approaching a plane of 30m×30m in size 2.8 m away (on which random dots were drawn) was simulated: the plane is tilted 45 deg, and the approaching speed is 2 m/s. Figure13(A) shows an image in which this plane is reflected on the retina of the eyeball. The normalized time-to-contact t_C_ is calculated as (2.8m/(2m/s))/50ms, i.e. 28.

Figure13(C) shows the cross sections at three heights (centered at height t=28) of the SDC column: a response of each SDC is displayed as the dot’s size. In the cross section of t=28, the SDC at a position of 45 deg from the center fired maximum, and thus correctly detected the plane parameters given (i.e. the normalized time-to-contact of 28 and the 3D orientation of 45 deg). In addition, in a cross section that deviates by plus or minus 2 from the height of t=28 (i.e. in a cross section that deviates by plus or minus 0.1 seconds in terms of the non-normalization time T), the SDC’s responses are sharply reduced to less than half, and thus a sharp detection of the plane is performed in the height direction of the column.

In order to examine the responses of the SDC column in detail, a vertical cross-section (Figure 13(D)) including the maximum-firing SDC and the column-central axis is shown in Figure 13(B) in a contour map. Red represents a positive (or excitable) response and blue represents a negative (or inhibitory) response. The horizontal axis is the radial direction β of the column, and the vertical axis is the column height t.

The half-width of the firing pattern centered on the maximum-firing SDC (β=45 deg, t=28) is sharp at 4 in the t direction, but broad at 10 deg in the β direction. The cause that the β direction (i.e. the direction of a 3D orientation-detection) is broad is due to the crescent region of Figure 25(A)(ii) described later, which is caused from dx/d τ_2D_ in equation(4.3-1) (see Section **4.3.3.1**(1)).

The above simulation confirmed that the neural network of Figure 9 correctly detects the normalized time-to-contact to the plane and its 3D orientation.

### 2.4. How is this cell model used?

How do we avoid obstacles like planes and then walk around? For example, when walking toward a wall (i.e. a plane) such as a corridor, we experience that we notice the wall at a certain moment and change the direction of movement to avoid it. Let us consider this for the SDC column of Figure 9(C).

When walking toward the wall (Figure 14(A)), the time to reach the wall (such as t_1_ and t_2_) is detected one after another as the height coordinate in the SDC column (Figure 14(B)). When it approaches the critical time t_critical_ that is set, we change the direction of movement so that it is perpendicular to the 3D orientation n of the wall that is detected at the same time, and then continue to walk parallel to the wall. In this way, we can walk through winding corridors without bumping into each other. At this time, the cells in the SDC column fire while drawing a trajectory as shown in Figure 14(B) (Kawakami et al., 2002).

**Figure 14.**
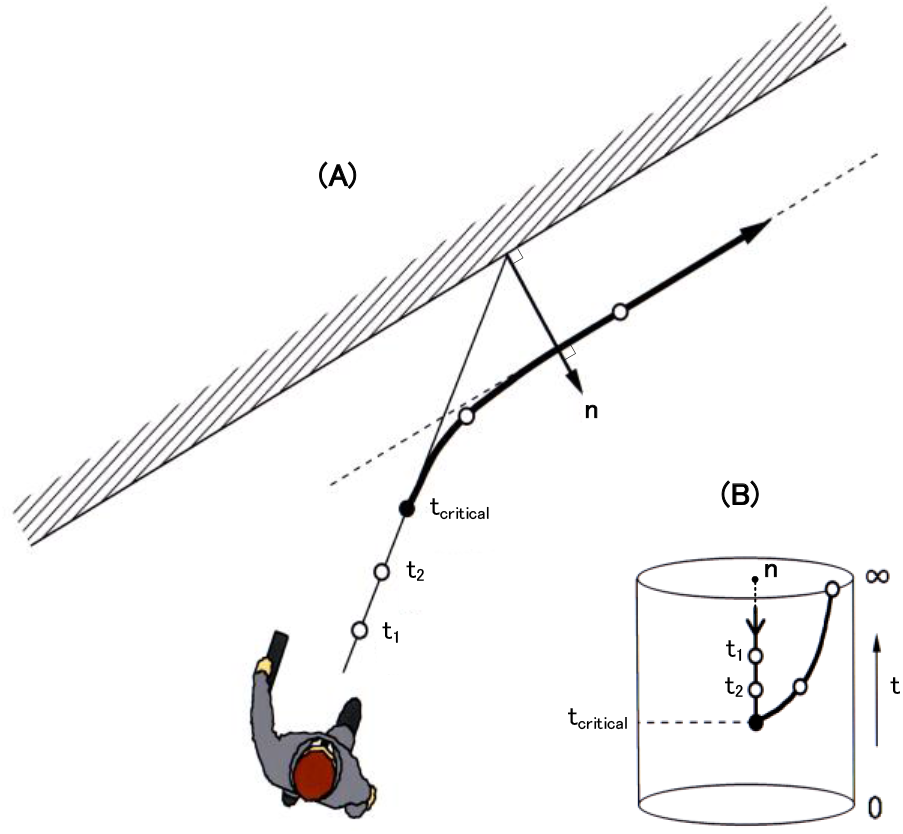
A firing trajectory within the SDC column during a wall avoidance. (A) When walking towards a wall (i.e. a plane), you may experience noticing the wall at a certain moment t_critical_ and changing the movement direction to avoid it. (B) In this experience, let us consider what is happening in the SDC column. As the height coordinate of each SDC that fires in the column, a time to reach the wall (such as t_1_ and t_2_) is detected one after another. When this time approaches the critical time t_critical_ that is set, the movement direction is changed so as to be perpendicular to the 3D orientation n of the wall detected at the same time, and continues to walk parallel to the wall (see (A)). At this time, SDCs in the column fire with a trajectory like the (B).

The landing of a airplane shown in Figure 1 can be done in the same way. That is, the time-to-contact to the runway (i.e. to the plane) is detected one after another as the height coordinate of the SDC column. When the critical time t_critical_ that is set is approached, the pilot raises the nose of the airplane in a direction perpendicular to the 3D orientation n of the runway, and then lands.

This model of Figure 9 enables the time-to-contact to a plane and its 3D orientation at each instant of movement (i.e. at every time t_d_ (=50 ms)) to be detected independently of movement speed and with a wide visual-field of 180 degrees. This speed independence is due to the fact that the speed V contained in equation(A1-4) in Appendix **1** is cancelled between the numerator and the denominator. In addition, the wide visual-field of 180 degrees is due to the use of the eyeball that is equivalent to a fisheye lens.

It is reasonable to detect the fixed critical time t_critical_ regardless of the movement speed and avoid obstacles. That is, since the time to start this avoidance is constant due to the response speed of the muscle and the like, it is difficult to change this avoidance time according to the speed. Therefore, the cell model of Figure 9, which can avoid obstacles in a fixed critical time t_critical_ regardless of speed, is reasonable. This makes it possible to start avoiding from a distance when the speed is fast, or to start avoiding after approaching when the speed is slow. Therefore, this enables the avoidance action to be automatically performed at the fixed critical time t_critical_ regardless of the speed. Based on the above considerations, I think that cells in the cerebrum may detect this time t_critical_. In support of this, Gannets and monkeys were reported to be able to avoid obstacles at a fixed time-to-contact (i.e. at the critical time t_critical_) regardless of the movement speed (see Section **4.2** later).

## 3. A cell model for detecting the shortest distance to a plane and its 3D orientation

A cell model that detects the shortest distance D and the orientation n in Figure 1 has been reported (Kawakami et al., 2003, 2000). However, the explanation is insufficient due to page limit. Therefore, it will be described in detail below.

Figure 15 shows a cross section of Figure 1 viewed from the side, and there is a relationship between the parameters in the following equation: V is a movement speed.

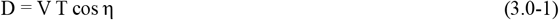

**Figure 15.**
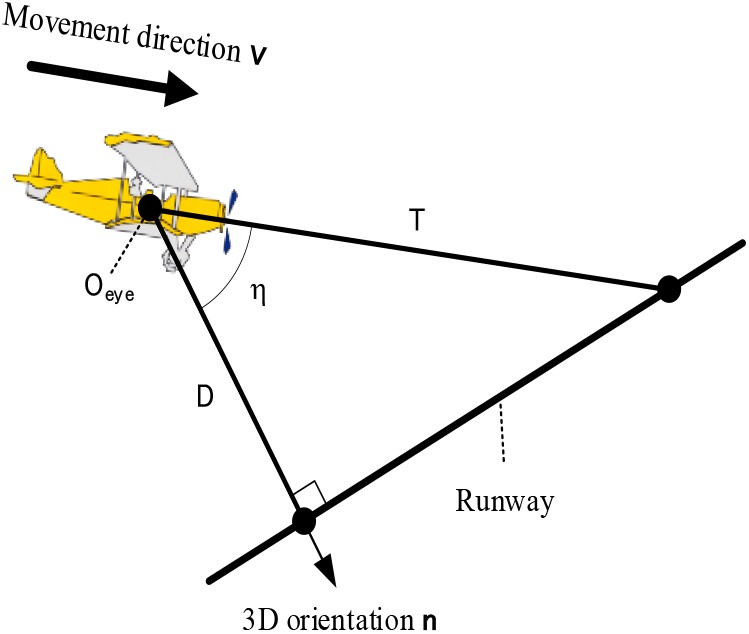
Relationship between parameters related to the plane. A cross section of Figure 1 viewed from the side is shown, and there is a relationship of equation(3.0-1) between the parameters related to the plane: the O_eye_ is the eyeball center of the pilot. Deforming this equation, there exists a relationship of equation(3.0-3) among the normalized time-to-contact t, the normalized shortest-distance d, the 3D orientation n, and the movement direction v. This relationship yields a small circle transform of equation(3.0-6) (see Appendix **6**). This transform enables a neural network, that detects the normalized shortest-distance d to a plane and its 3D orientation n, to be modeled in Section **3.1.2**.

By normalizing the shortest distance D by the distance Vt_d_ moved in the unit time t_d_, the normalized shortest distance d is defined by the following equation.

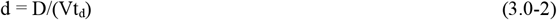

Here, since cosη=(n •v), combining equations(3.0-1 and 2) and equation(1.0-3) yields the following relationship, where (T–T_0_) in equation(1.0-3) is set to T. This equation allows us to convert d and t to each other.

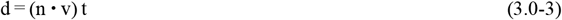

Combining this relationship with the polar and cross-ratio transforms in Section **1** enables the small-circle transform to be derived (Appendix **6**; Kawakami et al., 2003, 2000). This transform converts the current-time position P_0_ into a small circle with radius R centered on P_0_ (see Figure 16), and has an important property that the 3D orientation of a plane n lies on this small circle. The radius R above is calculated by the following equations, which were derived as equations(A6-6~8) in Appendix **6**.

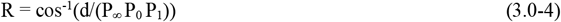

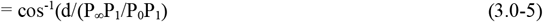

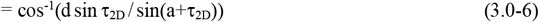

**Figure 16.**
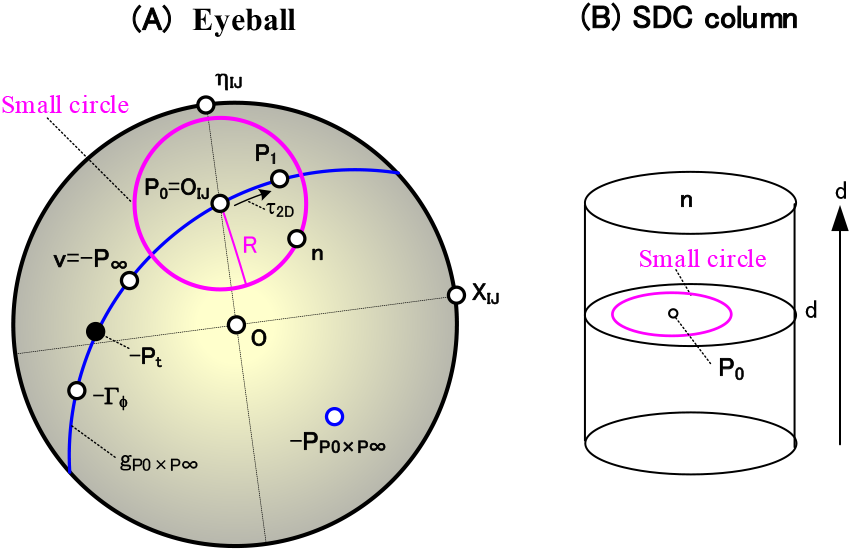
Detection of the normalized shortest-distance d and 3D orientation n using the small-circle transform. (A) The parameters on the eyeball are shown. The position P_0_ at a current time T_0_ moves as follows. That is, P_0_ moves to P_1_ after time t_d_, and reaches position P_∞_ opposite to the movement direction v after infinite time: the movement from P_0_ to P_1_ is the local motion τ_2D_, and P_p0×p∞_ is the pole of the blue movement trajectory. The small circle transform of equation(3.0-6) converts this P_0_ into a pink circle with radius R centered on P_0_. There is an important property that the 3D orientation n of a plane to be determined exists on this small circle. Here, X_IJ_ and η_IJ_ belong to each RF coordinate system (see equations(A4-4 and 10) in Appendix **4**) and P_0_=O_IJ_ (see equation(A4-3) in Appendix **4**). Γ_ϕ_ is the movement direction from P_0_ to P_1_ (see equation(A4-7) in Appendix **4**). (B) The SDC column is shown. The small-circle transform causes each position P_0_ on the cross section of the column at a height d to be converted into a pink small-circle with radius R centered on P_0_. When such small circle is drawn for all positions P_0_ and all heights d, these small circles intersect only at a certain height d_C_ (see Figure 17(C) for d_C_). The normalized shortest-distance d_C_ to the plane is detected as this height coordinate, and its 3D orientation n is detected as the polar coordinate of this intersection. The SDC neural network that performs this detection is expressed by mathematical equations in Section **3.2**

**Figure 17.**
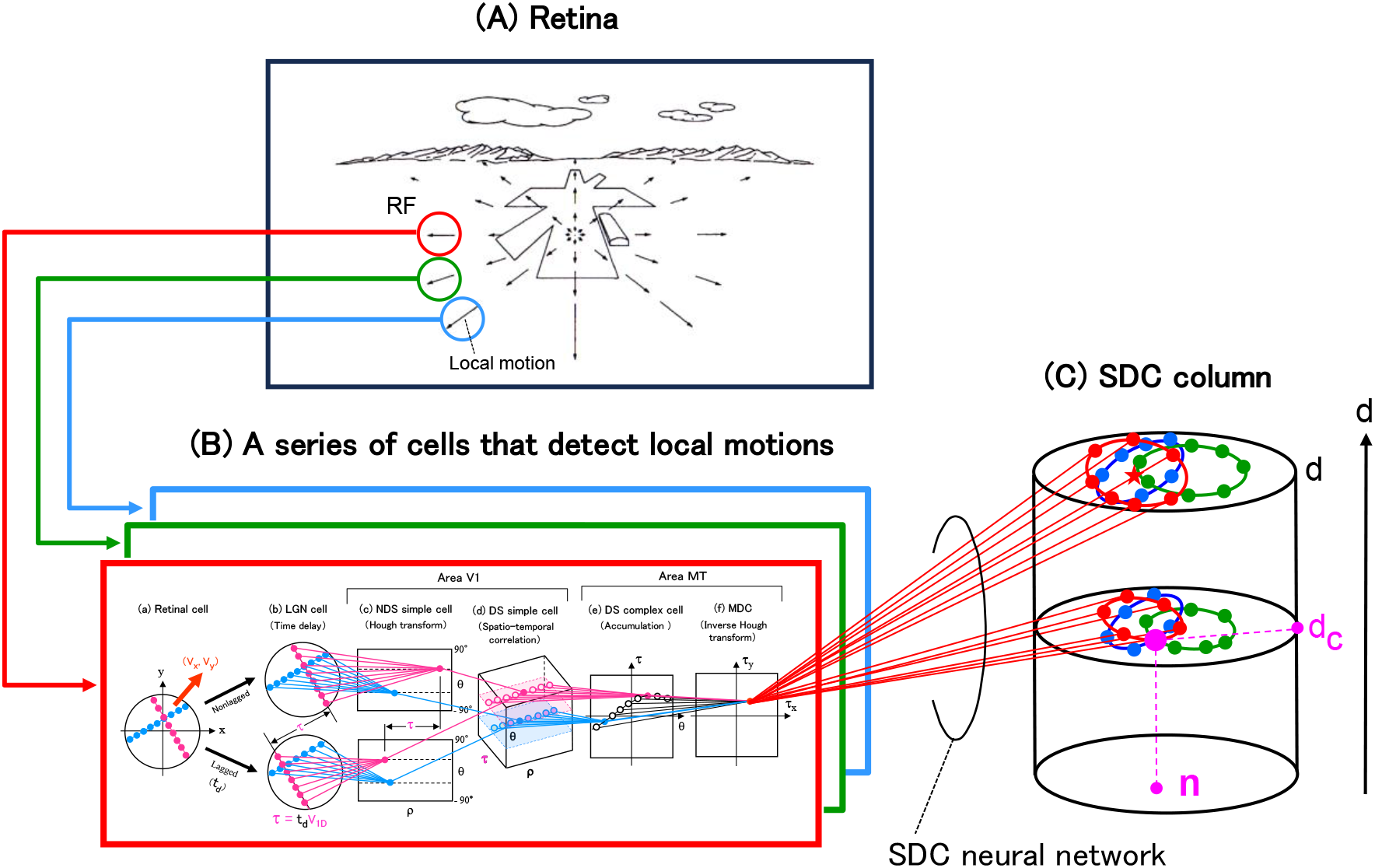
A cell model for detecting the shortest distance to a plane and its 3D orientation. A series of modeled cells that detect two parameters in Figure 1 (i.e. the normalized shortest-distance d to a plane and its 3D orientation n) is shown. (A) This is the same as Figure 9(A). An OFP on the retina consists of local motions, each of which occurs in a small region (i.e. a RF). (B) This is the same as Figure 9(B). This local motion is detected by a series of cells (i.e. LGN cells, NDS simple cells, DS simple cells, DS complex cells, and MDCs). In this series, a series of transforms (i.e. time delay, Hough transform, spatio-temporal correlation, accumulation, and inverse Hough transform) are performed, respectively: in LGN cells, the 2D convolution with a DOG filter is also performed at the same time to emphasize the contours of figures. The pink and blue straight lines represent the neural networks connecting each cell array, and perform the above transforms. (C) A column composed of SDCs is shown. Each SDC detects the normalized shortest-distance d to a plane as the height coordinate, and detects the 3D orientation n as the polar coordinate of the cross section at this height. The red straight lines represent the neural network connecting each MDC in the (B)(f) and the SDC column in the (C), and performs the small-circle transform.

Here, (P_∞_ P_0_ P_1_) is a simple ratio of the P_∞_, P_0_, and P_1_ on the eyeball in Figure 16(A). The P_∞_P_1_ and P_0_P_1_ are directed segments, the central angles (i.e. a and τ_2D_) were shown in Figure 8(B), and the variable a was calculated by equation(1.0-7).

Combining this small-circle transform with Section **2.1.4** enables the normalized shortest distance to a plane and its 3D orientation to be detected by the following steps (Figure 16).

[Step 1] Give any two parameters in Figure 16(A) (i.e. the position P_0_ on the eyeball at a current time and a local motion τ_2D_ from P_0_ to P_1_). Then, assuming that the movement direction v is known, calculate P_∞_ by equation(1.0-1).

[Step 2] Set a normalized shortest-distance d arbitrarily.

[Step 3] Convert this position P_0_ into a pink small circle with a radius R centered on P_0_ (Figure 16(A)): the radius R is calculated by equation(3.0-6). There is an important property that the 3D orientation n of a plane to be determined lies on this small circle.

[Step 4] Draw this small circle on the cross section of the height d in Figure 16(B): this circle is shown in pink.

[Step 5] Draw such small circle for all positions P_0_ at every height d. These small circles do not intersect at the general height d, but intersect only at a certain height d_C_. Due to the property described in the step 3, this intersection is the 3D orientation n of the plane. Also, the height at which this intersection occurs is the normalized shortest distance d_C_ to the plane.

In this way, knowing P_0_ and τ_2D_ enables the normalized shortest distance d_C_ to the plane and its 3D orientation n to be detected using the small-circle transform.

Section **3.1** will model a series of cells that performs these steps (see Figure 17), and Section **3.2** will enable these modeled cells to be expressed accurately by mathematical equations. Based on a computer simulation, Section **3.3** will confirm that the distance d_C_ and the orientation n are determined correctly by these equations. In addition, Supplementary materials show the procedures for simulating these modeled cells with a computer.

### 3.1. Cell model

Based on the above five steps, let us model a series of cells in Figure 17. This figure consists of the (A), (B), and (C), where the (A) and (B) are the same as in Figure 9. In the retina of the (A), an OFP is generated by the movement of an opponent (or object). A series of cells in the (B) detects a local motion within each RF: the motion constitutes this OFP (see Section **2.1.1**). The (C) is a cylindrical cell array named an SDC column. Each SDC in the column integrates all local motions (τ_X_, τ_Y_) detected in the (B)(f) to extract the OFP in the (A) and then to detect the shortest distance to a plane and its 3D orientation. This series of modeled cells are described below.

#### 3.1.1. SDC column for detecting planes

A cross section at each height of the SDC column in Figure 17(C) (or in Figure 16(B)) is equivalent to a disk to which the eyeball of Figure 16(A) has been converted by the equidistant projection in Figure 10: this conversion is the same as in Section **2.1.3**. A pink small circle on the eyeball (Figure 16(A)) is converted to a pink small circle on the disk (Figure 16(B)).

Stacking this disk with a normalized shortest-distance d as a height coordinate yields a cylinder (α,β,d). Since the 3D orientation n of the plane is expressed as (α_n_,β_n_) in the polar coordinate, this cylinder is expressed as (n,d), which is the SDC column of Figure 17(C). About 10,000 SDCs are arranged in this cylinder (n,d). Each SDC detects the normalized shortest-distance d as its height coordinate, and detects the 3D orientation n as its polar coordinate of the cross section.

#### 3.1.2. Neural network connecting MDCs and the SDC column

Based on the five steps above, let us determine a neural network that connects each MDC of Figure 17(B)(f) and the SDC column of Figure 17(C).

[Step 1] Give two parameters of Figure 16(A) (i.e. the position P_0_ on the eyeball at a current time and a local motion τ_2D_ from P_0_ to P_1_), as follows. First, give the position P_0_ as the RF center O_IJ_ based on equation(A4-3) in Appendix **4**. Next, give the local motion τ_2D_ by the following equation, where (τ_X_, τ_Y_) is the coordinate of the MDC in Figure 17(B)(f).

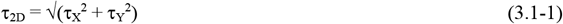

Further, assuming that the movement direction v is known, calculate P_∞_ by equation(1.0-1).

[Step 2] Set a normalized shortest-distance d arbitrarily.

[Step 3] Calculate a radius R used for the small circle transform by the following equation (see equation(3.0-6)). A variable a used in this equation is calculated by equation(1.0-7).

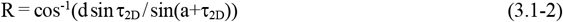

[Step 4] Plot the current position P_0_ on the cross section of height d in Figure 17(C): this plotted P_0_ is shown as a red mark ★. Then, draw a red small circle with radius R, centered on P_0_, on the cross section. There is an important property that the 3D orientation n of a plane to be determined exists on this small circle.

Finally, connect all SDCs (indicated as red dots) on this small circle to the MDC (τ_X_,τ_Y_) of Figure 17(B)(f). This connection shown as red straight lines represents an SDC neural network in Figure 17(C). The above is for the red RF in Figure 17(A), but for the green and blue RFs, the green and blue small circles and all cells SDCs on each small circle are drawn: the neural networks are omitted for ease of viewing. These small circles do not intersect at a single point.

[Step 5] When such a small-circle drawing is performed for all heights d, the small circles intersect at one point only at some height d_C_ (see Figure 17(C) for d_C_). A normalized shortest-distance d_C_ to the plane is detected as the height coordinate of the pink intersection cell, and its 3D orientation n is detected as the polar coordinate of this cell.

In this way, a neural network connecting each MDC in Figure 17(B)(f) and the SDC column in Figure 17(C) has been modeled. This network detects the normalized shortest-distance d to the plane and its 3D orientation n. Although the essential points of the network have been explained with an emphasis on comprehensibility, Section **3.2** will show that the network is expressed accurately using mathematical equations. Section **3.3** will also show that the validity of this network is confirmed by computer simulations.

### 3.2. Representation of the SDC neural network by mathematical equations

Let us express the neural network, described in Section **3.1.2**, by mathematical equations using the form of a flowchart, as follows (see Figure 16). Here, note that important properties such as the relationship of various parameters on the eyeball are shown in Appendix **4**.

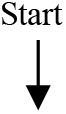

1. Give a movement direction v (see Figure 16(A)) Calculate P_∞_ using the following equation (see equation(1.0-1)).

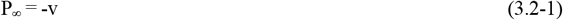
2. Sweep a RF center O_IJ_ 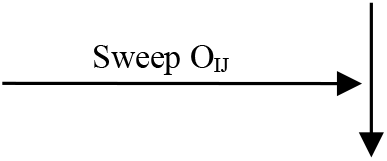 Calculate P_0_ using the following equation (see equation(A4-3) in Appendix **4**).

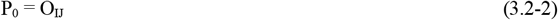
3. Sweep a normalized shortest-distance d 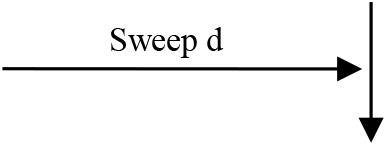
4. Sweep a MDC array (τ_X_,τ_Y_) 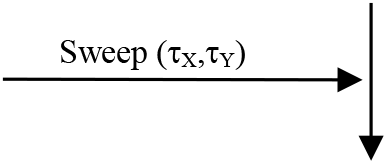 Convert this (τ_X_, τ_Y_) to its polar coordinate (τ_2D_,ϕ) by the following equation (see Figure A4(D) in Appendix **4**).

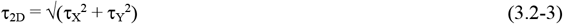

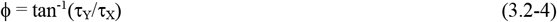
5. Calculate the radius R used for the small-circle transform by the following equation (see equation(3.0-6)).

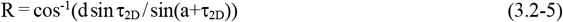 The variable a used above is calculated by the following equations (see equations(1.0-7 and 9)).

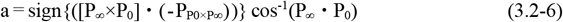

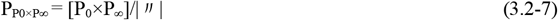
6. Calculate a small circle with radius R centered at position P_0_ (Figure 16(A)) First, P_0_ expressed by equation(3.2-2) is represented in an orthogonal coordinate system by the following equation.

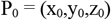 Convert the P_0_ to polar coordinate (α_0_,β_0_) by the following equation.

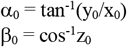 Next, when an arbitrary point on the small circle is represented as a polar coordinate (α,β), there is the following relationship (see equation(A7-1) in Appendix **7**).

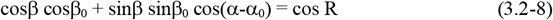 This is the mathematical expression of the pink small-circle with radius R centered at position P_0_ (see Figure 16(A)). There is an important property that the 3D orientation n of a plane to be determined lies on this small circle.
7. Draw this small circle in the SDC column Using the equidistant projection of Figure 10, convert this small circle on the eyeball into a pink small-circle on the cross section at the height d of the SDC column in Figure 16(B). Then, draw this converted small circle on the cross section. Note that the small circle on the cross section is represented by the same equation(3.2-8) as that on the eyeball (see Section **A7.2** in Appendix **7**).
8. Connect all cells SDC on this small circle to the MDC (τ_X_,τ_Y_) Connect all SDCs (indicated as red dots in Figure 17(C)) on this small circle at height d to the MDC (τ_X_,τ_Y_) of Figure 17(B)(f). This connection shown as red straight lines represents an SDC neural network in Figure 17(C). 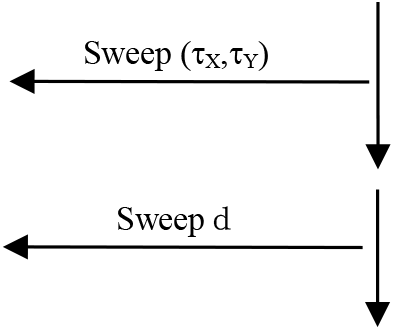
9. Up to this point, the red neural network in Figure 17(C) have been created. That is, the neural network for the red RF in Figure 17(A) has been completed. 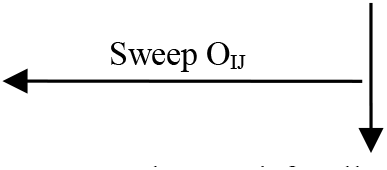
10. A neural network for all RFs in Figure 17(A), including red, green, and blue, has been completed. Each connection of the neural network (Figure 17(C)) connecting the MDCs and the SDC column has been accurately expressed one by one using mathematical equations. 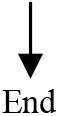

In this way, the SDC neural network of Figure 17(C) has been accurately expressed by mathematical equations at the level of the neural connection of which cell is connected to which cell. This neural connection is shown in detail as the forward connection table in Section **S2.4.2** of Supplementary materials.

The above is represented by the complex equations such as equation(3.2-5), and you may think that it is difficult for cerebral cells to execute them. However, that worry is not a problem for the same reasons as in Section **2.2**(12).

Since I have shown that the cells in Figure 17 are arranged in 2D (or 3D) arrays, you may think that it is impossible for the cerebrum to form such an orderly array. However, that worry is not a problem for the same reasons as in Section **2.2**(12). That is, no matter how deformed the array of each cell types in Figure 17 is, if the neural network is correct, the shortest distance to a plane and its 3D orientation can be detected.

### 3.3. Simulation results

#### 3.3.1. Case where MDCs are arranged continuously

Consider the case where the MDCs in Figure 17(B)(f) are continuously arranged. That is, the array (τ_X_, τ_Y_) of MDCs has a continuous address of the real number.

Using the mathematical expression in Section **3.2**, a computer simulation was performed to detect the shortest distance to a plane and its 3D orientation (Kawakami et al., 2003, 2000).

The result is shown in Figure 18 (Kawakami et al., 2003, 2000). A case approaching a perpendicular plane (which has eight points on it) 3 m away was calculated: this approaching speed is 1 m/s, and n (i.e. the 3D orientation) = v (i.e. the movement direction) = O (i.e. the center of the visual field). Based on physiological data (Mastronarde, 1987a, 1987b; Kawakami & Okamoto, 1996a), 50 ms was used as the unit time t_d_ in equation(1.0-2). Therefore, the normalized shortest-distance d_C_ to the plane is calculated as 3m/(1m/s×50ms), i.e. 60.

**Figure 18.**
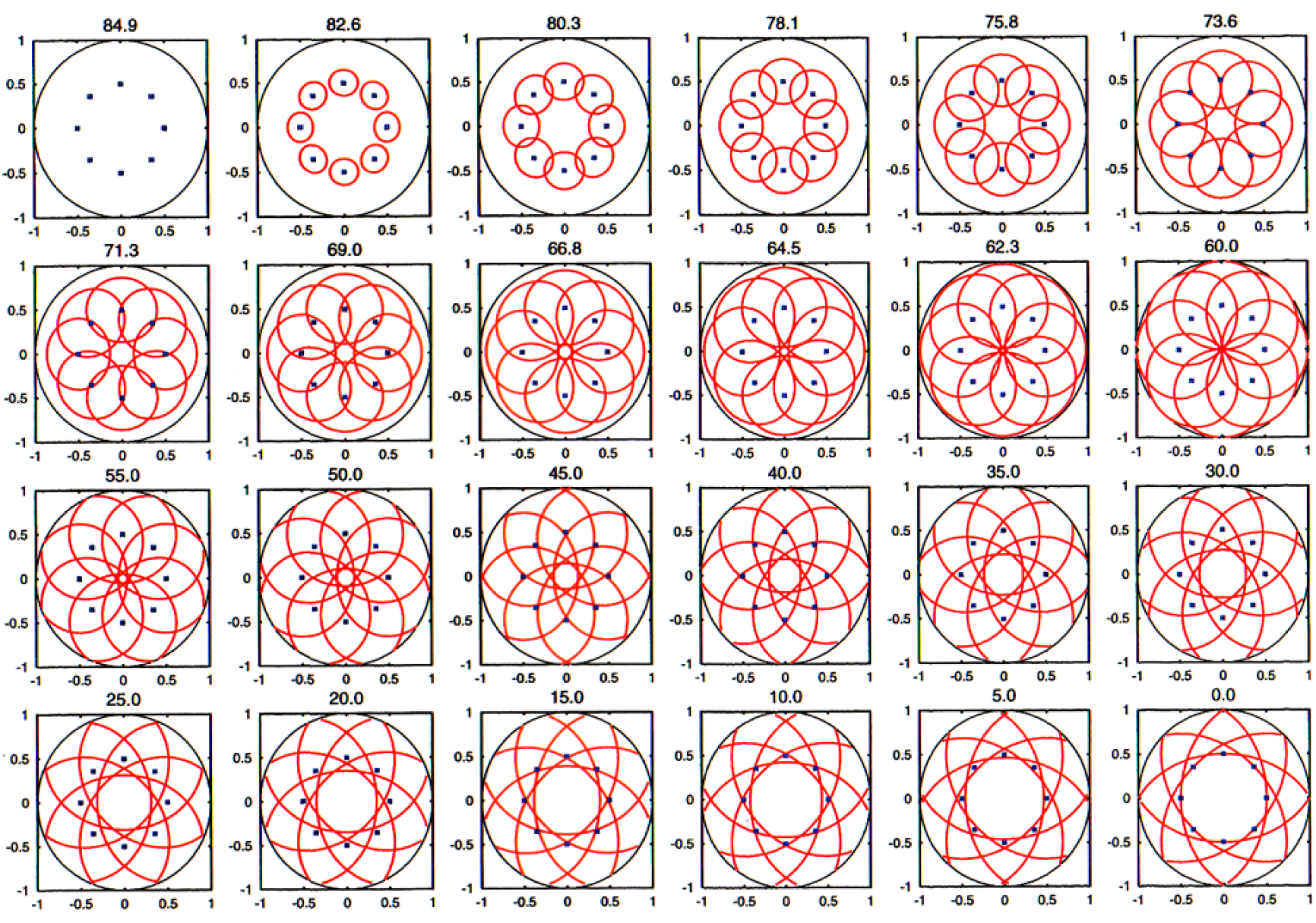
Simulation results detecting the normalized shortest-distance d and 3D orientation n when MDCs are arranged continuously. The simulation results when the MDCs of Figure 17(B)(f) are continuously arranged are shown. Cross sections at all heights d in the SDC column of Figure 17(C) were arranged. In a cross section at each height d, the blue small-square shows the position P_0_ at a current time that corresponds to Figure 16(B) and is constant regardless of the height. Also, the red curve shows a small circle with radius R of equation(3.0-6) centered on each P_0_: this circle corresponds to the pink small-circle in Figure 16(B). As the height d increased, a circle enveloped by these small circles became smaller, and became zero at d=60. That is, the small circles intersected at one point with the given normalized shortest-distance d=60. In addition, this intersection is at the visual field center O and coincides with the given 3D orientation n of the plane. Therefore, the given normalized shortest-distance d and 3D orientation n were detected as the coordinates of the intersection at d=60 of the SDC column. When d exceeds 60, the circle enveloped by these small circles becomes larger.

**Figure 19.**
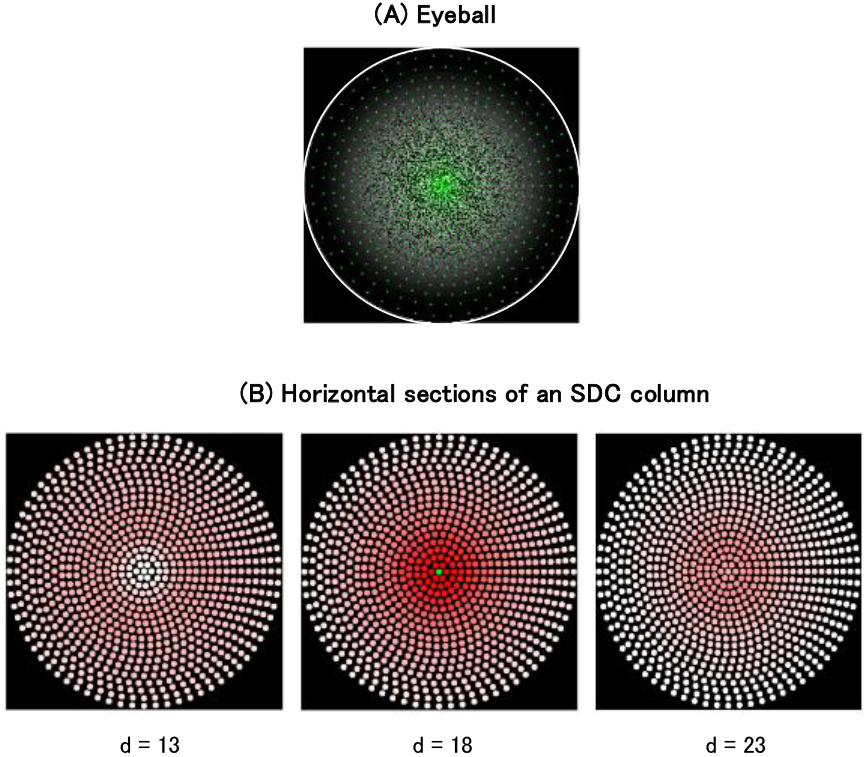
Simulation results detecting the normalized shortest-distanct d and 3D orientation n when MDCs are arranged discretely. The simulation results when the MDCs of Figure 17(B)(f) are discretely arranged are shown. (A) An image of a perpendicular plane on which random dots are drawn is shown on the eyeball. (B) Responses of two cross sections d=18±5 are shown centered on the horizontal cross section (d=18) of the SDC column: each dot on these cross sections represents a SDC. In the height direction, the cross section at d=18 had the largest response, and in that section, the center shown as a green dot had the largest response. This indicates that the SDC shown as the green dot detected the normalized shortest-distance of d=20 and 3D orientation of n corresponding to the visual field center.

**Figure 20.**
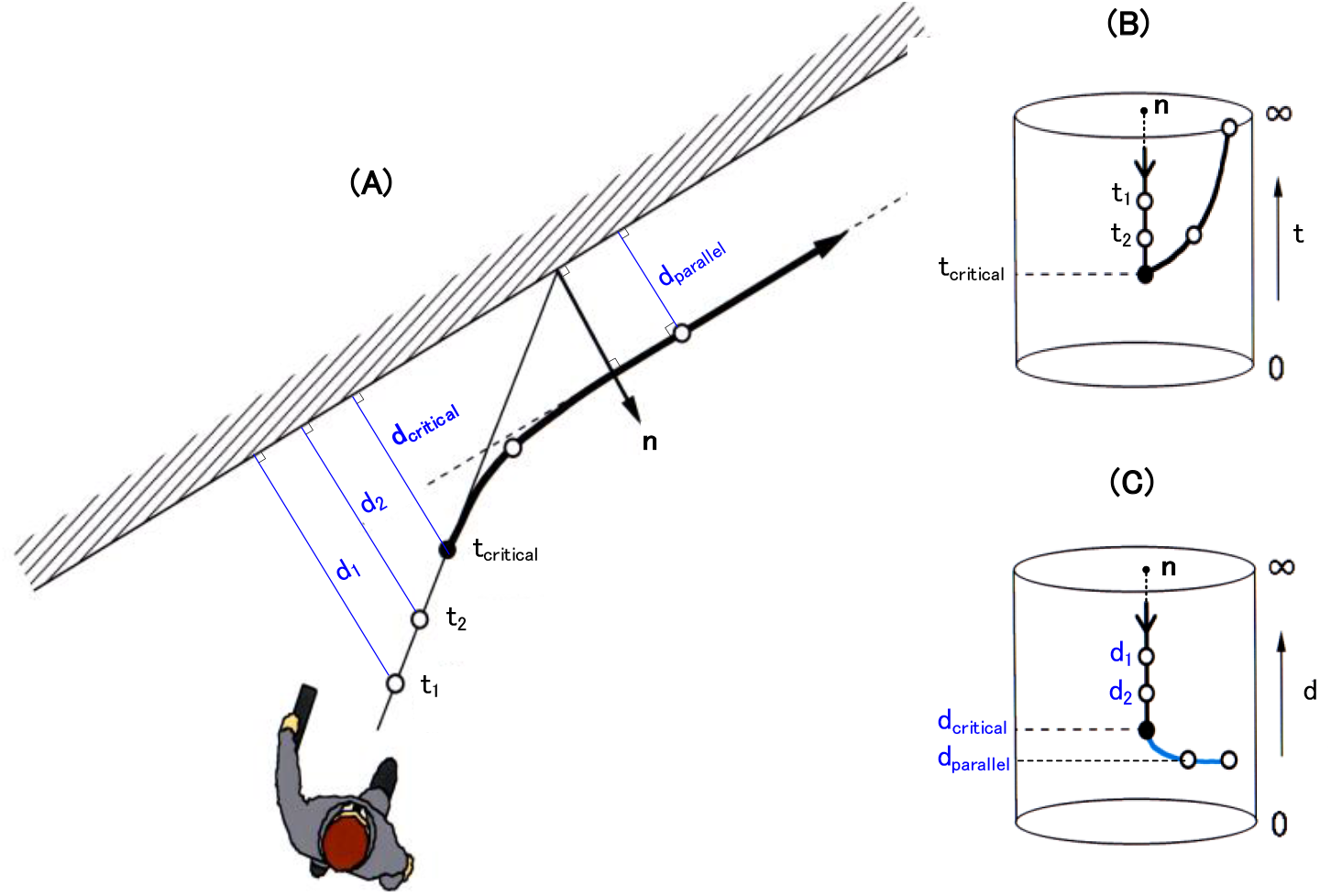
A firing trajectory within the SDC column during a wall avoidance. (A) In Figure 14, it was mentioned that the normalized time-to-contact t is essential to avoid a plane such as a wall. Consider how it would be useful if the normalized shortest-distance d was added to this. (B) A time to reach the wall is detected as a height coordinate of the SDC column, t_1_, t_2_, and so on. When the time approaches the critical time t_critical_ that is set, as shown in the (A), the movement direction is changed so as to be perpendicular to the 3D orientation n of the wall detected at the same time, then continuing to walk parallel to the wall. (C) Consider what happens when the normalized shortest-distance d is added to this. As a height coordinate of the SDC column, the shortest distances to the wall are detected one after another as d_1_, d_2_, and d_critical_. However, this distance does not directly lead to an avoidance behavior. The shortest distance d is useful only after you change the movement direction and keep walking parallel to the wall. As shown in the (A), while keeping the shortest distance d_parallel_, you can keep walking parallel to the wall. The SDC draws such a trajectory within the column and fires. Thus, the normalized shortest-distance d is important for walking parallel to the wall.

The cross sections of all heights d of the SDC column in Figure 17(C) are arranged in Figure 18. In a cross section of each height d, blue small-squares show the positions P_0_ at a current time (Figure 16(A)): the positions are constant regardless of the height. Also, each red curve shows a small circle with radius R (calculated by equation(3.2-5)) centered on P_0_ (Figure 16(A)).

As the height d increased, a circle enveloped by these small circles became smaller, and became zero at d=60. That is, these small circles intersected at one point in the given shortest-distance (d=60). In addition, their intersection is at the center of the visual field O, which coincides with the 3D orientation n of the given plane. Therefore, the given shortest-distance d_C_ and 3D orientation n were detected correctly as the coordinates of this intersection at d=60 of the SDC column. When d exceeds 60, the circle enveloped by the small circles becomes larger.

In this way, it has been confirmed that the normalized shortest-distance d_C_ to the plane and its 3D orientation n are accurately detected using the mathematical expressions in Section **3.2**.

#### 3.3.2. Case where MDCs are arranged discretely

Consider the case where the MDCs in Figure 17(B)(f) are discretely arranged (see Section **4.3.3** later for details). That is, the array (τ_X_, τ_Y_) of MDCs has a discrete address of the integer: this discrete array will be shown in Figure 26(A)(i) later. The procedure for calculating the responses of discretely arranged cells (Figure 17) is shown in Supplementary materials. Its overview is described as follows.

First, the procedures for calculating the neural networks (i.e. for calculating the connection tables representing these networks) of the NDS simple cells (Figure 17(B)(c)) and MDCs (Figure 17(B)(f)) are shown in Sections **S2.1** and **S2.2** of Supplementary materials, respectively. Next, using these tables, the procedure for calculating the series of cell responses in Figure 17(B) is shown in Section **S3.1** of Supplementary materials. These procedures were also used in previous simulations (Kawakami & Okamoto, 1996a, 1995a, 1992d; Kawakami, 1996b; Kawakami et al., 2003, 2000, 1992b).

Next, the procedure for calculating the neural network (i.e. for calculating the connection table representing this network), which connects each MDC in Figure 17(B)(f) and the SDC column in Figure 17(C), is shown in Section **S2.4** of Supplementary materials. Using this table, the procedure for calculating the cell responses of the SDC column (Figure 17(C)) is shown in Section **S3.3** of Supplementary materials. These procedures were also used in previous simulations (Kawakami et al., 2003, 2000).

All these procedures were installed on a computer to simulate the responses of all cells in Figures 17(B and C). The simulation results are shown in Figure 19, which are quoted from a report (Okamoto, 2006) that was jointly simulated using the Supplementary materials. It is shown below that the normalized shortest-distance to a plane and its 3D orientation can be detected correctly. Figures 17(B and C) are consisted of the about 5 million cells and about 120 million neural connections between them.

A case approaching a perpendicular plane 1 m away (on which random dots were drawn) was simulated: the approaching speed is 1 m/s. Figure 19(A) shows an image in which this plane is reflected on the retina of the eyeball. The normalized shortest-distance d_C_ to the plane is calculated as 1m/(1m/s×50ms), i.e. 20.

Although Figure 19(B) was intended to show the cross sections at three heights (centered at height d=20) of the SDC column, there was no d=20 in this discrete array in the height direction. Therefore, the cross-sections of d=18±5 were shown in the (B) centered on d=18, which was closest to d=20: it was 90±25 cm when converted to the shortest distance D. Each dot in these cross sections represents a SDC.

In the height direction, the cross section at d=18 had the largest response, and within this cross-section, a central SDC (indicated by a green dot) had the largest response. This indicates that this SDC detected the normalized shortest-distance (i.e. d=20) and orientation (i.e. n=the center of the visual field O) of the given plane. The shortest-distance detection can identify 90±25 cm, but the orientation detection is broad. The cause of the orientation detection being broad is due to the red ring-region of Figure 26(A)(ii) described later, which is caused from the dR/d τ_2D_ in equation(4.3-4) (see Section **4.3.3.2**(1)).

The above simulation has confirmed that the neural network of Figure 17 can detect the normalized shortest-distance to the plane and its 3D orientation.

### 3.4. How is this cell model used?

In Section **2.4**, it was mentioned that the normalized time-to-contact t is essential to avoid the plane such as a wall. Let us consider how it would be useful to add the normalized shortest-distance d to this.

When walking toward the wall (Figure 20(A)), the time to reach the wall (such as t_1_ and t_2_) is detected one after another as the height coordinate of the SDC column (Figure 20(B)). When it approaches the critical time t_critical_ that is set, you change the direction of movement so that it is perpendicular to the 3D orientation n of the wall detected at the same time, and then continue to walk parallel to the wall. At this time, the cells in the SDC column (n,t) fire in a trajectory as shown in Figure 20(B) (Kawakami et al., 2002). This is the avoidance of the wall using the normalized time-to-contact t, described in Section **2.4**.

Let us consider what happens when the normalized shortest-distance d is added to this. When walking toward the wall (see (A)), a shortest distance to the wall is detected one after another as the height coordinates of the SDC column (n,d) in the (C), such as d1, d2, and d_critical_, but this distance does not directly lead to avoidance behavior. This shortest distance d is useful when you continue to walk parallel to the wall after changing the movement direction. As shown in the (A), you can continue to walk in parallel while keeping the shortest distance d to the wall at a constant distance d_parallel_. At this time, the cell in the SDC column (n,d) draws a trajectory like the (C) and fire.

In this way, the normalized shortest distance d is important for walking parallel to the wall. In addition, the pilot in Figure 1 needs this distance d_parallel_ to keep flying while maintaining a constant distance to the runway.

Here, note the following. According to relationship D=(Vt_d_)d of equation(3.0-2), the actual shortest distance D changes in proportion to the speed V even if the normalized shortest distance d is kept constant: this normalized distance d may be detected by cells in the cerebrum. Thus, at high speed you walk apart at a large distance D, but at low speed you walk close at a small distance D. This is reasonable as follows. Since the risk is high at a high speed, it is reasonable to walk apart. On the other hand, since the risk is low at a low speed, it is reasonable to walk close. It is thought that we are doing the same action as above.

In addition, when driving a car, the scene appears to shrink at high speeds and expand at low speeds. This appearance may be considered to be related to the property of this normalized shortest-distance d (i.e. to be related to d=D/(Vt_d_)). In other words, I think that our brain detects the normalized shortest-distances d instead of the shortest distances D, so the scene looks like it shrinks or expands depending on speeds V.

## 4. Discussion

### 4.1. Lee’s detection of time-to-contact is a special case of this model

#### 4.1.1. Lee’s detection of time-to-contact

Lee was the first to propose an equation for detecting the time-to-contact (Lee, 1976, 1980). Let us explain the equation with reference to Figure 21(A). This equation detects the time-to-contact to a point instead of that to a plane in Figure 1.

**Figure 21.**
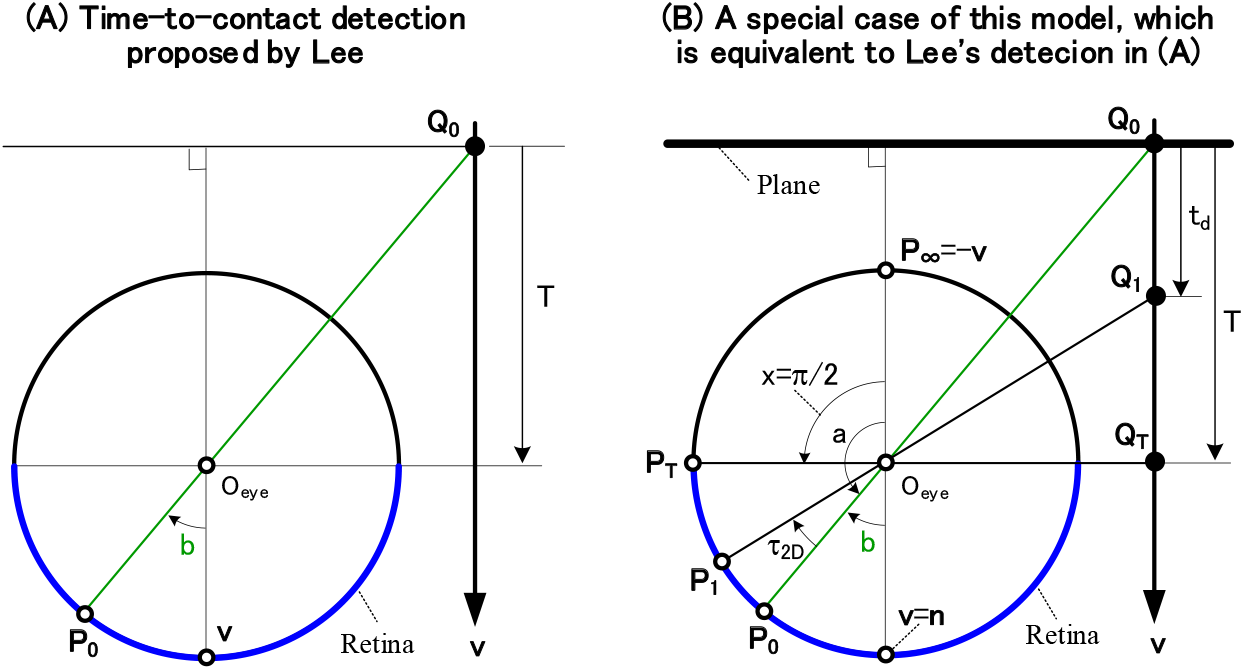
Lee’s detection of the time-to-contact is a special case of this model. (A) A figure for explaining Lee’s detection is shown. There is a point Q_0_ in space, which is reflected on the eyeball as a point P_0_. A cross section of the eyeball passing both this point and the eyeball is shown. This Q_0_ moves in the direction v and reaches just beside the eyeball center O_eye_ at a time T. Using this figure, equation(4.1-1) for obtaining the time-to-contact is explained in Section **4.1.1**: this equation was proposed by Lee (1976, 1980). (B) This figure is shown to prove that Lee’s detection is a special case of this model in Figure 9. Considering the case where a plane moves in direction v, we show a cross section of the plane and the eyeball. A point Q_0_ on the plane moves to Q_1_ in the unit time t_d_, and moves to position Q_T_ just beside the eyeball center O_eye_ in time T. These Q_0_, Q_1_, and Q_T_ are reflected on the eyeball as P_0_, P_1_, and P_T_. Using this figure, Section **4.1.2** proves that Lee’s equation of equation (4.1-1) is equivalent to a special case of this model.

There is a point Q_0_ in space at a current time, which is reflected as a point P_0_ on the eyeball. Figure 21(A) shows a cross section of this point and the eyeball. This Q_0_ moves toward direction v and reaches just beside the eyeball center O_eye_ at a time T. Lee proposed that this time-to-contact T is determined by the following equation (Lee, 1976, 1980).

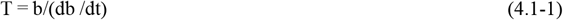

Here, the variable b used above is the central angle between the movement direction v and the point P_0_, and db/dt is the time derivative of b: note that t is not the normalized time-to-contact of equation(1.0-3), but T. This is Lee’s equation for detecting the time-to-contact. However, this equation is an approximation that holds when b is small: b in Figure 21(A) is drawn large for easy viewing.

Although Lee derived equation(4.1-1) using the time derivative, Appendix **8** shows that it can be derived geometrically without using this derivative.

#### 4.1.2. Derivation of Lee’s detection from this model

It is shown as follows that equation(4.1-1) can be derived as a special case of this model in Figure 9.

Figure 21(B) shows a cross-section of a plane and the eyeball: the plane moves in the direction v. Assume that a point Q_0_ on the plane moves to Q_1_ at the unit time t_d_ and moves to Q_T_ at a time T: Q_T_ is just beside the eyeball center O_eye_. These Q_0_, Q_1_, and Q_T_ are reflected on the eyeball as P_0_, P_1_, and P_T_. The central angle (a,x, and τ_2D_) is the same as in Figure 8(B), and the central angle b of Figure 21(A) is added to it.

Although this model of Figure 9 can detect planes with arbitrary orientation, consider a special case where the plane is perpendicular and its orientation n is equal to the movement direction v.

Equation(4.1-2) is obtained by combining equation(1.0-3) with the cross-ratio transform of equation(1.0-6): for clarity of expression, represent T–T_0_ in equation(1.0-3) as T. Since x=π/2 due to the above special case, equation(4.1-3) is obtained. Since a=π+b based on Figure 21(B), equation(4.1-4) is obtained. Since b≫τ_2D_, equation(4.1-5) is obtained. Since b and τ_2D_ are small, equation(4.1-6) is obtained. Finally, since t_d_ is the time interval between P_0_ and P_1_ (see equation(1.0-2)) and lim_td→0_(τ_2D_/t_d_)=db/dt, the same equation(4.1-7) as Lee has been obtained.

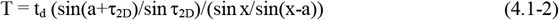

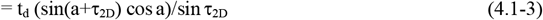

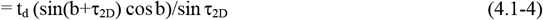

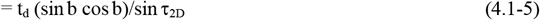

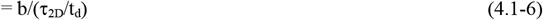

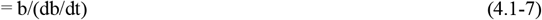

In this way, Lee’s equation (i.e. equation(4.1-1)) is equivalent to a special case of the cross-ratio transform (i.e. equation(4.1-2)), and also to a special case of of this model. Therefore, this equivalence has revealed that Lee’s equation calculates the time-to-contact of a point object near the center of the visual field, rather than a plane.

### 4.2. Comparison with Ethology, Psychology, and Physiology

#### 4.2.1. Comparison with Ethology and Psychology

Lee reported ethological experiments on gannets (Lee & Reddish, 1981) to verify his above proposal that the the time-to-contact T is detected by equation(4.1-1), as follows. The gannet dives into the surface of the sea at high speed in order to prey on fish, but just before that, it folds its wings so as not to hurt it. When this dive was photographed with a high-speed camera and the folding start time was measured, this time was almost constant (average 820 ms) regardless of the speed. Assuming that the gannets start folding their wings at a fixed time before contacting the sea, equation (4.1-1) that detects this time (i.e. the time-to-contact) regardless of speed is reasonable.

In addition, ethological experiments on flies were reported (Wagner, 1982), as follows. The time at which the flies begin to decelerate for landing was measured with a high-speed camera, and the times measured were almost constant regardless of speed. Assuming that the flies start decelerating at a fixed time before the landing, equation (4.1-1) that detects this time (i.e. the time-to-contact) regardless of speed is reasonable.

Since Lee’s equation above is equivalent to the special case of the cell model in Figure 9(C) (see Section **4.1.2**), this model also can explain these ethological experiments. Considering further, since the sea surface into which gannets dive is a planar surface (i.e. a plane), this cell model that can detect the time-to-contact to that sea surface seems more appropriate than Lee’s equation.

These ethological experiments on the gannets and flies suggest that there are the modeled SDCs of Figures 9(C) in their brains.

The above has described ethological experiments regarding the detection of time-to-contacts for points rather than for planes. However, no ethological experiments for the planes, which are more difficult to detect than the points, have been conducted. In this way, detecting the time-to-contacts of the planes seems to remain at a stage of psychological consideration (Gibson, 1950) that precedes ethological experiments. Gibson (1950) psychologically considered and discovered that the time-to-contact to a plane was detected based on the magnitude of each local motion that constituted the OFP (Figure 2), and that its 3D orientation was detected based on the shape of an OFP. However, it seems that this discovery was not developed, not being carried over by experiments in ethology and psychology.

It is surprising that the series of modeled cells in Figure 9 enables this psychological discovery of Gibson to be explained neurophysiologically at the level of the neural connection of which cell is connected to which cell (see Section **2.2**). This neurophysiological explanation is, to my knowledge, the first time except for our reports (Kawakami et al., 2003, 2000).

#### 4.2.2. Comparison with Physiology

##### 4.2.2.1. Pigeons

In nucleus rotundus of pigeons, cells that detect time-to-contacts were reported (Wang & Frost, 1992; Sun & Frost, 1998). They found that the cells responded strongly to light stimuli with a fixed time-to-contact (about 1 second) regardless of speed. This result is consistent with Lee’s equation(4.1-1) that detects this time regardless of the speed. The modeled SDCs in Figure 9(C), which detect the times regardless of the speed, also can explain these experiments.

In addition, cells that detect the shortest distance to a plane were reported in thalamic neurons of pigeons (Liu et al., 2008). The modeled SDCs in Figure 17(C), which detect the distances, can explain this experiment. Note that Lee (1976, 1980) above did not describe this distance detection.

##### 4.2.2.2. Monkeys

Cells in the frontal eye field (FEF) of monkeys that detect the time-to-contacts to planes regardless of the speed were reported (Fujiwara et al., 2014). The modeled SDCs in Figure 9(C), which detect these time-to-contacts regardless of the speed, can explain this experiment neurophysiologically. In addition, cells in area MST of the monkeys that detect the shortest distances to planes regardless of the speed were reported (Fujiwara et al., 2014). The modeled SDCs in Figure 17(C), which detect these shortest distances regardless of the speed, can explain this experiment neurophysiologically.

In addition, cells in area MST of monkeys that responded selectively to the OFP such as Figure 2 were reported (Saito et al., 1986). The series of modeled cells in Figure 9, which has the same selectivity for the OFP as in the above, can explain this experiment neurophysiologically.

In collaboration with Saito and Hida of Saito Laboratory in Tamagawa University, we conducted a preliminary verification experiment of this model in area MST of monkeys (see Okamoto (2006) who collaborated with us, for an outline). The results are shown as follows.

First, as shown in Figure 22, time responses of a modeled SDC to the planar stimulus was simulated on a computer using a series of cells in Figure 9. Figure 22(A) shows a plane used: the plane with random dots on it is tilted by a 3D orientation n=(0°,30°), and is approached vertically to the eyeball, where v (i.e. the movement direction) = O (i.e. the center of the visual field). The horizontal axis of Figure 22(B) shows the time course of the stimulus movement, as follows. At 6 seconds of time-to-contact, this plane that was stopped was presented for 1 second as a stimulus. Then, at 5 seconds, the plane started to move and was presented until the plane reached the eyeball.

**Figure 22.**
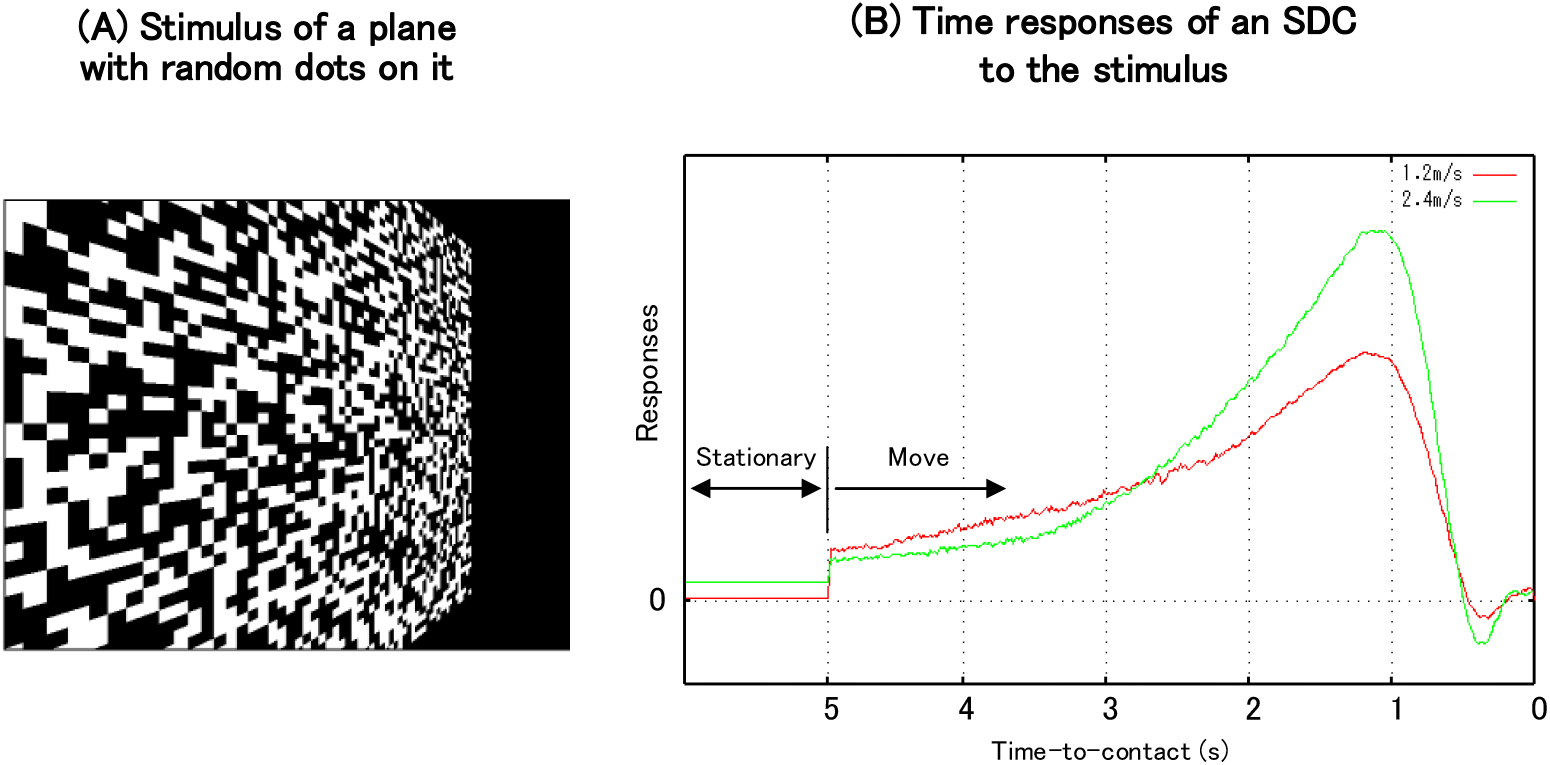
Time responses of a SDC to a stimulus. Time responses of the modeled SDC in Figure 9(C) to the stimulus were simulated. (A)The stimulus of a plane on which random dots are drawn was used. Its 3D orientation n is (0°,30°). (B) Time responses of a SDC to the stimulus were simulated. The cell response increased with shorter time-to-contacts, reaching a maximum response at 1.2 seconds. Even if the speed was changed from 1.2m/s to 2.4m/s, this time of 1.2 seconds to reach the maximum response did not change.

Figure 22(B) shows time responses of a SDC (n=(0°,30°), t=24) to the stimulus above. Responses of the SDC increased with decreasing the time-to-contact, and reached a maximum response at 1.2 seconds (i.e. 24×50 ms): the time-to-contact to reach this maximum response corresponds to the height t_C_ in Figure 9(C). Changing the speed from 1.2 m/s to 2.4 m/s does not change this time of 1.2 seconds. This independence on speed is due to the fact that the velocity (or the speed) V contained in equation(A1-4) in Appendix **1** was canceled in the numerator and denominator. This independence is an important property and is compared with the following experiments of a monkey. Figure 23 shows the experimental results for a cell in area MST of a monkey. The planar stimulus used is that with the same random dots as in Figure 22(A): the movement direction is the same vertically, but the 3D orientation of the plane is n = (0°,0°), which is different from Figure 22(A). The time course of the stimulus movement (i.e. the horizontal axis of Figure 23) is the same as that of Figure 22(B). That is, at 6 seconds of time-to-contact, this plane that was stopped was presented for 1 second as a stimulus. Then, at 5 seconds, the plane started to move and was presented until the plane reached the eyeball: however, in this experiment, after the stimulus arrived, the stimulus was hidden and the measurement continued for 1 second. A microelectrode was inserted in the vicinity of the cell in area MST, and its responses to the stimulus above were extracellularly recorded. This stimulation cycle was repeated 10 times, and the responses were added to form the vertical axis of Figure 23.

**Figure 23.**
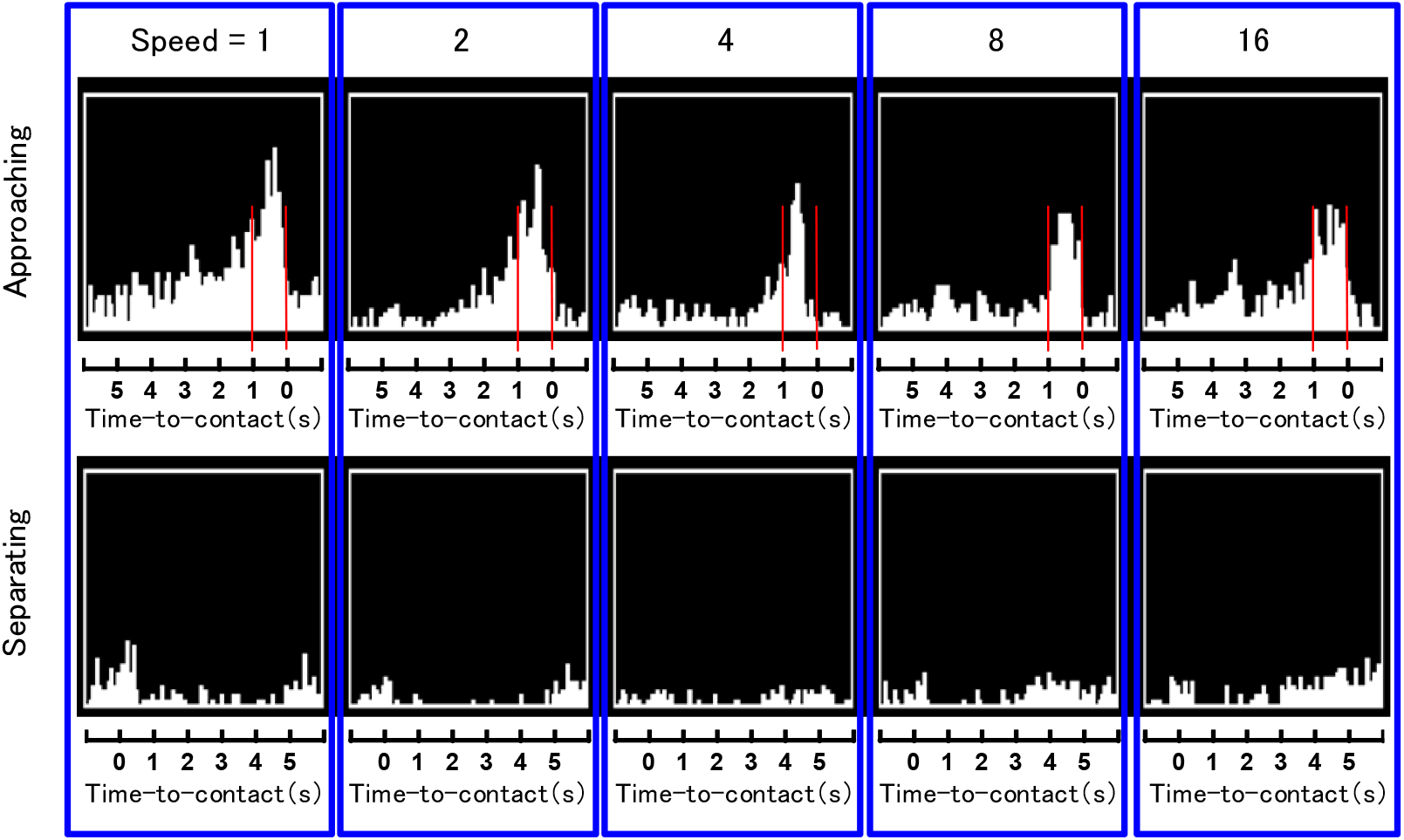
Time responses of a cell in area MST of a monkey to the stimulus. The results of electrophysiological experiments on a cell in area MST of a monkey are shown. The planar stimulus used is almost the same as in Figure 22(A). The time responses of the cell to the approaching movement are shown in the upper part. At speed=1, the cell responses increased with shorter time-to-contacts, reaching a maximum response at approximately 0.7 seconds. Even if the speed was increased by 2, 4, 8, and 16 times, the time to reach the maximum response remained almost the same. These are consistent with the simulation results for the modeled SDC of Figure 22(B). The time responses to the separating movement are shown in the lower part. Not only was the response significantly reduced, but the response was suppressed by presenting the stimulus.

The time responses of the cell to the approaching movement, in which the planar stimulus approachs the monkey, are shown in the upper part of Figure 23. At speed=1, the cell response increases with shorter time-to-contact, reaching a maximum response at about 0.7 seconds. This is the same as the time response of the modeled SDC in Figure 22(B). Even if the speed is increased by 2, 4, 8, and 16 times, the time of maximum response does not change much. This speed independence is also the same as in the time response of the modeled SDC in Figure 22(B).

Next, the time responses of the cell to separating movement, in which the planar stimulus moves away from the monkey, are shown in the lower part of Figure 23. Not only were the responses significantly reduced, but the responses were suppressed by presenting the stimulus. Note that this significant decrease due to the separating movement was not described in Figure 22(B), but is the same as that of the modeled SDC previously reported (see Figure 3.19 in Okamoto (2006)).

Thus, it has been confirmed that the modeled SDCs, which detect the time-to-contacts regardless of speed, exist in area MST. These experimental results (Figure 23) are consistent with the responses of the modeled SDC (Figure 22(B)), and thus supported the validity of the series of modeld cells in Figure 9.

### 4.3. How to use the forward and backward methods properly according to their characteristics

There are two methods for creating the SDC neural networks of Figure 9(C), which connect MDCs in Figure 9(B)(f) and the SDC column in Figure 9(C). These methods are named the forward (Figure 24(A)) and backward (Figure 24(B)) methods: only retinal cells ((ii)(a)) and MDCs ((ii)(f)) are shown, and other cells in Figure 9(B) are omitted for viewability. The forward method is how to create forwardly a SDC neural network from each MDC to the SDC column, and the backward method is how to create backwardly a network from each SDC of the column to the MDCs.

**Figure 24.**
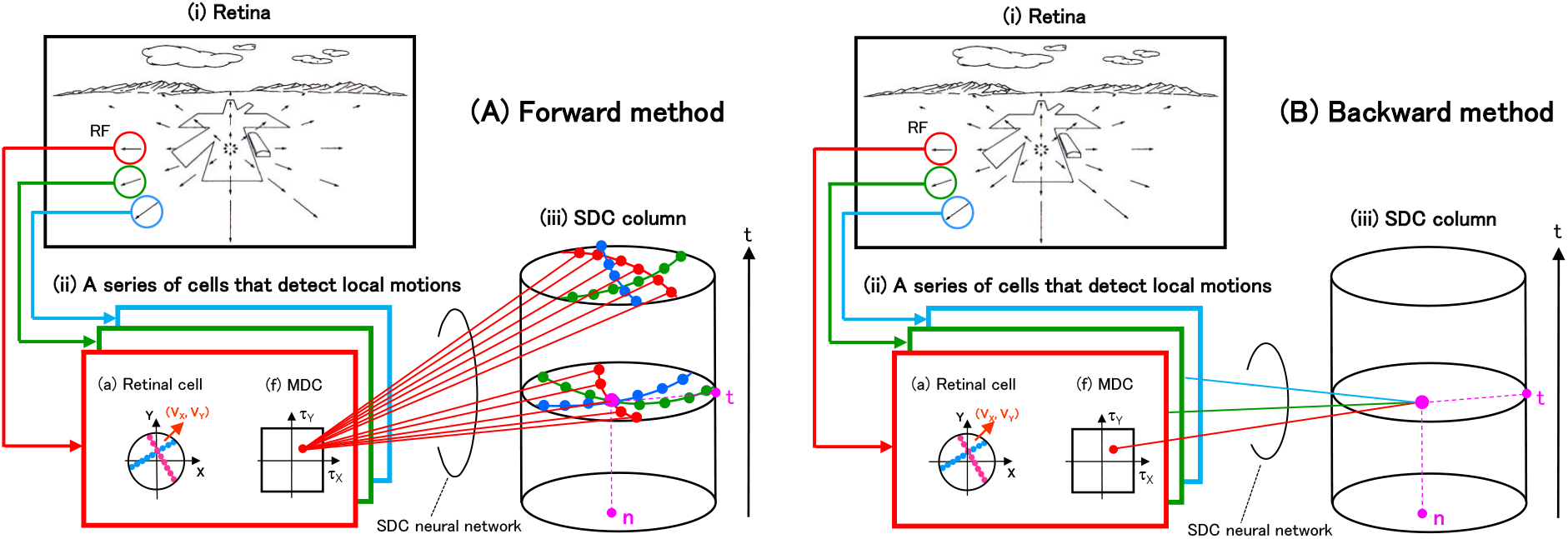
Forward and backward methods. There are two types of methods (i.e. the forward and backward methods) for creating of the SDC neural network that connects the MDCs in the Figure 9(B)(f) and the SDC column in the Figure 9(C). Note that in the (A)(ii) and (B)(ii) of Figure 24, only the retinal cells in the (a) and the MDCs in the (f) are shown, and the other cells in Figure 9(B) are omitted for ease of viewing. (A) A forward method is shown to create a SDC neural network that connects each MDC to the SDC column. (B) A backward method is shown to create a SDC neural network that connects each SDC in the column to the MDCs.

**Figure 25.**
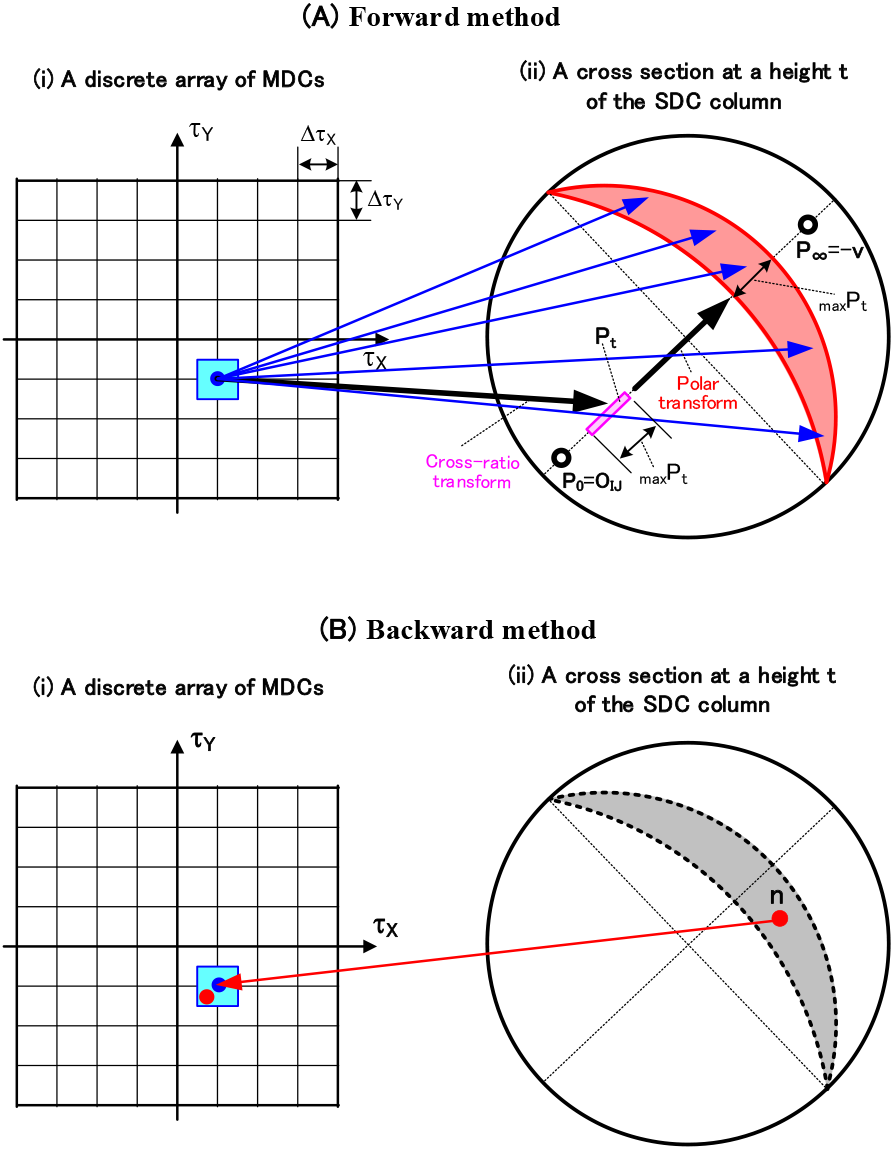
Influence of a discrete MDC array on a forward and backward SDC-neural networks that detect the normalized time-to-contact and 3D orientation. Consider a case where MDCs in Figures 24(A)(ii)(f) and 24(B)(ii)(f) are arranged discretely. (A) A forward SDC neural network is described as follows. Consider how a MDC at each lattice point indicated by a blue dot in (i) is connected to SDCs within a cross section of the column in (ii). Since this MDC represents all local motions {(τ_X_, τ_Y_)} within a blue square, this square is converted, by the cross-ratio transform of equations(2.2-5 and 8), to all positions P_t_ within a pink elongated rectangle in (ii). Then, this rectangle is converted into SDCs within a red crescent of (ii) by the polar transform of equation(2.2-10). Thus, a MDC on each blue lattice point in (i) is connected to all SDCs within this crescent in (ii). These connections are indicated as blue straight lines. Therefore, these straight lines represent the forward SDC neural network. (B) A backward SDC neural network is described as follows. Consider how each SDC indicated by the red dot in (ii) is connected to MDCs in (i). As a candidate for connecting this SDC n, a MDC indicated by the red dot in (i) is calculated using Section **A9.1** in Appendix **9**. However, since this MDC is not on any lattice point, a MDC on the blue lattice point closest to this dot is selected. The MDC on this lattice point is connected to the red SDC n in (ii), and the connection is indicated as a red straight line. This straight line represents the backward SDC neural network.

Brief overviews are provided below to highlight the content of the following sections. First, Section **4.3.1** will show that the SDC neural network obtained by the forward or backward method is expressed by mathematical equations. Next, Section **4.3.2** will show that when MDCs are continuously arranged, the networks obtained by both methods is the same. Finally, Section **4.3.3** will show that when MDCs are arranged discretely, both methods are different from each other and thus are necessary to be used properly according to their characteristics. The main points of this necessity are as follows. First, in terms of computer simulation, the backward method is easier than the forward method because there is a difference in complexity in the procedures of both methods. Second, in terms of ease for explaining mathematical structure of the neural network, the forward method can explain its structure, but the backward method cannot explain its structure. In this way, since there are advantages and disadvantages between both methods, it is necessary to use them appropriately as follows. The forward method is suitable for explaining the mathematical structure of neural networks, and the backward method is suitable for simulating the network (Sections **4.3.3.1**(3) and **4.3.3.2**(3)).

#### 4.3.1. Representation of the SDC neural network by mathematical equations

Let us express the forward and backward SDC-neural networks by mathematical equations.

1. Normalized time-to-contact The forward SDC-neural network of Figure 24(A) was expressed by mathematical equations in Section **2.2**. The backward SDC-neural network of Figure 24(B) was expressed by mathematical equations in Section **A9.1** of Appendix **9**.
2. Normalized shortest distance

In order to consider the normalized shortest distance, let us replace the height axis t of the SDC column (Figures 24(A)(iii) and 24(B)(iii)) for the normalized time-to-contact with the height axis d of the SDC column for the normalized shortest distance.

When such replacement has been performed, the forward SDC-neural network of Figure 24(A) was expressed by mathematical equations in Section **3.2**. The backward SDC-neural network of Figure 24(B) was expressed by mathematical equations in Section A**9.2** of Appendix **9**.

#### 4.3.2. Case where MDCs are arranged continuously

Consider the case where the MDCs in Figures 24(A)(ii)(f) and 24(B)(ii)(f) are continuously arranged. That is, the array (τ_X_, τ_Y_) of MDCs has a continuous address of the real number.

(1) Normalized time-to-contact The forward SDC network in Section **2.2** is the same as the backward SDC network in Section **A9.1** of Appendix **9**, although their equations are different. The sameness of these SDC networks has been confirmed by computer simulations.
(2) Normalized shortest distance The forward SDC network in Section **3.2** is the same as the backward SDC network in Section **A9.2** of Appendix **9**, although their equations are different. The sameness of these SDC networks was confirmed by computer simulations.

#### 4.3.3. Case where MDCs are arranged discretely

Consider the case where the MDCs in Figures 24(A)(ii)(f) and 24(B)(ii)(f) are discretely arranged. That is, the array (τ_X_, τ_Y_) of MDCs has a discrete address of the integer: this discrete array is shown in Figures 25(A)(i) and 26(A)(i). The SDCs are also arranged discretely, but the discrete effect is small and thus is not considered.

##### 4.3.3.1. Normalized time-to-contact

In Figure 25, the (i) shows a discrete array of MDCs, and the (ii) shows the cross section of the SDC column at a height t. The MDCs are discretely arranged, and each MDC exists at a lattice point.

(1) Forward method The forward method is shown in Figure 25(A). Consider how a MDC at ach lattice point ((A)(i)) indicated by the blue dot is connected to the SDCs ((A)(ii)) in the cross section at a height t. Since this MDC ((A)(i)) represents the all local motions {(τ_X_,τ_Y_)} in the blue square, the MDC is converted to all positions P_t_ in the pink elongated rectangle ((A)(ii)) by the cross-ratio transform of equations(2.2-5 and 8): {(τ_X_,τ_Y_)} represents a set of (τ_X_,τ_Y_). This elongation becomes more pronounced as the coordinate τ_2D_ of the lattice point becomes smaller (i.e. as the lattice point becomes closer to the origin). Finally, this rectangle is converted into a red crescent of the (A)(ii) by the polar transform of equation (2.2-10). Thus, the MDC at the blue lattice point is connected to all SDCs within this crescent: their connections are indicated by blue straight lines. These straight lines represent the forward SDC-neural network. The pink great circle g_Pt_ in Figure 11(A) has increased in width to become this crescent. The width _max_P_t_ of the crescent can be close to 90 degrees. Here, the width _max_P_t_ of this crescent, which is equal to the length _max_P_t_ of the rectangle, can be calculated by the following equation.

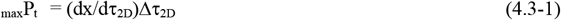 Here, Δτ_2D_ is calculated by the following equation using the resolution (Δτ_X_,Δ τ_Y_) of the MDC (see (A)(i)).

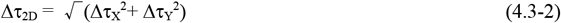 In addition, this derivative coefficient dx/d τ_2D_ is calculated as equation(4.3-3) that is obtained by differentiating equation(2.2-5) with τ_2D_: the variable a used was calculated by equation(2.2-6).

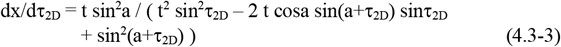
(2) Backward method The backward method is shown in Figure 25(B). Consider how each SDC in the (B)(ii), indicated by a red dot, is connected to MDCs in the (B)(i). As a candidate for connecting this SDC n, a local motion (τ_X_, τ_Y_) indicated by a red dot ((B)(i)) within the blue square is calculated using mathematical equations described in Section **A9.1** of Appendix **9**. However, since this motion (τ_X_, τ_Y_) is not a lattice point, the MDC of a blue lattice point closest to this motion (τ_X_, τ_Y_) is selected. The MDC of this lattice point is connected to the SDC n of the (B)(ii), and the connection is indicated by a red straight line. This line is the backward SDC-neural network.
(3) How to use both methods properly according to their characteristics The backward SDC network with one connection was shown in Figure 25(B). Therefore, you might think it is significantly smaller than the forward SDC network in the (A), but they are actually about the same as follows. Since one red SDC n was extracted in the (B)(ii), the backward SDC network had one connection, but in fact, all SDCs within the gray crescent in the (B)(ii) are connected to the MDC of this blue lattice point in the (B)(i). Why this is so is explained as follows. As described in subsection (2) above, as a candidate for connecting the red SDC within the crescent in (B)(ii), a local motion (τ_X_, τ_Y_) within the blue square of (B)(i) was calculated using mathematical equations in Section **A9.1** of Appendix **9**: this local motion was indicated by a red dot. Similarly, as a candidate for connecting every SDC within this crescent, a local motion (τ_X_, τ_Y_) within the blue square of (B)(i) can be calculated. However, since these local motions (τ_X_, τ_Y_) are not on any lattice point, the MDC of a blue lattice point closest to these local motions is selected. Therefore, all SDCs within the crescent of (B)(ii) are connected to the MDC at this blue lattice point of (B)(i). Therefore, the backward network thus obtained is substantially the same as the forward network that is indicated by blue straight lines in the (A): this sameness was confirmed by creating a simulator on a computer based on Supplementary materials. In this way, the neural networks by both methods are nearly identical. However, the complexity of the procedure required for the computer simulation differs greatly as follows. In the forward method of the (A), the blue square in the (A)(i) is converted to the pink enongated rectangle of P_t_ ((A)(ii)) by the cross-ratio transform of equations(2.2-5 and 8). Then, this rectangle is converted into the red crescent ((A)(ii)) by the polar transform of equation(2.2-10). Thus, this procedure is very complicated. On the other hand, in the backward method of (B), a MDC candidate connected to each SDC n in the (B)(ii) is simply calculated using mathematical equations described in Section **A9.1** of Appendix **9**. In this way, the procedure for the backward method is significantly simpler than that for the forward method. In terms of explaining the mathematical structure of the neural network, these two methods differ greatly. That is, in the forward method of the (A), the two transforms in Section **2.1.4** (i.e. the cross-ratio and polar transforms) explain its mathematical structure. On the other hand, in the backward method of the (B), the procedure is clear in Section **A9.1** of Appendix 9, but this method cannot explain its mathematical structure. Thus, the forward method explains its mathematical structure, but the backward method cannot explain its structure. Based on the advantages and disadvantages of these two methods, it is necessary to use them properly according to their characteristics, as follows. That is, the forward method is suitable for explaining the mathematical structure of the neural network, and the backward method, which is simple in procedure, is suitable for simulation.

##### 4.3.3.2. Normalized shortest distance

In Figure 26, the (i) shows a discrete array of MDCs, and the (ii) shows the cross section of the SDC column at a height d. The MDCs are discretely arranged, and each MDC exists at a lattice point.

(1) Forward method The forward method is shown in Figure 26(A). Consider how a MDC at each lattice point ((A)(i)) indicated by the blue dot is connected to the SDCs ((A)(ii)) in the cross section at a height d. Since this MDC ((A)(i)) represents the all local motions {(τ_X_, τ_Y_)} in the blue square, the MDC is converted to all SDCs within a red ring ((A)(ii)) with a width _max_R using the small-circle transform of equations(3.2-5 and 8). This width increases as the coordinate τ_2D_ of the lattice point becomes smaller (i.e. as the lattice point becomes closer to the origin). Thus, the MDC at the blue lattice point is connected to all SDCs within this ring: their connections are indicated by blue straight lines. These straight lines represent the forward SDC-neural network. The pink small circle in Figure 16(A) has increased in the width _max_R to become this ring. The width _max_R of this ring can be calculated by the following equation using the derivative coefficient dR/d τ_2D_.

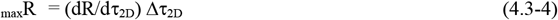 Here, this dR/d τ_2D_ is obtained by differentiating equation(3.2-5) with τ_2D_: Δτ_2D_ is calculated by equation(4.3-2).
(2) Backward method The backward method is shown in Figure 26(B). Consider how each SDC ((B)(ii)), indicated by a red dot, is connected to MDCs ((B)(i)). As a candidate for connecting this SDC n, a local motion (τ_X_, τ_Y_) indicated by a red dot ((B)(i)) within the blue square is calculated using mathematical equations described in Section **A9.2** of Appendix **9**. However, since this motion (τ_X_, τ_Y_) is not a lattice point, the MDC of a blue lattice point closest to this motion (τ_X_, τ_Y_) is selected. The MDC at this lattice point is connected to the SDC n of the (B)(ii), and the connection is indicated by a red straight line. This line represents the backward SDC-neural network.
(3) How to use both methods properly according to their characteristics The backward SDC network with one connection was shown in Figure 26(B). Therefore, you might think it is significantly smaller than the forward SDC network in the (A), but they are actually about the same as follows. Since one red SDC n was extracted in the (B)(ii), the backward SDC network had one connection, but in fact, all SDCs within the gray ring in the (B)(ii) are connected to the MDC of this blue lattice point in the (B)(i).

Why this is so is explained as follows. As described in subsection (2) above, as a candidate for connecting the red SDC within the ring in (B)(ii), a local motion (τ_X_, τ_Y_) within the blue square of (B)(i) was calculated using mathematical equations in Section **A9.2** of Appendix **9**: this local motion is indicated by a red dot. Similarly, as a candidate for connecting every SDC within this ring, a local motion (τ_X_, τ_Y_) within the blue square of (B)(i) can be calculated. However, since these local motions (τ_X_, τ_Y_) are not on any lattice point, the MDC of a blue lattice point closest to these local motions is selected. Therefore, all SDCs within the ring of (B)(ii) are connected to the MDC at this blue lattice point of (B)(i). Therefore, the backward network thus obtained is substantially the same as the forward network that is indicated by blue straight lines in the (A): this sameness was confirmed by creating a simulator on a computer based on Supplementary materials.

In this way, the neural networks by both methods are nearly identical. However, the complexity of the procedure required for the computer simulation differs greatly as follows. In the forward method of the (A), the blue square in the (A)(i) is converted to a red ring ((A)(ii)) with the width _max_R. Thus, this procedure is complicated. On the other hand, in the backward method ((B)), a MDC candidate connected to each SDC n in the (B)(ii) is simply calculated using mathematical equations described in Section **A9.2** of Appendix **9**. In this way, the procedure for the backward method is significantly simpler than that for the forward method.

In terms of explaining the mathematical structure of the neural network, these two methods differ greatly. That is, in the forward method of the (A), the mathematical structure of the SDC network can be explained by the small-circle transform. On the other hand, in the backward method of the (B), the procedure is clear in Section **A9.2** of Appendix **9**, but this method cannot explain its mathematical structure. Thus, the forward method can explain its mathematical structure, but the backward method cannot explain its structure.

Based on the advantages and disadvantages of these two methods, it is necessary to use them properly according to their characteristics, as follows. That is, the forward method is suitable for explaining the mathematical structure of the neural network, and the backward method, which is simple in procedure, is suitable for simulation.

### 4.2. A cell model for detecting the shortest distance to a plane and its 3D orientation by binocular stereo

#### 4.4.1. Binocular stereo is the same as motion stereo at each moment

It is shown in Figure 27 that the parameters for binocular stereo are the same as those for motion stereo at each moment. Specifically, the (A) shows a cross section of binocular stereo (i.e. a cross section that passes between a green dot (i.e. an object) and the centers of both eyes), and this dot is reflected at a position P_L_ on the left eye and a position P_R_ on the right eye. An angle at which the dot is seen with both eyes is the binocular parallax σ_2D_. Also, the (B) shows a cross section of motion stereo (i.e. a cross section that passes between a green dot and the centers of the eyeballs at the current and next times), and this dot is reflected at the position P_0_ on a current-time eyeball and the position P_1_ on a next-time eyeball. An angle at which the dot is seen by the eyeballs at both times is the motion parallax (i.e. the local motion) τ_2D_. Note that the plane and the distance D_Bino_ shown in the (A) will be used in Section **4.4.3**.

**Figure 26.**
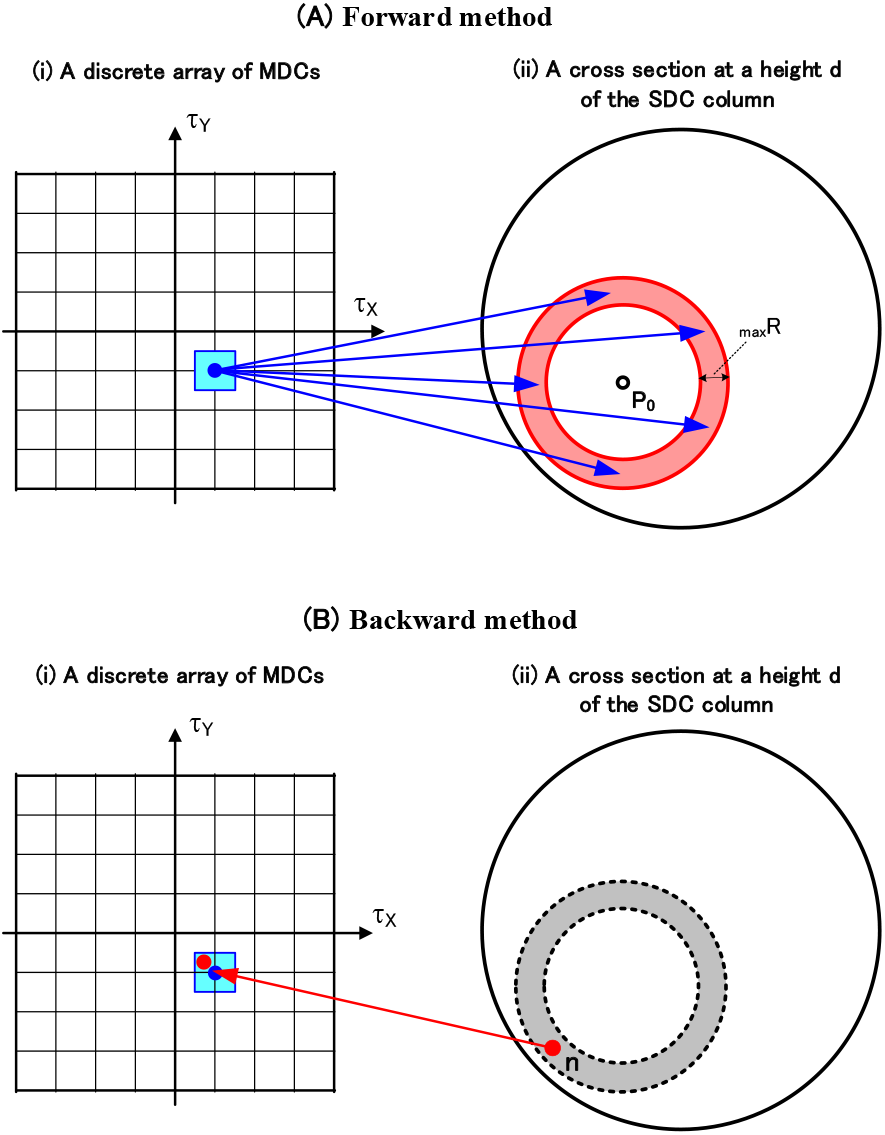
Influence of a discrete MDC array on a forward and backward SDC-neural networks that detect the normalized shortest-distance and 3D orientation. Consider a case where MDCs in Figures 24(A)(ii)(f) and 24(B)(ii)(f) are arranged discretely. (A) A forward SDC neural network is described as follows. Consider how a MDC at each lattice point indicated by a blue dot in (i) is connected to SDCs within a cross section of the column in (ii). Since this MDC represents all local motions {(τ_X_, τ_Y_)} within a blue square in (i), by the small-circle transform of equations(3.2-5 and 8), the square is converted into a red ring ((ii)) with a width _max_R centered on P_0_: this P_0_ is a position at a current time (see Figure 16(A)). Therefore, a MDC on each blue lattice point in (i) is connected to all SDCs within this ring. These connections are indicated as blue straight lines. These straight lines represent the forward SDC neural network. (B) A backward SDC neural network is described as follows. Consider how each SDC indicated by the red dot in (ii) is connected to MDCs in (i). As a candidate for connecting this SDC n, a MDC indicated by the red dot in (i) is calculated using Section **A9.2** in Appendix **9**. However, since this MDC is not on any lattice point, a MDC on the blue lattice point closest to this dot is selected. The MDC on this lattice point is connected to the red SDC n in (ii), and the connection is indicated as a red straight line. This straight line represents the backward SDC neural network.

**Figure 27.**
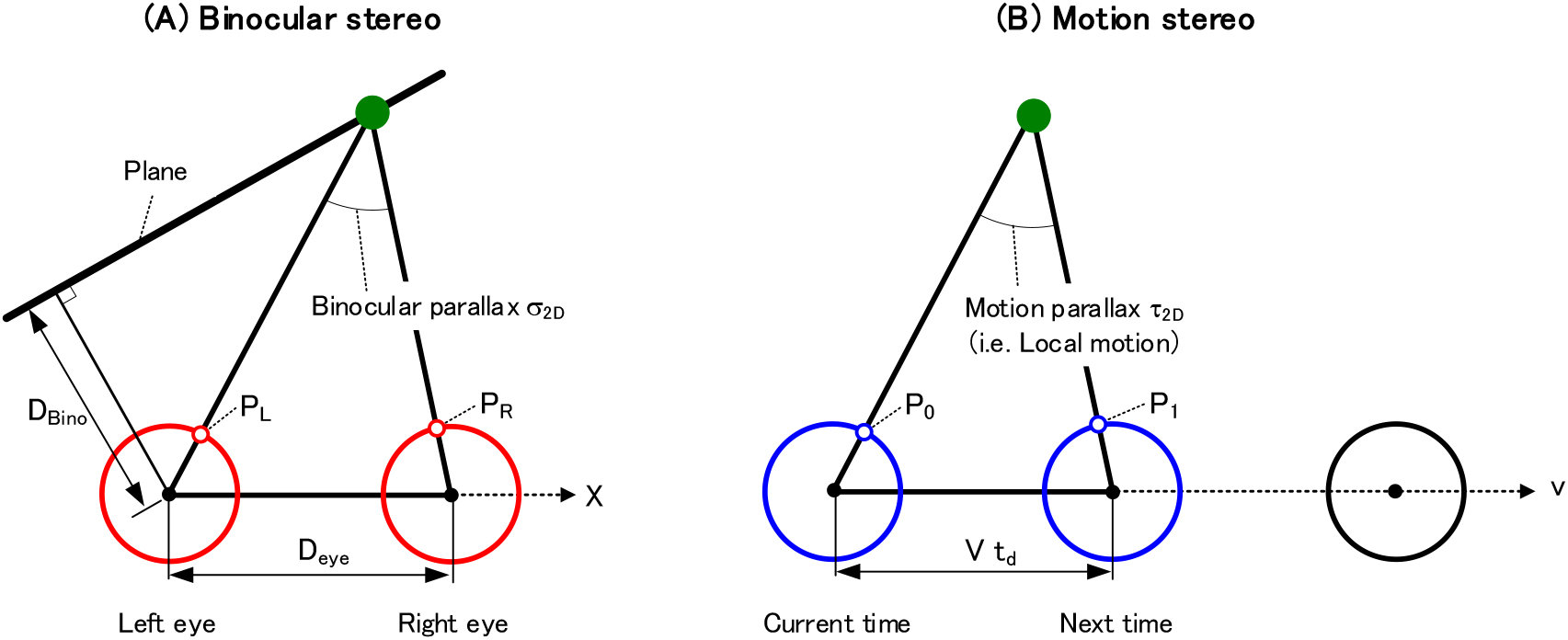
Binocular stereo is the same as motion stereo at each moment. Section **4.4.1** clarified that the parameters for binocular stereo are the same as those for motion stereo at each moment. Parameters necessary for this clarification are shown. (A) The parameters for binocular stereo are shown as follows. A cross section of binocular stereo (i.e. a cross section that passes between a green dot (i.e. an object) and the centers of both eyes) is shown. This dot is reflected in a position P_L_ (or P_R_) on the left (or right) eye. An angle at which the dot is seen with both eyes is the binocular parallax σ_2D_. The distance between both eyes, the direction from left eye to right eye, and the shortest distance to a plane are D_eye_, X, and D_Bino_, respectively. (B) The parameters for motion stereo are shown as follows. A cross section of motion stereo (i.e. a cross section that passes between a green dot and the centers of the eyes at the current and next times) is shown. This dot is reflected in a position P_0_ (or P_1_) on the eyeballs at the current (or next) time. An angle at which the dot is seen by the eyeballs at both times is the motion parallax (i.e. the local motion) τ_2D_. The distance between the eyeballs at the current and next times, and the movement direction of the eyeball from the current time to the next time are Vt_d_ and v, respectively.

**Figure 28.**
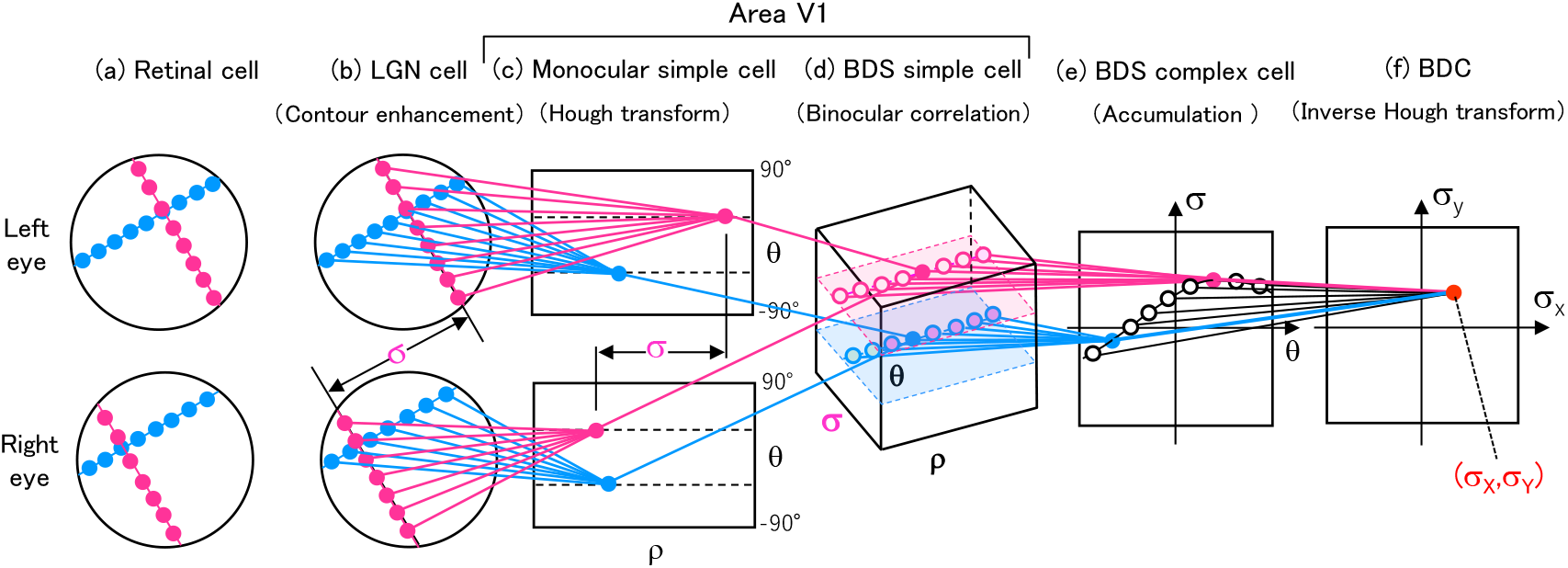
A series of modeled cells for detecting the binocular parallaxes (or disparities). A binocular parallax is detected by a series of modeled cells (i.e. LGN cells, monocular simple cells, BDS simple cells, BDS complex cells, and BDCs): here, BDS and BDC are an abbreviation for binocular-disparity selective and that for binocular-disparity-detection cell, respectively. The binocular parallax between the retinal cell arrays of the (a) is detected by a BDC firing in the (f) as its coordinate (σ_X_,σ_Y_). In these cell types, a series of transforms (i.e. the contour enhancement, Hough transform, binocular correlation, accumulation, and inverse Hough transform) are performed, respectively. The pink and blue straight lines represent the neural network connecting each cell array to perform the above transforms.

Comparing (A) and (B), binocular stereo is the same as motion stereo at each moment, and the terms used in (A) and (B) correspond as follows (Kawakami, 1996b; Kawakami et al., 2010, 1999).

Left eyeball vs. Eyeball at a current time

Right eyeball vs. Eyeball at a next time

Binocular parallax σ_2D_ vs. Motion parallax τ_2D_

Positions (i.e. P_L_ and P_R_) reflected in both eyeballs vs. Positions (i.e. P_0_ and P_1_) reflected in the eyeballs at the current and next times

Distance D_eye_ between both eyeballs vs. Distance Vt_d_ between the eyeballs at the current and next times

Direction X from the left eyeball to the right eyeball vs. Movement direction v of the eyeballs from a current time to the next time

Here, V is the movement speed and t_d_ is the unit time between the current time and the next time (see equation (1.0-2)).

In this way, the terms used for binocular stereo correspond to those for motion stereo. Therefore, replacing the terms (such as cell names and parameters) for motion stereo shown in Figure 17 to those for binocular stereo enables a series of cells, detecting binocularly the parameters of a plane, to be modeled. In Sections **4.4.3** and **4.4.4**, this series of cells will be modeled in Figure 30 using mathematical equations. In addition, the validity of this replacement will be confirmed in Sections **4.4.5** and **4.4.6**.

**Figure 29.**
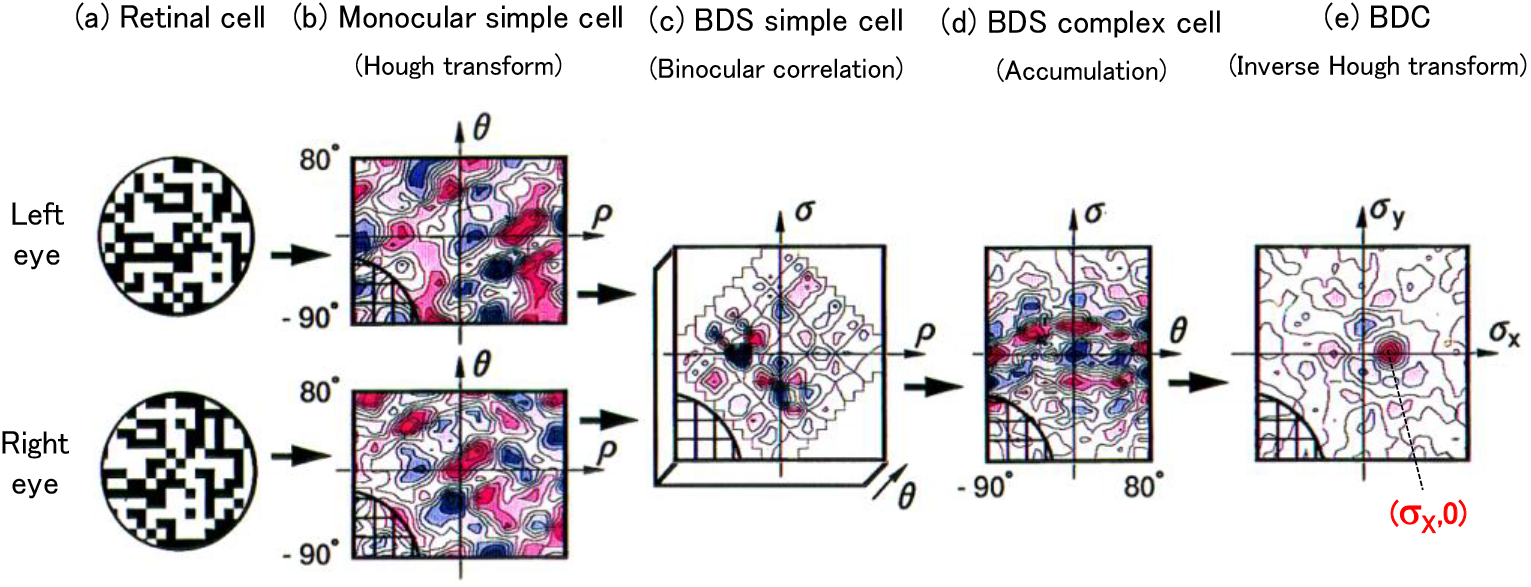
Simulation results of the binocular-parallax detection. A computer simulation confirmed that the series of modeled cells in Figure 28 correctly detects a binocular parallax. The random dots with a parallax (σ_X_,0) was given in the arrays of retinal cells ((a)). In the array of BDS complex cells in the (d), a sinusoidal firing pattern appeared in the same manner as DS complex cells of Figure 17(B)(e). The inverse Hough transform extracted this pattern to detect the parallax as the coordinate (σ_X_,0) of a BDC that had the isolated firing. Red is a positive response and blue is a negative response. Thus, it was confirmed that the given parallax was correctly detected by this series of cells. Each lattice in the lower left represents 2×2 cells.

**Figure 30.**
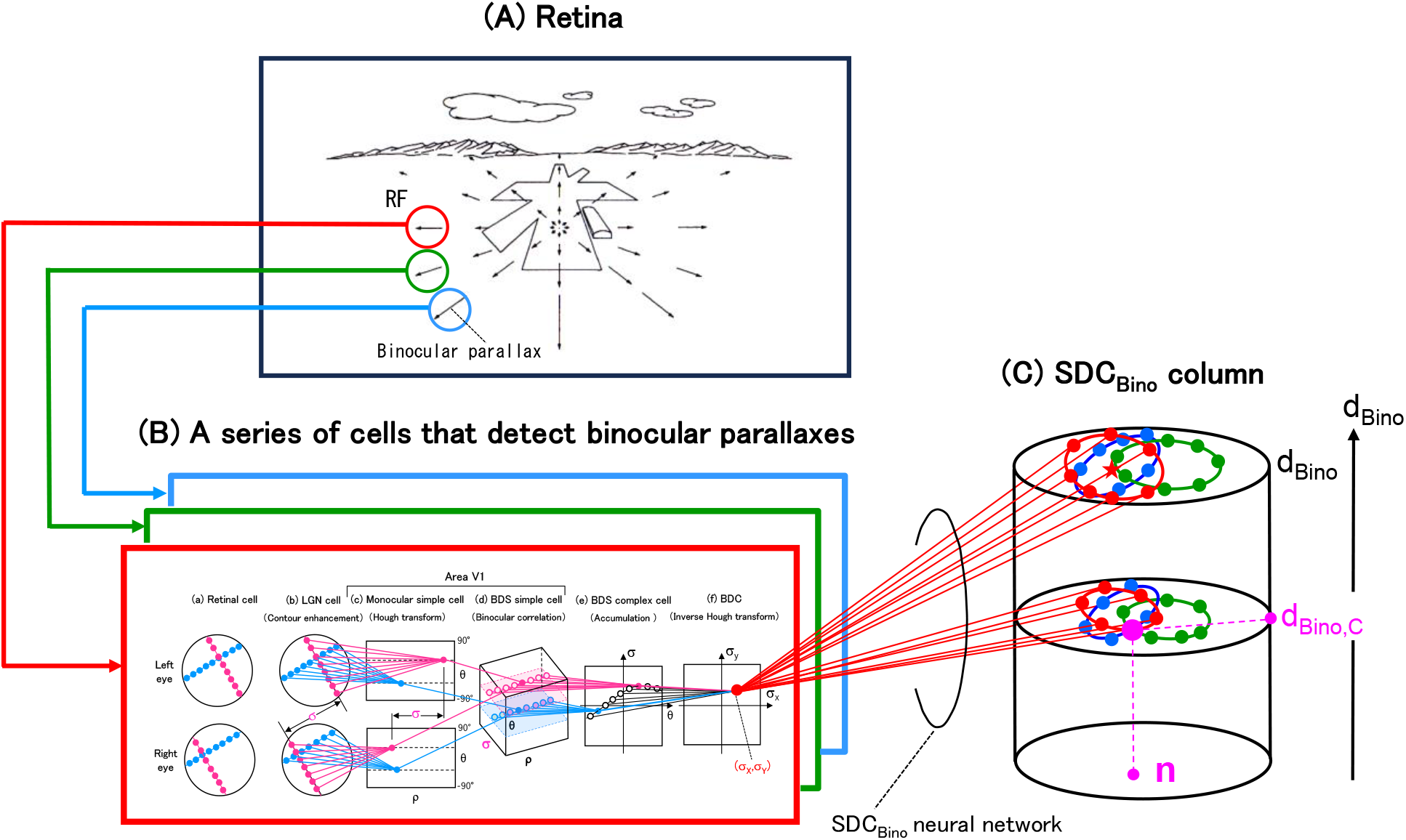
A cell model for detecting a plane with binocular stereo. A series of modeled cells that detects the parameters of a plane (i.e. the normalized shortest-distance d_Bino_ to a plane and its 3D orientation n) is shown. Based on the correspondence described in Section **4.4.1**, this series has been obtained by replacing the terms (such as cell names and parameters) of the series of cells for motion stereo in Figure 17 with those of a series of cells for binocular stereo. (A) A binocular parallax occurs in each small region of the retina (i.e. a RF): note that the (A) is a reuse of Figure 17(A) that shows the OFP of motion stereo, but it is not exactly correct. (B) This binocular parallax in the (a) is detected by the series of cells, which is shown enlarged in Figure 28. (C) A column composed of cells SDC_Bino_ is shown. Each cell SDC_Bino_ detects the normalized shortest-distance d_Bino_ to a plane as the height coordinate of the cell, and detects the 3D orientation n of the plane as the polar coordinate of the cell. The red straight lines represent a neural network connecting each BDC in the (B)(f) and the SDC_Bino_ column in the (C), and perform the small circle transform.

Before modeling this series, let us point out the following interesting relationship. That is, there is a interesting relationship of that the distance D_eye_ between both eyes is almost equal to the distance Vt_d_ between the eyeballs at the current and next times. Why this is so is explained as follows. The distance D_eye_ is said to be 6 cm on average for Japan people. In addition, assuming an average walking-speed V of 4 km/h, Vt_d_ is calculated to be 5.6 cm: here, t_d_ is assumed to be 50 ms based on physiological data (Mastronarde, 1987a, 1987b; Kawakami & Okamoto, 1996a). Therefore, these considerations yield the above relationship, and mean that the basic distances (i.e. D_eye_ and Vt_d_) for the binocular and motion stereos are almost the same. This sameness may be because these distances of D_eye_ and Vt_d_ are set to be the same in order to use both binocular and motion stereos simultaneously.

#### 4.4.2. A series of cells that detect binocular parallaxes

The correspondence in Section **4.4.1** enables a series of cells, which detect binocular parallaxes, to be modeled in Figure 28 by replacing the terms (such as cell names and parameters) in a series of cells that detect local motions in Figure 17(B) (see Section **2.1.2**) with the terms used in binocular stereo: this modeled cells were reported (Kawakami, 1996b; Okamoto & Kawakami, 1992; Kawakami & Okamoto, 1992d). This series is explained below.

First, the array of retinal cells in Figure 17(B)(a) has been divided into the arrays of the left and right eyes in Figure 28(a), and the arrays of the lagged and nonlagged LGN cells in Figure 17(B)(b) have been replaced with the LGN cells of the left and right eyes in Figure 28(b), respectively. Next, the cell types that detect the local motion in Figure 17(B) (i.e. NDS simple cell, DS simple cell, DS complex cell, and MDC) have been renamed to the cells that detect the binocular parallax in Figure 28 (i.e. monocular simple cell, BDS simple cell, BDS complex cell, and BDC): these cell types detecting the binocular parallax actually exist as described in Section **4.4.6** later. Here, BDS is an abbreviation for binocular-disparity selective, and BDC is that for binocular-disparity-detection cell.

The function (i.e. the transform) of each cell type is the same as that of motion stereo in Figure 17(B). That is, the Hough transform, accumulation, and inverse Hough transform have the same names, and the binocular correlation has the same function as the spatio-temporal correlation, although the names are different. The contour enhancement in Figure 28(b) is performed similarly to LGN cells for motion stereo in Figure 17(B)(b) (see Section **2.1.2**(2)). In addition, the network for binocular stereo in Figure 28 (shown by straight lines of pink and blue) is also the same as the network for motion stereo in Figure 17(B).

Therefore, since the function of each cell type and its neural network in Figure 28 are the same as those in Figure 17(B), the binocular parallax can be detected in the same way as the local motion. That is, the series of cells in Figure 28 enables the binocular parallax in Figure 28(a) to be detected as the coordinate (σ_X_,σ_Y_) of the BDC that fires in Figure 28(f).

Results of computer simulations shown in Figure 29 confirmed that this series of modeled cells in Figure 28 correctly detected the binocular parallax: this simulator was created by replacing the terms (such as cell names and parameters) for motion stereo in the Supplementary materials with the terms for binocular stereo. The results of the simulation in Figure 29 are described below. Note that LGN cells in Figure 28(b) have been omitted for ease of viewing.

Figure 29 is shown in a contour map, pink shows positive (or excitatory) responses, and blue shows negative (or inhibitory) responses. A random dot pattern with binocular parallax (σ_X_,0) was presented to the retinal cells of both eyes ((a)). In BDS complex cells ((d)), a sinusoidal firing pattern appeared similar to motion stereo in Figure 17(B)(e). Then, a BDC (σ_X_,0) in the (e) extracted this pattern by the inverse Hough transform and yielded the isolated firing. In this way, it was confirmed that the given binocular parallax was correctly detected as the coordinate (σ_X_,0) of the BDC.

#### 4.4.3. A cell model for detecting a plane with binocular stereo

Replacing the terms (such as cell names and parameters) for motion stereo in Figure 17 with those for binocular stereo enables cells that detect a plane by binocular stereo to be modeled in Figure 30, as follows.

The series of cells for detecting a binocular parallax in Figure 30(B) is the same as in Figure 28. A SDC_Bino_ column in Figure 30(C) has been obtained by replacing the height axis d of the SDC column in Figure 17(C) with the normalized shortest distance d_Bino_ for binocular stereo. Here, d_Bino_ is defined by the following equation, and has been obtained by normalizing the shortest distance D_Bino_ with the distance D_eye_ between both eyes (see Figure 27(A)).

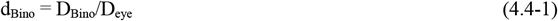

Figure 31 has been obtained by replacing the parameters (P_0_, P_1_, v, P_p0×p∞_, d, τ_2D_, and SDC) for motion stereo of Figure 16 with the parameters (P_L_, P_R_, X, P_pL×p∞_, d_Bino_, σ_2D_, and SDC_Bino_) for binocular stereo, respectively. That is, the positions P_0_ and P_1_ at the current and next times in Figure 16 were replaced with the positions P_L_ and P_R_ on each eye, and the movement direction v were replaced with the direction X from left eye to right eye. Figure 31 is used below to determine a SDC_Bino_ neural network that connects each BDC of Figure 30(B)(f) and the SDC_Bino_ column of Figure 30(C).

**Figure 31.**
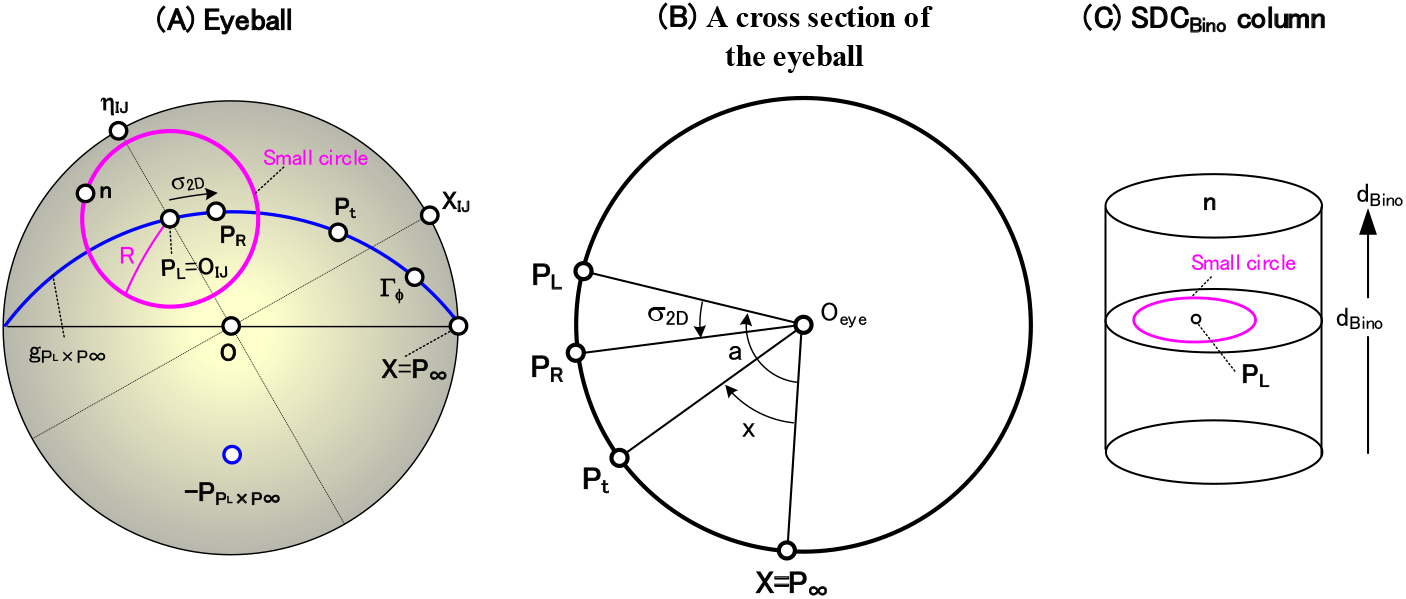
Detection of a plane by binocular stereo using the small-circle transform. (A) Parameters on the eyeball are shown. Positions P_L_ and P_R_ correspond to those for the left and right eyes, respectively, and the distance between P_L_ and P_R_ is the binocular disparity σ_2D_. The small-circle transform of equation(4.4-4) converts P_L_ into a pink small circle with radius R centered on P_L_: the 3D orientation n of a plane to be determined lies on this small circle. Here, X_IJ_ and η_IJ_ belong to each RF coordinate system (see equations(A4-4 and 10) in Appendix **4**), and P_L_=O_IJ_ (see equation(A4-3)). Γ_ϕ_ is the direction from P_L_ to P_R_ (see equation(A4-7) in Appendix **4**), where P_∞_=X. P_pL×P∞_ is the pole of the great circle g_pL×P∞_ and is calculated by equation(4.4-6). (B) A cross section of of the eyeball passing both the great circle g_pL×P∞_ in the (A) and the eyeball center O_eye_ is shown. The central angles (σ_2D_ and a) are used in the small-circle transform of equation(4.4-4). (C) The SDC_Bino_ column in Figure 30(C) is shown. The pink circle with radius R centered on a position P_L_ is drawn. When such small circle is drawn for all positions P_L_ and all heights, these small circles intersect only at a certain height d_Bino,C_ (see Figure 30(C) for d_Bino,C_). The normalized shortest-distance d_Bino,C_ to a plane is detected as this height coordinate, and its 3D orientation n is detected as the polar coordinate of this intersection.

Replacing the terms of the five steps in Section **3.1.2** with the terms for binocular stereo enables the SDC_Bino_ neural network to be determined. It is described below using Figures 30 and 31.

[Step 1] Give two parameters in Figure 31(A) (i.e. a position P_L_ on the left eye and a binocular parallax σ_2D_ between P_L_ and P_R_), as follows. First, give the position P_L_ as the RF center O_IJ_ based on equation(A4-3) of Appendix **4**: in this equation, P_0_ was replaced with P_L_. Next, give the binocular parallax σ_2D_ by the following equation, where (σ_X_,σ_Y_) is the coordinate of the BDC in Figure 30(B)(f).

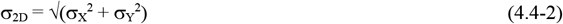

The P_∞_ is calculated by the following equation, where X = (1,0,0).

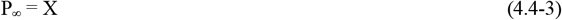

[Step 2] Set a normalized shortest-distance d_Bino_ arbitrarily.

[Step 3] Calculate a radius R used for the small circle transform by the following equation (see equation(3.0-6)).

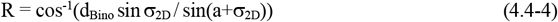

The variable a used above is calculated by the following equation (see equations(1.0-7 and 9) and Figure31(B)).

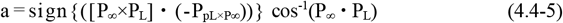

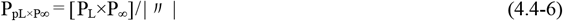

[Step 4] Plot the left eye’s position P_L_ on the cross section of height d_Bino_ in Figure 30(C): this plotted P_L_ is shown as a red mark ★. Then, draw a red small circle with radius R, centered on P_L_, on the cross section. The 3D orientation n of a plane to be determined exists on this small circle (see Section **3.1.2**).

Finally, connect all cells SDC_Bino_ (indicated as red dots) on this small circle to the BDC (σ_X_,σ_Y_) in Figure 30(B)(f). This connection shown as red straight lines represents an SDC_Bino_ neural network in Figure 30(C). The above is for the red RF in Figure 30(A), but for the green and blue RFs, the green and blue small circles and all cells SDC_Bino_ on each small circle are drawn: the neural network is omitted for ease of viewing. These small circles do not intersect at a single point.

[Step 5] When such a small-circle drawing is performed for all heights d_Bino_, the small circles intersect at one point only at some height d_Bino,C_ (see Figure 30(C) for d_Bino,C_). A normalized shortest-distance d_Bino,C_ to the plane is detected as the height coordinate of the pink intersection cell, and its 3D orientation n is detected as the polar coordinate of this cell.

In this way, a neural network connecting each BDC in Figure 30(B)(f) and the SDC_Bino_ column in Figure 30(C) has been modeled. This network detects the normalized shortest-distance d_Bino_ to the plane and its 3D orientation n. Although the essential points of the network have been explained with an emphasis on comprehensibility, Section **4.4.4** will show that the network is expressed accurately using mathematical equations. Section **4.4.5** will also show that the validity of this network is confirmed by computer simulations.

#### 4.4.4. Representation of the SDC_Bino_ neural network by mathematical equations

Let us express the neural network, described in Section **4.4.3**, by mathematical equations using the form of a flowchart, as follows (see Figures 30 and 31). This has been obtained by replacing the series of equations for motion stereo in Section **3.2** with those for binocular stereo.

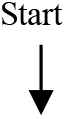

(1) Give the direction X from the left eye to the right eye in Figure 27(A) Calculate P_∞_ using the following equation (see equation(4.4-3)), where X = (1,0,0).

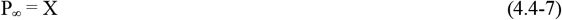
(2) Sweep a RF center O_IJ_ Calculate P_L_ using the following equation.

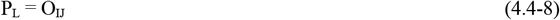 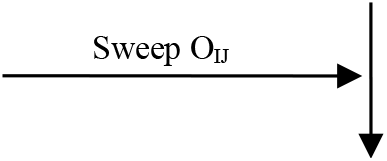
(3) Sweep a normalized shortest-distance d_Bino_ 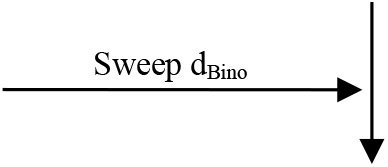
(4) Sweep a BDC array (σ_X_,σ_Y_) 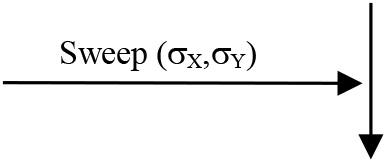 Convert this (σ_X_,σ_Y_) to its polar coordinate (σ_2D_,ϕ).

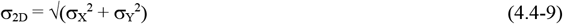

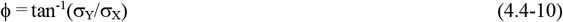
(5) Calculate the radius R used for the small-circle transform by the following equation (see equation(3.0-6)).

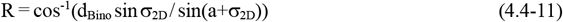 The variable a used above is calculated by the following equations (see equations(4.4-5 and 6)).

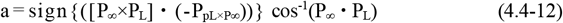

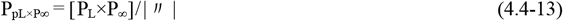
(6) Calculate a small circle with radius R centered at position P_L_ (Figure 31(A)) First, P_L_ expressed by equation(4.4-8) is represented in an orthogonal coordinate system by the following equation.

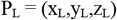 Convert the P_L_ to a polar coordinate (α_L_,β_L_) by the following equation.

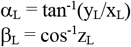 Next, when an arbitrary point on the small circle is represented as a polar coordinate (α,β), there is the following relationship (see equation(A7-1) in Appendix **7**).

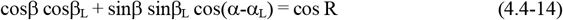 This is the mathematical expression of the pink small-circle with radius R centered at position P_L_ (see Figure 31(A)). There is an important property that the 3D orientation n of a plane to be determined lies on this small circle.
(7) Draw this small circle in the SDC column Using the equidistant projection of Figure 10, convert this small circle on the eyeball into a pink small-circle on the cross section at the height d_Bino_ of the SDC_Bino_ column in Figure 31(C). Then, draw this converted small circle on the cross section. Note that the small circle on the cross section is represented by the same equation(4.4-14) as that on the eyeball (see Section **A7.2** in Appendix **7**).
(8) Connect all cells SDC_Bino_ on this small circle with the BDC (σ_X_,σ_Y_) Connect all cells SDC_Bino_ (indicated as red dots in Figure 30(C)) on this small circle at height d_Bino_ to the BDC (σ_X_,σ_Y_) of Figure 30(B)(f). This connection shown as red straight lines represents an SDC_Bino_ neural network in Figure 30(C). 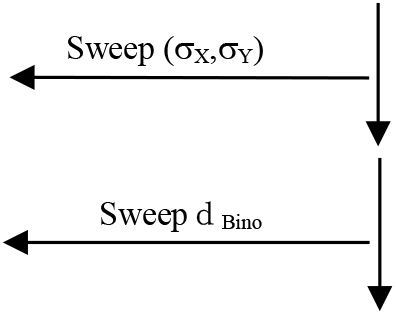
(9) Up to this point, the red neural network in Figure 30(C) have been created. That is, the neural network for the red RF in Figure 30(A) has been completed. 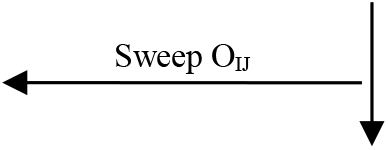
(10) A neural network for all RFs in Figure 30(A), including red, green, and blue, has been completed. Each connection of the neural network (Figure 30(C)) connecting the BDCs and the SDC_Bino_ column has been accurately expressed one by one using mathematical equations. 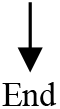

In this way, the SDC_Bino_ neural network of Figure 30(C) has been accurately expressed by mathematical equations at the level of the neural connection of which cell is connected to which cell. The above is represented by the complex equations such as equation(4.4-11), and you may think that it is difficult for cerebral cells to execute them. However, that worry is not a problem for the same reasons as in Section **2.2**(12).

#### 4.4.5. Simulation results

Using the mathematical expression in Section **4.4.4**, a computer simulation of the cell model (Figure 30) for detecting the normalized shortest distance to a plane and its 3D orientation was performed: this simulator was created by replacing the terms for motion stereo in Supplementary materials with the terms for binocular stereo. The simulation result is shown in Figure 32. This result is taken from a joint simulation (Okamoto, 2006) based on Supplementary materials.

**Figure 32.**
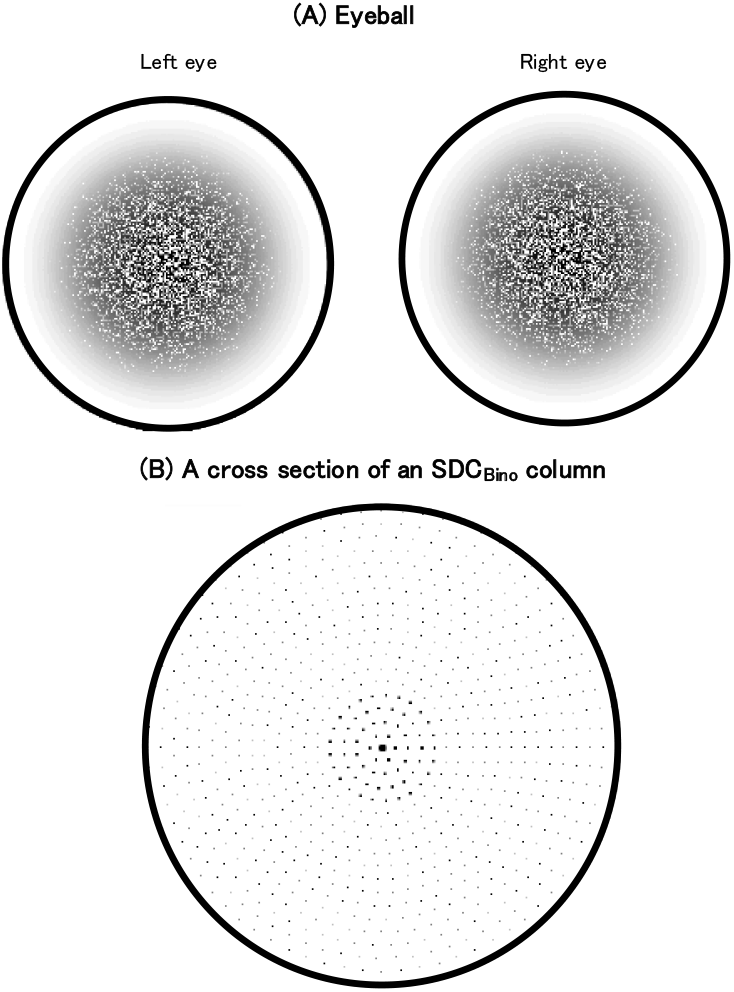
Simulation results detecting the normalized shortest-distance and 3D orientation by binocular stereo. (A) Perpendicular plane on which random dots are drawn are reflected on the left and right eyes. (B) A cross section of the SCD_Bino_ column (see Figure 30(C)) at a height d_Bino,C_ that corresponds to the given normalized-shortest-distance is shown: a response of each SCD_Bino_ is indicated by a dot’s size. The cell SCD_Bino_ corresponding to the 3D orietation n=(0°,0°) showed the maximum response, and thus it was confirmed that the shortest distance d_Bino_ to the given plane and its 3D orientation n were detected correctly.

A stimulus image in which a random dot pattern was drawn on a plane having a 3D orientation n (0°,0°) was presented to both eyes. The images reflected on the left and right eyeballs are shown in Figure 32(A). The responses of the SCD_Bino_ column in Figure 30(C) to this image were performed by the above simulator.

A cross section of the SCD_Bino_ column at the given normalized-shortest-distance d_Bino_ is shown in Figure 32(B): a response of each SCD_Bino_ is displayed as the dot’s size. It was confirmed that the SCD_Bino_ with the given orientation (0°,0°) had the maximum response and thus could correctly detect the shortest distance to the plane and its 3D orientation.

#### 4.4.6. Comparison with Psychology and Physiology

##### 4.4.6.1. Comparison with psychology

Psychological experiments (Julesz, 1960, 1971, 1986) discovered that the binocular stereo enables the shortest distances to planes and their 3D orientations of random dot stereograms (which are difficult to be recognized with monocular vision) to be recognized.

It is surprising that the series of modeled cells in Figure 30 enables this psychological discovery to be explained neurophysiologically at the level of the neural connection of which cell is connected to which cell (see Section **4.4.4**). This neurophysiological explanation is, to my knowledge, the first time.

##### 4.4.6.2. Comparison with physiology

The series of modeled cells in Figure 30 is compared with physiological experiments below.

Binocular cells that respond selectively to the 3D orientation of a plane were reported in the caudal intra-parietal area (i.e. the area CIP) of monkeys (Tsutsui et al., 2002). The modeled cell SDC_Bino_ in Figure 30(C), which has the same selectivity, can explain these actual cells neurophysiologically at the level of the neural connection of which cell is connected to which cell.

However, experiments on cells that detect the shortest distance to a plane have not been reported, and thus the modeled cell SDC_Bino_ that detects this distance D_Bino_ to the plane cannot be verified. On the other hand, it was described in Sections **4.4.2** and **4.4.4** that the neural networks of the binocular and motion stereos are the same. Therefore, this modeled SDC_Bino_ that detects the shortest distance by binocular stereo may exist in the cerebrum, similar to existing the modeled SDC in area MST that was described in Section **4.2.2.2**.

In area V1, BDS simple cells (Poggio & Ficsher, 1977; Ohzawa et al., 1990: Freeman & Ohzawa, 1990) and BDS complex cells (Ohzawa et al., 1990) were reported. Also, BDCs in area V2 (Hubel & Livingston, 1987, 1990; Poggio et al., 1988) were reported. The selective responses of these cells to light stimuli are the same as those of the series of modeled cells in Figure 30(B) (see Section **6.4** of Kawakami (1996b)). Thus, this series enables the above reports to be explained neurophysiologically.

Therefore, it has been shown that the series of modeled cells in Figure 30 actually exists.

##### 4.4.7. How to use the forward and backward methods properly according to their characteristics

In Figures 24(A)(iii) and 24(B)(iii), let us change the name of the SDC column for motion stereo to the SDC_Bino_ column for binocular stereo, and replace the height axis t with the height axis d_Bino_ for binocular stereo. When such change and replacement have been made, the forward SDC_Bino_ neural network in Figure 24(A) was expressed by mathematical equations in Section **4.4.4**. In addition, the backward SDC_Bino_ neural network in Figure 24(B) was expressed by mathematical equations in Appendix **10**.

Similar to the explanation for motion streo in Section **4.3.3.2**, these forward and backward methods for binocular stereo need to be used properly according to their characteristics, as follows. That is, the forward method is suitable for ex plaining the mathematical structure of the neural network, and the backward method, which is simple in procedure, is suitable for simulation.

### 4.5. The Hough transform is a special case of the polar transform

The Hough transform played an important role for modeling the NDS simple cells in Figure 9(B)(c) (see Section **2.1.2**(3)). The polar transform also played an important role for modeling the SDCs in Figure 9(C) (see [Step 4] in Section **2.1.4**). It is described below that there is a close relationship between these two important transforms (i.e. there is a relationship that the Hough transform is a special case of the polar transform).

#### 4.5.1 Hough transform

With reference to Figure 33, let us explain the Hough transform, as follows. This transform that was proposed by Hough (1962) converts any point P in the (x,y) plane of the (A) to a sine wave in the (ρ,θ) plane of the (C) by the following equation (Hough, 1962: Duda & Hart, 1972, 1973; Kawakami & Okamoto, 1996a, 1995a; Kawakami, 1996b).

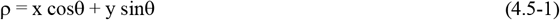

This equation is called Hesse’s standard form among the various representation forms of a straight line (Izumi et al., 1987).

Although Hough (1962) and Duda & Hart (1972, 1973) derived this algebraically, it is shown below that this transform can be derived geometrically.

First, convert any point P in the (x,y) plane to all straight lines through that point ((A)): as an example, the red, blue and green straight lines are shown. Each straight line can be represented using a perpendicular foot (ρ,θ) from origin 0 ((B)): as an example, the red, blue and green perpendicular-foots (ρ,θ) are shown. These perpendicular foots lie on a circle with a diameter passing through 0 and P.

Next, plot the red, blue, and green perpendicular-foots (ρ,θ) in the (ρ,θ) plane of the (C). These foots (ρ,θ) are on a sine wave with an amplitude of 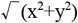 and a phase of tan^-1^(y/x). This sine wave is derived in the (D), as follows: only the red straight line of the (B) is extracted and shown in the (D). Focusing on the right triangle HP0 yields the following equation: here, ξ = tan^-1^(y/x).

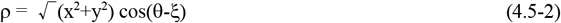

This equation represents the sine wave of the (C). Expand the above equation to become equation(4.5-3). Since 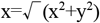 cosξ and 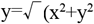 sinξ, the Hough transform has been obtained as equation (4.5-4).

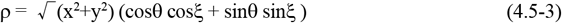

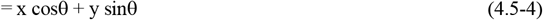

In this way, the Hough transform has been derived using three types of geometrical conversions. That is, first, a point (x,y) of the (A) was converted to all straight lines passing through that point. Second, these straight lines were converted to a circle ((B)) whose diameter passes through 0 and (x,y). Finally, this circle was converted to a sine wave, expressed by equation(4.5-4), in the (ρ,θ) plane of the (C). Thus, the Hough transform of equation (4.5-1) has been derived geometrically.

Here, note the following. The sine wave in the (C) can be detected as the point P(=(x,y)) in the (D).This is because the perpendicular foot 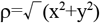, θ=tan^-1^(y/x)) of a straight line at this point P is equal to the amplitude and phase of the sine wave, respectively, as shown in the (D). This equality (or identity) derives the inverse Hough transform, which detects the sine wave (with the amplitude 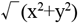 and phase tan^-1^(y/x)) on the (C) as the coordinate (x,y) on the (D): note that this (x,y) corresponds to the (τ_X_, τ_Y_) in Figure 9(B)(f). Thus, the inverse Hough transform has been derived geometrically.

The Hough transform and the inverse Hough transform can be summarized as follows. The Hough transform converts a point P in the (A)to the corresponding sine wave in the (C). On the other hand, the inverse Hough transform converts the a sine wave in the (C) to the corresponding point P in the (D), and thus detects the this sine wave as the point P.

Finally, let us consider as follows how a red straight line in the (x,y) plane ((D)) is detected as the corresponding red point (ρ,θ) in the (ρ,θ) plane ((C)) by the Hough transform. Each point on this straight line is converted into a corresponding sine wave in the (ρ,θ) plane by the Hough transform, as described above. When such conversion is performed for all points on this line, the resulting sine waves intersect at one point. This is because the coordinate (ρ,θ) of this intersection ((C)) is equal to the red perpendicular-foot (ρ,θ) of the line ((D)). Therefore, it has been indicated that the red straight line in the (x,y) plane ((D)) is detected as the red point in the (ρ,θ) plane ((C)) by the Hough transform. This can be generalized as follows. Any straight line in the (A) is detected as the corresponding point (ρ,θ) in the (C) by the Hough transform.

#### 4.5.2. The Hough transform is a special case of the polar transform

We reported that the Hough transform is a special case of the polar transform (Kawakami et al., 1991, 1992a). It is described below using Figure 34.

**Figure 33.**
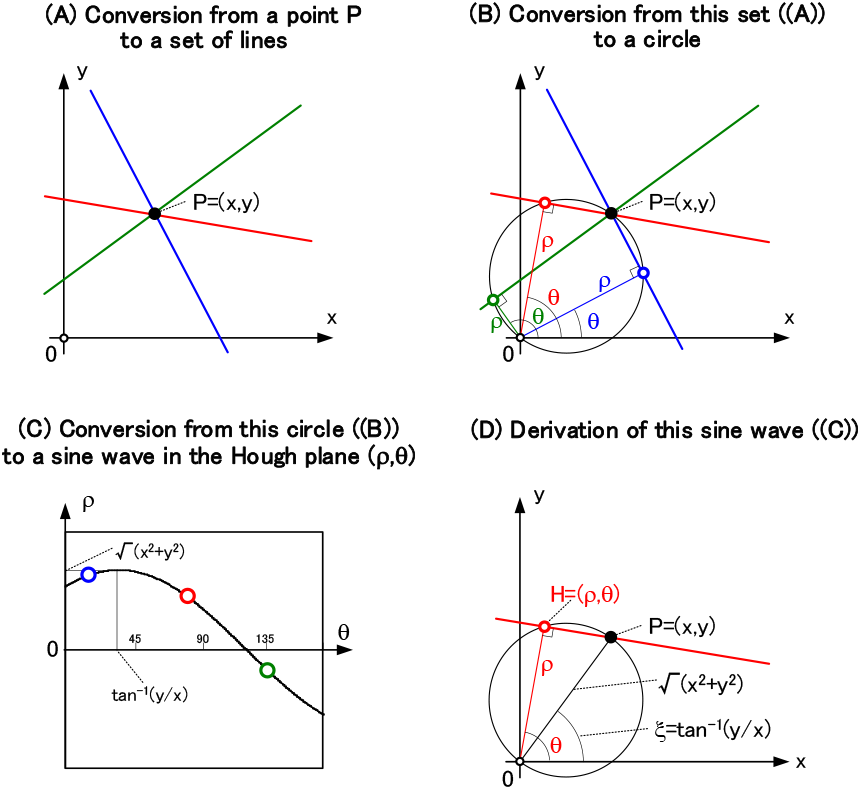
Derivation of the Hough transform. The Hough transform can be derived geometrically, as follows. (A) First, convert any point P in the (x,y) plane to all straight lines through that point: the red, blue, and green lines are shown as examples. (B)Next, each of these straight lines can be represented as a perpendicular foot (ρ,θ) from the origin 0: red, blue, and green perpendicular foots (ρ,θ) are shown as examples. These foots lie on a circle with a diameter passing through 0 and P. Therefore, all straight lines passing through the point P in the (A) are converted into this circle. (C) Finally, convert this circle in the (B) into a sine wave, in the (ρ,θ) plane, that is represented by equation(4.5-4). In this way, these three types of conversions have yielded geometrically the Hough transform. (D) Only the red straight line in the (B) was extracted. Focusing on the right triangle HP0, the sine wave of the (C) has been derived as equation(4.5-2).

**Figure 34.**
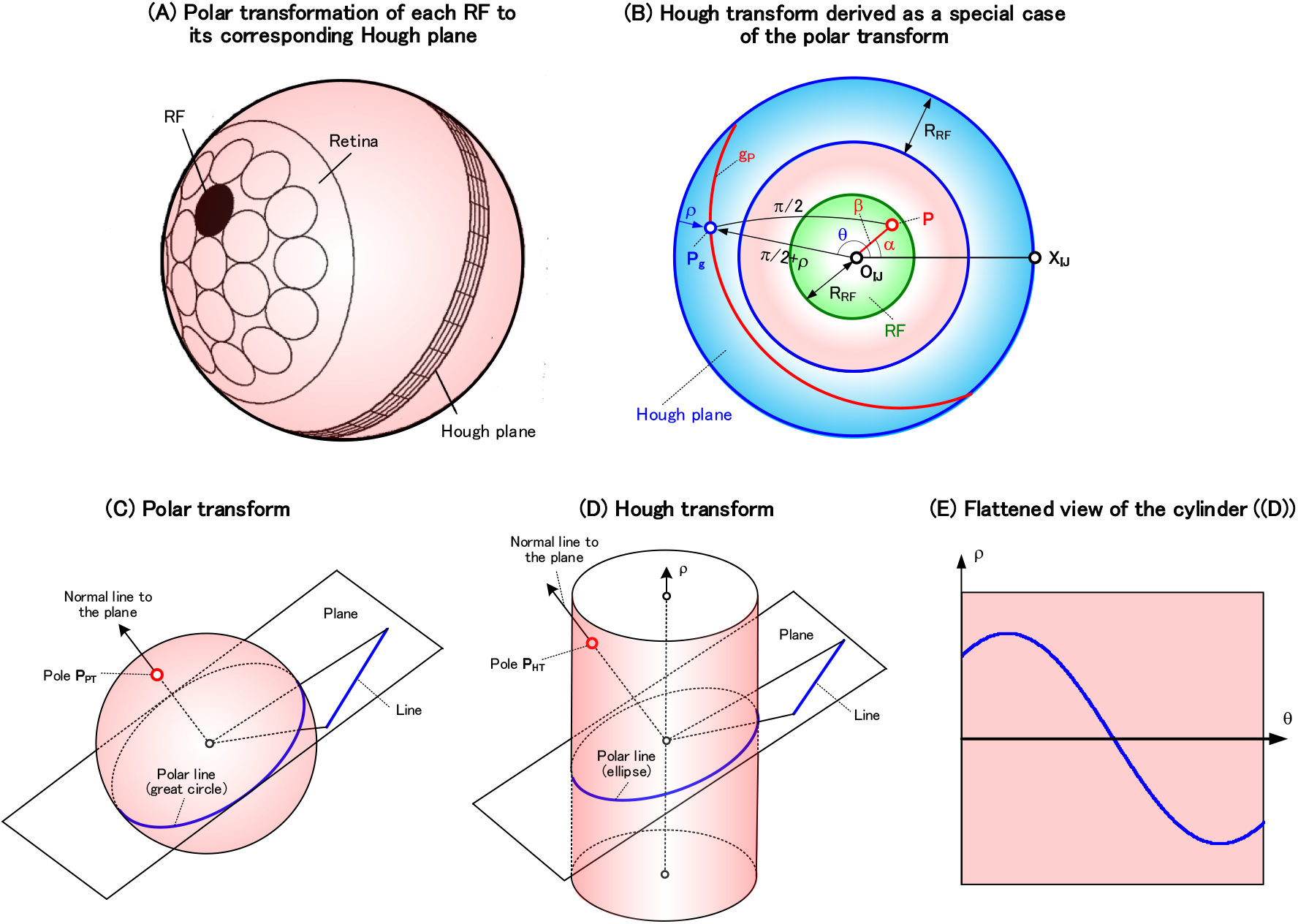
The Hough transform is a special case of the polar transform. (A) The relationship, on the eyeball, between each RF and its corresponding Hough plane is shown. This Hough plane has been obtained as the polar transformation of the RF, and corresponds to the (ρ,θ) plane of Figure 33(C). (B) The display in the (A) has been changed to center on the RF center O_IJ_. The RF is shown in green, the Hough plane is shown in blue, and R_RF_ is the radius of the RF. Expressing red any point P on the RF as (α,β) in a polar coordinate, we obtain a red great circle g_P_ in the Hough plane as the polar transformation of this point (see equation(4.5-7)). Since the radius R_RF_ of the RF is small, equation(4.5-7) is approximated to equations(4.5-8 and 10), and thus the Hough transform has been derived as a special case of the polar transform when the radius R_RF_ is small. (C) The polar transform is expressed as follows. Using a spherical surface enables the intersection point between a normal line and the surface to be represented as the pole P_PT_, and enables the intersection line between the plane perpendicular to this normal line and the surface to be represented as a polar line (i.e. a great circle). This representation allows the polar transform to be expressed as a conversion from the pole P_PT_ to the great circle. (D) The Hough transform is expressed as follows. Using a cylinder enables the intersection point between a normal line and the cylinder to be represented as the pole P_HT_, and enables the intersection line between the plane perpendicular to this normal line and the cylinder to be represented as a polar line (i.e. an ellipse). This representation allows the Hough transform to be expressed as a conversion from the pole P_HT_ to the ellipse. (E) Splitting the cylinder of the (D) vertically and developing it into a plane cause this ellipse to be converted to a sine wave on the (ρ,θ) plane that corresponds to the Hough plane in Figure 33(C). Note that the range θ=0~π is shown for contrast with Figure 33(C).

First, the relationship on the eyeball between a RF and its corresponding Hough plane is shown in the (A): this Hough plane corresponds to the (ρ,θ) plane of Figure 33(C). The polar transformation of this RF gives the Hough plane (ρ,θ) as a ring, whose width and circumferential directions are respectively ρ and θ of the (ρ,θ) plane.

The (B) is obtained by displaying the (A) at the center of the RF center O_IJ_, where R_RF_ is the radius of a RF. In the (B), the RF in the (A) is shown in green and its Hough plane is shown in blue: see Figure S2-2 of Supplementary materials for the correspondence between the RF and the Hough plane. Representing a red arbitrary point P within the RF as (α,β) in the polar coordinate, convert this point P to a red great circle g_P_ by the polar transform. Let us express this great circle with an equation, as follows.

Denote any point P_g_ on this red great circle as the polar coordinate (θ,π/2+ρ): this π/2 is due to the definition that the origin of ρ is on the blue outermost circle (i.e. on the equator when O_IJ_ is the north pole). Applying the cosine rule (Bronshtein & Semendyayev, 1978; Izumi et al., 1987) to a spherical triangle P_g_O_IJ_P yields equation(4.5-5), and then rearrange it to become equations(4.5-6 and 7). Equation(4.5-7) is the mathematical expression of this great circle g_P_: the trajectory of (θ,ρ) represents the great circle.

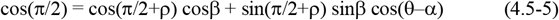

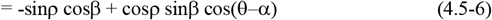

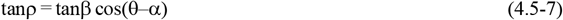

Since the diameter of the RF is less than 5 degrees (Kawakami & Okamoto, 1996a; Kawakami, 1996b), ρ and β are also small, and thus equation(4.5-7) is approximated as equation(4.5-8). The relations of x=βcosα and y=βsinα (see (B)) yields equation(4.5-10), and thus the Hough transform of equation(4.5-1) has been derived geometrically.

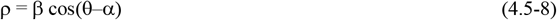

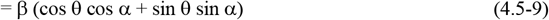

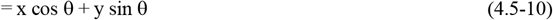

In this way, the Hough transform has been shown to be a special case of the polar transform when RF is small. The same applies to the inverse Hough transform of Figure 9(B)(f).

The cell model of Figure 9 uses the Hough, the inverse Hough, and the polar transforms. Since the Hough and inverse Hough transforms are special cases of the polar transform, renaming the Hough and inverse Hough transforms to the polar transform allows this model of Figure 9 to be consist of a time delay, spatio-temporal correlation, accumulation, cross-ratio transform, and three polar transforms.

#### 4.5.3. Geometrical relationship between the polar and Hough transforms

Let us go back to the origin of the polar and Hough transforms to consider their geometrical relationships using Figure 34. The fundamental 3D parameters that stipulate each transform are a plane and its normal line in the (C) or (D), and the essence of each transform is the transformation from the normal line to the plane (Kawakami et al., 1991). This transformation yields a polar transform in the (C) or a Hough transform in the (D), as follows.

To represent these 3D parameters (of the plane and its normal line) in two dimension, the polar transform uses a spherical surface and the Hough transform uses a cylinder (Kawakami et al., 1991). That is, using the spherical surface for the polar transform ((C)) enables the intersection point between the normal line and the surface to be expressed two-dimensionally as a pole P_PT_ on the spherical surface, and enables the intersection line between the plane and the surface to be expressed two-dimensionally as a polar line (i.e. a great circle) on it. Thus, the polar transform is a transformation, on the spherical surface, from the pole P_PT_ to the great circle: these pole and great circle correspond to the red P and the red great circle g_P_ in the (B), respectively. In addition, using the cylinder for the Hough transform ((D)) enables the intersection point between the normal line and the cylinder to be expressed two-dimensionally as a pole P_HT_ on the cylinder, and enables the intersection line between the plane and the cylinder to be expressed two-dimensionally as a polar line (i.e. an ellipse) on it. Thus, the Hough transform is a transformation, on the cylinder, from the pole P_HT_ to the ellipse. Note the following. The reason why the (D) is the Hough transform is that if the cylinder is split vertically and expanded on a plane ((E)), this ellipse yields a sine wave on the (ρ,θ) plane that corresponds to the Hough plane of Figure 33(C). In this way, using the spherical surface or the cylinder has enabled the 3D parameters, representing a plane and its normal line, to be expressed as the 2D parameters (i.e. the polar line and the pole). Note that the polar transform (or the Hough transform) from the pole to the polar line results from the transformation in space from the normal line to the plane.

The polar transform is superior to the Hough transform. That is, while the polar transform can represent any plane as a great circle ((C)), the Hough transform has a disadvantage that the diameter of the ellipse changes when the inclination of the plane changes ((D)). In extreme cases, this diameter becomes infinite on a plane parallel to the ρ axis and cannot be expressed.

Note the following. The inverse polar transform is the transformation from the great circle to the pole P_PT_ (see (C)), and the inverse Hough transform is the transformation from the ellipse to the pole P_HT_ (see (D)). Opposite to the above, this transform from the great circle (or the ellipse) to the pole P_PT_ (or P_HT_) results from the transformation in space from the plane to its normal line. Also note that the relationship between the polar transform (or Hough transform) and the inverse polar transform (or inverse Hough transform) belongs to a type of duality (The Mathematical Society of Japan, 1990).

Finally, let us point out an interesting point that the series of cells in Figure 9(B) is processed on the spherical surface (or on the eyeball). This processing on the spherical surface, abbreviated as the sphere, can be summarized in the following four parts. First, retinal cells and LGN cells (Figures 9(B)(a) and 9(B)(b)) are processed in the (x,y) plane of

Figure 33(A): this processing in the (x,y) plane is processed in each RF on the sphere (or the eyeball) of Figure 34(A), and is precisely processed on the tangent plane of Figure S2-3(A) of Supplementary materials. Next, NDS simple cells of Figure 9(B)(c) perform the Hough transform, and thus the (x,y) plane above is transformed into the Hough plane (i.e. (ρ,θ) plane) of Figure 33(C): this (ρ,θ) plane performed by the NDS simple cells corresponds to the blue ring in Figure 34(B), and thus is processed on the sphere. Furthermore, processings of DS simple cells and DS complex cells (Figures 9(B)(d) and 9(B)(e)) are also performed in this Hough plane on the sphere. Finally, MDCs of Figure 9(B)(f) perform the inverse Hough transform, and thus the Hough plane (i.e. the (ρ,θ) plane) above is transformed into the (x,y) plane of Figure 33(A): this (x,y) plane corresponds to the (τ_X_, τ_Y_) array in Figure 9(B)(f) (see Figure A4(D) in Appendix **4**); this (τ_X_, τ_Y_) array performed by the MDCs correspond to the green RF in Figure 34(B), and thus is processed on the sphere. In this way, it has been clarified that the series of cells of Figure 9(B) is topologically processed on the sphere. This clarification has been described in more detail in the caption of Figure 35.

**Figure 35.**
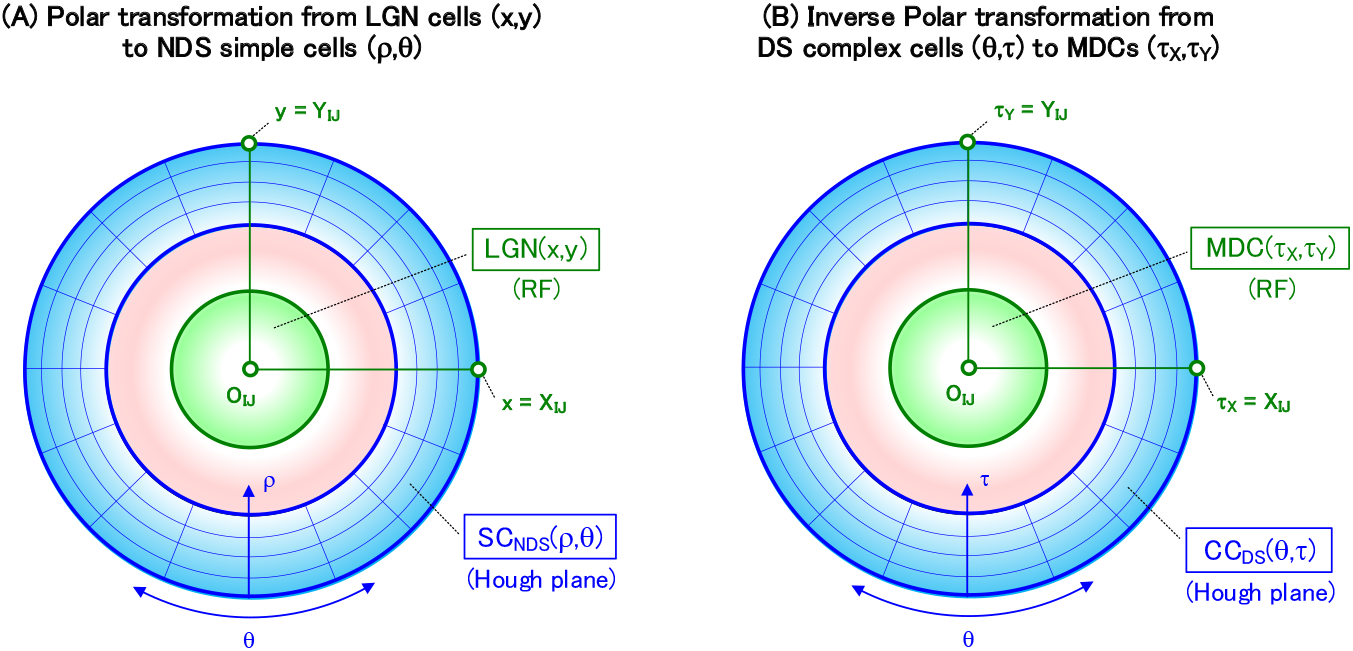
Conversion between the RF and the Hough plane. (A) This is the same as Figure 34(B), but its purpose is different as follows. While the purpose of Figure 34(B) is to explain that the Hough transform is a special case of the polar transform, the purpose of this figure is to explain how LGN cells of Figure 9(B)(b) are converted to NDS simple cells of Figure 9(B)(c). Let us explain this conversion as follows. The Hough transform (i.e. the polar transform above) performed by NDS simple cells converts the LGN cells LGN(x,y) in the green RF into the NDS simple cells SC_NDS_(ρ,θ) in the blue Hough plane: the local coordinate system (O_IJ_,X_IJ_,Y_IJ_) of each RF was described in Section **S1.2.3** of Supplementary materials. In this way, the LGN and NDS simple cells are processed on the spherical surface (i.e. on the eyeball). (B) The inverse Hough transform (i.e. the inverse polar transform) performed by MDCs converts the DS complex cells CC_DS_(θ, τ) in the blue Hough plane into the MDCs MDC (τ_X_, τ_Y_) in the green RF. In this way, the DS complex cells and MDCs are processed on the spherical surface. Extending this, it is shown below that the series of cells in Figure 9(B) is processed on the spherical surface (i.e. on the eyeball). First, LGN and NDS simple cells were processed on the eyeball, as shown in the (A). Second, the DS simple cells SC_NDS_(ρ,θ, τ) are processed in the Hough plane (θ, τ) (i.e. the eyeball) of the (B): the coordinate ρ is processed in the radial direction of the sphere. Third, the DS complex cells CC_DS_(θ, τ) are processed in the blue Hough plane (θ, τ) (i.e. the eyeball) in the (B). Finally, the MDCs MDC (τ_X_, τ_Y_) are processed in the green RF, on the eyeball, of the (B): this RF corresponds to both the RF in Figure A4(B) of Appendix **4** and the MDC array in Figure A4(D). In this way, it has been indicated that all cell types in Figure 9(B) is processed on the spherical surface (i.e. the eyeball).

In addition, by combining this processing on the sphere in Figure 9(B) with the cross-ratio and polar transforms performed by SDCs in Figure 9(C), it has been finally clarified that all cell types in Figure 9 are topologically processed on the sphere: note that these transforms are processed on the sphere as described in Section **2.1.4**. This clarification is interesting. I point out that all cell types in Figure 17 are also processed on the sphere.

These processings on the sphere yield the following two features. First, on the plane, circles and straight lines must be treated separately from each other, but on the sphere, they correspond to small circles and great circles, respectively, so they can be treated unifiedly as circles. In other words, since any figure on the sphere is a composite of circles, all figures can be handled unifiedly only with circles. The other is that there is no infinity unlike on the plane, and it can be processed with finiteness (i.e. with the maximum value π) on the sphere. Therefore, these two features enables all figures on the sphere to be handled by a combination of circles that are finite.

This geometry on the sphere belongs to a type of non-Euclidean geometry (i.e. elliptic geometry) (The Mathematical Society of Japan, 1990). Thus, in this spherical geometry, there are no exceptional figures such as straight lines and parallel lines, and all figures can be handled uniformly with only circles.

Furthermore, an arbitrary 3D world can be projected onto the sphere as a 2D image. This projection has made it possible to handle the transforms (i.e. the polar, cross-ratio, and small-circle transforms), which are used for spatial vision such as motion stereo, with finite processing on the sphere (see Sections **2** and **3**).

Two types of algorithms using this projection onto the sphere have been reported for motion stereo. One is an algorithm for detecting three parameters of a plane in Figure 1 (i.e. the time-to-contact to the plane, its 3D direction, and its shortest distance) (Kawakami et al., 2003, 2000): this paper has detailed the algorithm from a different point of view. The other is an algorithm for detecting two parameters of a straight line in space (i.e. the shortest distance to the line and its 3D orientation) (Sugie et al., 2006; Sugie, 2007; Kawakami, 1 987; Morita et al., 1989).

As described above, it has been clarified that the models for space recognition (see Figures 9, 17, and 30) are processed on a spherical surface. In addition, it was reported that a model for form recognition (more precisely, for curvature detection, which plays an important role in form recognition) is also processed on a spherical surface (Kawakami et al., 2020). Therefore, it has been clarified that all cell types that make up four types of model for the space and form recognitions (see Figures 19 and 20 in Kawakami et al. (2020)) are processed on a spherical surface.

## 5. Conclusion

Cells that detect the plane parameters of Figure 1 (i.e. the time-to-contact T to a plane, its 3D orientation n, and its shortest distance D) were modeled by dividing them into two. One is a series of cells that detect the time-to-contact T and 3D orientation n, and was modeled in Figure 9. The other is a series of cells that detect the shortest distance D and 3D orientation n, and was modeled in Figure 17.

The series of cells that detect local motions in Figures 9(B) and 17(B) are the same: these series were previously reported (Kawakami & Okamoto, 1996a, 1995a; Kawakami, 1996b). What is different in Figures 9 and 17 is the SDCs and SDC neural networks in Figures 9(C) and 17(C). That is, each SDC in Figure 9(C) performs the cross-ratio and polar transforms to detect the time-to-contact and 3D orientation (see Section **2**). Also, each SDC in Figure 17(C) performs the small-circle transform to detect the shortest distance and 3D orientation (see Section **3**). The SDC neural network that connects the MDCs in Figure 9(B) (or 17(B)) and the SDCs in Figure 9(C) (or 17(C)) was accurately modeled one by one with mathematical equations in Sections **2.2** (or **3.2**). These equations are quite complex, and you may think that it is difficult for cerebral cells to execute them. However, Section **2.2**(12) showed that this worry is not a problem.

In order to confirm the validity of these modeled cells in Figure 9 (or Figure 17), a simulator consisting of about 16 (or 5) million cells and about 460 (or 120) million neural connections was created on a computer based on procedures in Supplementary materials. This simulator confirmed that these modeled cells detected correctly the parameters of the plane (i.e. the normalized time-to-contact t, 3D orientation n, and normalized shortest-distance d) (see Figures 13 and 19).

The selectivities of the SDCs to the parameters of planes were compared with those of the ethological and physiological experiments (Section **4.2**). The selectivities of SDCs were consistent with those of the experiments. This consistency clarified that the series of modeled cells, constituting Figures 9 and 17, actually exist and enable to explain these experiments neurophysiologically.

An equation for detecting the time-to-contact to an opponent (or object) was first proposed by Lee (1976, 1980). This equation was clarified to be a special case of the cross-ratio transform that was used for modeling the SDCs in Figure 9(C) (see Section **4.1**). Thus, this Lee’s equation, unlike our model of detecting planes, detects the time-to-contact to a point object near the center of the visual field, not a planar object.

The parameters for binocular stereo were clarified to be the same as those for motion stereo at each moment (see Section **4.4.1**). Based on this sameness, replacing the terms (such as cell names and parameters) for motion stereo shown in Figure17 to those for binocular stereo yielded a series of modeled cells (shown in Figure 30) that detects binocularly the parameters of a plane (see Section **4.4.4**). Section **4.4.6** showed that this series actually existed in the cerebrum.

There are two methods to create the SDC neural networks of Figure 9(C) (or Figure 17(C)), each of which connects the MDCs and the SDC column (see Section **4.3**): these methods were named the forward and backward methods. The forward method is how to create forwardly the SDC neural network from each MDC to the SDC column, and the backward method is how to create backwardly the network from each SDC of the SDC column to MDCs. These two types of method need to be used properly according to their characteristics, as follows (see Section **4.3.3.1**(3) (or **4.3.3.2**(3))). That is, the forward method is suitable for explaining the mathematical structure of the neural network, and the backward method, which is simple in procedure, is suitable for simulation.

The series of modeled cells in Figures 9 and 17 uses the Hough, inverse Hough, and polar transforms. These Hough and inverse Hough transforms were clarified to be a special case of the polar transform that was used for modeling the SDCs in Figure 9(C) (see Section **4.5.2**). Therefore, renaming the Hough and inverse Hough transforms to the polar transform allows this series of modeled cells to consist of a time delay, spatio-temporal correlation, accumulation, cross-ratio transform, small-circle transform, and three polar transforms.

Two psychological discoveries have been known for the detection of planes. One is the discovery of detecting the time-to-contact to a plane and its 3D orientation by motion stereo (Gibson, 1950). The other is the discovery of detecting the shortest distance to a plane and its 3D orientation by binocular stereo (Julesz, 1960, 1971, 1986). It is surprising that the series of modeled cells in Figures 9 and 30 enable these psychological discoveries to be explained neurophysiologically at the level of the neural connection of which cell is connected to which cell (see Sections **4.2.1** and **4.4.6.1**). These neurophysiological explanations are, to my knowledge, the first time except for our reports (Kawakami et al., 2003, 2000).

## Acknowledgements

I thank the following researchers for their joint creation of the first challenging simulators based on Supplementary materials. Namely, I thank Masahiro Matsuoka, Hiroaki Okamoto, and Sinya Hosogi for creating the SDC simulator, and Hiroaki Okamoto for creating the simulator of a series of cells from retinal cells to MDCs, adding that of a series of cells from retinal cells to BDCs. Also, I thank Hisanao Akima for precisely recreating all the simulators from retinal cells to SDCs, and for validating equation(4.3-1) by simulations. Furthermore, I thank Hisanao Akima, Shigeo Sato, Masafumi Yano and Koji Nakajima for discussing the creation of these simulators from the perspective of realizing this neural network with LSI.

I thank Hide-aki Saito and Eiki Hida for their joint experiments to validate electrophysiologically the modeled SDCs in area MST of monkeys. I would like to express my deep gratitude to Masafumi Yano and Kennichi Abe for his great support in improving my research environment at Tohoku University, and to Masafumi Yano for making it possible for me to continue my research by strongly supporting the treatment of my illness.

## Appendix Appendix 1

### Proof of the cross-ratio transform

Let us prove the cross-ratio transform of equation(1.0-4) using Figure A1 (see Kawakami et al., 2003, 2000).

The (A) is the same as Figure 8(A), and shows how an arbitrary point P_0_ on the eyeball moves. P_0_ moves to P_1_ and P_T_, and reaches P_∞_ at infinity time: since the P_∞_ is not visible behind the eyeball, - P_∞_ is indicated. This movement trajectory is shown as a blue great circle. Here, P_0_, P_1_, and P_T_ are the positions at times T_0_, T_1_, and T, respectively. P_∞_ is the direction opposite to the movement direction v.

The (B) shows a cross section of the eyeball passing both this movement trajectory and the eyeball center O_eye_. Corresponding to positions P_0_, P_1_ and P_T_ on the eyeball, positions Q_0_, Q_1_, and Q_T_ in space move. In fact, the opposite is true: the movement of Q_0_, Q_1_, and Q_T_ is reflected on the retina of the eyeball as P_0_, P_1_, and P_T_. Here, although the (B) appears to be a special case where the movement direction v is directly downward, in reality, this direction v is arbitrary as shown in the (A). This is because in the (A), the direction from the eyeball center O_eye_ toward the movement direction v has been displayed directly downward for easy viewing.

According to the fundamental theorem of projective geometry, cross ratios are invariant to the projective transformation including the central projection of the (B) (Yanaga & Hirano, 1959; Iwata, 1981). Thus, the cross ratio (P_∞_P_0_P_1_P_T_) on the eyeball is equal to the cross ratio (Q_∞_Q_0_Q_1_Q_T_) on the straight line, and thus equation(A1-1) is obtained: here, Q_∞_ is infinity and corresponds to the movement direction v. Based on the definition of cross ratio (Izumi et al., 1987: Iwata, 1981), rewriting equation(A1-1) using directed segments Q_∞_Q_1_, Q_0_Q_1_, etc. yields equation(A1-2).

Since Q_∞_ is infinity, (Q_∞_Q_1_/Q_∞_Q_T_) converges to 1, and thus equation(A1-2) is transformed into equation(A1-3). Furthermore, since Q_0_Q_1_=Vt_d_ and Q_0_Q_T_=V(T–T_0_), equation (A1-4) is obtained: V is the movement speed. Then, the velocity V cancels out and equation(A1-5) is obtained. Finally, using t=(T–T_0_)/t_d_ of equation(1.0-3), equation(A1-6) is obtained. Thus, the cross-ratio transform represented by equation(1.0-4) has been proved.

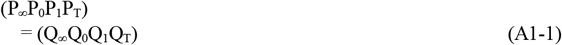

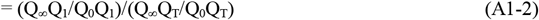

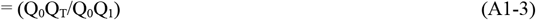

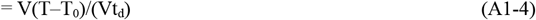

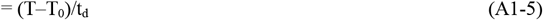

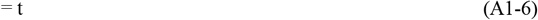

**Figure A1.**
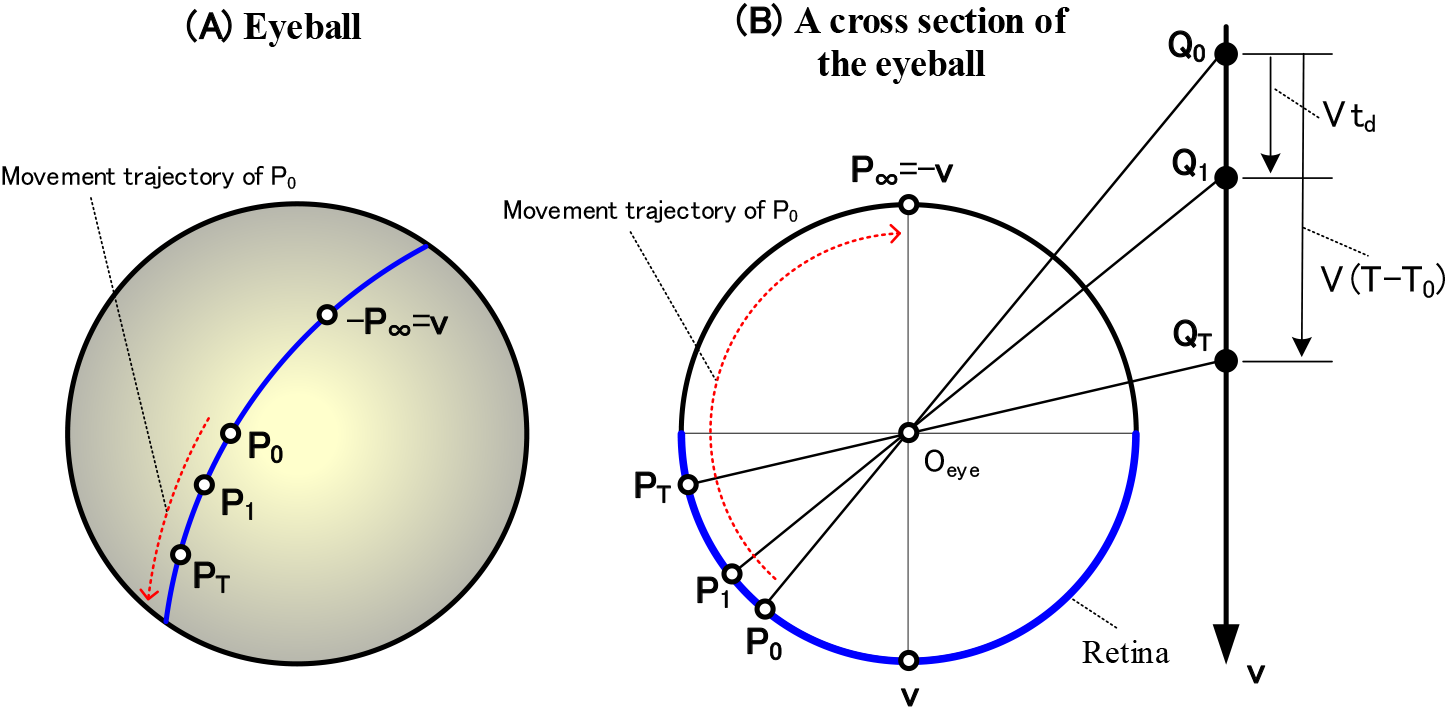
Proof of the cross-ratio transform. Parameters necessary for this proof are shown. (A) This is the same as Figure 8(A), and depicts how an arbitrary point P_0_ on the eyeball moves. The P_0_ moves to P_1_ and then P_T_, reaching P_∞_ after infinite time. The great circle, which is the trajectory of this movement, is drawn in blue. The P_∞_ is in the opposite direction to the movement direction v. (B) A cross section of the eyeball passing both the movement trajectory ((A)) and the eyeball center O_eye_ is shown. Movements Q_0_, Q_1_, and Q_T_ in space are reflected on the retina of the eyeball as P_0_, P_1_, and P_T_. Since the cross ratio is invariant to a projective transformation by the fundamental theorem of projective geometry, (P_∞_P_0_ P_1_ P_T_)=(Q_∞_Q_0_ Q_1_ Q_T_). Deforming it, equation(1.0-4) is obtained, and the cross-ratio transform has been proved (see Appendix **1**). Here, Q_∞_ is infinity in the movement direction v.

A feature of this cross-ratio transform is that it is independent of the velocity V. This feature enables the normalized time-to-contact t of equation(1.0-3) and the normalized shortest distance d of equation(3.0-2) to be detected independently of the velocity (see Sections **2.4** and **3.4**).

## Appendix 2

Proof of equation(1.0-6)

To prove equation(1.0-6), it is necessary to go back to equation(1.0-4) of its root. However, since the cross ratio (P_∞_P_0_P_1_P_T_) in equation(1.0-4) is subscripted, the proof process is complicated and difficult to see. Therefore, let us make (P_∞_P_0_P_1_P_T_) a simple symbol (ABCD) and express the cross ratio (ABCD) as a central angle (α,β,γ) in Figure A2(A). This expression will be shown in Section **A2.1**. Using this expression, the cross ratio (P_∞_P_0_P_1_P_T_) in Figure A2(B) can be expressed by the central angle (a, x, and τ_2D_), and thus equation(1.0-6) can be proved in Section **A2.2**.

### A2.1. Expression of the cross ratio (ABCD) on the eyeball using a central angle (α, β, and γ)

A cross ratio (ABCD) on the eyeball is projected on a straight line to obtain the cross ratio (abcd) (see Figure A2(A)). Since the cross ratio is invariant to the projection, equation(A2-1) is obtained. According to the definition of the cross ratio, equation(A2-2) is obtained by describing it in terms of directed segments (ac, bc, etc.), and equation(A2-3) is obtained by deforming it.

Since the triangles acp and adq are similar, ac/ad=cp/dq is obtained, and since the triangles bcr and bds are similar, bc/bd=cr/ds is obtained. Substituting these relationships into equation(A2-3) yields equation(A2-4).

Then, cp=cO_eye_ sin(α+β) is obtained by focusing on the right triangle cpO_eye_, dq=dO_eye_ sin (α+β+γ) is obtained by focusing on the right triangle dqO_eye_, cr=cO_eye_ sinβ is obtained by focusing on the right triangle crO_eye_, and ds=dO_eye_ sin(β+γ) is obtained by focusing on the right triangle dsO_eye_. Equation(A2-5) is obtained by substituting these relationships into equation(A2-4). By arranging it, equation(A2-6) is obtained. Thus, the cross ratio (ABCD) has been expressed using the central angle (α, β, and γ).

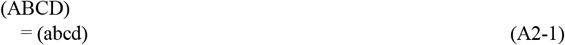

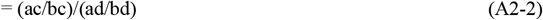

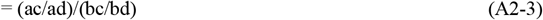

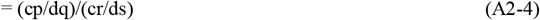

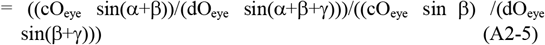

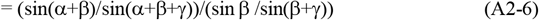

#### A2.2. Proof of equation(1.0-6)

Let us express the cross ratio (P_∞_P_0_P_1_P_T_) of equation(1.0-4) using the central angles (a, x, and τ_2D_) in Figure 8(B), as follows. The (B) shows a cross section of the eyeball passing both the blue moving trajectory in Figure 8(A) and the eyeball center O_eye_, and corresponds to Figure 8(B), except that the position of P_∞_ is different. The central angles and positions of the (B) correspond to those of the (A).

Comparing the central angles (α, β, and γ) of the (A) and the central angles (a, x, and τ_2D_) of the (B) yields a relationship of α=a, β=τ_2D_, and γ=x-a-τ_2D_. Rewriting the variables in equation (A2-6) using this relationship yields a relationship of α+β=a+τ_2D_, α+β+γ=x, β=τ_2D_, and β+γ=x-a. Substituting this relationship into equation (A2-6) and arranging it gives the following equation.

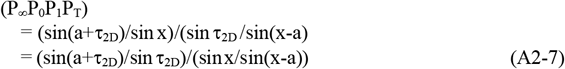

Therefore, the cross ratio (P_∞_P_0_P_1_P_T_) has been expressed using the central angles (a, x, and τ_2D_), and thus equation(1.0-6) has been proved by substituting equation(A2-7) into equation(1.0-4). In addition, the reason why the sine functions are necessary has been shown by equation(A2-5).

## Appendix 3

Great circle and its equidistant projection

### A3.1. Mathematical representation of a great circle

Figure A3 shows the eyeball viewed from directly above, where O is the center of the visual field: this figure was obtained by adding parameters necessary for the mathematical expression to Figure 10(A). Consider a great circle g_P_ on the eyeball. The pole P of this great circle is expressed as (α_P_,β_P_) in polar coordinates: the pole corresponds to the north pole when the great circle is the equator. In addition, express any point Q on the great circle as (α_Q_,β_Q_) in polar coordinates. Applying the cosine rule (Bronshtein & Semendyayev, 1978; Izumi et al., 1987) to the spherical triangle QPO gives the following equation.

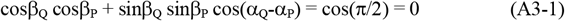

This is the mathematical expression of the great circle, and the trajectory of (α_Q_,β_Q_) represents the great circle. Here, the radius of the eyeball is 1, and π/2 is the radius of the great circle. Note that the conversion from the pole P to the polar line (i.e. the great circle g_P_) or vice versa (i.e. the conversion from the polar line g_P_ to the pole P) is called the polar transform (Gurevic, 1962).

### A3.2. Mathematical representation for the equidistant projection of the great circle

Using the equidistant projection in Section **2.1.3**, let us convert this great circle on the eyeball of Figure 10(A) (equal to Figure A3) into a curve on the disk of Figure 10(B). By the definition of this projection, the polar coordinates (i.e. (α_P_,β_P_) and (α_Q_,β_Q_)) of the above two points P and Q on the eyeball are the same as those on the disk. Therefore, the curve on the disk (Figure 10(B)), into which the great circle is converted by this projection, is expressed by the same equation(A3-1) as the great circle on the eyeball.

**Figure A2.**
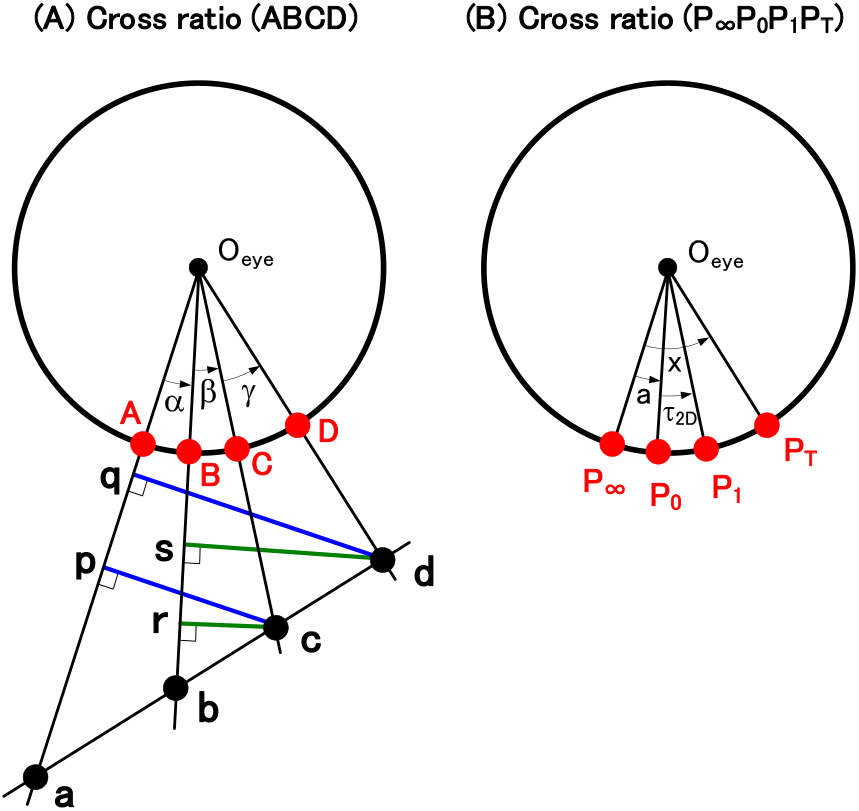
Proof of equation(1.0-6). Parameters necessary for this proof are shown. (A) The cross ratio (ABCD) on the eyeball is projected onto a straight line to obtain the ratio (abcd). Since the cross ratio is invariant to the projective transformation, (ABCD)=(abcd). By deforming it, the cross ratio (ABCD) is expressed as in equation(A2-6) of Appendix **2** using the central angles (α, β, and γ). (B) Using this equation(A2-6), the cross ratio (P_∞_P_0_P_1_P_T_) is expressed by the central angles (a, x, and τ_2D_), thereby proving equation(1.0-6).

**Figure A3.**
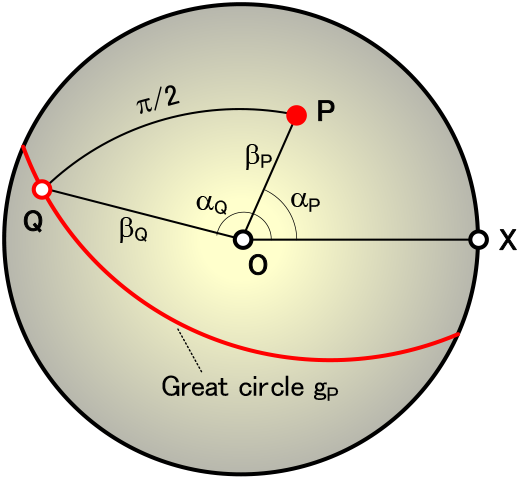
Mathematical representation of a great circle. Let us express a great circle on the eyeball with a mathematical expression. This figure was obtained by adding the necessary parameters to Figure 10(A). The pole P of the great circle and any point Q on this great circle are expressed in polar coordinates as (α_P_,β_P_) and (α_Q_,β_Q_), respectively. Applying the cosine rule to a spherical triangle QPO, the mathematical expression of the great circle is obtained as equation(A3-1).

Since the curve is thus expressed by the same equation(A3-1), this curve on the disk will be called a great circle in the same way as that on the eyeball.

Note that this can be generalized as follows. When an arbitrary figure on the eyeball is expressed in polar coordinates, the figure on a disk converted by the equidistant projection is also expressed by the same equation. This property is the same as in Section **A2** of Appendix **A** of Kawakami et al. (2020).

## Appendix 4

Correspondence between local motions on the eyeball and the MDC array, and various relationships

### A4.1. Correspondence between local motions on the eyeball and the MDC array

The following describes how the local motion τ_2D_ on the eyeball in Figure 11(A) is detected by the MDC array (τ_X_,τ_Y_) in Figure 9(B)(f). This motion τ_2D_ (i.e. the motion vector from the current position P_0_ to the next position P_1_) is shown in Figure A4(A) that is the same as Figure 11(A).

The process in which this local motion τ_2D_ on the eyeball of Figure A4(A) is detected as the coordinate (τ_X_,τ_Y_) of the MDC array of Figure A4(D) is shown as follows.

First, change the (A) (displayed centering on the center of the visual field O) to the (B) (displayed centering on the RF center O_IJ_): here, the RF is shown large for easy viewing, but its actual diameter is less than 5 degrees. Next, in the (C), display a tangent plane at the RF center O_IJ_, in which the local motion from P_0_ to P_1_ (i.e. its magnitude τ_2D_ and direction ϕ) is shown: here, Γ_ϕ_ is the movement direction from P_0_ to P_1_. Finally, arrange MDCs in this tangent plane ((D)). This arrangement corresponds to the MDC array (τ_X_,τ_Y_) in Figure 9(B)(f). About 1000 MDCs are arranged in this array. A MDC at position P_1_ fires to detect the local motion (τ_2D_,ϕ) based on its coordinate (τ_X_,τ_Y_), where the following relationship exists between the polar coordinate (τ_2D_,ϕ) and the orthogonal coordinate (τ_X_,τ_Y_).

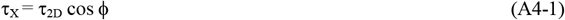

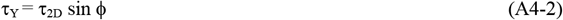

**Figure A4.**
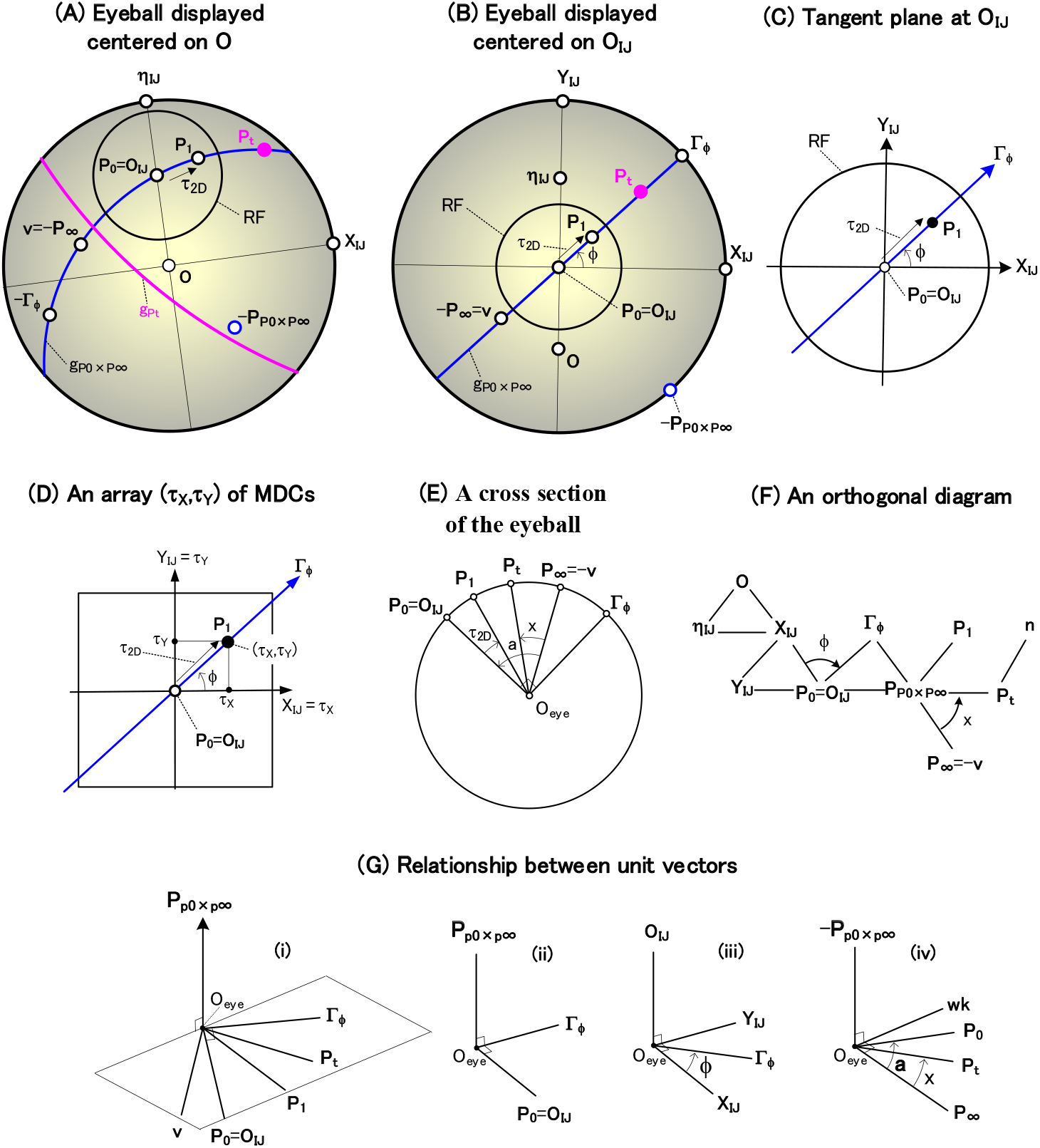
Relationships between the local motion τ_2D_ on the eyeball and the MDC array (τ_X_,τ_Y_). (A) This is the same as Figure11(A). How this local motion τ_2D_ on the eyeball (i.e. this motion on the eyeball from the position P_0_ at a current time to the position P_1_ at the next time) is detected by the MDC array (τ_X_,τ_Y_) of Figure 9(B)(f) is described below. (B) First, change the (A) to a display centered on the RF center O_IJ_. (C) Next, display a tangent plane in contact with O_IJ_ of the (B). (D) Finally, arrange MDCs on this tangent plane. This arrangement represents the MDC array (τ_X_,τ_Y_) shown in Figure 9(B)(f). In this way, the local-motion τ_2D_ of the (A) on the eyeball, mentioned above, is detected as the coordinate (τ_X_,τ_Y_) in this MDC array. (E) A cross section of the eyeball ((A)) passing both the great circle g_P0×P∞_ and the eyeball center O_eye_ is shown. It is used to express the cross-ratio transform as equation(1.0-6) using central angles (a, x, and τ_2D_). (F) This is called an orthogonal diagram and shows the orthogonal relationships between the various unit-vectors in the (A). Each straight line indicates that the vectors at both ends of that line are orthogonal. Appendix **5** explains how to use this diagram. (G) The orthogonal relationship of unit vectors and the angles (a, x, and ϕ) between them are shown. This is useful for deriving equations(A4-5, 6, 7, and 12).

Therefore, it has been clarified that each local motion τ_2D_ in a RF O_IJ_ on the eyeball ((A)) is detected as the coordinate (τ_X_,τ_Y_) of the corresponding MDC ((D)) on a tangent plane that touches the eyeball at the RF center O_IJ_. Here, note the following. Since this local motion is detected wherever P_0_ is within the RF (see Section **2.1.2**(6)), P_0_ can be considered to be at the RF center O_IJ_ and thus is expressed by the following equation.

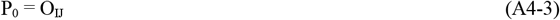

The parameters used in the (A) to (D) are described as follows. First, the coordinate axis X_IJ_ of the MDC array in the (D) is calculated by the following equation using a vector product.

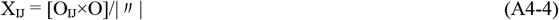

Here, this X_IJ_ has been defined as the pole of a great circle that passes through O_IJ_ and O in the (A). Then, the coordinate axis Y_IJ_ orthogonal to X_IJ_ is calculated by the following equation.

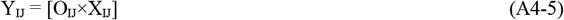

Here,|″| is the absolute value of [O_IJ_×O], and is used to normalize X_IJ_ to a unit vector.

In addition, the method for calculating the two important parameters (i.e. P_P0×P∞_ and Γ_ϕ_) is described as follows (see (B)). First, P_P0×P∞_ is calculated by the following equation as the pole of the great circle that passes through P_0_ and P_∞_.

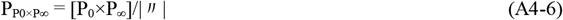

Here, this P_P0×P∞_ determines the blue movement trajectory g_P0×P∞_ as its polar line. The other is Γ_ϕ_ in the (B), which indicates the movement direction ϕ of P_0_. This Γ_ϕ_ is calculated by the following equation (see (G)(ii)).

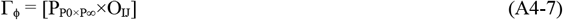

Also, the position P_∞_ that P_0_ reaches after an infinite time is calculated by the following equation (see equation(1.0-1)), where v is the movement direction.

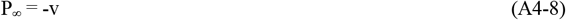

Other figures are described below. The (E) shows a cross section of the eyeball passing both the great circle g_P0×P∞_ of the (A) and the eyeball center O_eye_. This cross section is used for expressing the cross-ratio transform by central angles (a, x, and τ_2D_) (see equation(1.0-6)).

The (F) is named an orthogonal diagram and shows the orthogonal relationship between the various unit vectors of the (A): each straight line indicates that the vectors at both ends of that line are orthogonal. This diagram is useful in two ways. First, a complex relationship between vectors such as the (A) can be expressed exactly as an orthogonal relationship between all vectors by only checking whether each two vectors are orthogonal. Second, as shown in Appendix **5**, even if a vector c is unknown, if you know two vectors a and b that are orthogonal to the vector c, you can easily calculate the vector c using the following equation.

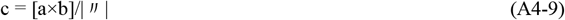

Calculating this unknown vector as described above enables a next unknown vector to be calculated one after another to expand the calculation range of the orthogonal diagram.

Knowing angles ϕ and x in the (F) can be helpful as follows. First, knowing the angle ϕ and the vectors X_IJ_ and O_IJ_ enables an unknown vector Γ_ϕ_ to be calculated by equations(A4-12) (see (G(iii)). Next, knowing the angle x and the vectors P_∞_ and P_P0×P∞_ enables an unknown vector P_t_ to be calculated by equations(2.2-8 and 9) (see (G(iv)). In this way, by adding these known angles to the orthogonal diagram, the computable range of this diagram can be further expanded.

This orthogonal diagram was devised by the author in 1980 to create a control algorithm for multi-articulated robots that consist of many vectors, and has been applied to this series of modeled cells (Figure 9 or 17) that consists of many vectors such as the (A).

The (G) shows the orthogonal relationship between vectors used above and the angles (a, x, and ϕ) between these vectors, and was used to derive equations(A4-5, 6, 7, 12, etc.).

### A4.2. Various relationships

The various relationships used in Figure A4 are summarized below. The sign{a} is a function that determines the sign, and if a is positive, it is 1, if it is negative, it is -1.

1. Basic coordinate system

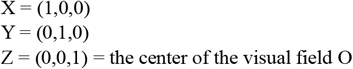
2. η_IJ_ related to each RF coordinate (see (A))

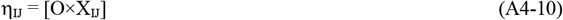
3. Γ_ϕ_ related to the movement direction from P_0_ to P_1_ (see (F) and G(iii))

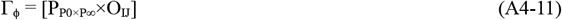

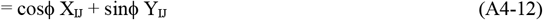
4. ϕ related to the MDC, and τ_2D_ related to cross-ratio transform Based on G(iii), ϕ is calculated by the following equation.

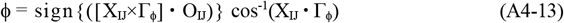 Solving equation(1.0-6) for τ_2D_ yields the following equation.

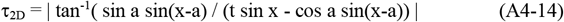

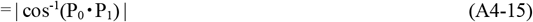 Here, a and x used above are calculated by the following equations (see equations(1.0-7 and 8)).

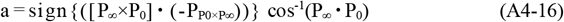

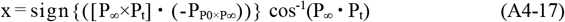

## Appendix 5

How to use orthogonal diagrams

Even if you do not know a certain vector c, if you know two vectors a and b orthogonal to the vector c, you can draw the polar transformations of the vectors a and b on the eyeball and can determine the vector c, as follows (Figure A5(A)). That is, drawing great circles g_a_ and g_b_ (obtained by the polar transformations of these vectors) enables the vector c to be determined as the intersection of these great circles. This is a geometrical method that uses the drawing. This method enables unknown vectors to be determined while confirming the geometrical image, but requires the tedious work of drawing the great circles.

**Figure A5.**
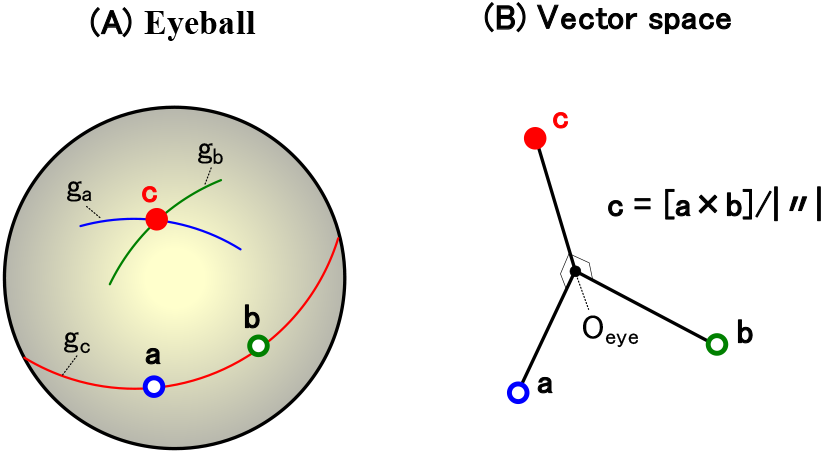
How to use orthogonal diagrams such as Figure A4(F). (A) This shows a geometrical method, as follows. Even if a certain vector c is unknown, if two vectors a and b orthogonal to the vector c are known, it is possible to obtain the vector c by drawing the vectors on the sphere. That is, drawing great circles g_a_ and g_b_ (obtained by the polar transformations of the vectors a and b) enables the vector c to be determined as the intersection of these great circles. (B) This shows an algebraic method, as follows. The unit vectors a, b, and c of the (A) are extracted and the relationship between these vectors is shown. Assume that the vectors a and b, orthogonal to the vector c, are known. The vector c can be calculated as c=[a×b]/|″| using the vector product. Appendix 5 will show that this algebraic method is simpler than the geometrical method in the (A). Orthogonal diagrams such as Figure A4(F) enables this algebraic method to be used effectively.

A algebraic method shown in the (B) is simpler than this geometrical method in the (A), as follows. Extracting the unit vectors a, b, and c from the (A), the relationship between these vectors is shown in the (B). Knowing vectors a and b orthogonal to a vector c enables this unknown vector c to be easily calculated as the following vector product.

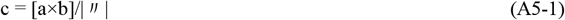

This algebraic method can be used effectively if there is the orthogonal diagram such as Figure A4(F). That is, each unknown vector in the diagram can be determined as a vector product between any two known vectors that are orthogonal to that vector. Therefore, this method for determining an unknown vector enables next unknown vectors in the diagram to be calculated one after another. In this way, this method allows us to successively reduce unknown vectors in the diagram.

## Appendix 6

Derivation of the Small-Circle Transform

### A6.1. Derivation of the small-circle transform

The small-circle transform of Figure 16(A) is derived as follows: this derivation elaborates the previous report (Kawakami et al., 2003, 2000). This transform converts the position P_0_ at a current time to a small circle with radius R centered on P_0_: the radius R is determined by equation(3.0-4). Also, this transform has an important property that the 3D orientation n of a plane to be determined lies on this small circle.

In order to make the derivation process easier to see, Figure 16(A) was changed in two points to obtain Figure A6-1. First, change the display centered on the visual field center O in Figure 16(A) to a display centered on the movement direction v. Second, enlarge the size of the pink small-circle for easier viewing, change the positions of P_0_ and P_t_, and omitt P_1_.

Consider a blue small-circle with radius r centered on v, and assume that the 3D orientation n of a plane to be determined exists on this circle. Since (n•v) in equation(3.0-3) is cos r, this equation is deformed into d=t cos r. Thus, using the following equation, convert the normalized shortest-distance d that is given to a normalized time-to-contact t, where r is currently an unknown variable.

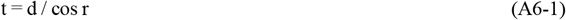

Therefore, substituting this conversion into the cross-ratio transform of equation(1.0-11) enables the central angle x (corresponding to the position P_t_ on the eyeball at a time t) to be calculated by the following equation.

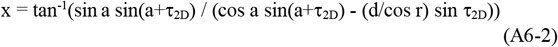

Based on this x, P_t_ is calculated by equation(2.2-8). Then, draw a green great-circle g_Pt_ on the eyeball, which is obtained as the polar transformation of this P_t_. This drawing indicates that the 3D orientation n of a plane to be determined exists on this great circle. On the other hand, above we assumed that this orientation lies on the blue small-circle with radius r. Therefore, points that satisfies these two conditions (or assumptions) are the 3D orientations n that should be determined. That is, the pink intersections n_+_ and n_-_ of the green great-circle g_Pt_ and the blue small-circle satisfy these conditions, and thus the 3D orientations n are determined as these intersections.

Calculate this intersection as follows. First, applying the cosine rule to a spherical triangle n_+_P_t_v yields the following equation based on the spherical trigonometry (Bronshtein & Semendyayev, 1978; Izumi et al., 1987).

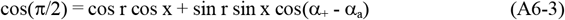

**Figure A6-1.**
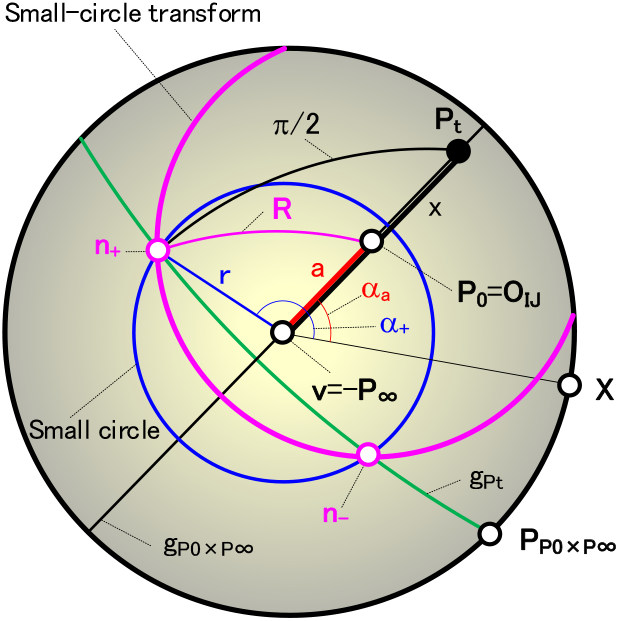
Derivation of the small-circle transform. In order to make the derivation process easier to see, Figure 16(A) showing this transform was rewritten at two points to obtain this figure. First, the display centered on the visual field center O of Figure 16(A) was rewritten to a display centered on the movement direction v. Second, the size of the small circle was enlarged to make it easier to see, the positions of P_0_ and P_t_ were changed, and the position P_1_ was omitted. Let us derive the small-circle transform as follows. First, consider a blue small-circle with radius r centered at v, and assume that the 3D orientation n of a plane exists on this small circle. Next, a position P_t_ corresponding this radius r can be calculated using equation(A6-2) in Appendix 6. Drawing a green great-circle g_Pt_ obtained by the polar transformation of this P_t_ causes the 3D orientation n of a plane to exist on this great circle. Therefore, pink two points satisfying these two considerations (i.e. pink intersections n_+_ and n_-_ of the green great-circle g_Pt_ and the blue small-circle) are the 3D orientations n of the plane. In this way, the small-circle transform of equation(A6-6) has been derived, which is described in Section **A6.1** of Appendix **6**.

Here, the polar coordinates of n_+_ and P_0_ are shown as (r,α_+_) and (a,α_a_), respectively. Substituting equation(A6-3) into equation(A6-2) to erase x and further deforming it, express the intersection point (r,α_+_) (i.e. the n_+_) as the following equation that represents a curve consisting of the intersections n_+_ and n_-_.

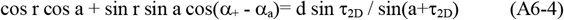

Thus, the unknown variable r above can be determined by solving this equation for α_+_. On the other hand, applying the cosine rule to the spherical triangle n_+_P_0_v, express a pink small circle with radius R centered on P_0_ as the following equation.

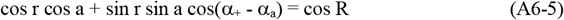

By comparing equations(A6-4 and 5), it has been clarified that equation(A6-4) consisting of the intersections n_+_ and n_-_ is a mathematical expression of a small circle with radius R centered on P_0_, where this comparison also enables the radius R to be determined by the following equation.

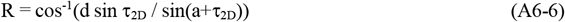

Rewriting equation(A6-6) using the positions P_0_, P_1_, and P_∞_ on the eyeball yields equations(A6-7 and 8) (see equation(A2-6) of Appendix **2** for requiring sine functions): (P_∞_P_0_P_1_) is a simple ratio of P_∞_, P_0_, and P_1_. Therefore, the small-circle transform of equation(3.0-4) has been derived.

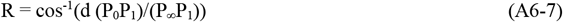

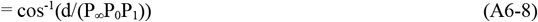

In this way, the 3D orientation n of a plane to be determined has been shown to lie on this small circle with radius R centered on P_0_.

### A6.2. Derivation of the small-circle transform from a geometrical perspective

Another derivation is given below using Figure A6-2, from a different geometrical point of view than Section **A6.1**.

#### A6.2.1. Derivation of the normalized distance d_0_ to the position Q_0_ at a current time

Figure A6-2(A) shows a cross section passing through P_0_ and P_1_ in Figure 16(A). A point Q_0_ in space moves in the direction of P_∞_ at a speed V and reaches Q_1_ after a unit time t_d_. The Q_0_ and Q_1_ are reflected as P_0_ and P_1_ on the eyeball. The distance between the O_eye_ and Q_0_ is D_0_, and the distance between Q_0_ and Q_1_ is Vt_d_. The P_∞_ is the direction opposite to the movement direction v.

Applying the sine rule to the triangle Q_0_O_eye_Q_1_ yields the following equation.

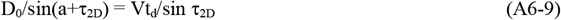

A normalized distance d_0_ is represented as equation(A6-10) based on the definition of equation(3.0-2). Substituting equation(A6-9) into equation(A6-10) yields equation(A6-11), expressing by directed segments of P_∞_P_0_ and P_0_P_1_ on the eyeball yields equation(A6-12), and using simple ratio (P_∞_P_0_P_1_) yields equation(A6-13). In this way, the distance d_0_ has been derived.

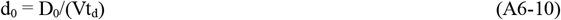

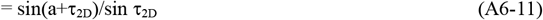

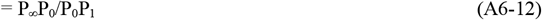

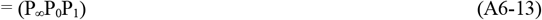

#### A6.2.2. Derivation of a radius R used by the small-circle transform

Figure A6-2(B) shows a cross section passing through the P_0_ and 3D orientation n in Figure 16(A). The Q_n_ is the perpendicular foot to a plane, and its reflection on the eyeball is the vector n. A radius R to be derived is the angle at which the vectors P_0_ and n are viewed from the O_eye_ (see Figure A6-1). Also, the distance from the O_eye_ to Q_n_ (i.e. the shortest distance from the O_eye_ to the plane) is D_n_.

**Figure A6-2.**
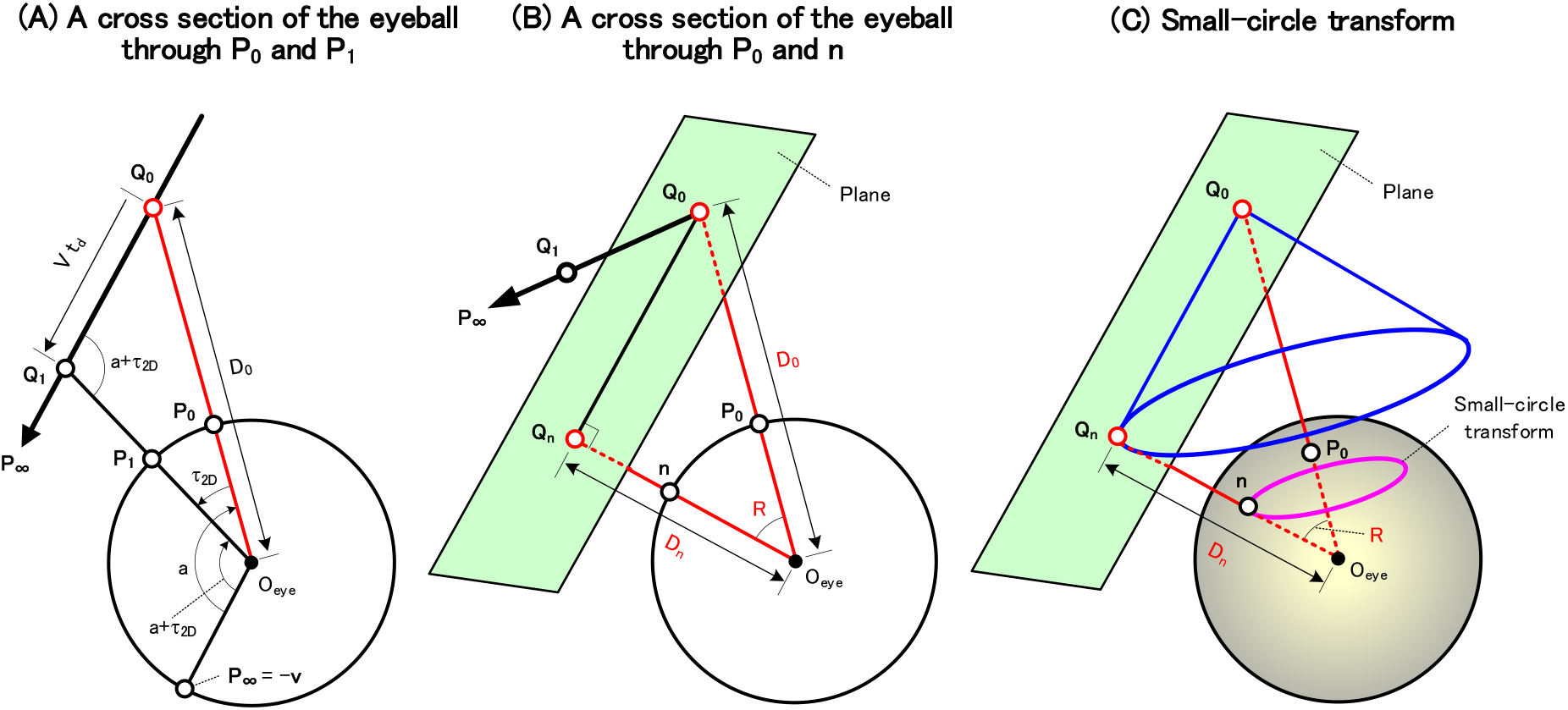
Another derivation of the small-circle transform. From a geometrical point of view, another derivation can be given. Parameters necessary for this derivation are shown. (A) A cross section of the eyeball in Figure 16(A) passing through P_0_ and P_1_ is shown. A point Q_0_ in space moves with velocity V in the direction of P_∞_ and reaches Q_1_ after the unit time t_d_. The P_0_ and P_1_ on the eyeball are the positions where Q_0_ and Q_1_ are reflected on it. The distance between O_eye_ and Q_0_ is D_0_, and the distance between Q_0_ and Q_1_ is Vt_d_. The P_∞_ is in the opposite direction to the movement direction v. A relationship of equation(A6-9) in Appendix **6** can be obtained using the central angle (τ_2D_, a, and a+τ_2D_). (B) A cross section of the eyeball in Figure 16(A) passing through the P_0_ and the 3D orientation n is shown. The Q_n_ is the perpendicular foot to a plane, and the n is the position where Q_n_ is reflected on the eyeball. The distance from the O_eye_ to Q_n_ is D_n_. (C) The small circle transform is derived as follows. A curved surface enveloped by the green planes forms a blue right cone with Q_0_ as its apex. The base of this right cone is the locus of Q_n_, which is shown as the blue circle in space. Projecting this circle onto the eyeball generates a pink small-circle with radius R centered at P_0_. Thus, transforming the P_0_ into this pink small-circle yields the small-circle transform. In this way, a geometrical derivation of the small-circle transform can be obtained.

Focusing on the right triangle Q_0_O_eye_Q_n_ yields the following equation.

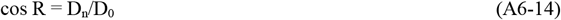

Replacing d and D in equation(3.0-2) with d_n_ and D_n_, respectively, yields a relationship d_n_ = D_n_/(Vt_d_). Substituting this relationship and equation(A6-10) into equation(A6-14) yields the relationship cosR=d_n_/d_0_. Deforming this relationship yields equation(A6-15), and substituting equation(A6-11) into equation(A6-15) yields equation(A6-16). This is the same as equation(A6-6) and thus the radius R used by the small-circle transform has been derived geometrically.

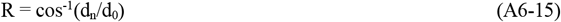

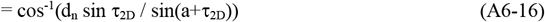

Notice that the cross-ratio transform of equation(1.0-11) has been not used to derive this radius R of equation(A6-16). This is different from the Section **A6.1** where the cross-ratio transform was used in equation(A6-2).

#### A6.2.3. Derivation of the small-circle transform

Let us derive below the small-circle transform using Figure A6-2(C). A blue right cone with Q_0_ as the vertex is formed by a curved surface that is enveloped by the green planes shown in the (B). The base of this right cone is the locus of the point Q_n_ in the (B) and constitutes a blue circle in space. Projecting this circle onto the eyeball yields a pink small-circle with radius R centered on P_0_: this radius R was calculated by equation(A6-16). A transformation that converts P_0_ into this pink small circle represents the small-circle transform, because this circle corresponds to the blue right cone above. Thus, the small-circle transform has been derived geometrically.

In the above, it was considered the case where the green plane is away from the eyeball for ease of viewing (see Figures A6-2(B) and (C)). Here, let us assume that this plane is in contact with the eyeball. This assumption causes its contact Q_n_ (i.e. its perpendicular foot Q_n_) to be equal to the 3D orientation n on the eyeball, and thus causes the green plane in the (C) to touch the eyeball at the point n. Thus, it has been clarified that the small circle transform is equivalent to (or corresponds to) the transformation of the point P_0_ into a right cone that is composed of this plane touching the eyeball at position n: the angle R at which the positions P_0_ and n are viewed from the O_eye_ is calculated by (A6-16). The pink small circle of (C) used for the small circle transform is obtained as the locus of the contact point n between this right cone and the eyeball.

In this way, the small circle transform has been derived from a geometric perspective, without using the cross-ratio transform of equation(1.0-11).

## Appendix 7

Small circle and its equidistant projection

### A7.1. Mathematical representation of a small circle

Figure A7 shows the eyeball viewed from directly above, where O is the center of the visual field. Consider a small circle with radius R centered on P and any point S on that small circle. P and S are expressed in polar coordinates as (α_P_,β_P_) and (α_S_,β_S_), respectively. Applying the cosine rule to the spherical triangle SPO (Bronshtein & Semendyayev, 1978; Izumi et al., 1987) yields the following equation.

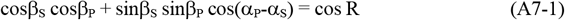

This is the mathematical expression of the small circle on the eyeball, and the trajectory of (α_S_,β_S_) represents the small circle.

### A7.2. Mathematical representation for the equidistant projection of a small circle

Let us convert this small circle on the eyeball (Figure A7) into a curve on the disk: this conversion to the curve is performed by the equidistant projection of Figure 10 (see Section **2.1.3**). By the definition of this projection, the polar coordinates (i.e. (α_P_,β_P_) and (α_S_,β_S_)) of the above two points P and S on the eyeball are the same as those on the disk. Therefore, this curve on the disk, into which the small circle is converted by this projection, is expressed by the same equation(A7-1) as the small circle on the eyeball.

Since the curve is thus expressed above by the same equation(A7-1), this curve on the disk will be called a small circle in the same way as that on the eyeball.

## Appendix 8

Geometrical derivation of Lee’s time-to-contact detection

Let us derive Lee’s time-to-contact detection of equation(4.1-1) geometrically, without using derivatives.

Figure A8 shows a cross section of the eyeball passing a point Q_0_ and the eyeball center O_eye_, where the point Q_0_ moves in the direction v at the velocity V. The Q_0_ moves to Q_1_ at the unit time t_d_, and the Q_0_ and Q_1_ are reflected on the eyeball as the P_0_ and P_1_. The P_0_ reaches P_∞_ in infinite time. The P_0_ and P_1_ are expressed as central angles a and τ_2D_, and the central angle b of Figure 21(A) is added to them.

Applying the sine rule to the triangle Q_0_O_eye_Q_1_, the following equation is obtained.

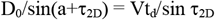

By deforming this, equation(A8-1) is obtained: note that this equation is the same as equations(A6-10 and 11). Since a=π-b, equation(A8-2) is obtained. Equation(A8-3) is obtained because b>>τ_2D_. Equation(A8-4) is obtained because b and τ_2D_ are small.

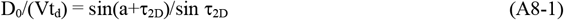

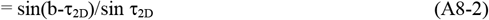

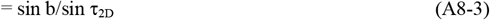

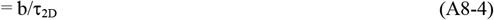

**Figure A7.**
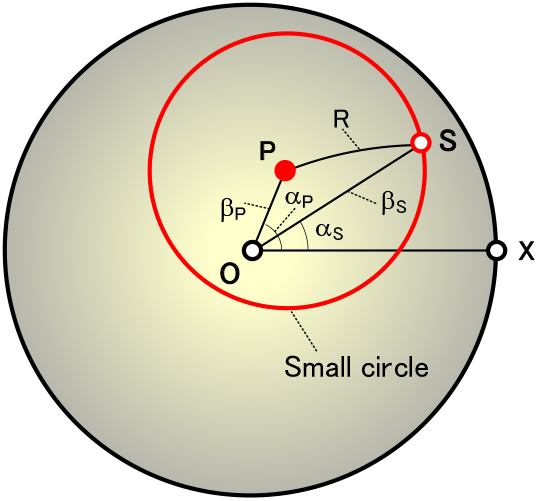
Mathematical representation of a small circle. This is drawn from directly above the eyeball: O is the visual field center. Consider a small circle with radius R centered at P and any point S on this small circle. Denote the P and S in polar coordinates as (α_P_,β_P_) and (α_S_,β_S_), respectively. Applying the cosine rule to a spherical triangle SPO enables the mathematical expression of the small circle to be obtained as equation(A7-1).

**Figure A8.**
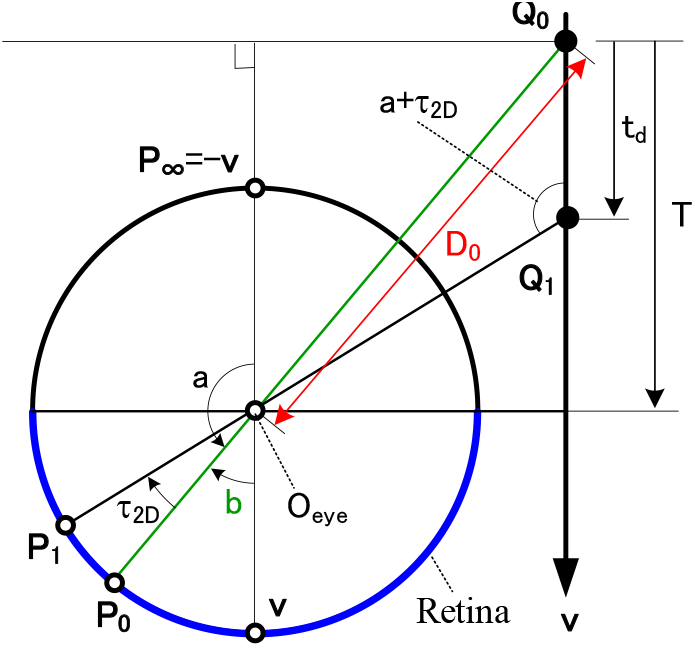
Another derivation of Lee’s time-to-contact detection. Parameters necessary for this derivation are shown. Considering the case where a point Q_0_ moves in the direction v at a speed V, we show a cross section of the eyeball passing through this point and the eyeball. A point Q_0_ moves to Q_1_ at the unit time t_d_, and these Q_0_ and Q_1_ are reflected on the eyeball as P_0_ and P_1_. The P_0_ reaches P_∞_ after infinite time. Represent P_0_ and P_1_ using central angles a and τ_2D_, and add the central angle b of Figure 21(A) to them. Applying the sine rule to the triangle Q_0_O_eye_Q_1_ and performing a deformation enables Lee’s time-to-contact detection to be geometrically derived as equation(A8-6) (see Appendix **8**).

Here, when b is small, it can be approximated as D_0_≒VT. Combining this approximation with equation(A8-4) yields equation(A8-5). Finally, since t_d_ is the time interval between P_0_ and P_1_, and lim_td→0_(τ_2D_/t_d_)=db/dt, equation (A8-6) is obtained: note that t is not the normalized time-to-contact of equation(1.0-3), but corresponds to T in Figure A8.

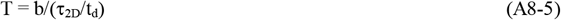

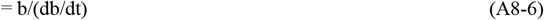

In this way, Lee’s time-to-contact detection expressed by equation(4.1-1) has been derived geometrically.

## Appendix 9

Representation of the backward method by mathematical equations

For the definitions of the forward and backward methods, see Figure 24 and Section **4.3**.

### A9.1. Detection of the normalized time-to-contact to a plane

The forward SDC neural network shown in Figure 24(A) was expressed in Section **2.2** by mathematical equations. Here, let us express the backward SDC neural network shown in Figure 24(B), using Figure A9, by mathematical equations: Figure A9 was obtained by adding a 3D orientation n and its great circle g_n_ to Figure 11(A) and by omitting the RF and great circle g_Pt_ in Figure 11(A).

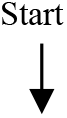

(1) Give a movement direction v (see Figure A9(A)) Calculate P_∞_ using the following equation (see equation(1.0-1)).

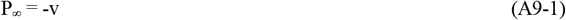 **Figure A9.**
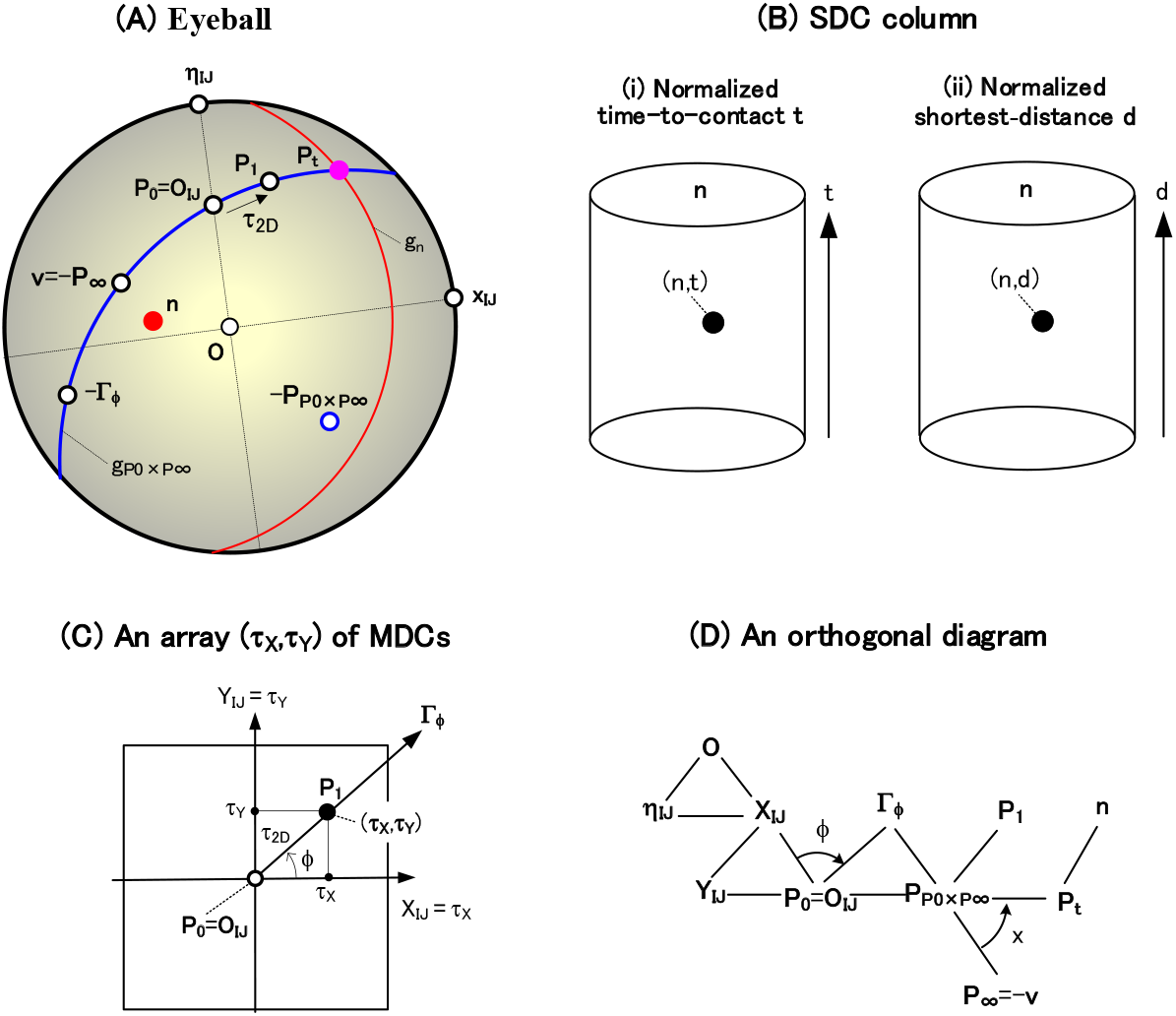
Detection of the time-to-contact and shortest-distance using the backward method. By using this figure, Sections **A9.1** and **A9.2** in Appendix **9** shows that a SDC neural network by the backward method can be expressed as mathematical equations. (A) This has been obtained by adding the following to Figure 11(A) that represents the forward method. That is, a red 3D orientation n and a red great circle g_n_ are added to it, although the great circle g_Pt_ used in Figure 11(A) is omitted. This circle g_n_ is the polar transformation of the position n. (B) Two types of SDC columns are shown. The (i) is a SDC column (n,t) for detecting the normalized time-to-contact t, and the (ii) is a SDC column (n,d) for detecting the normalized shortest-distance d. (C) This shows a MDC array (τ_X_,τ_Y_) of Figure 9B(f) (or Figure 17B(f)), where various parameters are added. The reason why a local motion τ_2D_ in the (A) is detected in this array was explained in Section **A4.1** of Appendix **4**. (D) This is the same as the orthogonal diagram in Figure A4(F) in Appendix **4**.
(2) Sweep a normalized time-to-contact t (see Figure A9(B)(i)) 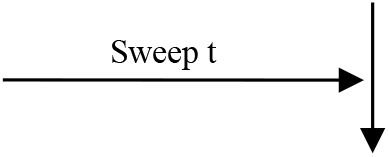
(3) Sweep the 3D orientation n of a plane (see Figure A9(B)(i)) 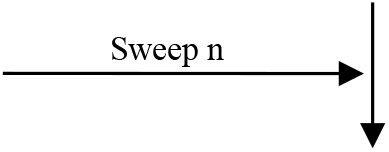
(4) Sweep a RF center O_IJ_ (see Figure A9(A)) 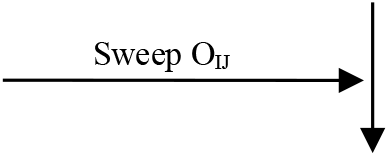 Calculate P_0_ using the following equation (see equation(A4-3) in Appendix **4**).

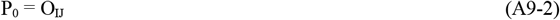
(5) Calculate P_t_ (see Figures A9(A) and (D), and Appendix **5**)

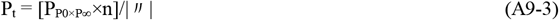 Here, P_P0×P∞_ used above is calculated by the following equation (see equation(A4-6))

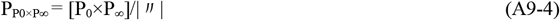
(6) Calculate the polar coordinate τ_2D_ of a MDC by the cross-ratio transform Using the cross-ratio transform of equation(1.0-4), the coordinate τ_2D_ is calculated by the following equation (see equation(A4-14) in Appendix **4**).

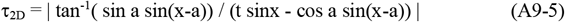 Here, the central angles a and x used above are calculated by the following equations (see equations(A4-16 and 17)).

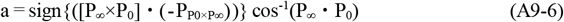

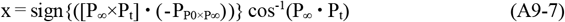
(7) Calculate the polar coordinate ϕ of the MDC (see equation(A4-13))

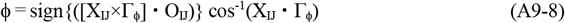 Here, X_IJ_ and Γ_ϕ_ used above are calculated by the following equations (see equations(A4-4 and 7)).

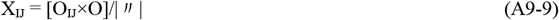

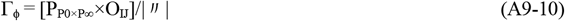
(8) Convert the polar coordinate (τ_2D_,ϕ) to the orthogonal coordinate (τ_X_,τ_Y_) by the following equations (see Figure A9(C))

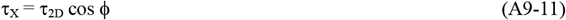

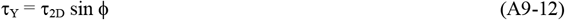
(9) Connect the SDC (n,t) to the MDC (τ_X_,τ_Y_) (see Figure 24(B)) 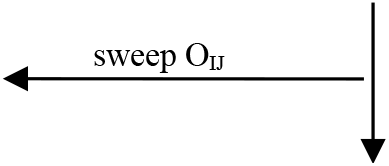
(10) The neural network for all RFs including red, green, and blue in Figure 24(B)(i) has been completed. 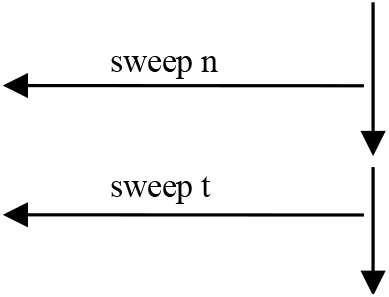
(11) The entire neural network backwardly connecting the SDC column to MDCs (Figure 24(B)(iii)) has been accurately expressed by mathematical equations. That is, each of the neural network that detects the normalized time-to-contacts and 3D orientations by the backward method has been expressed one by one by a mathematical equation. 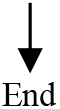

In this way, the neural network of Figure 9(C) (or Figure 24(B)(iii)) has been accurately expressed by mathematical equations at the level of the neural connection of which cell is connected to which cell. This neural connection is shown in detail as the backward connection table in Section **S2.3.1** of Supplementary materials.

The above is represented by the complex equations such as equation(A9-5), and you may think that it is difficult for cerebral cells to execute them. However, that worry is not a problem for the same reasons as in Section **2.2**(12).

### A9.2. Detection of the normalized shortest-distance to a plane

In order to consider the normalized shortest distance to a plane, let us replace the height axis t (of the SDC column (in Figures 24(A)(iii) and 24(B)(iii)) for the normalized time-to-contact with the height axis d for the normalized shortest distance. After making such a replacement, let us explain the following.

The forward SDC neural network shown in Figure 24(A) was expressed in Section **3.2** by mathematical equations. Here, let us express the backward SDC neural network shown in Figure 24(B), using Figure A9, by mathematical equations.

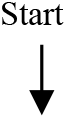

(1) Give a movement direction v (see Figure A9(A)) Calculate P_∞_ using the following equation (see equation(1.0-1)).

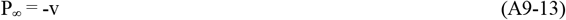
(2) Sweep a normalized shortest-distance d (see Figure A9(B)(ii)) 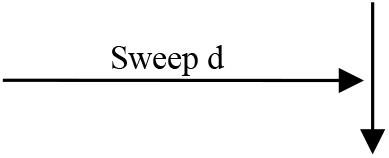
(3) Sweep the 3D orientation n of a plane (see Figure A9(B)(ii)) 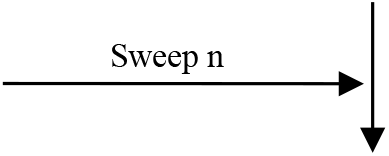
(4) Convert the normalized shortest-distance d to the normalized time-to-contact t by the following equation (see equation(3.0-3))

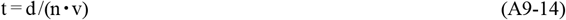 Since the shortest distance d has been converted to the time-to-contact t, the rest of the explanation is the same as Section **A9.1**.
(5) Sweep a RF center O_IJ_ (see Figure A9(A)) 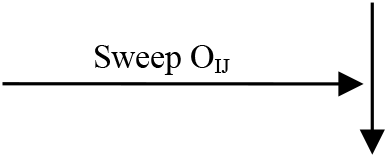 Calculate P_0_ using the following equation (see equation(A4-3) in Appendix **4**).

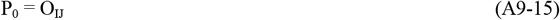
(6) Calculate P_t_ (see Figures A9(A) and (D), and Appendix **5**)

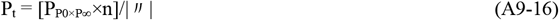 Here, P_P0×P∞_ used above is calculated by the following equation (see equation(A4-6))

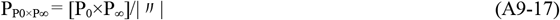
(7) Calculate the polar coordinate τ_2D_ of a MDC by the cross-ratio transform Using the cross-ratio transform of equation(1.0-4), the coordinate τ_2D_ is calculated by the following equation (see equation(A4-14) in Appendix **4**).

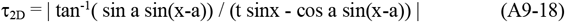 Here, the central angles a and x used above are calculated by the following equations (see equations(A4-16 and 17)).

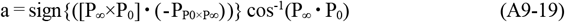

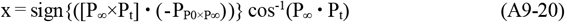
(8) Calculate the polar coordinate ϕ of the MDC (see Figures A9(C) and (D), and equation(A4-13))

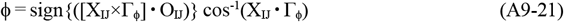 Here, X_IJ_ and Γ_ϕ_ used above are calculated by the following equations (see equations(A4-4 and 7)).

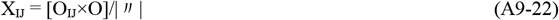

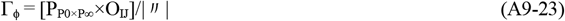
(9) Convert the polar coordinate (τ_2D_,ϕ) to the orthogonal coordinate (τ_X_,τ_Y_) by the following equations (see Figure A9(C))

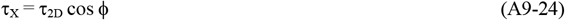

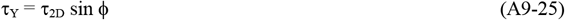
(10) Connect the SDC (n,d) to the MDC (τ_X_,τ_Y_) (see Figure 24(B)) 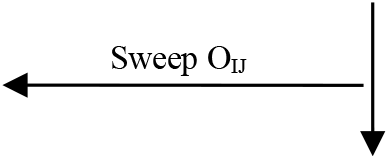
(11) The neural network for all RFs including red, green, and blue in Figure 24(B)(i) has been completed 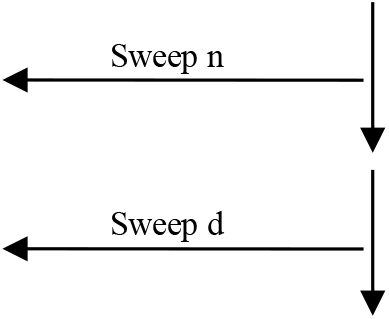
(12) The entire neural network backwardly connecting the SDC column to MDCs (Figure 24(B)(iii)) has been accurately expressed by mathematical equations. That is, each of the neural network that detects the normalized normalized shortest-distance and 3D orientations by the backward method has been expressed one by one by a mathematical equation. 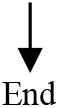

In this way, the neural network of Figure 17(C) (or Figure 24(B)(iii)) has been accurately expressed by mathematical equations at the level of the neural connection of which cell is connected to which cell. This neural connection is shown in detail as the backward connection table in Section **S2.4.1** of Supplementary materials.

The above is represented by the complex equations such as equation(A9-18), and you may think that it is difficult for cerebral cells to execute them. However, that worry is not a problem for the same reasons as in Section **2.2**(12).

## Appendix 10

Representation of the backward method for binocular stereo by mathematical equations

The mathematical representation for binocular stereo by the forward method was explained in Section **4.4.4**. Here, using Figure A10, let us explain the mathematical representation for binocular stereo by the backward method.

Based on the correspondence in Section **4.4.1**, replacing the terms (such as cell names and parameters) in the mathematical expression for motion stereo (which is described in Section **A9.2** of Appendix **9**) with those for binocular stereo enables binocular stereo by the backward method to be expressed in mathematical equations: Figure A10 was obtained by replacing the parameters (P_0_, P_1_, v, P_p0×p∞_, d, τ_X_, τ_Y_, τ_2D_, and SDC) in Figure A9 with the parameters (P_L_, P_R_, X, P_pL×p∞_, d_Bino_, σ_X_, σ_Y_, σ_2D_, and SDC_Bino_), respectively. The backward method for binocular stereo is described as follows.

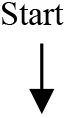

(1) Give the direction vector X of the visual axis (see Figure 27(A)) **Figure A10.**
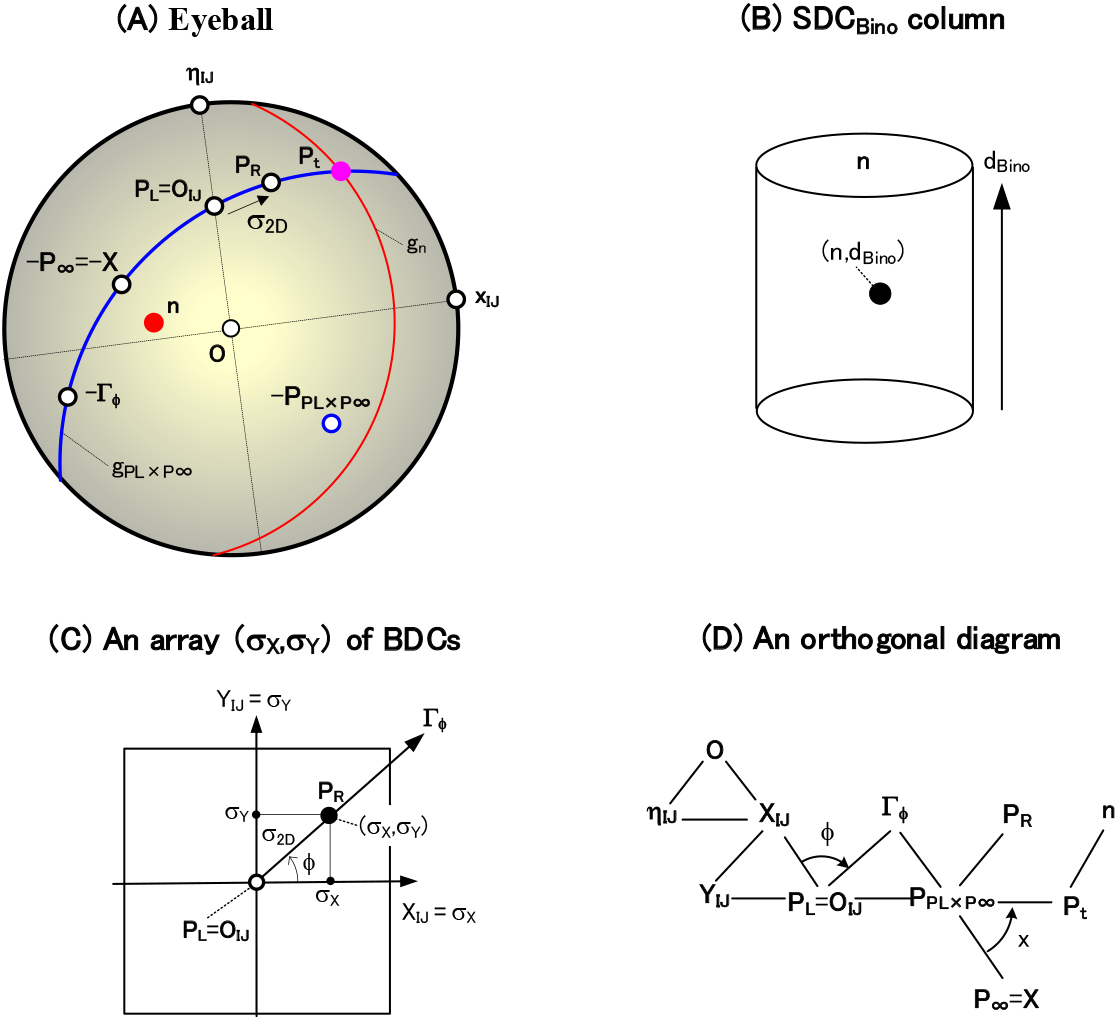
Detection of a plane by binocular stereo using the backward method. Based on the correspondence in Section **4.4.1**, this figure has been obtained by replacing the parameters (P_0_, P_1_, P_p0×p∞_, τ_2D_, τ_X_, τ_Y_, d, and v) for motion stereo of Figure A9 in Appendix **9** with parameters (P_L_, P_R_, P_pL×P∞_, σ_2D_, σ_X_, σ_Y_, d_Bino_, and X) for binocular stereo, respectively. This replacement yields (A), (B), (C), and (D): the (i) in Figure A9(B) is unnecessary and thus was deleted. Calculate P_∞_ using the following equation (see equation(4.4-3)).

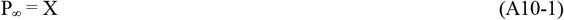 Here, X = (1,0,0).
(2) Sweep a normalized shortest-distance d_Bino_ (see Figure A10(B)) 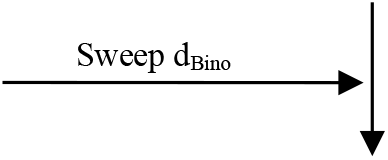
(3) Sweep the 3D orientation n of a plane (see Figure A10(B)) 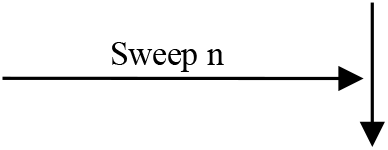
(4) Convert the normalized shortest-distance d_Bino_ to the normalized time-to-contact t by the following equation (see equation(3.0-3))

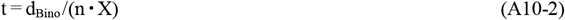 Since the shortest distance d_Bino_ has been converted to the time-to-contact t, the rest of the explanation is the same as Section **A9.1** of Appendix **9**. Note that this conversion to the time-to-contact t is for convenience, and this time t does not have the meaning of time. Also, P_t_ in equation(A10-4) later is for convenience, and does not have the meaning of time.
(5) Sweep a RF center O_IJ_ 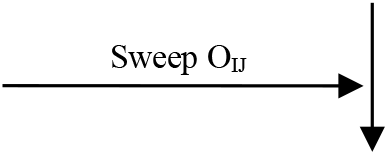 P_L_ is calculated using the following equation by replacing P_0_ in equation(A9-2) with P_L_.

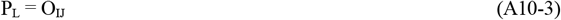
(6) Calculate P_t_ (see equations(A9-3 and 4))

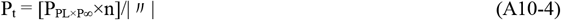 Here, P_PL×P∞_ used above is calculated by the following equation.

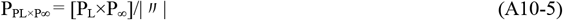
(7) Calculate the polar coordinate σ_2D_ of a BDC by the cross-ratio transform (see equations(A9-5~7)) Using the cross-ratio transform of equation(1.0-4), the coordinate σ_2D_ is calculated by the following equation.

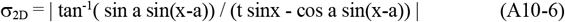 Here, the central angles a and x used above are calculated by the following equations.

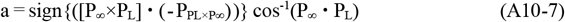

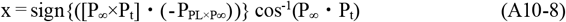
(8) Calculate the polar coordinate ϕ of the BDC (see Figures A10(C) and (D), and equation(A9-8))

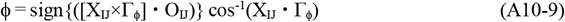 Here, X_IJ_ and Γ_ϕ_ used above are calculated by the following equations (see equations(A9-9 and 10)).

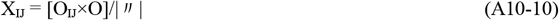

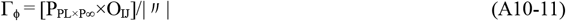
(9) Convert the polar coordinate (σ_2D_,ϕ) to the orthogonal coordinate (σ_X_,σ_Y_) by the following equations (see Figure A10(C))

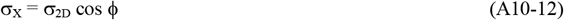

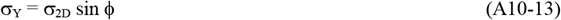
(10) Connect the SDC_Bino_ (n,d_Bino_) to the BDC (σ_X_,σ_Y_) (see Figure 24(B)) Note that the height axis t of the SDC column in Figure 24(B)(iii) is replaced with d_Bino_, and the MDCs in Figure 24(B)(ii)(f) is replaced with the BDCs. The same applies to subsections (11) and (12) below. 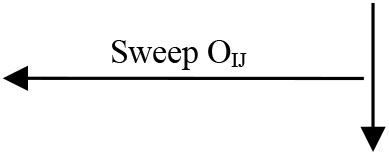
(11) The neural network for all RFs including red, green, and blue in Figure 24(B)(i) has been completed 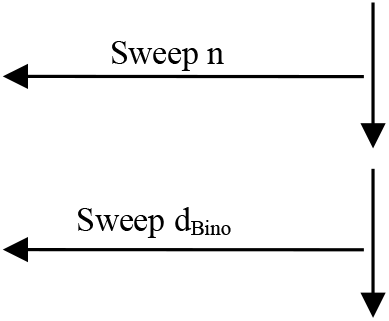
(12) The entire neural network backwardly connecting the SDC_Bino_ column to BDCs (Figure 24(B)(iii)) has been accurately expressed by mathematical equations. That is, each of the neural network that detects the normalized normalized shortest-distance d_Bino_ and 3D orientations by the backward method has been expressed one by one by a mathematical equation. 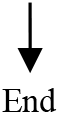

In this way, the neural network of Figure 30(C) (or Figure 24(B)(iii)) has been accurately expressed by mathematical equations at the level of the neural connection of which cell is connected to which cell. The above is represented by the complex equations such as equation(A10-6), and you may think that it is difficult for cerebral cells to execute them. However, that worry is not a problem for the same reasons as in Section **2.2**(12).

## Supplementary materials

### How to create a series of connection tables that represents neural networks for the detection of a planar surface by motion stereo

In order to validate the series of cells that detect planes by motion stereo in Figure 9 or 17, it is necessary to create a simulator on a computer. The main points of the procedures for creating the simulator are shown as follows. First, Section **S1** will show a procedure for determining the array sizes of each cell, receptive field (RF), etc. Next, Section **S2** will show a procedure for creating a connection table that represents each neural network: this table accurately represents the neural network of Figure 9 or 17 at the level of the neural connection of which cell is connected to which cell. Finally, Section **S3** will show a procedure for calculating the response of the series of cells in Figure 9 or 17 using these connection tables. Note that the simulator in Section **4.4.5** for validating the cells that detect planes by binocular stereo in Figure 30 was created on a computer in the similar procedures.

These procedures were created in the following history. First, in 1991, with the support of Okamoto, I created a procedure for calculating local motions of Figure 9(B) or 17(B). Next, in 1998, with the support of Matsuoka, I created a procedure for calculating the detection of a plane based on these local motions (Figures 9(C) or 17(C)), followed by partial improvements. Based on these procedures, we created simulators (Kawakami & Okamoto, 1996a, 1995a, 1992d; Kawakami, 1996b; Akima et al., 2017; Kawakami et al., 1992b; Kawakami et al., 2003, 2000, 1992b). Some of the simulated results were shown in Figures 13, 19, 22(B), 29, and 32.

Note the following. The recognition performed in our brain consists of the space recognition and and form recognition. First, this paper has modeled cells that perform the space recognition (i.e. the motion and binocular stereos). Next, we reported a cell model related to the form recognition (Kawakami et al., 2020): this cell detects curvatures of figures that play an important role in the form recognition; the simulator for validating the cells that detect these curvatures was also created on a computer in the similar procedures.

### S1. Array sizes of the cells and RFs

#### S1.1. Composition of subsequent sections

In Section **S1.2**, array’s sizes of the retinal cells and RFs will be determined, and then an array’s size of each cell type in Figure 9(B) or 17(B) will be determined. Section **S1.3** will determine the array’s size of a SDC column in Figures 9(C) or 17(C) that detects of planes.

#### S1.2. Each array size from retinal cells to MDCs

In the last section (i.e. in Section **S1.2.6**), a size of each array that constitutes the series of cells from retinal cells to MDCs in Figure 9(B) or 17(B) will be determined. Various preparations for this purpose will be made in Sections **S1.2.1** to **S1.2.5**.

##### S1.2.1. Array P_ij_ of retinal (or LGN) cells on the eyeball and its address (i,j)

(1) Array P_ij_ of retinal (or LGN) cells (see Figure S1-1(A)) **Figure S1-1.**
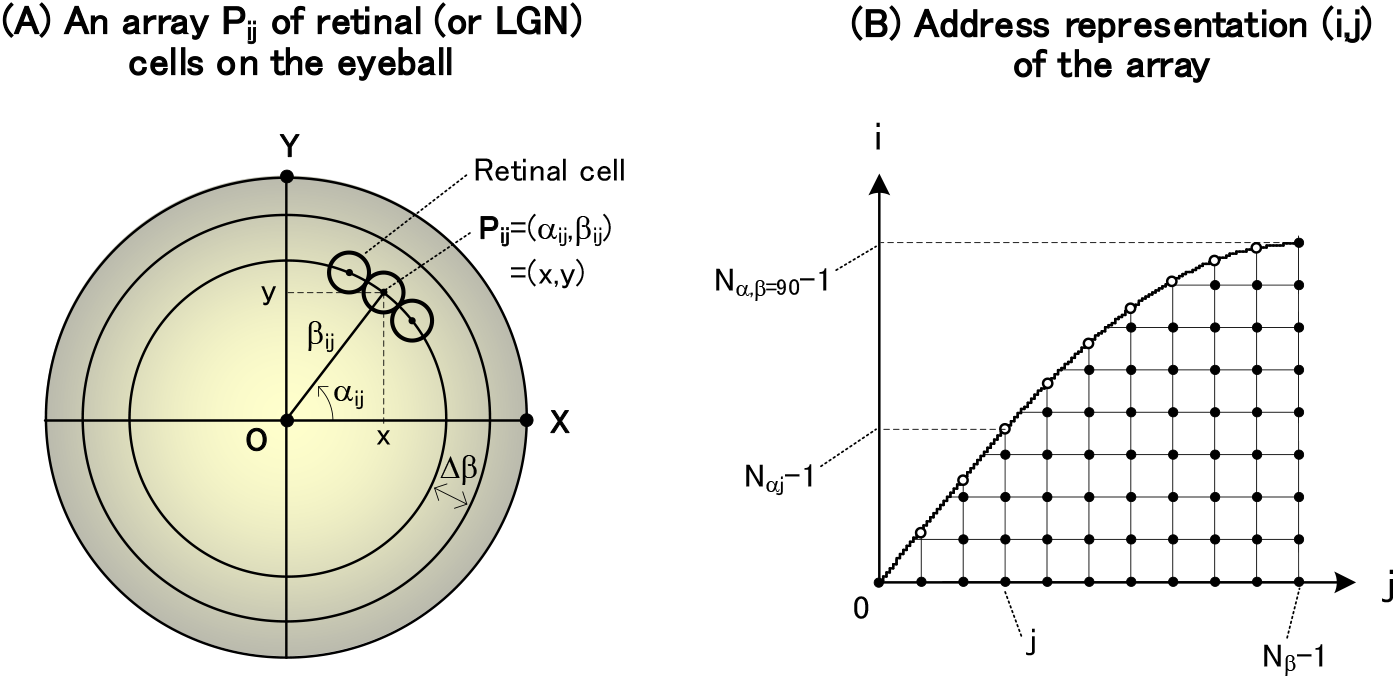
Array P_ij_ of retinal (or LGN) cells on the eyeball and its address (i,j). (A) Each retinal (or LGN) cell P_ij_ is arrayed at an equal interval Δβ that is calculatd by equation(S1.2.1-6). (B) This array is represented by its address (i,j). Each retinal (or LGN) cell P_ij_ is arranged, as follows, on the eyeball at an equal interval Δβ that will be calculated by equation(S1.2.1-6). In this way, since the retinal and LGN cells are arranged at the same interval Δβ, all the parameters for the retinal and LGN cells are represented by the same equations(S1.2.1-1~10c) below.

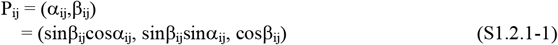 Here, (α_ij_,β_ij_) is the polar coordinate of P_ij_, and parameters used above are shown as follows.

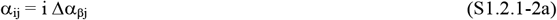

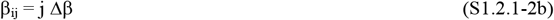

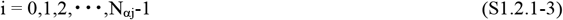

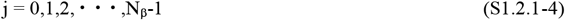 **Figure S1-2.**
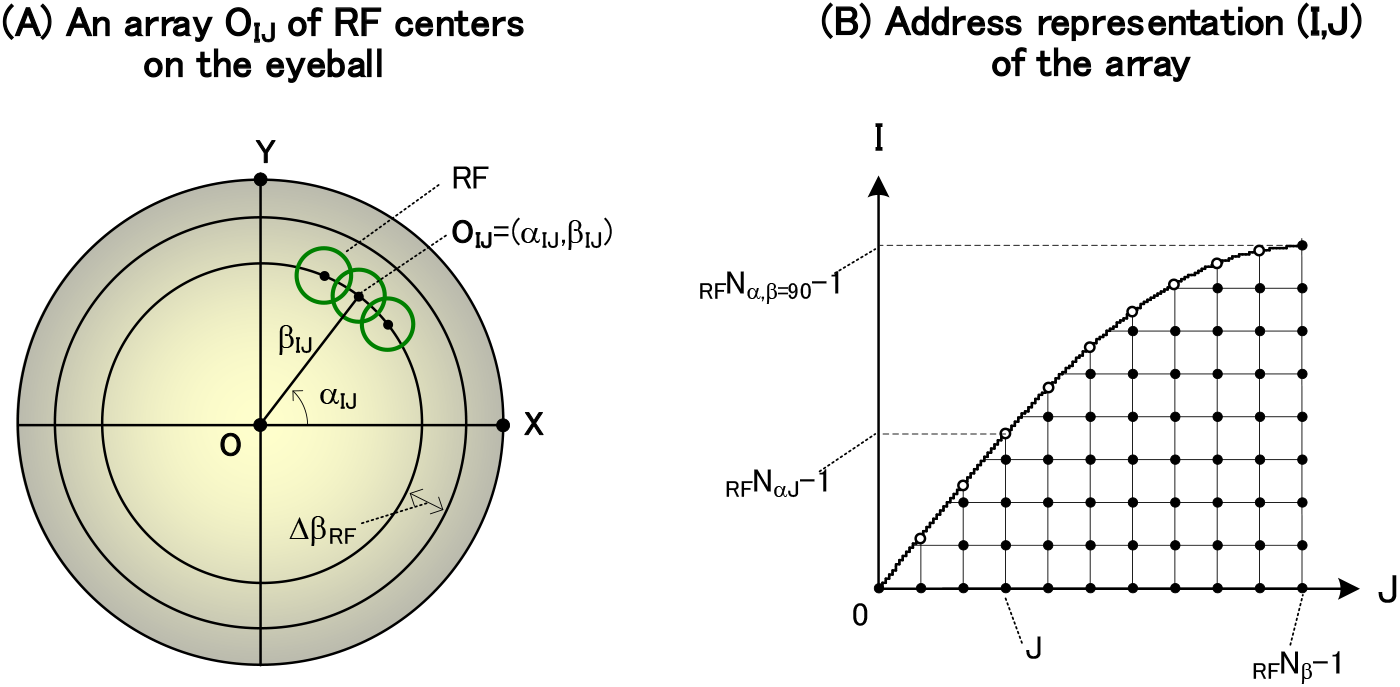
Array O_IJ_ of RF centers on the eyeball and its address (I,J). (A) Each RF center O_IJ_ is arrayed at an equal interval Δβ_RF_ that is calculatd by equation(S1.2.2-6). (B) This array is represented by its address (I,J). The main parameters (N_β_, Δβ, N_αj_, and Δα_βj_) used above are calculated in subsection (2) below. Figure S1-1(B) is obtained by expressing each cell P_ij_ with its address (i,j).
(2) Calculate the parameters (N_β_, Δβ, N_αj_, and Δα_βj_) as follows (see Figure S1-1)
  a. Temporarily set the latitudinal pitch Δβ’ of retinal (or LGN) cells
  b. Calculate the number N_β_ of retinal (or LGN) cells in the latitudinal direction based on this Δβ’

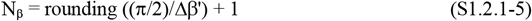
  c. Calculate finally the latitudinal pitch Δβ of retinal (or LGN) cells

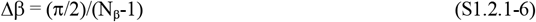
  d. Calculate the maximum number N_α,β=90_ of cells in the longitude direction

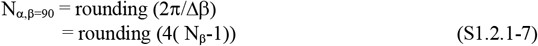 **Figure S1-8.**
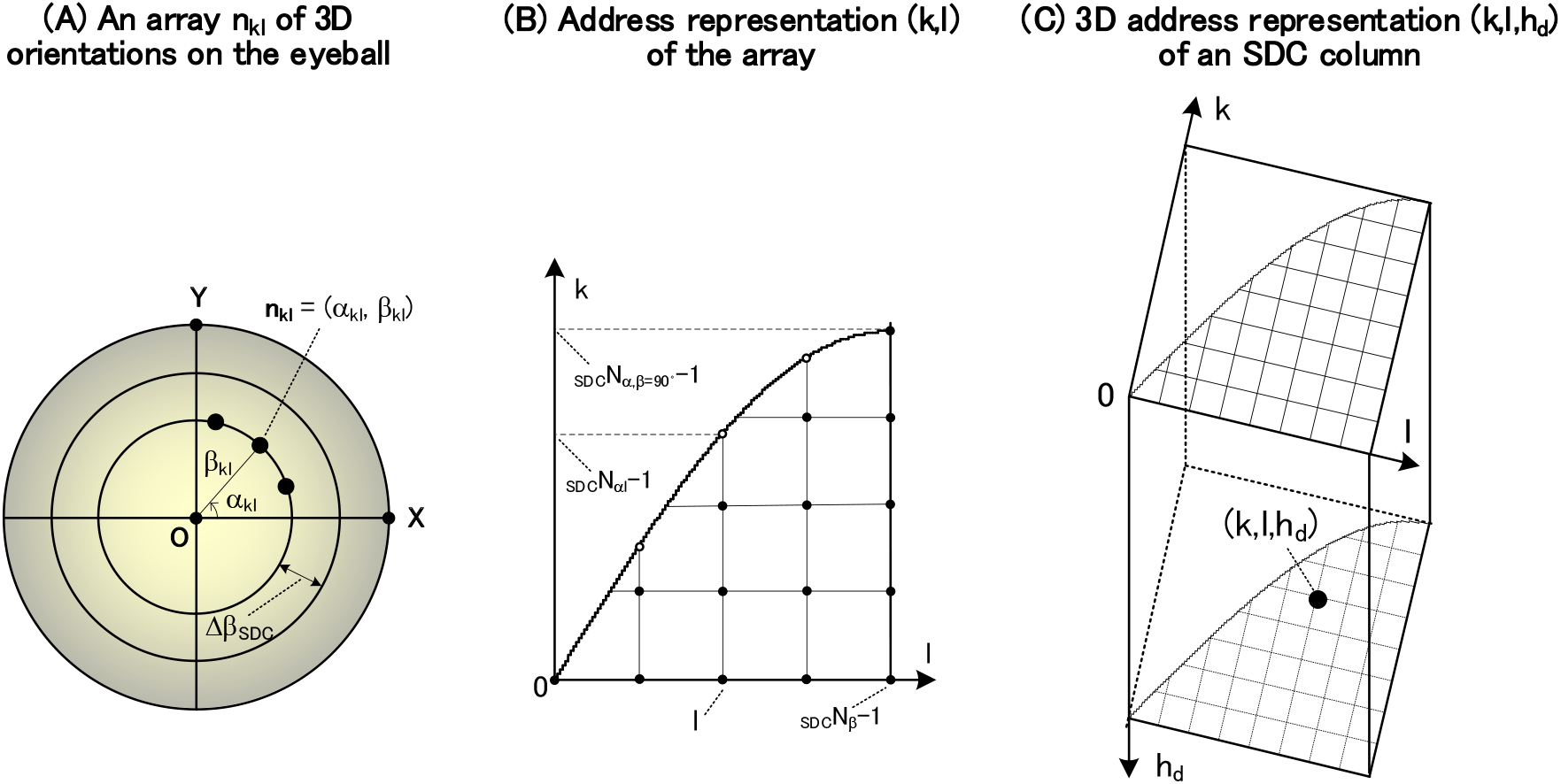
3D address representation (k,l,h_d_) of a SDC column for detecting normalized shortest-distances and 3D orientations. (A) This array is the same as in Figure S1-7(A). (B) This address representation (k,l) is the same as in Figure S1-7(B). (C) A 3D address representation (k,l,h_d_) of the SDC column in Figure 17(C) is shown. That is, this representation has been obtained by stacking the address (k,l) in the (B) along the height address h_d_.
  e. Calculate, at the latitude address j, the number N_αj_ of cells and the pitch Δα_βj_ in the longitude direction (see Figure S1-1(B))

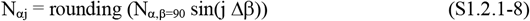

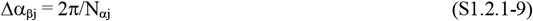 Note: These roundings that allow for equally spaced coordinates are necessary to determine the coordinates of the retinal (or LGN) cells by a modulo operation.
(3) Calculate the total number of retinal (or LGN) cells N_total_

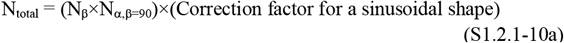

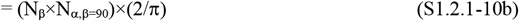 Here, this correction factor is obtained as the ratio of the rectangular area N_β_×N_α,β=90_ to the area of the sine wave in Figure S1-1(B), and is given by the following equation.

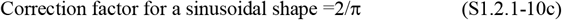

#### S1.2.2. Array O_IJ_ of RF centers on the eyeball and its address (I,J)

(1) Array O_IJ_ of RF centers (see Figure S1-2(A)) Each RF center O_IJ_ is arranged, as follows, on the eyeball at an equal interval Δβ_RF_ that will be calculatd by equation(S1.2.2-6).

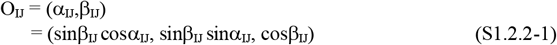 Here, (α_IJ_,β_IJ_) is the polar coordinate of O_IJ_, and parameters used above are shown as follows.

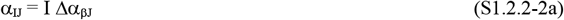

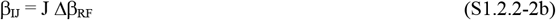

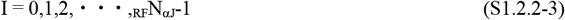

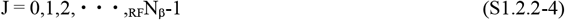 The main parameters (_RF_N_β_, Δβ_RF_, _RF_N_αJ_, Δα_βJ_) used above are calculated in subsection (2) below. Figure S1-2(B) is obtained by expressing each RF center O_IJ_ with its address (I,J).
(2) Calculate the parameters (_RF_N_β_, Δβ_RF_, _RF_N_αJ_, Δα_βJ_) as follows (see Figure S1-2)
  a. Temporarily set the latitudinal pitch Δβ_RF_’ of RF centers
  b. Calculate the number _RF_N_β_ of RF centers in the latitudinal direction based on this Δβ_RF_’

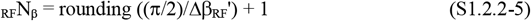
  c. Calculate finally the latitudinal pitch Δβ_RF_ of RF centers

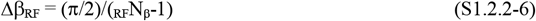
  d. Calculate the maximum number _RF_N_α,β=90_ of RF centers in the longitude direction

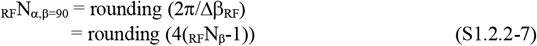
  e. Calculate, at the latitude address J, the number _RF_N_αJ_ of RF centers and the pitch Δα_βJ_ in the longitude direction (see Figure S1-2(B))

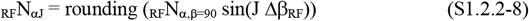

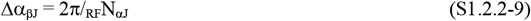 Note: These roundings that allow for equally spaced coordinates are necessary to determine the coordinates of RF centers by a modulo operation.
(3) Calculate the total number of RF centers _RF_N_total_

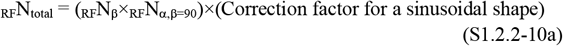

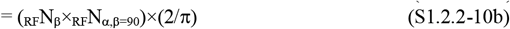 Here, this correction factor is the same as in equation (S1.2.1-10c).

#### S1.2.3. Local coordinate system (O_IJ_,X_IJ_,Y_IJ_) of each RF

Its definition is shown in Figure S1-3, where O = (0,0,1).

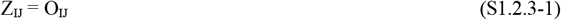

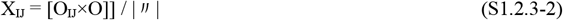

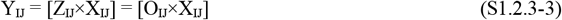

Here, when O_IJ_=O, the following equations are obtained.

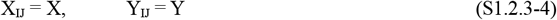

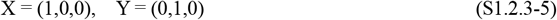

#### S1.2.4. Cutting out of retinal (or LGN) cells belonging to each RF and creating of a cutout table _I,J_T_RF_[m→(i_m_,j_m_)]

Before starting the cutting-out process, you should be aware that a cutout table created below is the same for retinal cells and LGN cells. This is because, as shown in Figure S1-1(A), retinal cells and LGN cells are arranged in the same array on the eyeball.

All retinal (or LGN) cells belonging to each RF are cut out as follows. First, scan a serial number m for each RF, and cut out a retinal (or LGN) cell corresponding to this number. Next, create a cutout table _I,J_T_RF_[m→(i_m_,j_m_)] by confirming whether this cell is in the RF. This table will be used in Sections **S2.1.1** and **S3.1.2.1**. The procedure for creating this cutout table will be described below: there are two ways to do this procedure, as described in Sections **S1.2.4.1** and **S1.2.4.2**.

Before that, the notation of this table is explained. Subscripts _RF_ and _I,J_ indicate a RF and its address (I,J), respectively, and m→(i_m_,j_m_) indicates that a retinal (or LGN) cell (i_m_,j_m_) corresponding to each serial number m is cut off.

#### S1.2.4.1. Method using equation(S1.2.4-10)

Cutting out of retinal (or LGN) cells and creating of a cutout table _I,J_T_RF_[m→(i_m_,j_m_)] are performed by the following procedure (see Figure S1-4).

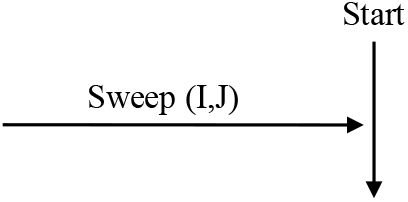

The sweep (I,J) of a RF is performed by equations(S2.1.1-1~3d) in Section **S2.1.1**.

1. Calculate a RF center O_IJ_ (see equations(S1.2.2-1~4) in Section **S1.2.2**)

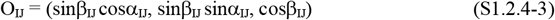

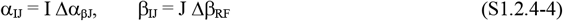

where Δβ_RF_ and Δα_βJ_ will be calculated by the equations(S2.1.1-3b and 3h) in Section **S2.1.1**.
2. Reset a serial number m

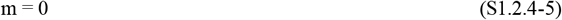 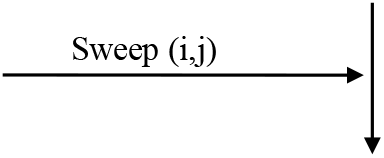 The sweep (i,j) of a retinal (or LGN) cell is performed by equations(S1.2.1-3~8) in Section **S1.2.1**.
3. Calculate a retinal (or LGN) cell P_ij_ (see equations(S1.2.1-1~2) in Section **S1.2.1**(1))

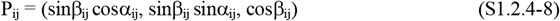

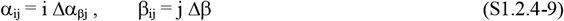

where Δβ and Δα_βj_ were calculated using equations(S1.2.1-6 and 9) in Section **S1.2.1**.
4. For an address (i,j) that satisfies equation(S1.2.4-10), perform the following ➀ and ➁ (see Figure S1-4).

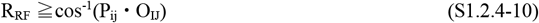 **Figure S1-3.**
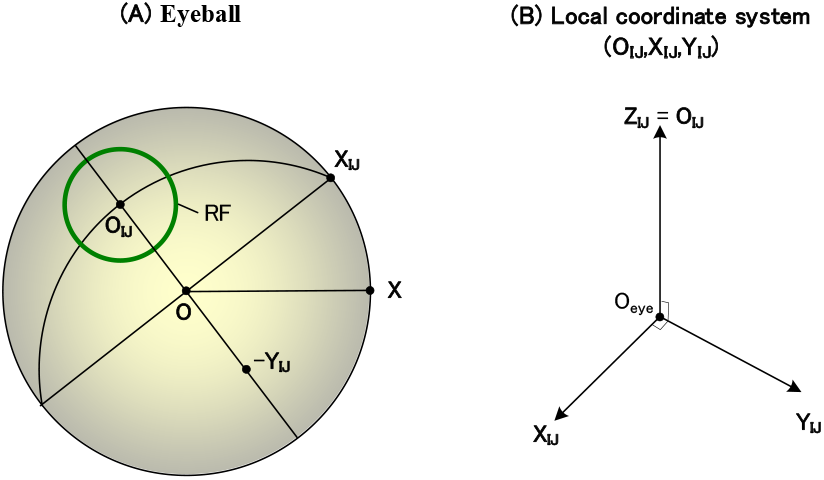
Local coordinate system (O_IJ_,X_IJ_,Y_IJ_) of each RF. (A) A local coordinate system (O_IJ_,X_IJ_,Y_IJ_) on the eyeball is shown. (B) The system (O_IJ_,X_IJ_,Y_IJ_) is displayed in an orthogonal coordinate system.

**Figure S1-4.**
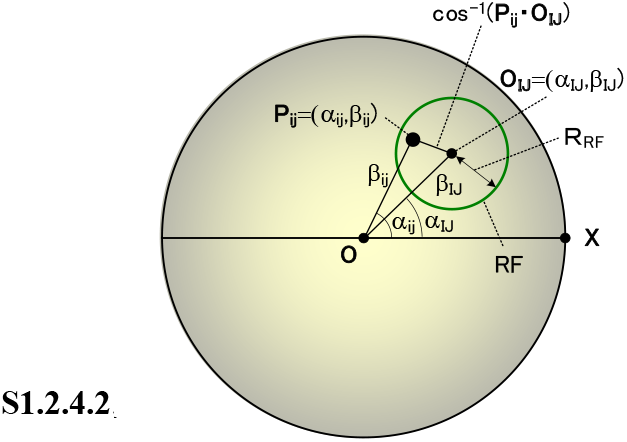
Cutting out of retinal (or LGN) cells within each RF. Parameters necessary for this cutting out are shown. A RF center O_IJ_ and a retinal (or LGN) cell P_ij_ are represented in polar coordinates as (α_IJ_,β_IJ_) and (α_ij_,β_ij_), respectively. R_RF_ is the RF radius.
  ➀ Write (i_m_,j_m_) to the content of an address m in the table

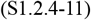 Note: The address (i,j) of the retinal (or LGN) cell above has been changed to (i_m_,j_m_) for the convenience of notation
  ➁ Add 1 to the serial number m

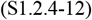

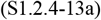 Here, a RF’s radius R_RF_ is calculated by the following equation (see equation(S1.2.6.1-35b)).

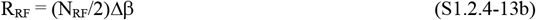

where N_RF_ will be shown in Figure S1-5(A) and will be calculated by equation (S1.2.6.1-22). 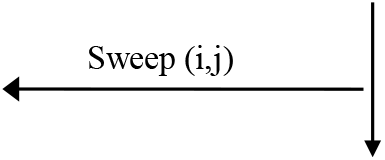
5. For each RF O_IJ_, write the maximum value m_max_ of the serial number m to m_max_(I,J)

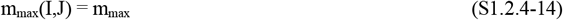 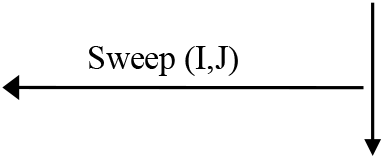
6. The cutout table _I,J_T_RF_[m→(i_m_,j_m_)] has been created. 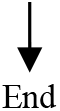

#### S1.2.4.2. Method using equation (S1.2.4-20)

Cutting out of retinal (or LGN) cells and creating of a cutout table _I,J_T_RF_[m→(i_m_,j_m_)] are performed by the following procedure (see Figure S1-4). This method using equation(S1.2.4-20) does not require the calculation of the O_IJ_ and P_ij_, and thus is simpler than Section **S1.2.4.1**.

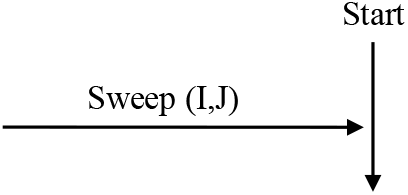

The sweep (I,J) of a RF is performed by equations(S2.1.1-1~3d) in Section **S2.1.1**.

1. Reset a serial number m

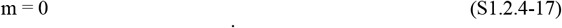

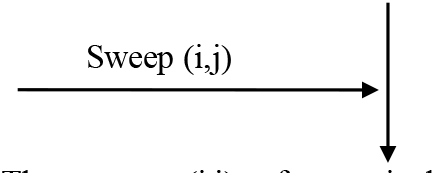 The sweep (i,j) of a retinal (or LGN) cell is performed by equations(S1.2.1-3~8) in Section **S1.2.1**.
2. For an address (i,j) that satisfies equation(S1.2.4-20), perform the following ➀ and ➁ (see Figure S1-4).

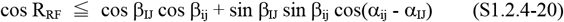 This equation has been obtained by applying the cosine rule (Bronshtein & Semendyayev, 1978; Izumi et al., 1987) to the spherical triangle P_ij_O_IJ_O in Figure S1-4. Here, the parameters used above are described as follows.

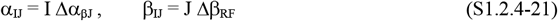

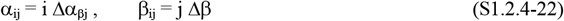

where Δβ_RF_ and Δα_βJ_ will be calculated by equations(S2.1.1-3b and 3h), and Δβ and Δα_βj_ will be calculated by the equations(S1.2.6.1-4 and 8). **Figure S1-5.**
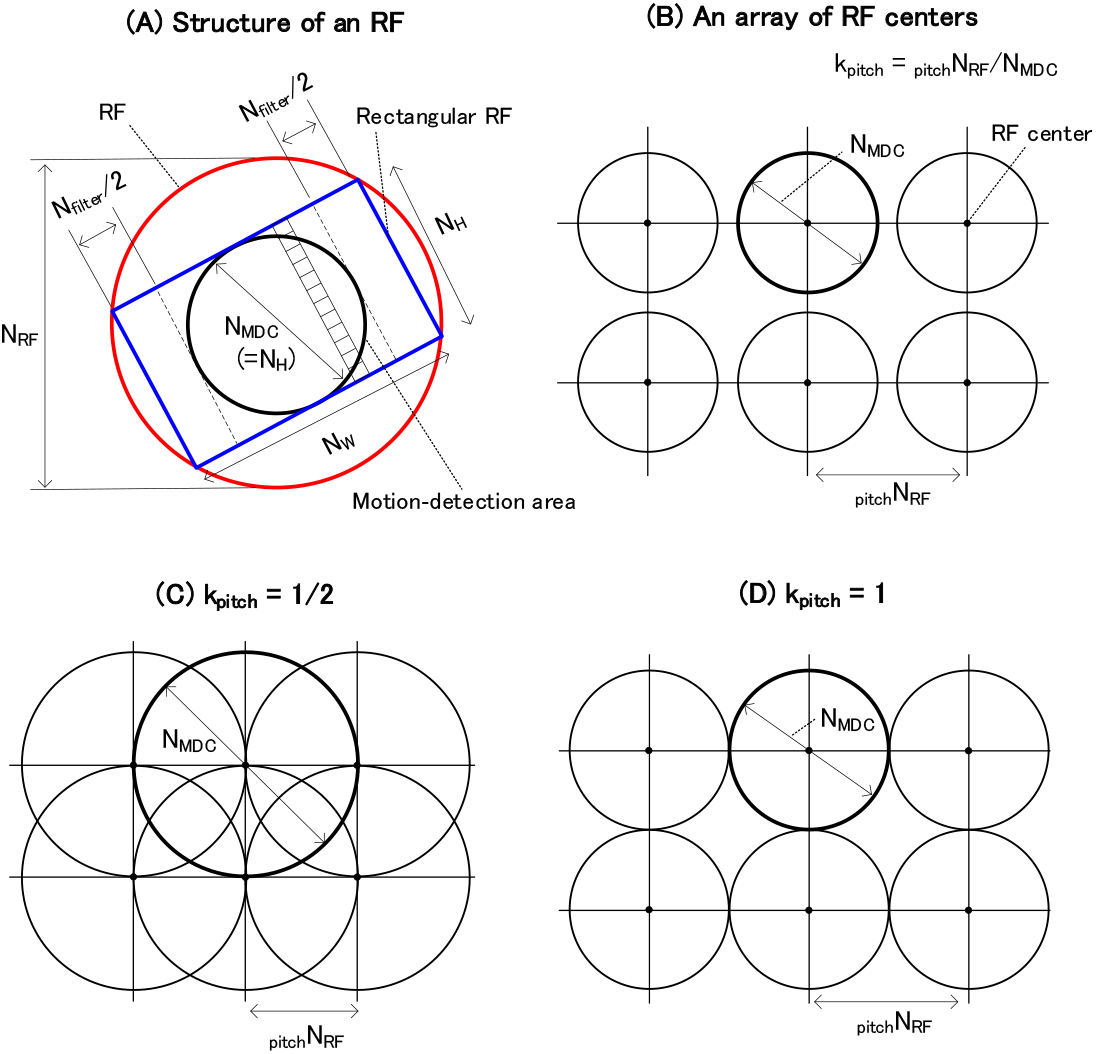
A structure of each RF and a pitch between RF centers. (A) The structure of a RF is shown. A red circle indicates a RF with a diameter N_RF_: the diameter is not a size but an integer in the unit of a retinal(or LGN)-cell pitch Δβ that is caluculated by equation(S1.2.1-6). A blue rectangular RF inscribed in the RF has a height of N_H_ and a width of N_W_. In the rectangular RF, every small rectangle aligned in the height direction represents a retinal (or LGN) cell. The N_filter_/2 at both ends in the width direction corresponds to the width N_filter_/2 of the DOG filter in equations(S3.1.1-6a and 6b) of Supplementary materials. A circle with a diameter N_MDC_ inscribed in this rectangular RF represents a detection region of local motion, and each MDC inside this region is guaranteed to detect a local motion. (B)An array of the RF centers is shown. The pitch of each array is represented as _pitch_N_RF_, and each circle indicates a motion-detection area with a diameter N_MDC_. A parameter k_pitch_ is _pitch_N_RF_ /N_MDC_. (C) An array with k_pitch_=1/2 is shown. This array is default, and there is no omission in each local-motion detection. (D) An array with k_pitch_=1 is shown. In this array, each local-motion detection region is in contact.
  ➀ Write (i_m_,j_m_) to the content of an address m in the table

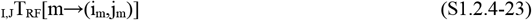 Note: The address (i,j) of the retinal (or LGN) cell above has been changed to (i_m_,j_m_) for the convenience of notation

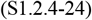

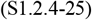

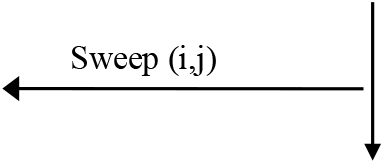
  ➁ Add 1 to the serial number m
3. For each RF O_IJ_, write the maximum value m_max_ of the serial number m to m_max_(I,J)

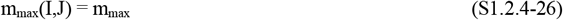 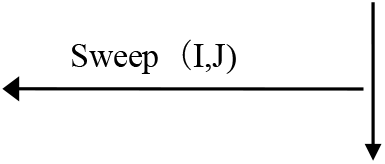
4. The cutout table _I,J_T_RF_[m→(i_m_,j_m_)] has been created.

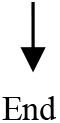

#### S1.2.5. RF structure and array sizes of various cell types

The structure of a RF is shown in Figure S1-5(A), and the array sizes of various cell types are shown in Figure S1-6 (see Figure A2 of Kawakami & Okamoto (1996a) and Figure A-2 of Kawakami (1996b)).

1. First, set the following basic parameters (see Figure S1-5(A))
  - N_MDC_: diameter of a motion-detection area
  - N_filter_: width for two dimensional (2D) filtering (see equations(S3.1.1-6a and 6b) for the 2D filtering)
2. Based on these parameters, calculate various parameters related to the RF, as follows (see Figure S1-5(A))
  ➀ Calculate the width N_W_ and height N_H_ of a rectangular RF

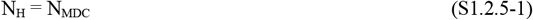

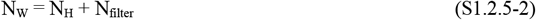
  ➁ Calculate the RF diameter N_RF_

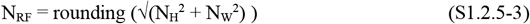 The RF’s radius R_RF_ is calculated by the following equation.

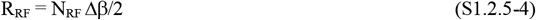 Here, Δβ was calculated by equation(S1.2.1-6).
3. Calculate parameters related to the RF pitch (see Figure S1-5(B))
  ➀ Calculate the RF’s pitch _pitch_N_RF pitch_N_RF_

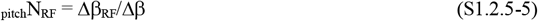 Here, Δβ_RF_ was calculated by equation(S1.2.2-6).
  ➁ Calculate the ratio k_pitch_ of RF’s pitch _pitch_N_RF_ to N_MDC_

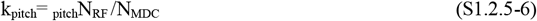

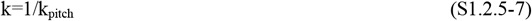
  Calculate parameters related to the arrays of various cell types (see Figure S1-6)

An array size of each cell type corresponding to Figure 9(B) (i.e. of NDS simple cells, DS simple cells, DS complex cells, and MDCs) is shown in Figure S1-6.

➀ Array of NDS simple cells (see Figure S1-6(A))
  a. Calculate an input array (_In_N_ρ_,N_θ_) for a 2D filtering

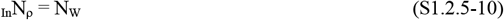

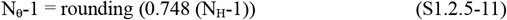 Note: this coefficient 0.748 refers to page 184 of Kawakami (1996b).
  b. Calculate an output array (_Out_N_ρ_,N_θ_) from this input array to a DS simple cell

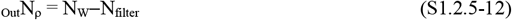 Note: N_θ_ is the same as equation(S1.2.5-11).
➁ Calculate an array (N_ρ_,N_θ_,N_τ_) of the DS simple cell (see Figure S1-6(B))

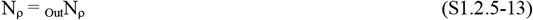

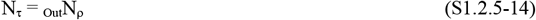

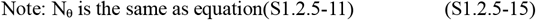

**Figure S1-6.**
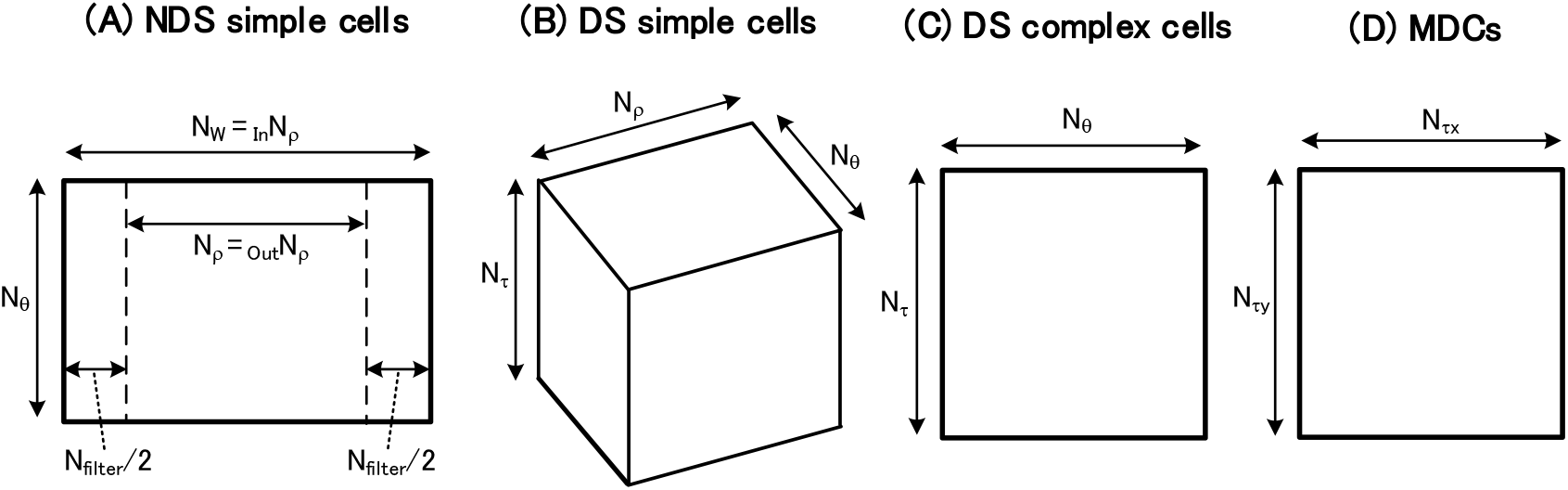
Array sizes of various cell types. Array sizes for the series of cell types in Figure 9(B) (or Figure 17(B)) are shown. (A) An array of NDS simple cells is shown, constituting a 2D array with N_θ_ height and _In_N_ρ_ width. Each region N_filter_/2 corresponds to the width N_filter_/2 of the DOG filter in equations(S3.1.1-6a and 6b) of Supplementary materials. The region _Out_N_ρ_ is delivered to DS simple cells. (B) An array of DS simple cells is shown, constituting a 3D array with height N_τ_, width N_ρ_, and depth N_θ_. (C) An array of DS complex cells is shown, constituting a 2D array with N_τ_ height and N_θ_ width. (D) An array of MDCs is shown, constituting a 2D array with N_τy_ height and N_τx_ width.
➂ Calculate an array (N_θ_,N_τ_) of the DS complex cell (see Figure S1-6(C)) N_θ_ and N_τ_ are the same as those of the DS simple cell.
➃ Calculate an array (N_τx_,N_τy_) of the MDC (see Figure S1-6(D)) N_τx_ = N_τy_ = _Out_N_ρ_ (S1.2.5-16)

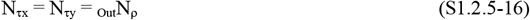

#### S1.2.6. Various parameters used for the simulation

Based on the above preparation, let us calculate various parameters required for the simulation, as follows.

##### S1.2.6.1. Calculation of various parameters used for the simulation

As described in Section **S1.2.1**(1), you should be aware that the parameters below (N_β_ and Δβ) for retinal cells and LGN cells are the same.

1. Basic parameters of the retinal (or LGN) cell array (see Figure S1-1 and Section **S1.2.1**) ➀ Temporarily set the number N_β_’ of retinal (or LGN) cells in the latitudinal direction and calculate Δβ’

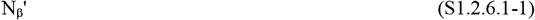

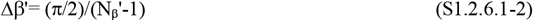 ➁ Calculate the following parameters based on Δβ’
  - N_β_: number of retinal (or LGN) cells in the latitudinal direction

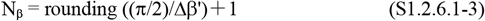
  - Δβ: pitch of retinal (or LGN) cells

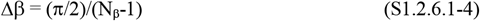
  - N_α,β=90_: maximum number of cells in the longitudinal direction at a latitude of 90 degrees

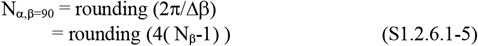
  - N_αj_: number of cells in the longitudinal direction at a latitude address j

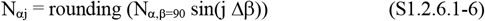

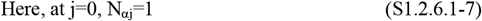
  - Δα_βj_: cell pitch in the longitudinal direction at a latitude address j

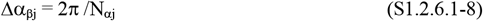
2. Basic parameters of the RF structure and the RF pitch (See Figure S1-5 and Section **S1.2.5**)
  ➀ Set the main parameters of the RF structure (see Figures S1-5(A) and (B))

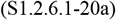

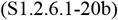

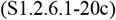
  ➁ Calculate the following parameters based on these settings (see Figure S1-5(A) and equations(S1.2.5-2, 3, and 6))
    - N_MDC_: diameter of a circle inscribed in the rectangular RF

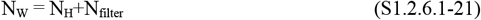
    - N_RF_: RF diameter

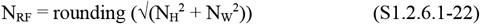
    - N_MDC_: diameter of a circle inscribed in the rectangular RF

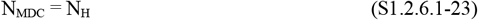
    - _pitch_N_RF_: RF pitch

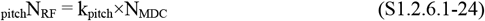
3. Calculate the following parameters of the RF array based on _pitch_N_RF_ and Δβ obtained above (see Figure S1-2 and Section **S1.2.2**).
  ➀ Temporarily calculate the RF pitch Δβ_RF_’ using the following equation

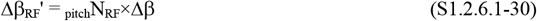
  ➁ Calculate the following parameters based on this Δβ_RF_’ (see Section **S1.2.2**(2))

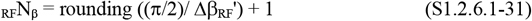

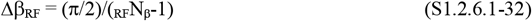

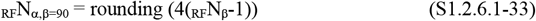

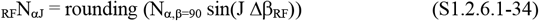

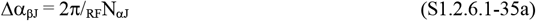

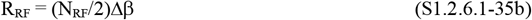 Here, R_RF_ is the radius of a RF.
  ➂ Calculate the number _RF_N_total_ of total RFs (see Section **S1.2.2**(3))

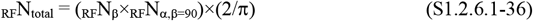 Here, this correction factor 2/π is the same as in equation(S1.2.1-10c).
4. Calculate array sizes of various cell types using N_H_ and N_W_ obtained above (see Figure S1-6 and Section **S1.2.5**(4))
  ➀ Array of NDS simple cells (see Figure S1-6(A))
    a. Calculate an input array (_In_N_ρ_,N_θ_) for 2D filtering

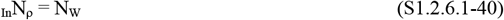

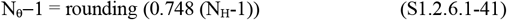

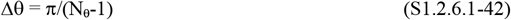 Here, Δθ is the resolution in θ orientation of the NDS simple, DS simple, and DS complex cells.
    b. Calculate an output array (_Out_N_ρ_,N_θ_) from this input array to a DS simple cell

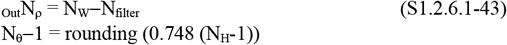
  ➁ Calculate an array (N_ρ_,N_θ_,N_τ_) of the DS simple cell (see Figure S1-6(B))

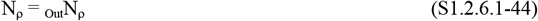

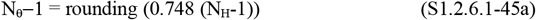

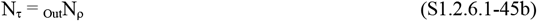
  ➂ Calculate an array (N_θ_,N_τ_) of the DS complex cell (see Figure S1-6(C))

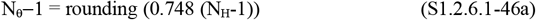

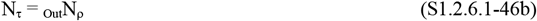
  ➃ Calculate an array (N_τx_,N_τy_) of the MDC (see Figure S1-6(D))

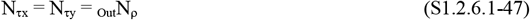
5. Summary of the above
  ➀ The basic parameters to be set are the following four
    - N_β_′: number of retinal (or LGN) cells in the latitudinal direction that is temporarily set (S1.2.6.1-50a)
    - N_H_: height of the rectangular RF (S1.2.6.1-50b)
    - N_filter_: width for a 2D filtering (S1.2.6.1-50c)
    - k_pitch_: coefficient for determining the RF pitch (S1.2.6.1-50d)
  ➁ Other parameters can be calculated based on these four (see Sections **S1.2.6.1**(1~4))
6. Features of these parameters

Before describing the features below, you should note the following. As explained in Section **S1.2.1**(1), the retinal and LGN cells have the same arrangement. Therefore, although it will be described as features of retinal cells hereafter, the following description holds even if it is replaced with LGN cells.

The features of these parameters are shown using Figure S1-5(A), as follows. Not only the main parameter of the RF’s diameter N_RF_ but also the internal structure (i.e. N_H_, N_W_, and N_MDC_) are determined by the number of retinal cells in each RF, not by the length of the RF. This yields a feature that the various RF’s parameters such Δθ, N_θ_, and N_τ_, including the main parameter and internal structure, are determined by the number of retinal cells in each RF and are independent of the RF’ size.

This feature enables the orientation resolution Δθ of the NDS simple ((A)), DS simple ((B)), and DS complex cells ((C)) in Figure S1-6 to remain unchanged when the number of retinal cells in the RF is constant, even if the size of the RF changes (Kawakami et al., 1991): this unchangeness results from equations(S1.2.6.1-41 and 42).

This feature allows the following discovery to be explained. It was discovered that even if the RF’s size changed about 30 times between the fovea and periphery of the visual field, the orientation resolution Δθ of the simple and complex cells remained almost constant at about 10 degrees (Hubel & Wiesel, 1984; Hubel et al, 1978). How this discovery (i.e. the invariance of Δθ) can be explained by this feature is described as follows.

If the number of retinal cells in the RF is unchanged between the fovea and periphery, the above feature allows the orientation resolution Δθ to remain constant even if the RF’s size changes. Physiological data that suggest this unchangeness in the numbers of retinal cells were reported (Wassle et al., 1990; Drasdo, 1977). Therefore, it has been clarified that this feature enables the above discovery of Hubel and Wiesel to be explained: this clarification was reported by Kawakami et al. (1991).

This assumption (suggested by the physiological data above) of that the number of retinal cells in each RF does not change significantly between the fovea and periphery can be extended as follows. That is, let us extend the above unchangeness, on the number of retinal cells contained in each RF, to an unchangeness on the cell number contained in each array of Figure S1-6 (i.e. in each array of NDS simple cells, DS simple cells, DS complex cells, and MDCs). This unchangeness of the cell number contained in each array allows the size of an array to be constant, and thus enables the area of areas V1 and MT in which these arrays exist (see Figure S2-1) to be constant between the fovea and periphery. This constancy of the area of areas V1 and MT is consistent with psychological experiments (e.g. Hubel & Wiesel, 1984; Hubel et al, 1978), and thus has explained these experiments.

Note the following. Despite this unchangeness on the numbers of retinal cells, I think that the pitch Δβ of retinal cells changes by about 30 times in the fovea and periphery.

##### S1.2.6.2. Examples of these parameters

Considering consistency with physiological data (Kawakami & Okamoto, 1996a: Kawakami, 1996b), set the basic parameters of N_β_’, N_H_, N_filter_, and k_pitch_, and then calculate other parameters using Section **S1.2.6.1**, as follows.

1. Parameters of the retinal (or LGN) cell array (see Figure S1-1 and Section **S1.2.6.1**(1))
  ➀ Temporarily set the number N_β_’ of retinal (or LGN) cells in the latitudinal direction and calculate Δβ’

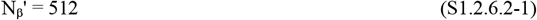

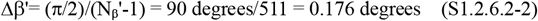
  ➁ Calculate the following parameters based on Δβ’
    - N_β_: number of retinal (or LGN) cells in the latitudinal direction

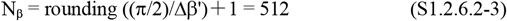
    - N_α,β=90_: maximum number of cells in the longitudinal direction at a latitude of 90 degrees

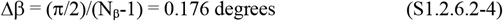
    - N_αj_: number of cells in the longitudinal direction at a latitude address j

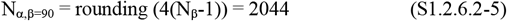 N_αj_ = rounding (N_α,β=90_ sin(j Δβ)) = rounding (2044 sin(j Δβ))

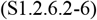

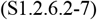 Here, at j=0, N_αj_=1
    - Δα_βj_: cell pitch in the longitudinal direction at a latitude address j

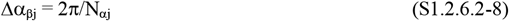
2. Parameters of the RF structure and the RF pitch (See Figure S1-5 and Section **S1.2.6.1**(2))
  ➀ Set the main parameters of the RF structure
    - N_H_: height of the rectangular RF

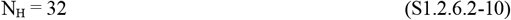
    - N_filter_: width for a 2D filtering

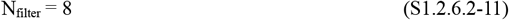
    - k_pitch_: coefficient for determining the RF pitch _pitch_N_RF_

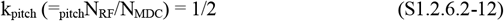 Note: This N_H_=32 has been set based on equations(S1.2.6.2-41~42) so that Δθ is close to 10 degrees (i.e. 7.826 degrees) of physiological data (see page 184 of Kawakami (1996b)).
  ➁ Calculate the following parameters based on these settings
    - N_W_: width of the rectangular RF

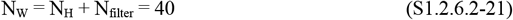
    - N_RF_: RF diameter

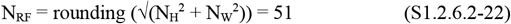
    - N_MDC_: diameter of a circle inscribed in the rectangular RF (see Figure S1-5(A))

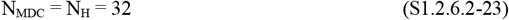
    - _pitch_N_RF_: RF pitch

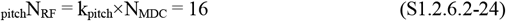
3. Calculate the following parameters of the RF array based on the _pitch_N_RF_ and Δβ obtained above (see Figure S1-2 and Section **S1.2.6.1**(3))
  ➀ Temporarily calculate the RF pitch Δβ_RF_’

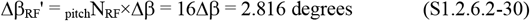
  ➁ Calculate the following parameters based on this Δβ_RF_’ (see Section **S1.2.2**(2))

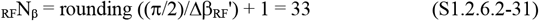

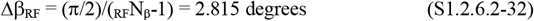

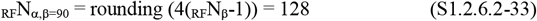

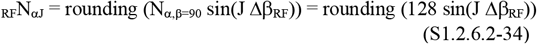

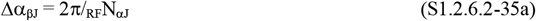

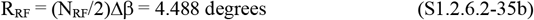
  ➂ Calculate the number _RF_N_total_ of total RFs (see Section **S1.2.2**(3))

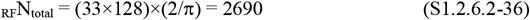 Here, this correction factor 2/π is the same as in equation(S1.2.1-10c).
4. Calculate array sizes of various cell types based on N_H_ and N_W_ obtained above (see Figure S1-6 and Section **S1.2.6.1**(4))
  ➀ Array of NDS simple cells
    a. Calculate an input array (_In_N_ρ_,N_θ_) for 2D filtering

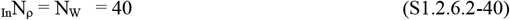

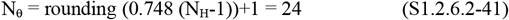

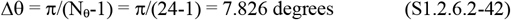 Here, Δθ is the resolution in θ orientation of the NDS simple, DS simple, and DS complex cells.
    b. Calculate an output array (_Out_N_ρ_,N_θ_) from this input array to a DS simple cell

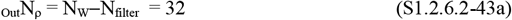

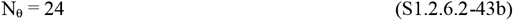
  ➁ Calculate an array (N_ρ_,N_θ_,N_τ_) of the DS simple cell

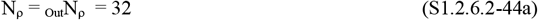

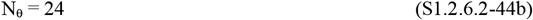

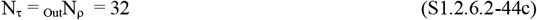
  ➂ Calculate an array (N_θ_,N_τ_) of the DS complex cell

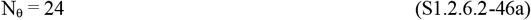

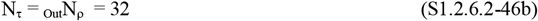
  ➃ Calculate an array (N_τx_,N_τy_) of the MDC

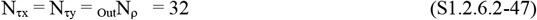

#### S1.3. Array size of SDC

The array sizes of the SDCs in Figures 9(C) and 17(C) are determined, as follows. The explanation is divided into SDCs for detecting the time-to-contact in Section **S1.3.1** and for detecting the shortest distance in Section **S1.3.2**.

##### S1.3.1. SDC array for detecting the time-to-contact and its address (k,l,h_t_)

1. The size of the SDC array shown in Figure S1-7(C) How this size is determined is described below. Each SDC_klt_ in the SDC column of Figure 9(C) is expressed by the following equation, where t is the normalized time-to-contact to a plane and n_kl_ is its 3D orientation.

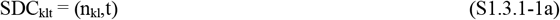
  ➀ Determine the time t by the following equation using the address h_t_ in Figure S1-7(C)

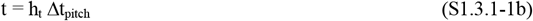 Here, parameters used above are shown as follows.

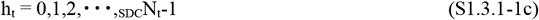

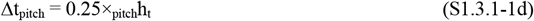

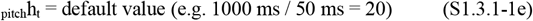

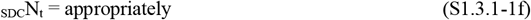
  ➁ Determine the orientation n_kl_ Figure S1-7(A) shows an eyeball that corresponds to the cross section at a height t of the SDC column in Figure 9(C) (see Figure 10). On this eyeball, SDCs are arranged at equal intervals Δβ_SDC_ that will be calculated by equation(S1.3.1-9). The (B) shows an address representation (k,l) of the (A). Stacking the (B) along the height address h_t_ yields the (C), and thus has yielded the 3D address representation (k,l,h_t_) of the SDC column in Figure 9(C). **Figure S1-7.**
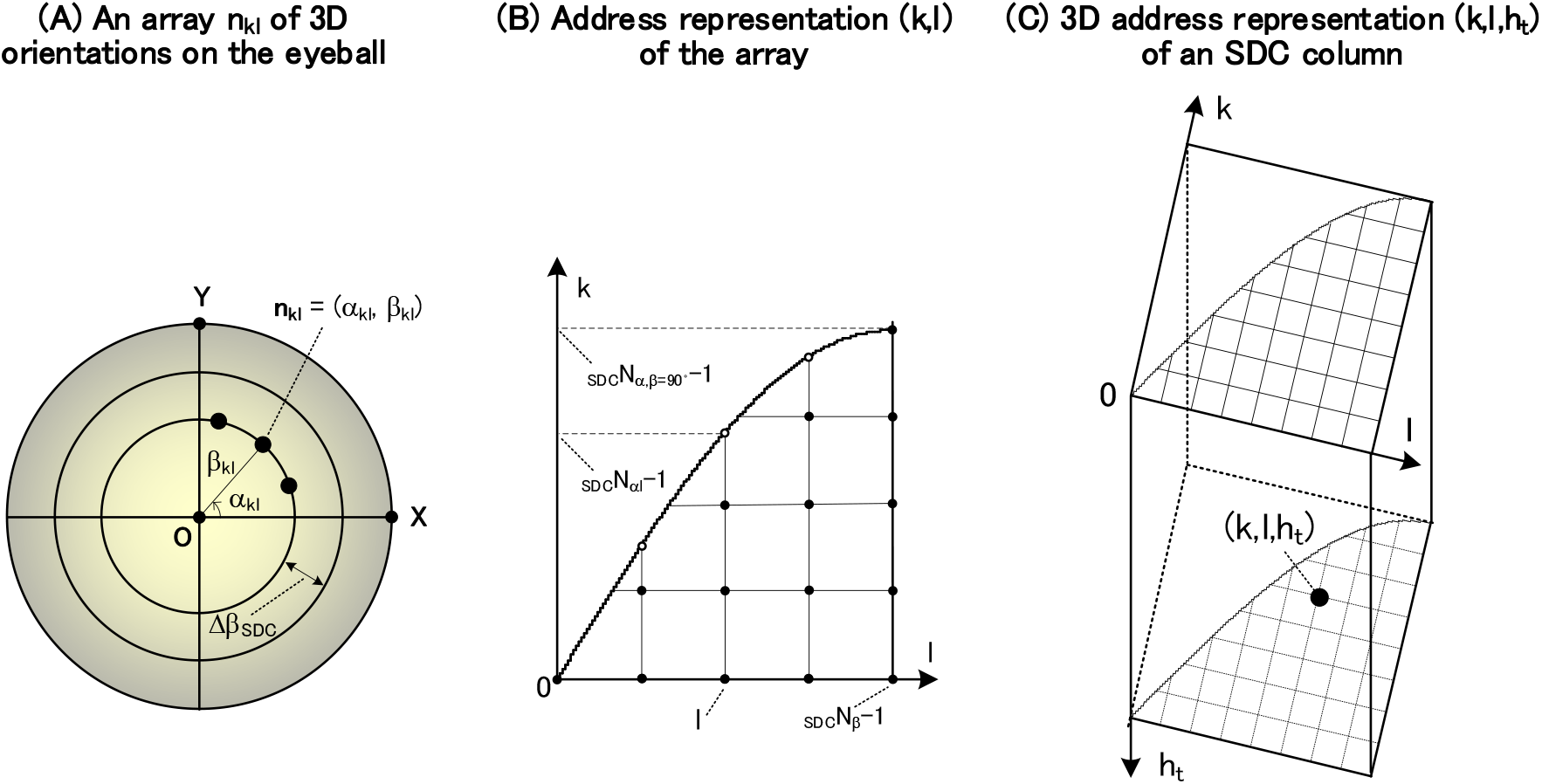
3D address representation (k,l,h_t_) of a SDC column for detecting normalized time-to-contacts and 3D orientations. (A) A SDC array for detecting the 3D orientations n_kl_ on the eyeball is shown. This array was obtained by converting each cross section of the SDC column in Figure 9(C) into the eyeball using the equidistant projection of Figure 10. On this eyeball, a 3D orientation n_kl_ (i.e. (α_kl_,β_kl_)) of each plane is arrayed at an equal interval Δβ_SDC_ that is calculatd by equation(S1.3.1-9). (B) This shows an address representation (k,l) of the array in the (A). That is, each orientation n_kl_ in the (A) is represented as an address (k,l) of it. (C) A 3D address representation (k,l,h_t_) of the SDC column in Figure 9(C) is shown. That is, this representation has been obtained by stacking the address (k,l) in the (B) along the height address h_t_. In the (A), a 3D orientation n_kl_ of each plane is expressed by the following equation.

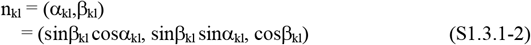 Here, (α_kl_,β_kl_) is the polar coordinate of n_kl_, (sinβ_kl_ cosα_kl_, sinβ_kl_ sinα_kl_, cosβ_kl_) is its orthogonal coordinate, and parameters used above are shown as follows.

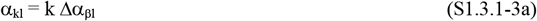

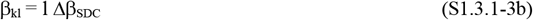

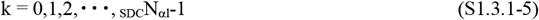

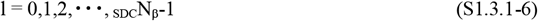 The parameters (_SDC_N_β_, Δβ_SDC_, _SDC_N_αl_, and Δα_βl_) used above will be calculated in subsection (2) below.
2. Calculate the parameters (_SDC_N_β_, Δβ_SDC_, _SDC_N_αl_, and Δα_βl_) This calculation is similar to Section **S1.2.2**(2), and Figure S1-7(A) and (B) corresponds to Figure S1-2(A) and (B) respectively.
  a. Temporarily set the latitudinal cell pitch Δβ_SDC_’
  b. Calculate the number _SDC_N_β_ of SDCs in the latitudinal direction based on this Δβ_SDC_’

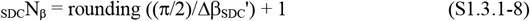
  c. Calculate finally the latitudinal pitch Δβ_SDC_ of SDCs

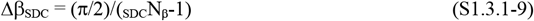
  d. Calculate the maximum number _SDC_N_α,β=90_ of SDCs in the longitude direction

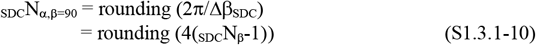
  e. Calculate the number _SDC_N_αl_ of SDCs (see Figure S1-7(B)) and the pitch Δα_βl_ in the longitude direction

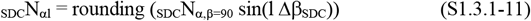

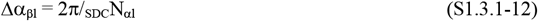 Note: These roundings that allow for equally spaced coordinates are necessary to determine the coordinates of SDCs by a modulo operation.
3. Calculate the total number _SDC_N_total_ of SDC columns at each cross-section h_t_ (see Figure S1-7(B))

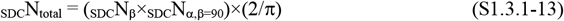 Here, this correction factor 2/π is the same as in equation(S1.2.1-10c).

##### S1.3.2. SDC array for detecting the shortest distance and its address (k,l,h_d_)

Replacing the normalized time-to-contact t in Section **S1.3.1** with the normalized shortest-distance d gives Section **S1.3.2**.

1. The size of the SDC array shown in Figure S1-8(C) How this size is determined is described below. Each SDC_kld_ in the SDC column of Figure 17(C) is expressed by the following equation, where d is the normalized shortest-distance to a plane and n_kl_ is its 3D orientation.

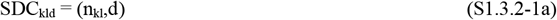
  ➀ Determine the distance d by the following equation using the address h_d_ in Figure S1-8(C)

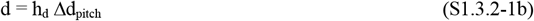 Here, parameters used above are shown as follows.

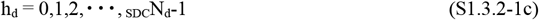

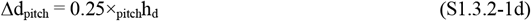

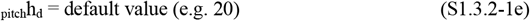

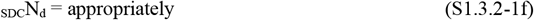
  ➁ Determine the orientation n_kl_ Figure S1-8(A) shows an eyeball that corresponds to the cross section at a height d of the SDC column in Figure 17(C) (see Figure 10). On this eyeball, SDCs are arranged at equal intervals Δβ_SDC_ that will be calculated by equation(S1.3.2-9). The (B) shows an address representation (k,l) of the (A). Stacking the (B) along the height address h_d_ yields the (C), and thus has yielded the 3D address representation (k,l,h_d_) of the SDC column in Figure 17(C). In the (A), a 3D orientation n_kl_ of each plane is expressed by the following equation.

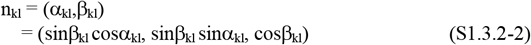 Here, (α_kl_,β_kl_) is the polar coordinate of n_kl_, (sinβ_kl_ cosα_kl_, sinβ_kl_ sinα_kl_, cosβ_kl_) is its orthogonal coordinate, and parameters used above are shown as follows.

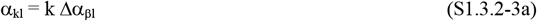

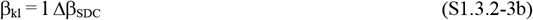

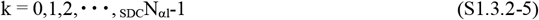

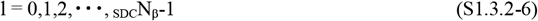 The parameters (_SDC_N_β_, Δβ_SDC_, _SDC_N_αl_, and Δα_βl_) used above will be calculated in subsection (2) below.
2. Calculate the parameters (_SDC_N_β_, Δβ_SDC_, _SDC_N_αl_, and Δα_βl_) This calculation is similar to Section **S1.2.2**(2), and Figure S1-8(A) and (B) corresponds to Figure S1-2(A) and (B) respectively.
  a. Temporarily set the latitudinal cell pitch Δβ_SDC_’
  b. Calculate the number _SDC_N_β_ of SDCs in the latitudinal direction based on this Δβ_SDC_’

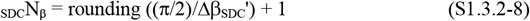
  c. Calculate finally the latitudinal pitch Δβ_SDC_ of SDCs

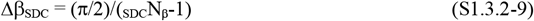
  d. Calculate the maximum number _SDC_N_α,β=90_ of SDCs in the longitude direction

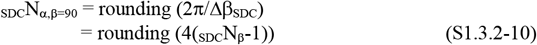
  e. Calculate the number _SDC_N_αl_ of SDCs and the pitch Δα_βl_ in the longitude direction (see Figure S1-8(B))

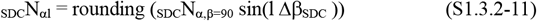

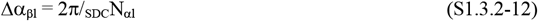 Note: These roundings that allow for equally spaced coordinates are necessary to determine the coordinates of SDCs by a modulo operation.
3. Calculate the total number _SDC_N_total_ of SDC columns at each cross-section h_d_ (see Figure S1-8(B))

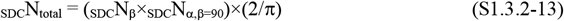

Here, this correction factor 2/π is the same as in equation(S1.2.1-10c).

### S2. Procedure for creating a connection table

The procedure for creating a connection table that represents each neural network is shown below in the form of a flowchart: this table accurately represents each network of Figure 9 or Figure 17 at the level of the neural connection of which cell is connected to which cell. This table will be used to calculate a series of cell responses in Section **S3**.

#### S2.1. Connection table for the Hough transform performed by NDS simple cells

The pink and blue straight lines in Figure S2-1(c) indicate the backward neural network from each NDS simple cell (ρ,θ) to the LGN cells {(x,y)}: {(x,y)} represents a set of (x,y). Here, Figure S2-1 has been obtained by enlarging Figure 9(B), and see Figure 24 for the forward and backward methods.

Two procedures are shown as follows. Section **S2.1.1** will show the procedure for creating a connection table that represents the backward neural network. Next, Section **S2.1.2** will show the procedure for transposing this table to create a connection table that represents the forward neural network.

Note the following. This connection table can also be expressed as a four-dimensional (4D) matrix A_SC,NDS_(ρ,θ;x,y), and then responses SC_NDS_(ρ,θ) of NDS simple cells are obtained by performing a matrix operation between this matrix and the responses LGN(x,y) of LGNcells (See page 190 of Kawakami (1996b), and Kawakami et al., 1992a). The algebraic geometry structure of this matrix is as follows.

**Figure S2-1.**
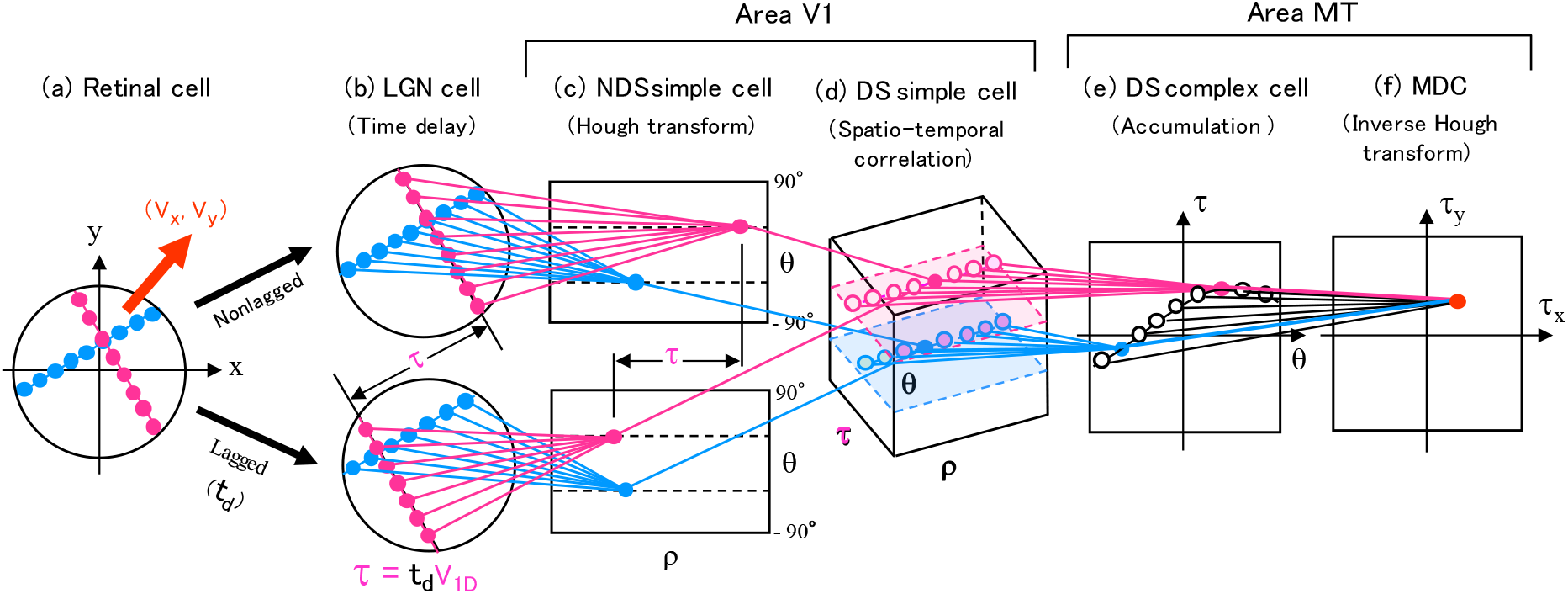
A series of modeled cells for detecting local motions. This has been obtained by enlarging Figure 9(B) (or Figure 17(B)).

This matrix constitutes a 3D hypersurface in the 4D space (ρ,θ,x,y) and is a covariant tensor of order 2 or a Grassmannian manifold of lowest order (Akizuki & Takizawa, 1957; Sasaki, 1957).

#### S2.1.1. Procedure for creating a backward connection table _I,J_T^BWD^ _SC,NDS_[(ρ,θ)→{(i,j)}]

Calculate the backward neural network from each NDS simple cell (ρ,θ) in Figure S2-1(c) to LGN cells {(x,y)} in Figure S2-1(b), and create a connection table _I,J_T^BWD^ _SC,NDS_[(ρ,θ)→{(i,j)}] representing this neural network: (i,j) is an address representation of (x,y) (see Figure S1-1). The creation procedure is shown below based on Figures S2-2 to S2-5.

Before that, the notation of this table is explained. Subscripts (_I,J_, ^BWD^, and _SC,NDS_) indicate the address of a RF O_I,J_, the backward neural network, and the NDS simple cell, respectively, and (ρ,θ)→{(i,j)} indicates a connection from each NDS simple cell (ρ,θ) in Figure S2-5(B) to the corresponding LGN cells {(i_m_’,j_m_’)} in Figure S2-5(A): note that this (i_m_’,j_m_’) is a representation on the tangent plane of the (i,j), as described below.

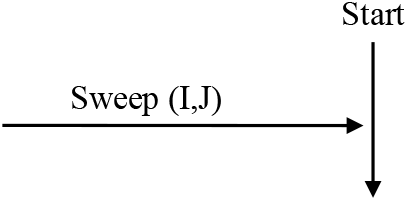

The sweep (I,J) of a RF is performed in the following using equation(S1.2.2-3~8) in Section **S1.2.2**.

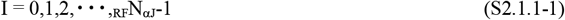

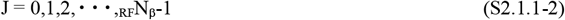

By giving Δβ_RF_’ temporarily, calculate the following (see Section **S1.2.2**(2)).

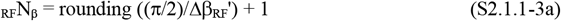

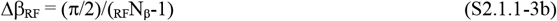

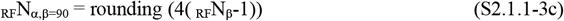

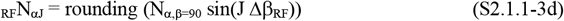

Also, calculate O_IJ_ used in equation(S2.1.1-17) in advance using the following equation (see equations(S1.2.2-1, 2, 6, and 9) in Section **S1.2.2**).

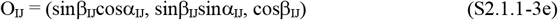

Here, (sinβ_IJ_cosα_IJ_, sinβ_IJ_sinα_IJ_, cosβ_IJ_) is the orthogonal coordinate of O_IJ_, and parameters used above are shown as follows.

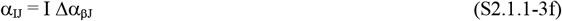

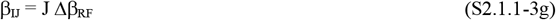

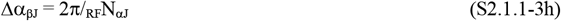

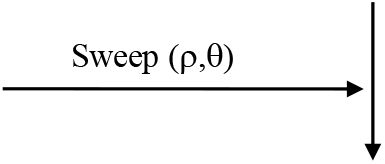

The sweep (ρ,θ) of a NDS simple cell is performed in the following.

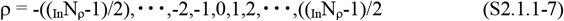

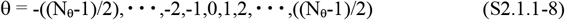

Here, the parameters (_In_N_ρ_ and N_θ_) required for this sweep were described by equations(S1.2.6.1-40 and 41), and the parameters Δθ required in subsection (4) later were described by equation(S1.2.6.1-42). These parameters are shown below.

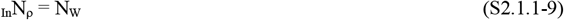

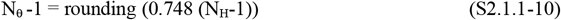

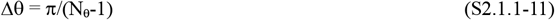

Note 1: _In_N_ρ_ is an odd number, and N_θ_ may be an even number

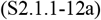

Note 2: The main parameters such as N_W_ and N_H_ were given in Section **S1.2.6.1**(2).

1. Reset a weight-evaluation variable _W_N_(ρ,θ)_ to 0

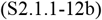 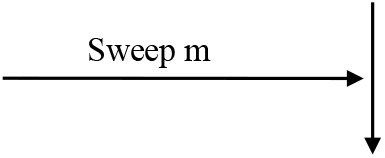 Sweep a cutout number m by the following equation.

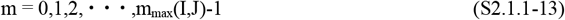 Here, m_max_(I,J) was calculated by equation(S1.2.4-14 or 26) in Section **S1.2.4**.
2. Using the cutout table _I,J_T_RF_[m→(i_m_,j_m_)] created in Section **S1.2.4**, read out the address (i_m_,j_m_) of the LGN cell that corresponds to this cutout number m.
3. Calculate the address (i_m_’,j_m_’) of a LGN cell on each tangent plane
  ➀ Calculate the red position P_m_ of the LGN cell on the eyeball in Figure S2-2 For convenience of notation, the parameters (P_ij_, α_ij_, β_ij_, i, j, Δα_βj_, and N_αj_) in Section **S1.2.1**(1) have been replaced with parameters (P_m_, α_m_, β_m_, i_m_, j_m_, Δα_βjm_, and N_αjm_), respectively. The replaced P_m_ is calculated by the following equation.

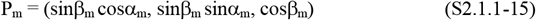

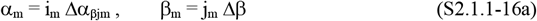 Here, the parameters (Δβ and Δα_βjm_) used above are described in the following. Temporarily give Δβ’ and calculate the following equations (see Section **S1.2.1**(2)).

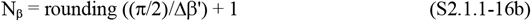

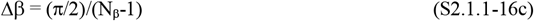

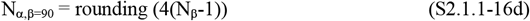

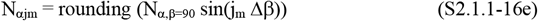

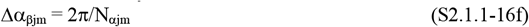
  ➁ Calculate a position P_m_’ obtained by projecting P_m_ onto the tangent plane (see Figure S2-3(A) and Figure S2-4(A))

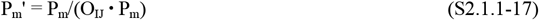
  ➂ Calculate the coordinate (x_m_’,y_m_’) of P_m_’ in Figure S2-3(B), where P_m_’ is expressed in the coordinate system (x’,y’) on the tangent plane

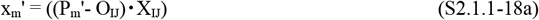

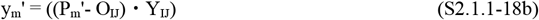 Here, X_IJ_ and Y_IJ_ used above are calculated by the following equations (see Section **S1.2.3** and Figure S1-3).

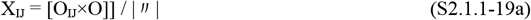

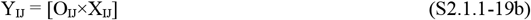 **Figure S2-2.**
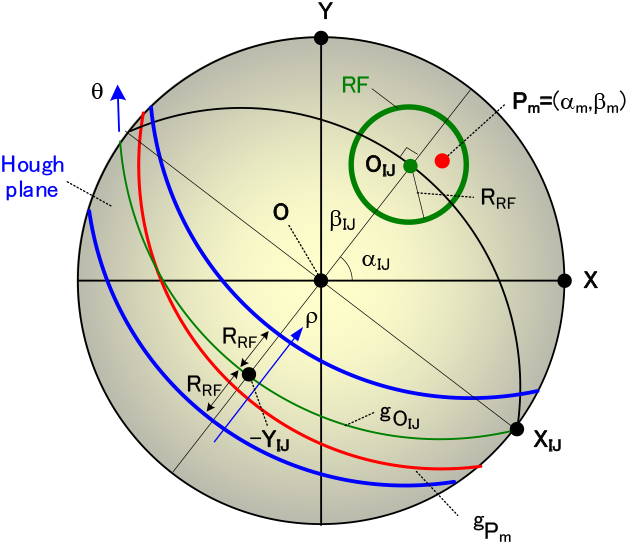
Correspondence on the eyeball between the RF and the Hough plane. This figure is used for two roles. One is used for calculating P_m_ using equation(S2.1.1-15) in Section **S2.1.1**. The other is used for supplementing the correspondence between the RF (shown in green) and the Hough plane (shown in blue) in Figure 34(B) from a different perspective. The O_IJ_ is the RF center, and the R_RF_ is the RF radius. The (O_IJ_,X_IJ_,Y_IJ_) is each RF coordinate system. The g_Pm_ and g_OIJ_ are great circles obtained by polar transformations of the P_m_ and O_IJ_, respectively.

**Figure S2-3.**
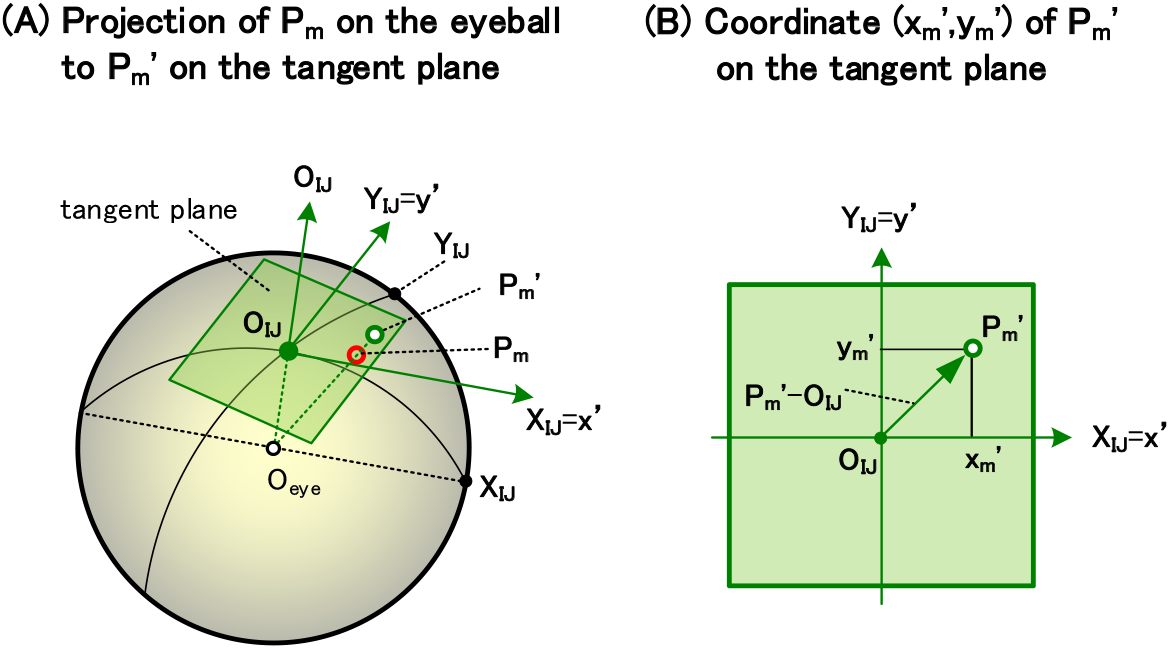
A position P_m_’ on a tangent plane and its coordinate (x_m_’,y_m_’). This figure is used for LGN cells. (A) The position P_m_’ is obtained by projecting the P_m_ of Figure S2-2 onto a tangent plane, and is calculated by equation(S2.1.1-17). (B) Calculate the coordinate (x_m_’,y_m_’) of the position P_m_’ by equations(S2.1.1-18a and 18b). Note: when O_IJ_=O, the following equations are obtained.

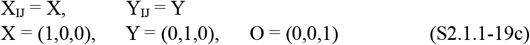
  ➃ Calculate the address representation (i_m_’,j_m_’) of the coordinate (x_m_’,y_m_’) (see Figure S2-4(D))

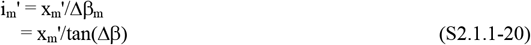

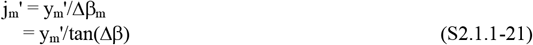 Note: (i_m_’,j_m_’) remains a real number, not integerized.
4. Confirm whether this (i_m_’,j_m_’) is inside the red elongated rectangle in Figure S2-5(A). To do this, check the following equation.

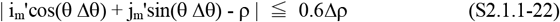

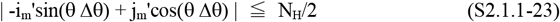 Here, Δρ used above is the default value of 1.

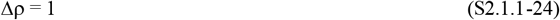 Note 1: This 0.6Δρ causes the fluctuation of the number of backward connections from each NDC simple cell to LGN cells to be less than 10%. The 0.5Δρ fluctuates considerably. Note 2: The slope of equation(S2.1.1-23) is obtained by rotating the slope θΔθ of equation(S2.1.1-22) by 90 degrees.
5. After confirming that (i_m_’,j_m_’) satisfies equations(S2.1.1-22 and 23), do the following.
  ➀ Write the address (i_m_,j_m_) of the LGN cell to the contents of the address (I,J,ρ,θ) in the table _I,J_T^BWD^ _SC,NDS_[(ρ,θ)→{(i,j)}] (see subsection (2) above for this address (i_m_,j_m_))

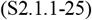
  ➁ Add 1 to the weight-evaluation variable _W_N_(ρ,θ)_

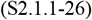 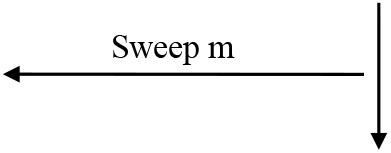
6. Calculate a weighting coefficient W_(I,J,ρ,θ)_ Calculate the following equation, where Δρ×N_H_ is the area of the red rectangle in Figure S2-5(A).

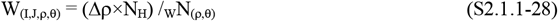 The reason for using this coefficient W_(I,J,ρ,θ)_ is to correct a variation in the area of each this red-rectangle. This weighting coefficient W_(I,J,ρ,θ)_ will be used in Section **S3.1.2**. That is, multiplying the coefficient W_(I,J,ρ,θ)_ and the responses _I,J_SC_NDS_(ρ,θ) of NDS simple cells enables these responses to be uniform. Note: If W_(I,J,ρ,θ)_ fluctuates too much, it is recommended to increase Δρ in equation(S2.1.1-24) a little (perhaps to about 1.2)

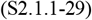 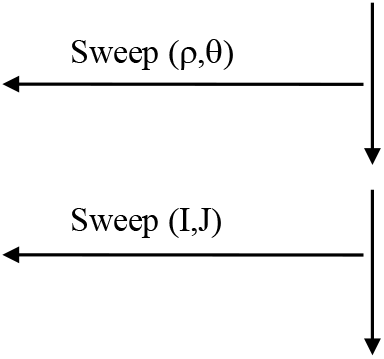
7. The backward connection table _I,J_T^BWD^ _SC,NDS_[(ρ,θ)→{(i,j)}] and weighting coefficient W_(I,J,ρ,θ)_ have been created. 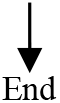

##### S2.1.2. Procedure for creating a forward connection table _I,J_T^FWD^_SC,NDS_[(i,j)→{(ρ,θ)}]

Transpose the backward connection table _I,J_T^BWD^_SC,NDS_[(ρ,θ)→{(i,j)}] obtained in Section **S2.1.1** to create a forward connection table _I,J_T^FWD^ _SC,NDS_[(i,j)→{(ρ,θ)}] that connects each LGN cell (x,y) in Figure S2-1(b) to NDS simple cells {SC_NDS_(ρ,θ)} in Figure S2-1(c): (i,j) is an address representation of (x,y) (see Figure S1-1). The creation procedure is shown below based on Figure S2-5.

**Figure S2-4.**
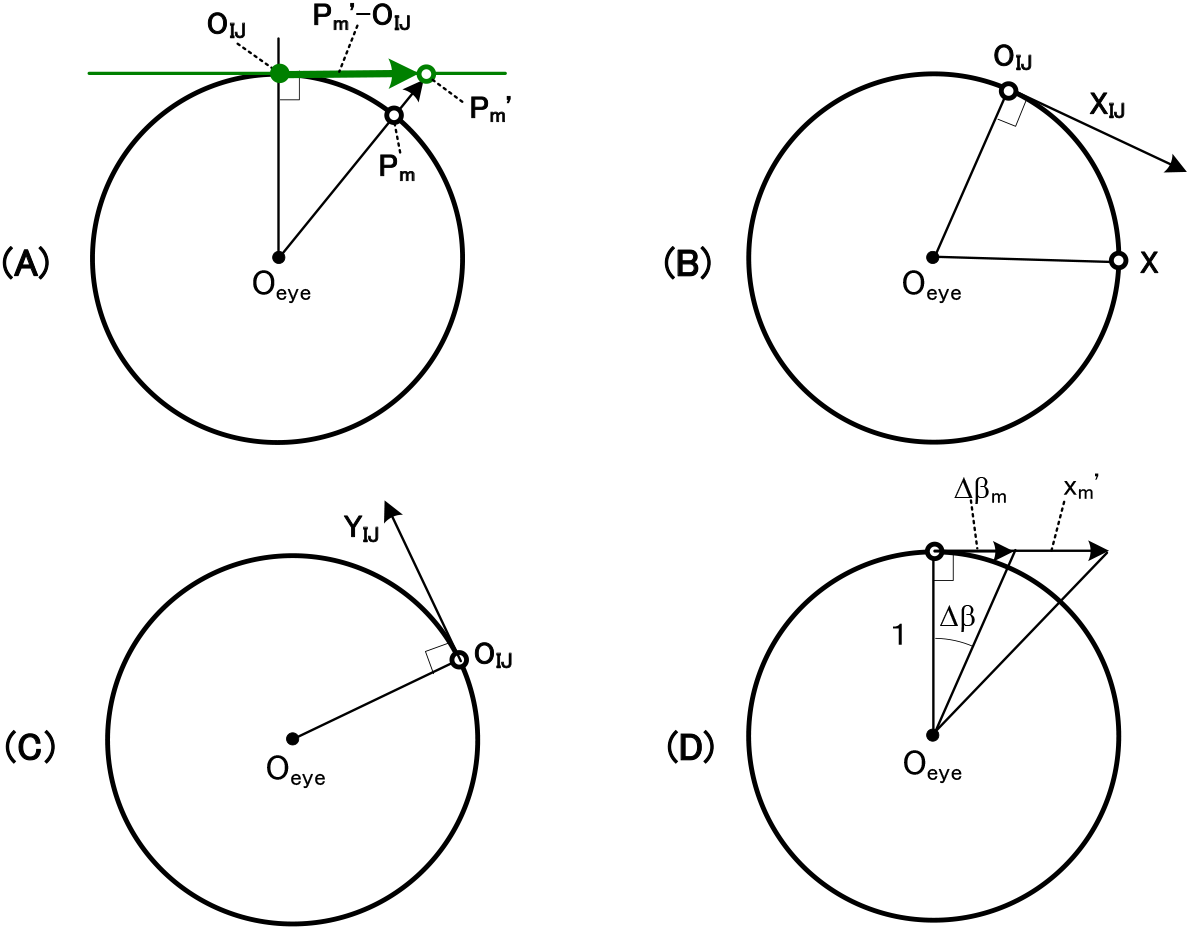
Projection onto the tangent plane. This figure is used for LGN cells. (A) Calculate the position P_m_’ in Figure S2-3(A) by equation(S2.1.1-17). (B) Calculate the RF coordinate X_IJ_ by equation(S2.1.1-19a). (C) Calculate the RF coordinate Y_IJ_ by equation(S2.1.1-19b). (D) Convert the the coordinate (x_m_’,y_m_’) of the position P_m_’ in Figure S2-3(B) to a real address (i_m_’,j_m_’) using equations(S2.1.1-20 and 21).

Before that, the notation of this table is explained. Subscripts (_I,J_, ^FWD^, and _SC,NDS_) indicate the address of a RF O_I,J_, the forward neural network, and the NDS simple cell, respectively, and (i,j)→{(ρ,θ)} indicates a connection from each LGN cell (i_m_’,j_m_’) in Figure S2-5(A) to the corresponding NDS simple cells {(ρ,θ)}: note that this (i_m_’,j_m_’) is a representation on the tangent plane of the (i,j), as described in Section **S2.1.1**.

**Figure S2-5.**
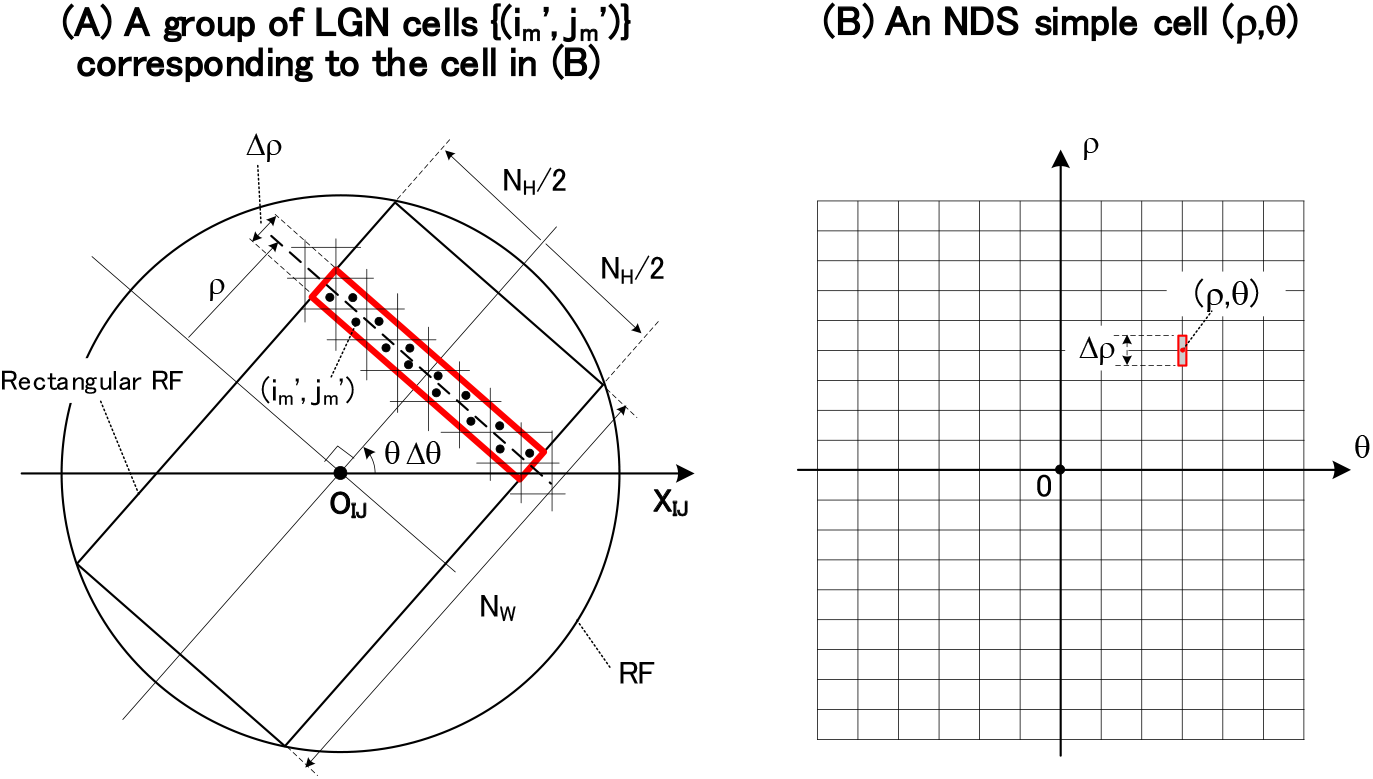
A group of LGN cells {(i_m_’, j_m_’)} corresponding to each NDS simple cell (ρ,θ). (A) A backward neural network from each NDS simple cell (ρ,θ) in the (B) to a red group of LGN cells {(i_m_’,j_m_’)} is calculated in Section **S2.1.1**. In this calculation, it is checked by equations(S2.1.1-22 and 23) whether each LGN cell (i_m_’,j_m_’) of this group is inside the red elongated-rectangle. The size of the rectangular RF (i.e. N_W_ and N_H_) is shown in Figure S1-5(A) and Δθ was calculated by equation(S1.2.6.1-42). (B) The NDS simple cell (ρ,θ) that has been used in the (A) is shown. This cell has a region Δρ in the direction ρ, which determines the width Δρ of the red rectangle in the (A).

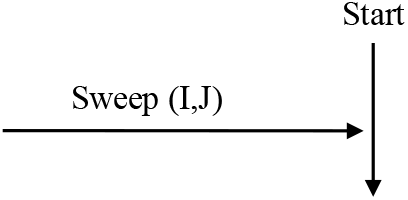

The sweep (I,J) of a RF is performed by equations(S2.1.1-1~3d) in Section **S2.1.1**.

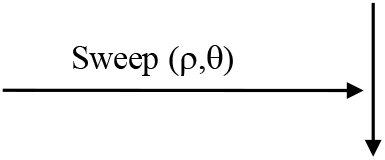

The sweep (ρ,θ) of a NDS simple cell is performed by equations(S2.1.1-7~11) of Section **S2.1.1**.

1. Using the backward connection table _I,J_T^BWD^ _SC,NDS_[(ρ,θ)→{(i,j)}] in Section **S2.1.1**, read the addresses {(i,j)} of the LGN cells corresponding to the address (I,J,ρ,θ) of each NDC simple cell.
2. Write this address (ρ,θ) of the NDS simple cell into the contents for addresses (I,J,{(i,j)}), which were read above, of the forward connection table _I,J_T^FWD^ _SC,NDS_[(i,j)→{(ρ,θ)}]. This writing has yielded the backward connection table to be converted into the forward connection table. 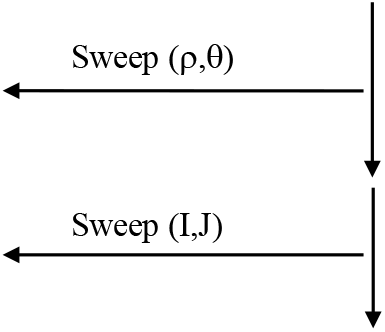
3. The forward connection table _I,J_T^FWD^ _SC,NDS_[(i,j)→{(ρ,θ)}] has been created. The descriptions of the subsections (1) and (2) above can be expressed in another way as follows. That is, by transposing (ρ,θ) and (i,j) of _I,J_T^BWD^ _SC,NDS_[(ρ,θ)→{(i,j)}] in Section **S2.1.1**, we can get_I,J_T^FWD^ _SC,NDS_ [(i,j)→{(ρ,θ)}]. 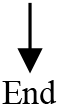

### S2.2. Connection table for the inverse Hough transform performed by MDCs

The straight lines containing pink and blue in Figure S2-1(f) indicate the backward neural network from each MDC (τ_x_,τ_y_) to the DS complex cells {(θ,τ)}.

Two procedures are shown as follows. Section **S2.2.1** will show the procedure for creating a connection table that represents this backward neural network. Next, Section **S2.2.2** will show the procedure for transposing this table to create a connection table that represents the forward neural network.

- Note 1: This table is common for every RF, so it does not depend on (I,J)

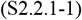
- Note 2: The properties of DS simple and DS complex cells are also independent of (I,J)

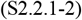 Note the following. This connection table for MDCs can also be expressed as a 4D matrix A_MDC_(τ_x_,τ_y_; θ,τ), and then responses MDC(τ_x_,τ_y_) of MDCs are obtained by performing a matrix operation between this matrix and the responses CC_DS_(θ,τ) of DS complex cells (See page 193 of Kawakami(1996b)). The algebraic geometry structure of this matrix is as follows. This matrix constitutes a 3D hypersurface in the 4D space (τ_x_,τ_y_,θ,τ) and is a contravariant tensor of order 2 or a Grassmannian manifold of lowest order (Akizuki & Takizawa, 1957; Sasaki, 1957).

#### S2.2.1. Procedure for creating a backward connection table T^BWD^_MDC_[(τ_x_,τ_y_)→{(θ,τ)}]

Calculate the backward neural network from each MDC (τ_x_,τ_y_) in Figure S2-1(f) to DS complex cells {(θ,τ)} in Figure S2-1(e), and create a connection table T^BWD^_MDC_[(τ_x_,τ_y_)→{(θ,τ)}] representing this neural network. The creation procedure is shown below based on Figure S2-6.

Before that, the notation of this table is explained. Subscripts (^BWD^ and _MDC_) indicate the backward neural network and the MDC, respectively, and (τ_x_,τ_y_)→{(θ,τ)} indicates a connection from each MDC (τ_x_,τ_y_) in Figure S2-6(B) to the corresponding DS complex cells {(θ,τ)} in Figure S2-6(A).

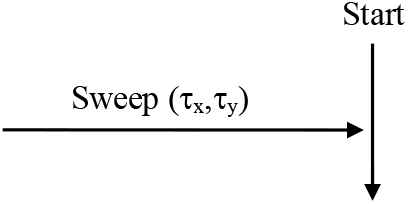

The sweep (τ_x_,τ_y_) of a MDC is performed in the following.

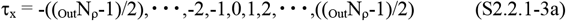

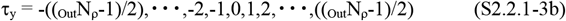

Here, the parameter _Out_N_ρ_ used was described in equation(S1.2.6.1-43). It is shown in the following equation.

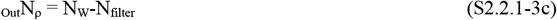

The main parameters such as N_W_ and N_filter_ were given in Section **S1.2.6.1**(2).

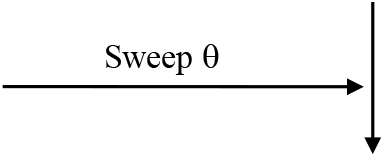

The sweep θ of a DS complex cell (θ,τ) is performed as follows.

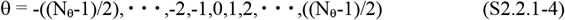

Here, the parameter N_θ_ used was described by equation(S1.2.6.1-41), and the parameter Δθ used in equation(S2.2.1-6) was described by equation(S1.2.6.1-42). These parameters are shown in the following equation.

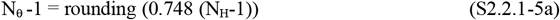

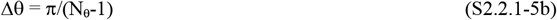

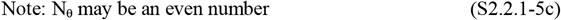

(2) By the following equation, calculate the intersections (τ^+^,τ^-^) of the red sine waves and each θ, where τ^+^ and τ^-^ refer to Figure S2-6(A).

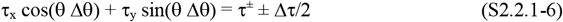

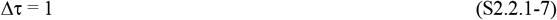 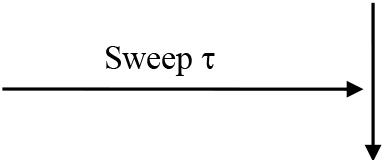 The sweep τ of a DS complex cell (θ,τ) is performed as follows.

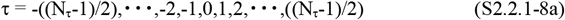 Here, the parameter N_τ_ used above was described in equation(S1.2.6.1-45b). It is shown in the following equation.

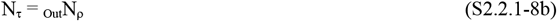

(3) For a lattice point τ that satisfy τ^-^≦τ≦τ^+^, do the following ➀ (see Figure S2-6(A)).

➀ Write the address (θ,τ) to the contents of address (τ_x_,τ_y_) of the connection table T^BWD^_MDC_ [(τ_x_, τ_y_)→{(θ,τ)}]

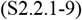 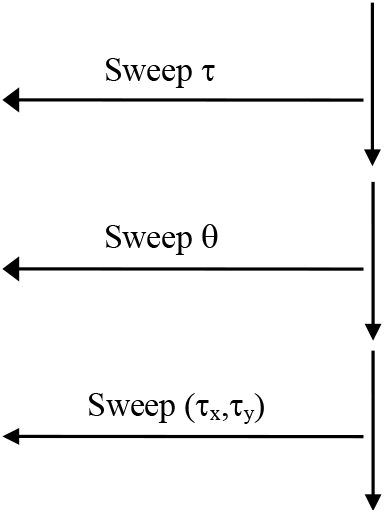

(4) The backward connection table T^BWD^ _MDC_ [(τ_x_, τ_y_)→{(θ,τ)}] has been created. 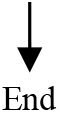

#### S2.2.2. Procedure for creating a forward connection table T^FWD^_MDC_[(θ,τ)→{(τ_x_,τ_y_)}]

Transpose the backward connection table T^BWD^_MDC_[(τ_x_,τ_y_)→{(θ,τ)}] obtained in Section **S2.2.1** to create a forward connection table T^FWD^ _MDC_ [(θ,τ)→{(τ_x_, τ_y_)}] that conncts each DS complex cell (θ,τ) to MDCs {MDC(τ_x_,τ_y_)}. The creation procedure is shown below.

**Figure S2-6.**
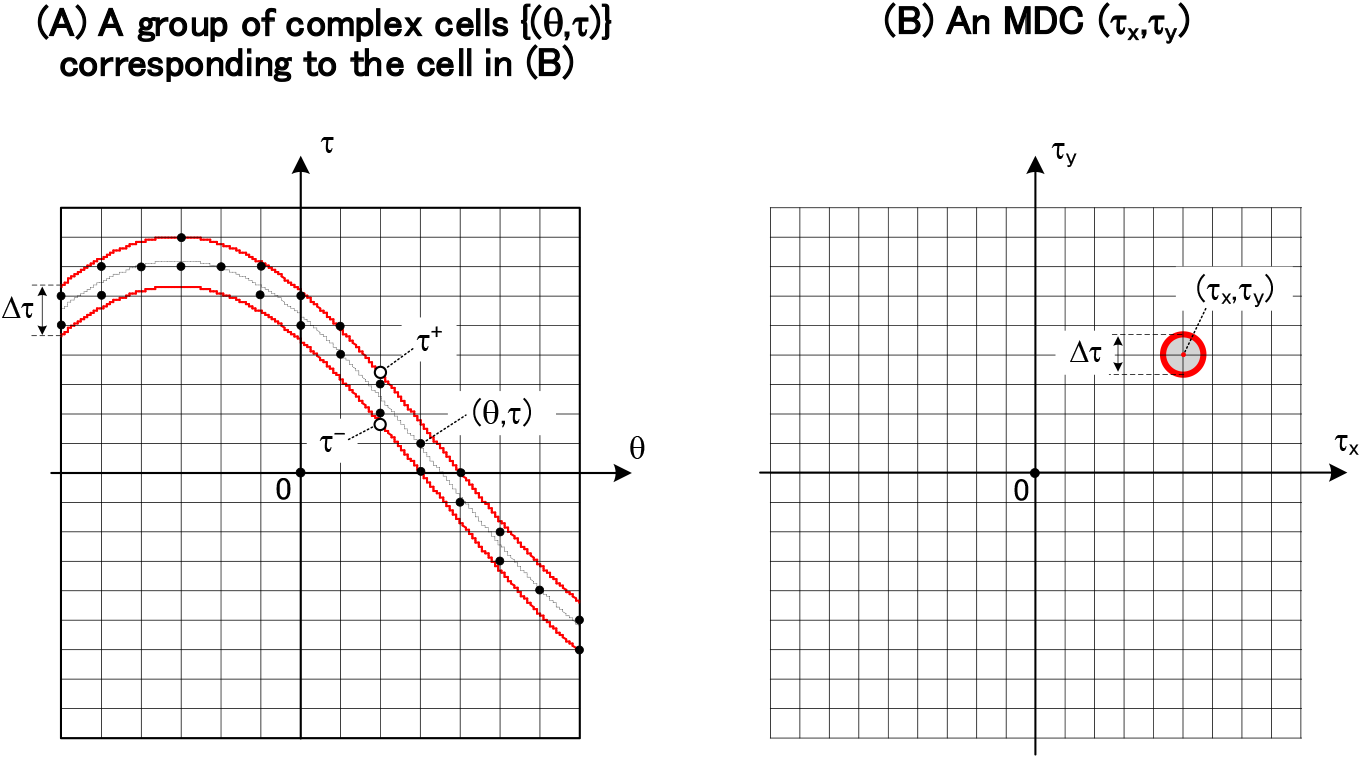
A group of complex cells {(θ,τ)} corresponding to each MDC (τ_x_,τ_y_). (A) A backward neural network from each MDC (τ_x_,τ_y_) in the (B) to a red group of DS complex cells {(θ,τ)} is calculated in Section **S2.2.1**: this group is inside the red sine-wave with a width Δτ. In this calculation, it is checked by τ^-^≦τ≦τ^+^ in Section **S2.2.1**(3) whether each DS complex cell (θ,τ) of this group is inside this red sine-wave. (B) The MDC (τ_x_,τ_y_) that has been used in the (A) is shown. This MDC has a region Δτ, which determines the width Δτ of the sine wave in the (A).

**Figure S2-7.**
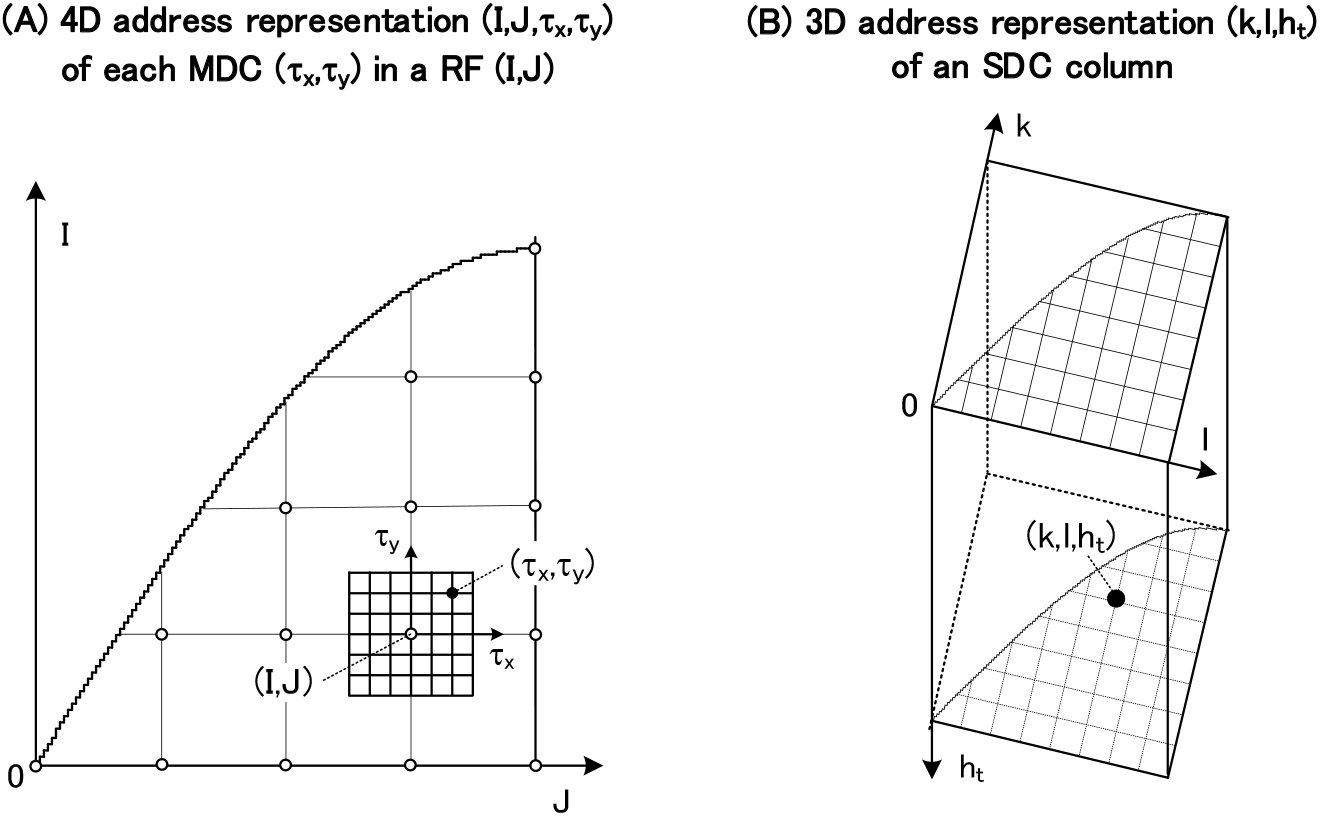
4D address representation (I,J,τ_x_,τ_y_) of each MDC and 3D address representation (k,l,h_t_) of an SDC column for detecting normalized time-to-contacts. (A) A 4D address representation (I,J,τ_x_,τ_y_) for each MDC (τ_x_,τ_y_) in a RF (I,J) is explained as follows: this MDC corresponds to that in Figure 9(B)(f), and this RF (I,J) corresponds to the RF center O_IJ_. Each white circle represents the RF center O_IJ_ at a grid point (I,J). Then, arrange an array of MDCs (τ_x_,τ_y_) centering on this grid point (I,J). Thus, this arrangement enables each MDC (τ_x_,τ_y_) in a RF (I,J) to be represented as the 4D address (I,J,τ_x_,τ_y_). (B) This is is the same as Figure S1-7(C), showing the 3D address representation (k,l,h_t_) of the SDC column in Figure 9(C).

**Figure S2-8.**
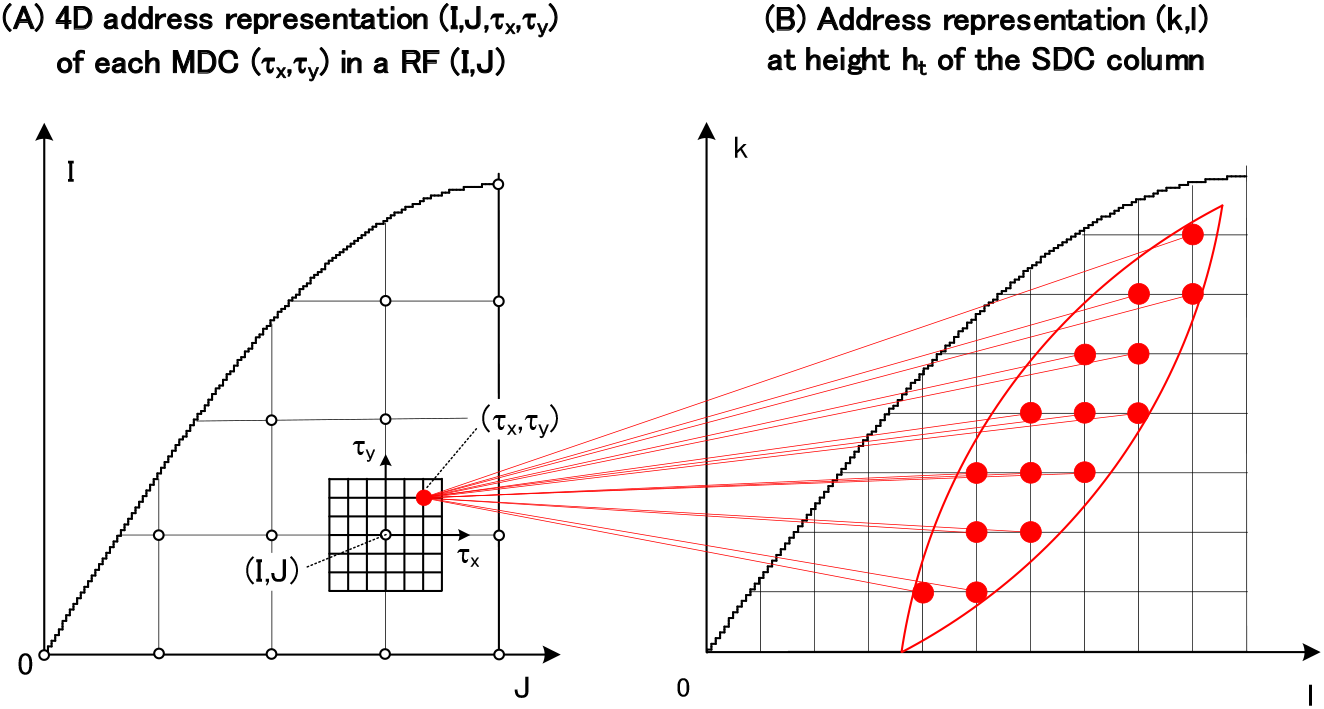
Forward connections from each MDC to a group of SDCs that detect normalized time-to-contacts. Red straight lines represent a forward neural network from each MDC (I,J,τ_x_,τ_y_) in the (A) to the group of SDCs {(k,l)} in the (B): this group {(k,l)} corresponds to the address represention of the red crescent in Figure 25(A)(ii). A connection table representing this neural network is calculated in Section **S2.3**.

**Figure S2-9.**
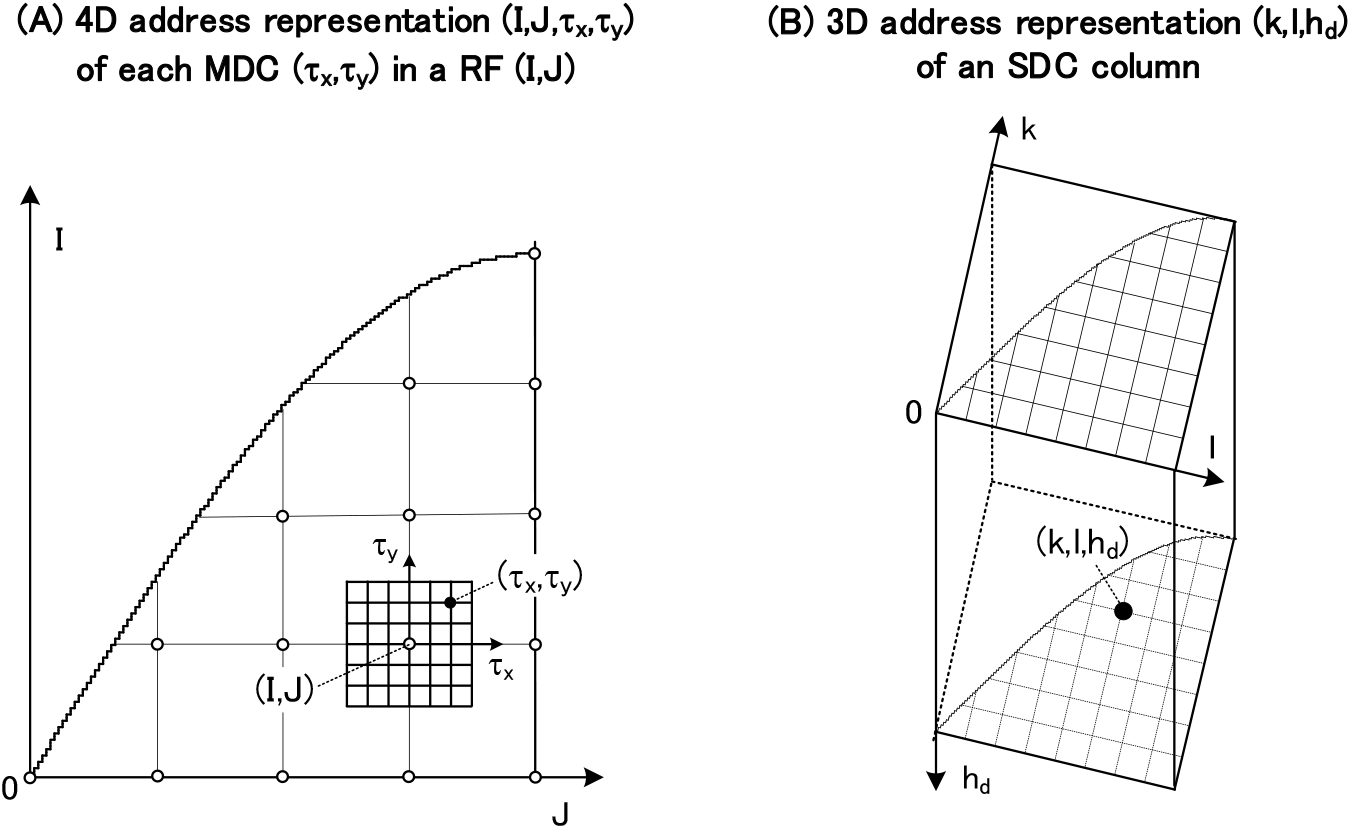
4D address representation (I,J,τ_x_,τ_y_) of each MDC and 3D address representation (k,l,h_d_) of an SDC column for detecting normalized shortest-distances. (A) This is is the same as Figure S2-7(A), showing the 4D address representation (I,J,τ_x_,τ_y_) of a MDC in Figure 17(B)(f). (B) This is is the same as Figure S1-8(C), showing the 3D address representation (k,l,h_d_) of the SDC column in Figure 17(C).

**Figure S2-10.**
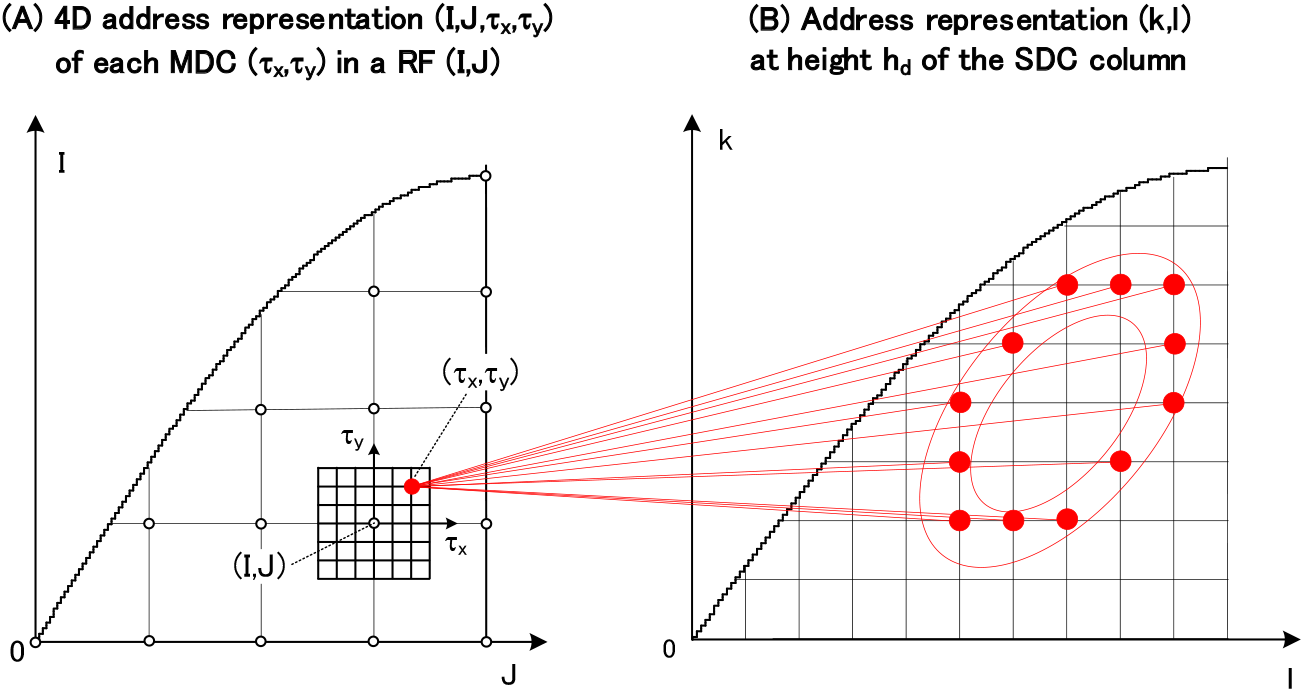
Forward connections from each MDC to a group of SDCs that detect normalized shortest-distances. Red straight lines represent a forward neural network from each MDC (I,J,τ_x_,τ_y_) in the (A) to the group of SDCs {(k,l)} in the (B): this group {(k,l)} corresponds to the address represention of the red ring in Figure 26(A)(ii). A connection table representing this neural network is calculated in Section **S2.4**.

Before that, the notation of this table is explained. Subscripts (^FWD^ and _MDC_) indicate the forward neural network and the MDC, respectively, and (θ,τ)→{(τ_x_,τ_y_)} indicates a connection from each DS complex cell (θ,τ) to the corresponding MDCs {(τ_x_,τ_y_)}.

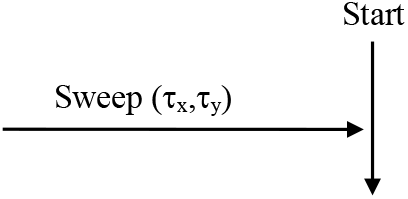

The sweep (τ_x_,τ_y_) of a MDC is performed by equations(S2.2.1-3a∼3c) in Section **S2.2.1**.

(1) Using the backward connection table T^BWD^ _MDC_ [(τ, τ)→{(θ,τ)}] in Section **2.2.1**, read the addresses {(θ,τ)} of the DS complex cells corresponding to the address (τ_x_,τ_y_) of each MDC.
(2) Write this address (τ_x_,τ_y_) of the MDC into the contents for addresses {(θ,τ)}, which were read above, of the forward connection table T^FWD^ _MDC_ [(θ,τ)→{(τ, τ)}]. This writing has yielded the backward connection table to be transposed into the forward connection table. 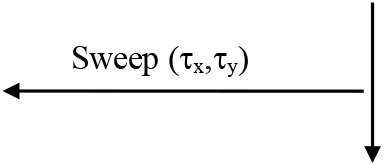
(3) The forward connection table T^FWD^ _MDC_ [(θ,τ)→{(τ, τ)}] has been created. The descriptions of the subsections (1) and (2) above can be expressed in another way as follows. That is, by transposing (τ_x_,τ_y_) and (θ,τ) of T^BWD^_MDC_ [(τ_x_,τ _y_)→{(θ,τ)}] in Section **2.2.1**, we can get T^FWD^ _MDC_ [(θ,τ)→{(τ, τ)}]. 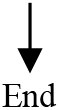

### S2.3. Connection table for the cross-ratio and polar transforms performed by SDCs that detect the time-to-contact and 3D orientation

Figure S2-8 has been obtained by drawing Figures 9(B)(f) and 9(C) (or Figure 25(A)) with the address representation. For details, Figure S2-8(A) shows that each MDC (τ_x_,τ_y_) in a RF O_IJ_ of Figure 9(B)(f) is represented as an address (I,J,τ_x_,τ_y_), and Figure S2-8(B) shows that each cross section of the SDC column (n,t) at a height t of Figure 9(C) is represented as an array of addresses (k,l). Such address representations enable a forward neural network in Figure S2-8 to be expressed as red straight lines that connects each MDC (I,J,τ_x_,τ_y_) to the corresponding SDCs {(k,l)}. Thus, these red straight lines represent the forward neural network, and the blue straight lines in Figure 25(A) has been represented as these red ones using the address representations above.

Inverting Figure S2-8 from forward to backward enables a backward neural network from each SDC (k,l) to MDCs {(I,J,τ_x_,τ_y_)} to be obtained (see Section **4.3.3.1**(3)). A series of equations, which determines this backward network, was described in Section **A9.1** of Appendix **9**.

Three procedures are performed as follows. First, the equations in Section **A9.1** enables a connection table representing the backward neural network to be created: why this backward method in Section **A9.1** has been used here was described in Section **4.3.3.1**(3); this is because the backward method is significantly simpler than the forward method. Based on these equations, a procedure for creating this backward table will be shown in Section **S2.3.1**. Next, by transposing this backward table, Section **S2.3.2** will show a procedure for creating a connection table representing a forward neural network network. Finally, the procedure for creating a weighting factor will be shown in Section **S2.3.3**.

#### S2.3.1. Procedure for creating a backward connection table _time_T^BWD^_SDC_ [(k,l,h_t_)→{(I,J,τ_x_,τ_y_)}]

Calculate the backward neural network from each SDC (k,l,h_t_) to MDCs {(I,J,τ_x_,τ_y_)} and create a connection table _time_T^BWD^_SDC_ [(k,l,h_t_)→{(I,J,τ_x_,τ_y_)}] representing this neural network (see Figure S2-8). The creation procedure is shown below based on Section **A9.1** of Appendix **9**.

Before that, the notation of this table is explained. Subscripts (_time_, ^BWD^, and _SDC_) indicate the time-to-contact, the backward neural network, and the SDC, respectively, and (k,l,h_t_)→{(I,J,τ_x_,τ_y_)} indicates a connection from each SDC (k,l,h_t_) to the corresponding MDCs {(I,J,τ_x_,τ_y_)}.

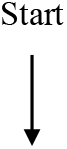

1. Give a movement direction v (see Figure 8)
  ➀ With respect to the center of the visual field O, if the movement direction v is (O • v) >0, it is an approaching movement, and if (O • v) <0, it is a separating movement

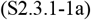
  ➁ Calculate the P_∞_ by equation(1.0-1)

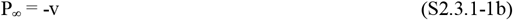 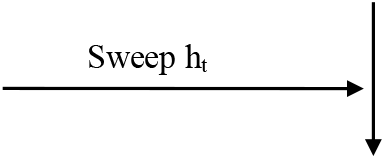 The sweep h_t_ of a SDC (k,l,h_t_) is performed by the following equation (see Section **S1.3.1**(1)).

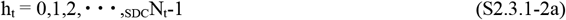

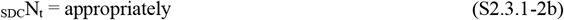
2. Convert h_t_ to t (see Section **S1.3.1**(1))

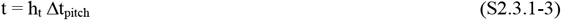 Here,

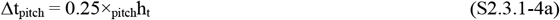

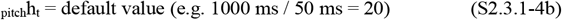 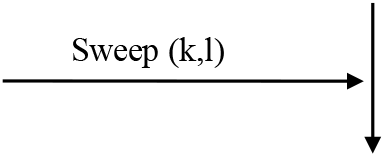 The sweep (k,l) of a SDC (k,l,h_t_) is performed as follows

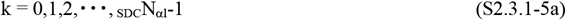

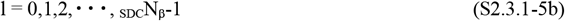 Here, the parameters (_SDC_N_β_and _SDC_N_αl_) required for this sweep were described by equations(S1.3.1-8 and 11), and the parameters required later (Δβ_SDC_ and Δα_βl_) were described by equations(S1.3.1-9 and 12). These parameters are shown as follows. Given Δβ_SDC_’ temporarily, calculate them as follows (see Section **S1.3.1**(2)).

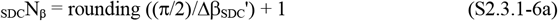

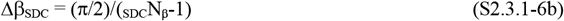

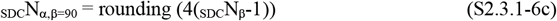

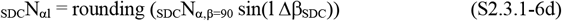 Here, if l=0, _SDC_N_αl_= 1

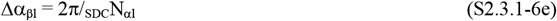 Note: These roundings that allow for equally spaced coordinates are necessary to determine the coordinates of SDCs by a modulo operation.
3. Calculate the 3D orientation n_kl_ (see Section **S1.3.1**(1))

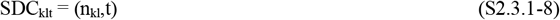

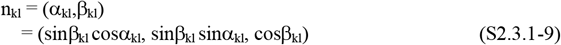 Here, (α_kl_,β_kl_) is the polar coordinate of n_kl_, (sinβ_kl_ cosα_kl_, sinβ_kl_ sinα_kl_, cosβ_kl_) is its orthogonal coordinate, and parameters used above are shown as follows.

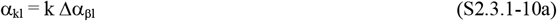

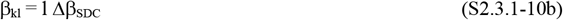 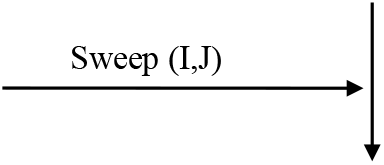 The sweep (I,J) of a RF is performed by equations(S2.1.1-1∼3d) of Section **S2.1.1**.
4. Calculate the RF center O_IJ_ (see equations(S1.2.2-1∼4) in Section

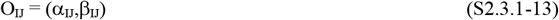

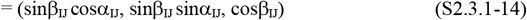 Here, (α_IJ_,β_IJ_) is the polar coordinate of O_IJ_, (sinβ_IJ_ cosα_IJ_, sinβ_IJ_ sinα_IJ_, cosβ_IJ_) is its orthogonal coordinate, and parameters used above are shown as follows.

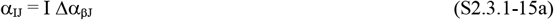

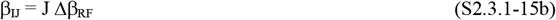

where Δβ_RF_ and Δα_βJ_ were calculated by the equations(S2.1.1-3b and 3h) in Section **S2.1.1**, respectively.
5. Calculate the position P_0_ at a current time (see equation(A9-2) in Appendix **9**)

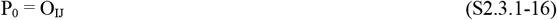
6. Calculate P_t_ (see equations(A9-3 and 4) in Appendix **9**)

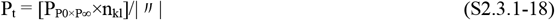

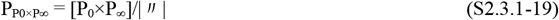
7. Calculate the polar coordinate ϕ of a MDC (see equations(A9-8∼10) in Appendix **9**)

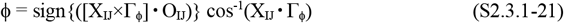 Here,

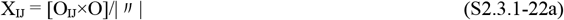

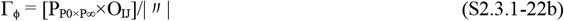
8. Calculate the polar coordinate τ_2D_ of the MDC by the cross-ratio transform (see equations(A9-5 to 7) in Appendix **9**)

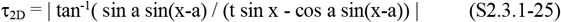 Here, the central angles a and x used above (see Figure 8(B)) are calculated by the following equations.

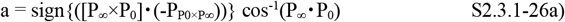

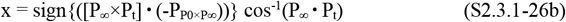
9. Calculate the address (τ_x_,τ_y_) of the MDC (see equations(A9-11 and 12) in Appendix **9**)

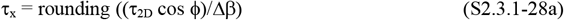

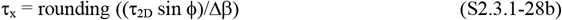 Note: This rounding serves to round to the nearest MDC, and the Δβ?used above was calculated by equation(S1.2.6.1-4).
10. When the following conditions are satisfied, write the address (I,J,τ_x_,τ_y_) to the contents of the address (k,l,h_t_) in the backward connection table _time_T^BWD^ _SDC_ [(k,l,h_t_)→{(I,J,τ_x_,τ_y_)}].

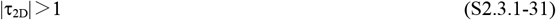

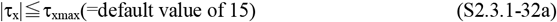

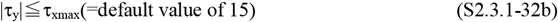 Note: This default value varies depending on the retinal cell resolution, RF size, etc. 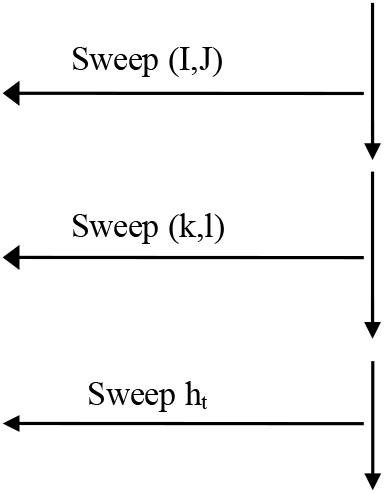
11. The backward connection table _time_T^BWD^ _SDC_ [(k,l,h_t_)→{(I,J,τ_x_,τ_y_)}] has been created (see Figure S2-8) 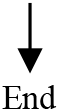

#### S2.3.2. Procedure for creating a forward connection table _time_T^FWD^_SDC_[(I,J,τ_x_,τ_y_)→{(k,l,h_t_)}]

Transpose the backward connection table _time_T^BWD^_SDC_[(k,l,h_t_)→{(I,J,τ_x_,τ_y_)}] obtained in Section **S2.3.1** to create a forward connection table _time_T^FWD^ _SDC_?[(I,J,?τ_x_ τ_y_) →{(k,l,h_t_)}] that conncts each MDC (τ_x_,τ_y_) to SDCs {(k,l,h_t_)}. The creation procedure is shown below.

Before that, the notation of this table is explained. Subscripts (_time_, ^FWD^, and _SDC_) indicate the time-to-contact, the forward neural network, and the SDC, respectively, and (I,J,τ_x_,τ_y_)→{(k,l,h_t_)} indicates a connection from each MDC (I,J,τ_x_,τ_y_) to the corresponding SDCs {(k,l,h_t_)}.

Note the following. Section **2.2** showed how to create the forward connection table using the forward method without going through this backward method. However, this forward method is more complex than the backward method, as described in Section **4.3.3.1**(3). Therefore, I decided in Section **S2.3.1** above to use the backward method in Section **A9.1** of Appendix **9**.

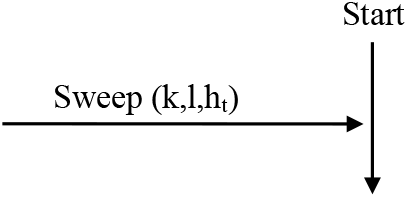

The sweeps h_t_ and (k,l) of a SDC (k,l,h_t_) are performed by equations(S2.3.1-2a and 2b) and equations(S2.3.1-5a∼6d) in Section **S2.3.1**, respectively.

1. Using the backward connection table _time_T^BWD^ _SDC_ [(k,l,h_t_)→{(I,J,τ_x_,τ_y_)}] in Section **S2.3.1**, read the addresses {(I,J,τ_x_,τ_y_)} of MDCs corresponding to the address (k,l,h_t_) of each SDC.
2. Write this address (k,l,h_t_) of the SDC into the contents for addresses {(I,J,τ_x_,τ_y_)}, which were read above, of the forward connection table _time_T^FWD^_SDC_ [(I,J,τ_x_,τ_y_)→{(k,l,h_t_)}]. This writing has yielded the backward connection table to be transposed into the forward connection table. 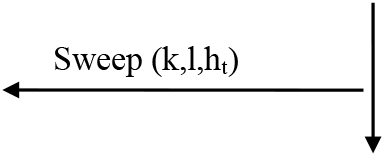
3. The forward connection table _time_T^FWD^ _SDC_ [(I,J,τ_x_,τ_y_)→{(k,l,h_t_)}] has been created. The descriptions of the subsections (1) and (2) above can be expressed in another way as follows. That is, by transposing (k,l,h_t_) and (I,J,τ_x_,τ_y_) of _time_T^BWD^ _SDC_ [(k,l,h_t_)→{(I,J,τ_x_,τ_y_)}] in Section **S2.3.1**, we can _time_T^FWD^ _SDC_ [(I,J,τ_x_,τ_y_)→{(k,l,h_t_)}]. 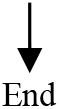

#### S2.3.3. Procedure for creating a weighting factor _time_W_(I,J,τx,τy,ht)_

The divergent neural network, indicated by red straight lines in Figure S2-8, varies considerably from MDC to MDC. This weighting factor _time_W_(I,J,τx,τy,ht)_ is needed to correct for that variation. That is, each SDC response is corrected so that it is not affected by the extent of this divergence (i.e. the number of red straight lines).

For that purpose, the number of these straight lines will be counted as a variable _W_N_(I,J,τx,τy,ht)_ in equation(S2.3.3-1) later: this variable corresponds to the area of the red crescent in Figure 25(A)(ii). Then, the weighting factor _time_W _(I,J,τx,τy,ht)_ will be calculated by equation(S2.3.3-2) as the reciprocal of this variable _W_N_(I,J,τx,τy,ht)_. The procedure for calculating this weighting factor will be shown below.

Note the following. This weighting factor will be used in Section **S3.2.1** as follows. By multiplying each MDC response by this weighting factor, the response is corrected so that it does not depend on the number of the red straight lines above. This correction is performed using Figure S2-8 as follows. Let us make the following correction so that each MDC response in the (A) is output to the red SDCs in the (B), regardless of the number of the red straight lines. When the number of straight lines is large, the MDC response is output weakened in inverse proportion to the number, and conversely, when the number of straight lines is small, the MDC response is output strengthened in inverse proportion to the number. In this way, a correction is made so that each MDC outputs a constant response to the red SDCs.

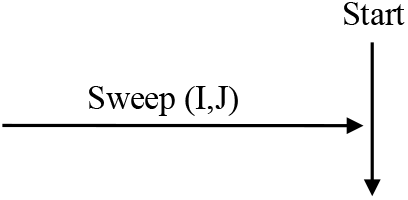

The sweep (I,J) of a RF is performed by equations(S2.1.1-1∼3d) in Section **S2.1.1**.

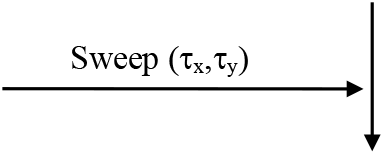

The sweep (τ_x_,τ_y_) of a MDC is performed by equations(S2.2.1-3a∼3c) in Section **S2.2.1**.

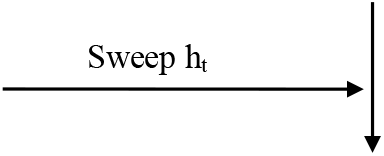

The sweep h_t_ of a SDC (k,l,h_t_) is performed by equations(S2.3.1-2a∼2b) in Section **S2.3.1**.

1. Using the forward connection table _time_T^FWD^_SDC_ [(I,J,τ_x_,τ_y_)→{(k,l,h_t_)}] in Section **S2.3.2**, read the addresses {(k,l,h_t_)} of the SDCs corresponding to the address (I,J,τ_x_,τ_y_) of each MDC.
2. Calculate the number of SDC addresses {(k,l)} at height h_t_, read above, and write the number to a variable _W_N_(I,J,τx,τy,ht)_. _W_N_(I,J,τx,τy,ht)_= the number of SDC addresses {(k,l)} at height h_t_

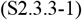
3. Determine the weighting factor _time_W_(I,J,τx,τy,ht)_ by the following equation

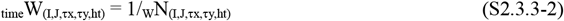 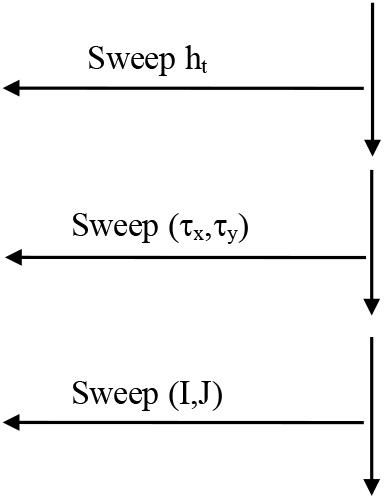
4. The weighting factor _time_W_(I,J,τx,τy,ht)_ has been created 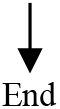

### S2.4. Connection table for the small-circle transform performed by SDCs that detect the shortest distance and 3D orientation

Figure S2-10 has been obtained by drawing Figures 17(B)(f) and 17(C) (or Figure 26(A)) with the address representation. For details, Figure S2-10(A) shows that each MDC (τ_x_,τ_y_) in a RF O_IJ_ of Figure 17(B)(f) is represented as an address (I,J,τ_x_,τ_y_), and Figure S2-10(B) shows that each cross section of the SDC column (n,d) at a height d of Figure 17(C) is represented as an array of addresses (k,l). Such address representations enable a forward neural network in Figure S2-10 to be expressed as red straight lines that connects each MDC (I,J,τ_x_,τ_y_) to the corresponding SDCs {(k,l)}. Thus, these red straight lines represent the forward neural network, and the blue straight lines in Figure 26(A) has been represented as these red ones using the address representations above.

Inverting Figure S2-10 from forward to backward enables the backward neural network from each SDC (k,l) to MDCs {(I,J,τ_x_,τ_y_)} to be obtained (see Section **4.3.3.2**(3)). A series of equation, which determines this backward network, was described in Section **A9.2** of Appendix **9**.

Three procedures are performed as follows. First, the equations in Section **A9.2** enables a connection table representing the backward neural network to be created: why this backward method in Section **A9.2** has been used here was described in Section **4.3.3.2**(3); this is because the backward method is significantly simpler than the forward method. Based on these equations, a procedure for creating this backward table will be shown in Section **S2.4.1**. Next, by transposing this backward table, Section **S2.4.2** will show a procedure for creating a connection table representing a forward neural network network. Finally, the procedure for creating a weighting factor will be shown in Section **S2.4.3**.

#### S2.4.1. Procedure for creating a backward connection table _distance_T^BWD^_SDC_ [(k,l,h_d_)→{(I,J,τ_x_,τ_y_)}]

Calculate the backward neural network from each SDC (k,l,h_d_) to MDCs {(I,J,τ_x_,τ_y_)} and create a connection table _distance_T^BWD^ _SDC_ [(k,l,h_d_)→{(I,J,τ_x_,τ_y_)}] representing this neural network (see Figure S2-10). The creation procedure is shown below based on Section **A9.2** of Appendix **9**.

Before that, the notation of this table is explained. Subscripts (_distance_, ^BWD^, and _SDC_) indicate the shortest distance, the backward neural network, and the SDC, respectively, and (k,l,h_d_)→{(I,J,τ_x_,τ_y_)} indicates a connection from each SDC (k,l,h_d_) to the corresponding MDCs {(I,J,τ_x_,τ_y_)}.

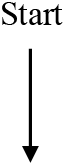

1. Give a movement direction v (see Figure A9(A))
  ➀ With respect to the center of the visual field O, if the movement direction v is (O •v)>0, it is an approaching movement, and if (O•v)<0, it is a separating movement

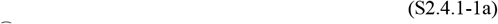
  ➁ Calculate the P_∞_ by equation(1.0-1)

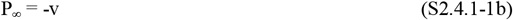 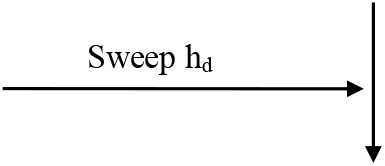 The sweep h_d_ of a SDC (k,l,h_d_) is performed by the following equation (see Section **S1.3.2**(1)).

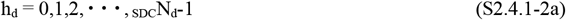

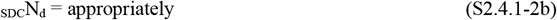
  Convert h_d_ to d (see Section **S1.3.2**(1))

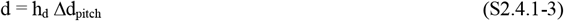 Here,

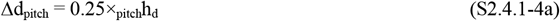

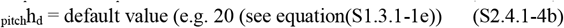 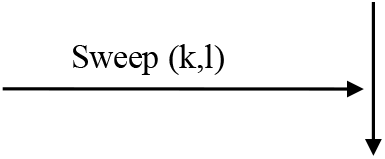 The sweep (k,l) of a SDC (k,l,h_d_) is performed by equations(S2.3.1-5a ∼6d) in Section **S2.3.1**.
  Calculate the 3D orientation n_kl_ (see Section **S1.3.2**(1))

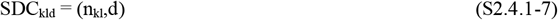

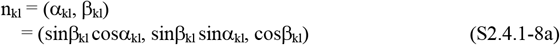 Here, (α_kl_,β_kl_) is the polar coordinate of n_kl_, (sinβ_kl_ cosα_kl_, sinβ_kl_ sinα_kl_, cosβ_kl_) is its orthogonal coordinate, and parameters used above are shown as follows.

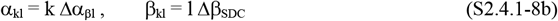

Δβ_SDC_ and Δα_βl_ were calculated by equations(S2.3.1-6b and 6e) in Section **S2.3.1**, respectively.
  Calculate the normalized time-to-contact t (see equation(A9-14) in Appendix **9**)

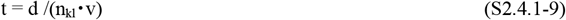 Note: The rest is the same as the normalized time-to-contact in Section **S2.3.1**. 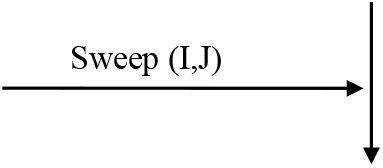 The sweep (I,J) of a RF is performed by equations(S2.1.1-1∼3d) of Section **S2.1.1**.
2. Calculate the RF center O_IJ_ (see the equations(S1.2.2-1∼4) in Section **S1.2.2**(1))

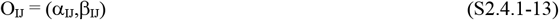

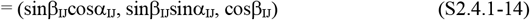 Here, (α_IJ_,β_IJ_) is the polar coordinate of O_IJ_, (sinβ_IJ_ cosα_IJ_, sinβ_IJ_ sinα_IJ_, cosβ_IJ_) is its orthogonal coordinate, and parameters used above are shown as follows.

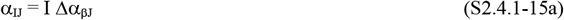

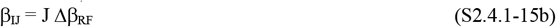

Δβ_RF_ and Δα_βJ_ were calculated by equations(S2.1.1-3b and 3h) in Section **S2.1.1**, respectively.
3. Calculate the position P_0_ at a current time (see equation(A9-15) in Appendix **9**)

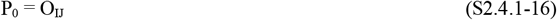
4. Calculate P_t_ (see equations(A9-16 and 17) in Appendix **9**)

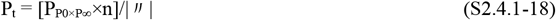

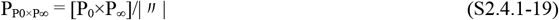
5. Calculate the polar coordinate ϕ of a MDC (see equations(A9-21∼23) in Appendix **9**)

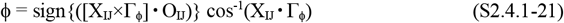 Here,

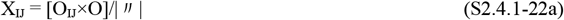

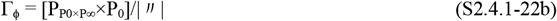
6. Calculate the polar coordinate τ_2D_ of the MDC by the cross-ratio transform (see equations(A9-18 to 20) in Appendix **9**)

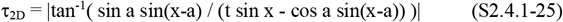 Here, the central angles a and x used above (see Figure 8(B)) are calculated by the following equations.

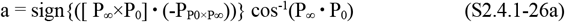

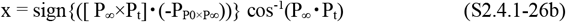
7. Calculate the address (τ_x_,τ_y_) of the MDC (see equations(A9-24 and 25) in Appendix **9**)

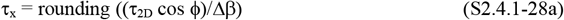

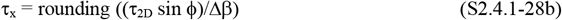 Note: This rounding serves to round to the nearest MDC, and the Δβ?used above was calculated by equation(S1.2.6.1-4).
8. When the following conditions are satisfied, write the address (I,J,τ_x_,τ_y_) to the contents of the address (k,l,h_d_) in the backward connection table _distance_T^BWD^_SDC_ [(k,l,h_d_)→{(I,J,τ_x_,τ_y_)}].

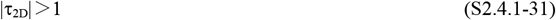

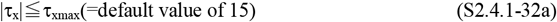

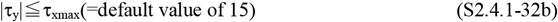 Note: This default value varies depending on the retinal cell resolution, RF size, etc. 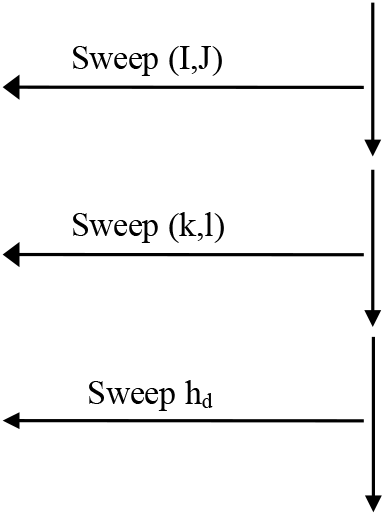
9. The backward connection table _distance_T^BWD^_SDC_ [(k,l,h_d_)→{(I,J,τ_x_,τ_y_)}] has been created (see Figure S2-10) 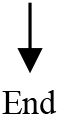

#### S2.4.2. Procedure for creating a forward connection table _distance_T^FWD^_SDC_ [(I,J,τ_x_,τ_y_)→{(k,l,h_d_)}]

Transpose the backward connection table _distance_T^BWD^ _SDC_ [(k,l,h_d_)→{(I,J,τ_x_,τ_y_)}] obtained in Section **S2.4.1** to create a forward connection table _distance_T^FWD^_SDC_ [(I,J,τ_x_,τ_y_)→{(k,l,h_d_)}] that conncts each MDC (τ_x_,τ_y_) to SDCs {(k,l,h_d_)}. The creation procedure is shown below.

Before that, the notation of this table is explained. Subscripts (_distance_, ^FWD^, and _SDC_) indicate the shortest distance, the forward neural network, and the SDC, respectively, and (I,J,τ_x_,τ_y_)→{(k,l,h_d_)} indicates a connection from each MDC (I,J,τ_x_,τ_y_) to the corresponding SDCs {(k,l,h_d_)}.

Note the following. Section **3.2** showed how to create the forward connection table using the forward method without going through this backward method. However, this forward method is more complex than the backward method, as described in Section **4.3.3.2**(3). Therefore, I decided in Section **S2.4.1** above to use the backward method in Section **A9.2** of Appendix **9**.

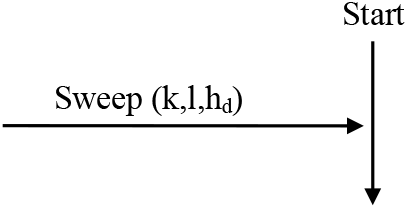

The sweeps h_d_ and (k,l) of a SDC (k,l,h_d_) are performed by equations(S2.4.1-2a and 2b) in Section **2.4.1** and equations(S2.3.1-5a ∼6d) in Section **S2.3.1**, respectively.

1. Using the backward connection table _distance_T^BWD^_SDC_[(k,l,h_d_)→{(I,J,τ_x_,τ_y_)}] in Section **S2.4.1**, read the addresses {(I,J,τ_x_,τ_y_)} of MDCs corresponding to the address (k,l,h_d_) of each SDC.
2. Write this address (k,l,h_d_) of the SDC into the contents for addresses {(I,J,τ_x_,τ_y_)}, which were read above, of the forward connection table _distance_T^FWD^_SDC_ [(I,J,τ_x_,τ_y_)→{(k,l,h_d_)}]. This writing has yielded the backward connection table to be transposed into the forward connection table. 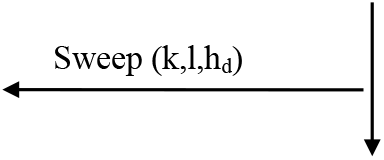
3. The forward connection table _distance_T^FWD^ [(I,J,τ_x_,τ_y_)→{(k,l,h_d_)}] has been created. The descriptions of the subsections (1) and (2) above can be expressed in another way as follows. That is, by transposing (k,l,h_d_) and (I,J,τ_x_,τ_y_) of _distance_T^BWD^_SDC_ [(k,l,h_d_)→{(I,J,τ_x_,τ_y_)}] in Section **S2.4.1**, we can get _distance_T^FWD^_SDC_ [(I,J,τ_x_,τ_y_)→{(k,l,h_d_)}]. 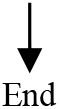

#### S2.4.3. Procedure for creating a weighting factor _distance_W_(I,J,τx,τy,hd)_

The divergent neural network, indicated by red straight lines in Figure S2-10, varies considerably from MDC to MDC. This weighting factor _distance_W_(I,J,τx,τy,hd)_ is needed to correct for that variation. That is, each SDC response is corrected so that it is not affected by the extent of this divergence (i.e. the number of red straight lines).

For that purpose, the number of these straight lines will be counted as a variable _W_N_(I,J,τx,τy,hd)_ in equation(S2.4.3-1) later: this variable corresponds to the area of the red ring in Figure 26(A)(ii). Then, the weighting factor _distance_W_(I,J,τx,τy,hd)_ will be calculated by equation(S2.4.3-2) as the reciprocal of this variable _W_N_(I,J,τx,τy,hd)_. The procedure for calculating this weighting factor will be shown below.

Note the following. This weighting factor will be used in Section **S3.3.1** as follows. By multiplying each MDC response by this weighting factor, the response is corrected so that it does not depend on the number of the red straight lines above. This correction is performed using Figure S2-10 as follows. Let us make the following correction so that each MDC response in the (A) is output to the red SDCs in the (B), regardless of the number of the red straight lines. When the number of straight lines is large, the MDC response is output weakened in inverse proportion to the number, and conversely, when the number of straight lines is small, the MDC response is output strengthened in inverse proportion to the number. In this way, a correction is made so that each MDC outputs a constant response to the red SDCs.

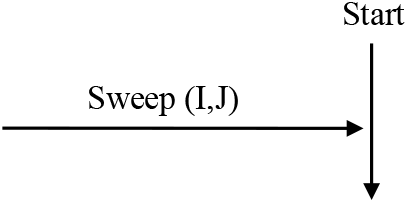

The sweep (I,J) of a RF is performed by equations(S2.1.1-1∼3d) of Section **S2.1.1**.

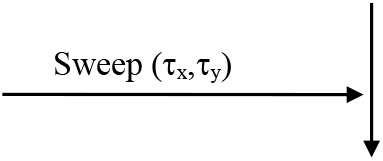

The sweep (τ_x_,τ_y_) of a MDC is performed by equations(S2.2.1-3a∼3c) of Section **S2.2.1**.

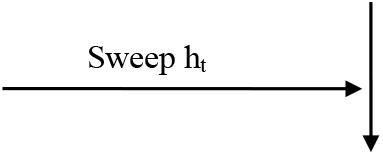

The sweep h_d_ of a SDC (k,l,h_d_) is performed by equations(S2.4.1-2a∼2b) of Section **S2.4.1**.

1. Using the forward connection table _distance_T^FWD^_SDC_[(I,J,τ_x_,τ_y_)→{(k,l,h_d_)}] in Section **S2.4.2**, read the addresses {(k,l,h_d_)} of the SDCs corresponding to the address (I,J,τ_x_,τ_y_) of each MDC.
2. Calculate the number of SDC addresses {(k,l)} at height h_d_, read above, and write the number to a variable _W_N_(I,J,τx,τy,hd)_. _W_N_(I,J,τx,τy,hd)_ = the number of SDC addresses {(k,l)} at height h_d_

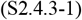
3. Determine the weighting factor _distance_W_(I,J,τx,τy,hd)_ by the following equation

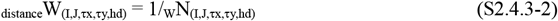 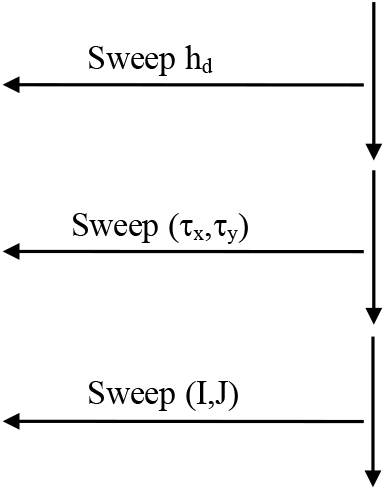
4. The weighting factor _distance_W_(I,J,τx,τy,hd)_ has been created 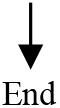

## S3. Procedure for calculating the responses of the series of cells

Using the connection table created in Section **S2**, a procedure for calculating responses of the series of cells in Figures 9 or 17 are described in two steps. In the first step, Section **S3.1** will show a procedure for calculating the series of cell responses up to MDC in Figure 9(B) or 17(B) in the form of a flowchart. In the second step, Sections **S3.2** and **S3.3** will show procedures for calculating the SDC responses in Figures 9(C) and 17(C), respectively, in the form of a flowchart.

### S3.1. Response of the series of cells from retinal cells to MDCs

Using the connection tables created in Sections **S2.1** and **S2.2**, a procedure for calculating responses of the series of cells in Figure S2-1 are shown below: Figure S2-1 was obtained by enlarging Figure 9(B) or 17(B). Previous simulation results (Kawakami & Okamoto, 1996a, 1995a; Kawakami, 1996b; Akima et al., 2017; Kawakami et al., 2003, 2000) were also calculated using this procedure.

#### S3.1.0. Responses Retina(i,j) of retinal cells

A response of each retinal cell in Figure S2-1(a) (or Figure 9(B)(a)) is represented as Retina(i,j) using the address representaion (i,j) of Figure S1-1(B). By giving this response Retina(i,j), responses of the series of cells in Figure 9(B) or 17(B) to that responses are calculated below.

#### S3.1.1. Responses of LGN cells: 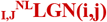 LGN(i,j) and 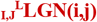 LGN(i,j)

LGN cells have two roles. One is the time delay, which causes the array of retinal cells in Figure S2-1(a) to diverge into two. That is, this array branches to an array ^NL^Retina(x,y) without time delay (i.e. nonlagged) in the upper part of (b) and an array ^L^Retina(x,y) with time delay (i.e. lagged) in the lower part: subscripts ^NL^ and ^L^ indicate Nonlagged and Lagged, respectively. Although these arrays are made up of LGN cells rather than retinal cells, the names of retinal cells have been kept to make equations(S3.1.1-1a and 1b) easier to see. The other is the 2D convolution, which is performed between each array of retinal cells and the DOG filter DOG(u,v) to emphasize the contours of figures in the (b). This convolution for the contour enhancement, which is expressed by the following equations, yields the LGN cell responses of ^NL^LGN(x,y) and ^L^LGN(x,y) (Kawakami & Okamoto, 1996a; Kawakami, 1996b).

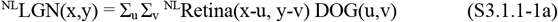

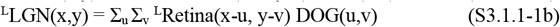

Here, the DOG filter used above is represented by the following equation.

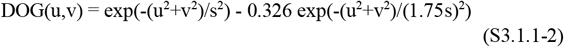

Since these convolutions are performed for each RF O_IJ_, attaching the subscript _IJ_ to ^NL^LGN(x,y) and ^L^LGN(x,y) yields ^NL^LGN(x,y) and _I,J_^L^LGN(x,y), respectively.

Expressing the responses of these LGN cells in the address representation of Figure S1-1(B) yields 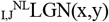 and 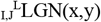. A procedure for calculating these cell responses are shown below in flowchart format: 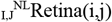 and 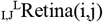 are the address representations of 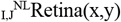 and 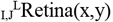, respectively.

Before that calculation, the following precaution is pointed out. In other words, it was pointed out that this 2D convolution can be executed with 1D convolution, and thus you can easily perform this 2D convolution. This pointing out is due to the fact that the Hough transformation after 2D convolution is equivalent to the 1D convolution after the Hough transform (page 162 of Kawakami (1996b); Kawakami et al., 1992c; Kawakami & Okamoto, 1996c). This 1D convolution can be executed by replacing the order (a), (b), and (c) in Figure S2-1 with the order (a), (c), and (b). Furthermore, this 1D convolution can be executed with a simpler convolution using a skeleton filter (Kawakami et al., 1992a, 1992c, 1991).

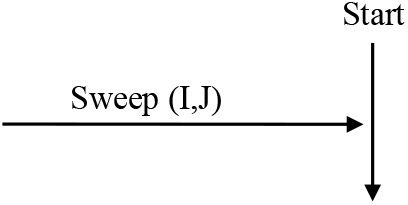

The sweep (I,J) of a RF is performed by equations(S2.1.1-1∼3d) in Section **S2.1.1**.

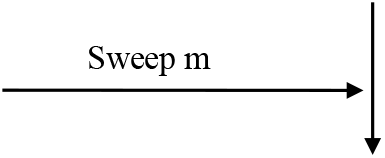

The sweep m of a serial number for cutting out is performed by equation(S2.1.1-13) of Section **S2.1.1**.

Using the cutout table _I,J_T_RF_[m→(i,j)] created in Section **S1.2.4**, read an address (i,j) of a retinal cell corresponding to the serial number m. Recall that this retinal cell is actually a LGN cell, but to make equations(S3.1.1-1a and 1b) easier to see, only the name has been kept to be retinal cells: this was described above.
Execute the 2D-convolution operation between the DOG filter of equation(S3.1.1-2) and the retinal-cell response _I,J_^L^Reina(i,j) and _I,J_^NL^Reina(i,j) using the following equations. This execution gives the cell responses _I,J_^NL^LGN(i,j) and _I,J_^L^LGN(i,j).

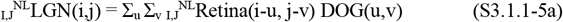

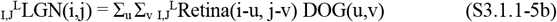 Here, Σ_u_ and Σ_v_ used above indicates an accumulation with respect to u and v, respectively. The accumulation ranges of u and v are shown as follows.

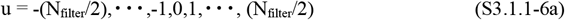

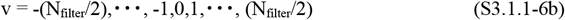 The parameter N_filter_ used above was given by equation(S1.2.6.1-20b) in Section **S1.2.6.1**(2). 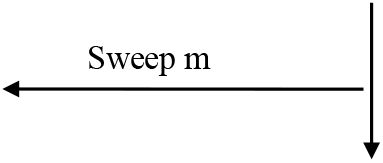

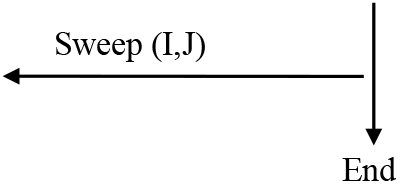

#### S3.1.2. Procedure for calculating responses of NDS simple cells: 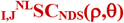 and 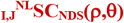

A response SC_NDS_(ρ,θ) of each NDS simple cell in Figure S2-1(c) is obtained as the Hough transformation of LGN cell responses LGN(i,j) in Figure S2-1(b): note that (i,j) is an address representation of (x,y) (see Figure S1-1). Sections **S3.1.2.1** and **S3.1.2.2** will show procedures for calculating the responses 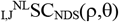 of NDS simple cells as the Hough transformation of responses 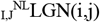 of LGN cells that were calculated in Section **S3.1.1**. Replacing the subscript ^NL^ below with the subscript ^L^ enables a procedure for calculating the responses 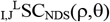 of NDS simple cells to be obtained.

##### S3.1.2.1. Forward method

A response 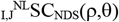 of each NDS simple cell to the LGN-cell responses 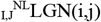 is calculated using the forward connection table 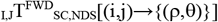 created in Section **S2.1.2**. The calculation procedure is as follows.

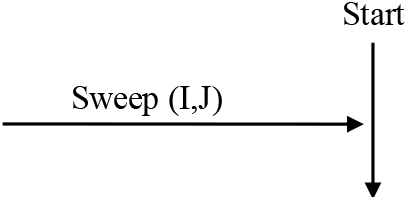

The sweep (I,J) of a RF is performed by equations(S2.1.1-1∼3d) in Section **S2.1.1**.

1. Reset responses _I,J_^NL^_NDS_ SC (ρ,θ) of all NDS simple cells for (ρ,θ)

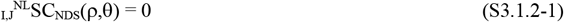 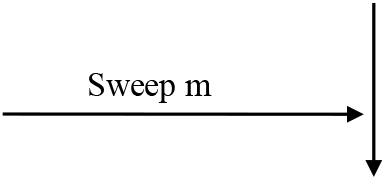 The sweep m of a serial number for cutting out is performed by equation(S2.1.1-13) of Section **S2.1.1**.
2. Using the cutout table _I,J_T_RF_[m→(i,j)] created in Section **S1.2.4**, read an address (i,j) of a LGN cell corresponding to the serial number m.
3. Using the forward connection table _I,J_T^FWD^_SC,NDS_ [(i,j)→{(ρ,θ)}] created in Section **S2.1.2**, read the addresses {(ρ,θ)} of NDS simple cells corresponding to the address (i,j) read above.
4. Add the response _I,J_^NL^LGN(i,j) of this LGN cell to the the responses {_I,J_^NL^SC_NDS_(ρ,θ)} of the NDS simple cells corresponding to the addresses {(ρ,θ)} read above. 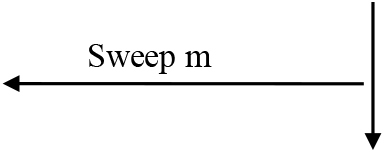
5. Responses _I,J_^NL^SC_NDS_ (ρ,θ) of NDS simple cells belonging to each RF O_IJ_ have been obtained. 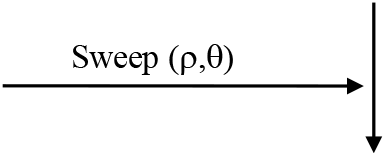 The sweep (ρ,θ) of a NDS simple cell is performed by equations(S2.1.1-7∼11) of Section **S2.1.1**.
6. Apply the weighting factor W_(I,J,ρ,θ)_ calculated by equation(S2.1.1-28) to the response _I,J_^NL^SC_NDS_(ρ,θ). That is, _I,J_^NL^SC_NDS_(ρ,θ) is multiplied by W_(I,J,ρ,θ)_ to obtain a new response of NDS simple cell.

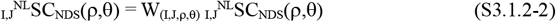 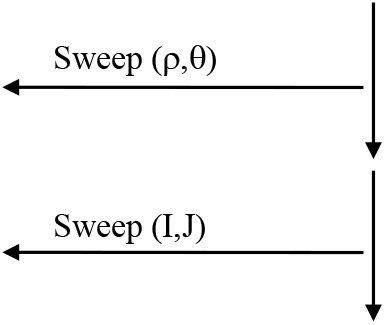
7. The responses _I,J_^NL^SC_NDS_ (ρ,θ) of NDS simple cells belonging to all RF O_IJ_ have been calculated. 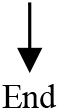

##### S3.1.2.2. Backward method

The response 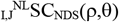 of each NDS simple cell is calculated by the following procedures using the backward connection table 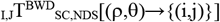 created in Section **S2.1.1**. This backward method is suitable for evaluating a specific NDS simple cell.

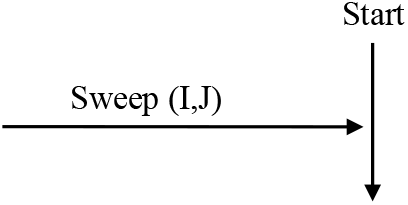

The sweep (I,J) of a RF is performed by equations(S2.1.1-1∼3d) in Section **S2.1.1**.

1. Reset responses 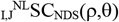 of all NDS simple cells for (ρ,θ)

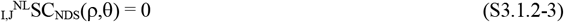 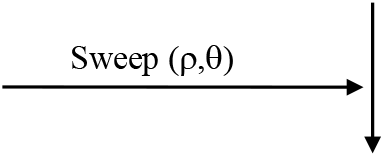 The sweep (ρ,θ) of a NDS simple cells is performed by equations(S2.1.1-7∼11) in Section **S2.1.1**.
2. Using the backward connection table _I,J_T^BWD^ _SC,NDS_ [(ρ,θ)→{(i,j)}] in Section **S2.1.1**, read the addresses {(i,j)} of LGN cells corresponding to the address (ρ,θ) of each NDS simple cell: these addresses {(i,j)} correspond to all addresses{(i_m_’,j_m_’)} within the red rectangle in Figure S2-5(A) (see equations(S2.1.1-22 and 23)).
3. Add the LGN-cell responses { _I,J_^NL^LGN(i,j)} corresponding to these addresses {(i,j)}, read above, to the response of _I,J_^NL^SC_NDS_ (ρ,θ) of this NDS simple cell.
4. Apply the weighting factor W_(I,J,ρ,θ)_ calculated in Section **S2.1.1**(6) to this response _I,J_^NL^SC_NDS_ (ρ,θ). That is, _I,J_^NL^SC_NDS_ (ρ,θ) is multiplied by W_(I,J,ρ,θ)_ to obtain a new response of NDS simple cell.

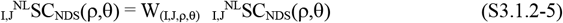 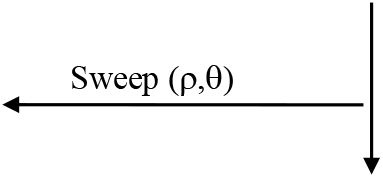
5. The responses _I,J_^NL^SC_NDS_ (ρ,θ) of NDS simple cells belonging to each RF O_IJ_ have been calculated. 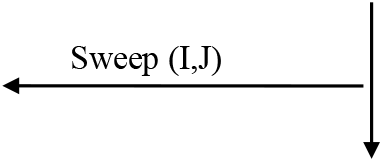
6. The responses _I,J_^NL^SC_NDS_ (ρ,θ) of NDS simple cells belonging to every RF O_IJ_ have been calculated.

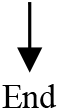

#### S3.1.3. Procedure for calculating responses of DS simple cells: _I,J_SC_DS_(ρ,θ,τ)

A response 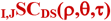 of each DS simple cell in Figure S2-1(d) is obtained as the spatio-temporal correlation between the responses (i.e. 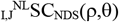 and 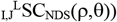 of NDS simple cells that were calculated in Section **S3.1.2**. The procedure for calculating this correlation is as follows.

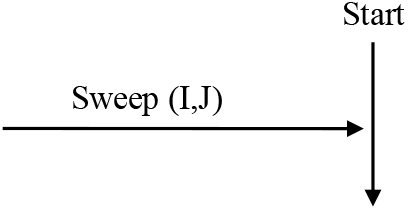

The sweep (I,J) of a RF is performed by equations(S2.1.1-1∼3d) in Section **S2.1.1**.

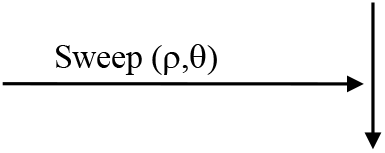

The sweep (ρ,θ) of a DS simple cell (ρ,θ,τ) is performed by equations(S2.1.1-7∼11) in Section **S2.1.1**.

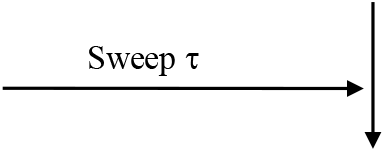

The sweep τ of a DS simple cell (ρ,θ,τ) is performed by equations(S2.2.1-8a ∼8b) in Section **S2.2.1**.

1. Using the following equation, perform a spatio-temporal correlation between the responses above (i.e. _I,J_^NL^SC_NDS_ (ρ,θ) and _I,J_^L^SC_NDS_ (ρ,θ)) to obtain the response SC (ρ,θ,τ) of a DS simple cell.

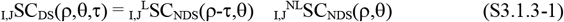 Note the following. Equation(S3.1.3-1) uses a multiplication for this correlation operation, but there is a problem in performing the multiplication physiologically. Therefore, we proposed a multiplication-like operation that combines the presynaptic inhibition and the postsynaptic excitation and inhibition (Kawakami, 1996b; Kawakami & Okamoto, 1995b). In this operation, addition of absolute values is performed with the same sign operation as multiplication. For this reason, this multiplication-like operation should be used for correlation, but the multiplication has been used in this flowchart because it is almost functionally equivalent. For details on these, see Section **2.1.2**(4). 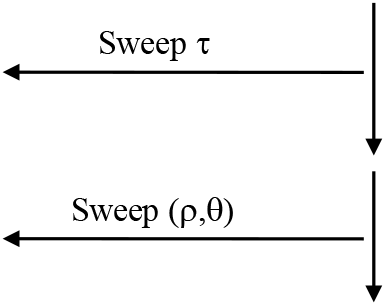
2. The responses _I,J_SC_DS_(ρ,θ,τ) of DS simple cells belonging to each RF O_IJ_ have been calculated. 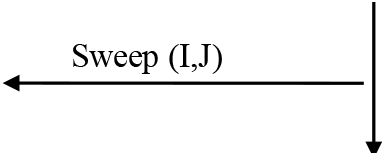
3. The responses _I,J_SC_DS_(ρ,θ,τ) of DS simple cells belonging to every RF O_IJ_ have been calculated. 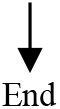

#### S3.1.4. Procedure for calculating responses of DS complex cells: _I,J_CC_DS_(θ,τ)

A response _I,J_CC_DS_(θ,τ) of each DS complex cell in Figure S2-1(e) is obtained as the accumulation of the responses _I,J_SC_DS_(ρ,θ,τ) of DS simple cells in the (d): this accumulation is performed for ρ. A neural network that is shown by the pink and blue straight lines of the (e) performs this accumulation to collect responses of all DS simple cells on each straight line of the (d), and causes the pink and blue cells in the (e) to fire strongly. Using this accumulation, the procedure for calculating the responses _I,J_CC_DS_(θ,τ) of the DS complex cells in the RF O_IJ_ is shown below.

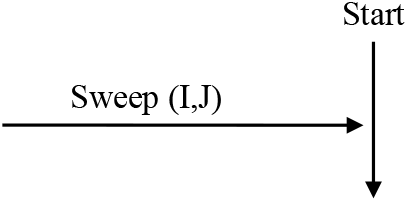

The sweep (I,J) of a RF is performed by equations(S2.1.1-1∼3d) in Section **S2.1.1**.

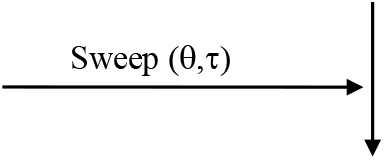

The sweeps θ and τ of a DS complex cell (θ,τ) are performed by equations(S2.2.1-4∼5b and S2.2.1-8a∼8b) in Section **S2.2.1**, respectively.

1. Accumulate _I,J_SC_DS_(ρ,θ,τ) for ρ by the following equation.

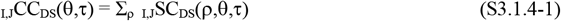 Here, the accumulation range of ρ is the following equation (see equation(S2.1.1-7)).

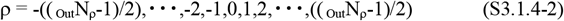

_Out_N_ρ_ used above was calculated by equation(S1.2.6.1-43). 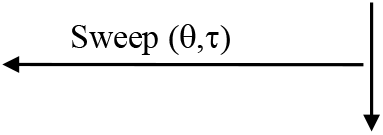
2. Responses _I,J_CC_DS_(θ,τ) of the complex cells belonging to each RF O_IJ_ have been calculated. 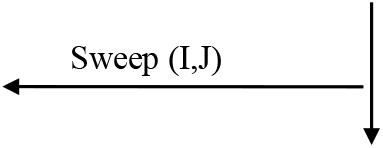
3. Responses _I,J_CC_DS_(θ,τ) of the complex cells belonging to every RF O_IJ_ have been calculated. 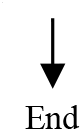

#### S3.1.5. Procedure for calculating responses of MDCs: _I,J_MDC(τ_x_,τ_y_)

A response _I,J_MDC(τ_x_,τ_y_) of each MDC in Figure S2-1(f) is obtained as the inverse Hough transformation of the DS-complex-cell responses _I,J_CC_DS_(θ,τ) in the (e). The neural network that is shown by straight lines including pink and blue in the (f) performs this transformation to collect the responses of all DS complex cells on the sine wave in the (e), and causes a red cell in the (f) to fire strongly. Using this transformation, the procedure for calculating the responses _I,J_MDC(τ_x_,τ_y_) of MDCs in every RF O_IJ_ is shown below.

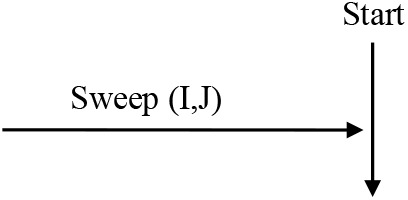

The sweep (I,J) of a RF is performed by equations(S2.1.1-1∼3d) in Section **S2.1.1**.

1. Reset responses _I,J_MDC(τ_x_,τ_y_) of all MDCs for (τ_x_,τ_y_)

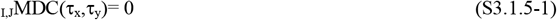

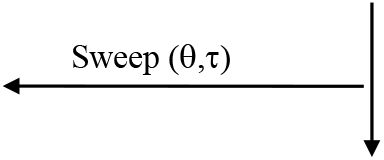 The sweeps θ and τ of a DS complex cell (θ,τ) are performed by equations(S2.2.1-4∼5b and S2.2.1-8a∼8b) in Section **S2.2.1**, respectively.
2. Using the forward connection table T^FWD^_MDC_[(θ,τ)→{(τ_x_,τ_y_)}] created in Section **S2.2.2**, read the addresses {(τ_x_,τ_y_)} of MDCs corresponding to the address (θ,τ) of each complex cell.
3. Add the response _I,J_CC_DS_(θ,τ) of this complex cell to the responses {_I,J_MDC(τ_x_,τ_y_)} of the MDCs corresponding to the addresses {(τ_x_,τ_y_)} read above. 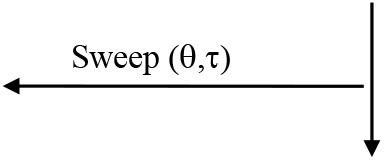
4. The responses _I,J_MDC(τ_x_,τ_y_) of MDCs belonging to each RF O_IJ_ have been obtained. 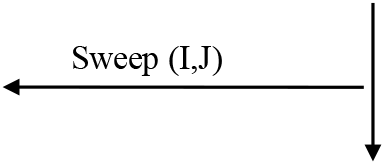
5. The responses _I,J_MDC(τ_x_,τ_y_) of MDCs belonging to every RF O_IJ_ have been calculated. 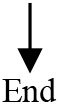

#### S3.2. Responses SDC(k,l,h_t_) of SDCs detecting the time-to-contact and 3D orientation

The procedure for calculating the responses SDC(k,l,h_t_) of SDCs using the following forward connection table and weighting factor is shown below. The simulations reported so far (Kawakami et al., 2003, 2000) were also performed using this procedure.

- Forward connection table created in Section **S2.3.2**: _time_T^FWD^_SDC_ [(I,J,τ_x_,τ_y_)→{(k,l,h_t_)}]
- Weighting factor created in Section **S2.3.3**: _time_W_(I,J,τx,τy,ht)_

#### S3.2.1. Procedure for correcting the MDC responses by the weighting factor

This weighting factor _time_W_(I,J,τx,τy,ht)_ was created as equation(S2.3.3-2) in Section **S2.3.3**. Although the divergence of the neural network (i.e. the number of red straight lines) in Figure S2-8 varies considerably from MDC to MDC, this factor can correct for that variation. That is, by multiplying the MDC response with this factor, the response can be corrected so that it is not affected by the variation. A procedure for performing this correction is shown below.

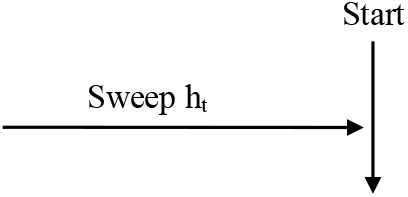

The sweep h_t_ of a SDC (k,l,h_t_) is performed by equations(S2.3.1-2a∼2b) of Section **S2.3.1**.

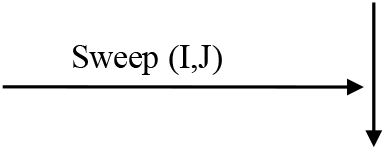

The sweep (I,J) of a RF is performed by equations(S2.1.1-1∼3d) in Section **S2.1.1**.

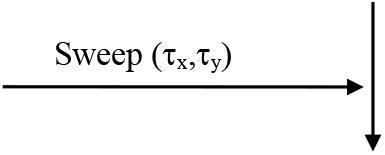

The sweep (τ_x_,τ_y_) of a MDC is performed by equations(S2.2.1-3a∼3c) of Section **S2.2.1**.

1. Calculate the corrected MDC responses _I,J,ht_^correct^MDC(τ_x_,τ_y_) by multiplying the MDC responses _I,J_MDC(τ_x_,τ_y_) calculated in Section **S3.1.5** with the weighting factor _time_W_(I,J,τx,τy,ht)_.

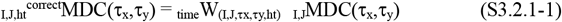 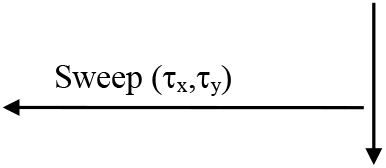

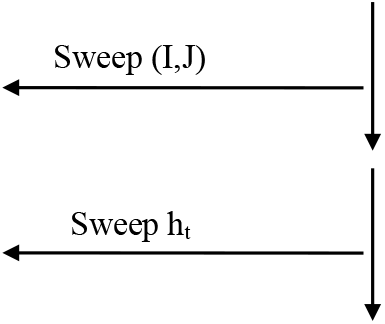
2. The corrected MDC responses _I,J,ht_^correct^MDC(τ, τ) have been calculated. 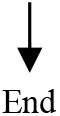

#### S3.2.2. Procedure for calculating SDC responses SDC(k,l,h_t_)

Calculate the SDC response SDC(k,l,h_t_) by applying the forward connection table _time_T^FWD^_SDC_ [(I,J,τ_x_,τ_y_)→{(k,l,h_t_)}] created in Section **S2.3.2** to the MDC response _I,J,ht_^correct^MDC(τ_x_,τ_y_) corrected in Section **S3.2.1**. The procedure for this calculation is as follows.

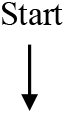

(1) Reset responses SDC(k,l,h_t_) of all SDCs for (k,l,h_t_)

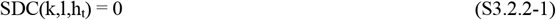 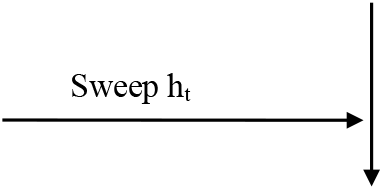 The sweep h_t_ of a SDC (k,l,h_t_) is performed by equations(S2.3.1-2a∼2b) of Section **S2.3.1**. 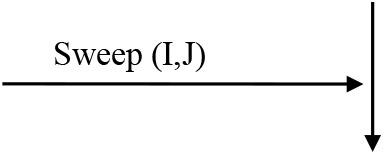 The sweep (I,J) of a RF is performed by equations(S2.1.1-1∼3d) in Section **S2.1.1**. 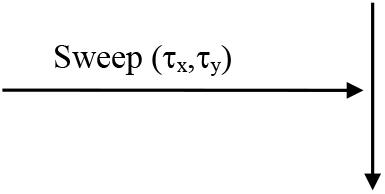 The sweep (τ_x_,τ_y_) of a MDC is performed by equations(S2.2.1-3a∼3c) of Section **S2.2.1**.
(2) Using the forward connection table _time_T^FWD^_SDC_ [(I,J,τ_x_,τ_y_)→{(k,l,h_t’_)}] created in Section **S2.3.2**, read the addresses {(k,l,h_t’_)} of SDCs corresponding to the address (I,J,τ_x_,τ_y_) of each MDC: for convenience of notation, h_t_ used so far has been changed to h_t’_.
(3) Among the above addresses {(k,l,h _t_^‘^)}, select addresses ({(k,l)},h_t_) that correspond to the cross section of the column (k,l,h_t’_) at h_t’_= h_t_: these addresses ({(k,l)},h_t_) correspond to a group of the addresses {(k,l)} shown in red dots in Figure S2-8(B).
(4) Add the MDC response _I,J,ht_^correct^MDC(τ_x_,τ_y_) corrected in Section **S3.2.1** to the SDC responses {SDC(k,l,h_t_)} corresponding to the addresses ({(k,l)},h_t_) selected above. 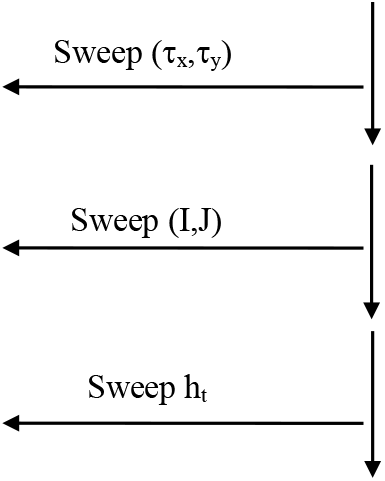
(4) The SDC responses SDC(k,l,h_t_) have been calculated. 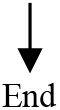

### S3.3. Responses SDC(k,l,h_d_) of SDCs detecting the shortest distance and 3D orientation

The procedure for calculating the responses SDC(k,l,h_d_) of SDCs using the following forward connection table and weighting factor is shown below. The simulations reported so far (Kawakami et al., 2003, 2000) were also performed using this procedure.

- Forward connection table created in Section **S2.4.2**: _distance_T^FWD^_SDC_ [(I,J,τ_x_,τ_y_)→{(k,l,h_d_)}]
- Weighting factor created in Section **S2.4.3**: _distance_W_(I,J,τx,τy,hd)_

#### S3.3.1. Procedure for correcting MDC responses by the weighting factor

This weighting factor _distance_W_(I,J,τx,τy,hd)_ was created as equation(S2.4.3-2) in Section **S2.4.3**. Although the divergence of the neural network (i.e. the number of red straight lines) in Figure S2-10 varies considerably from MDC to MDC, this factor can correct for that variation. That is, by multiplying the MDC response with this factor, the response can be corrected so that it is not affected by the variation. A procedure for performing this correction is shown below.

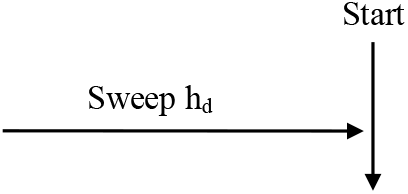

The sweep h_d_ of a SDC (k,l,h_d_) is performed by equations(S2.4.1-2a∼2b) of Section **S2.4.1**.

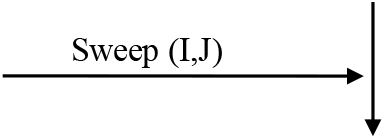

The sweep (I,J) of a RF is performed by equations(S2.1.1-1∼3d) in Section **S2.1.1**.

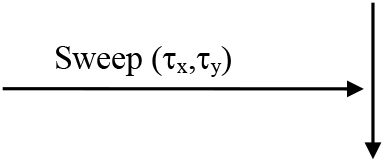

The sweep (τ_x_,τ_y_) of a MDC is performed by equations(S2.2.1-3a∼3c) of Section **S2.2.1**.

1. Calculate the corrected MDC responses _I,J,hd_^correct^MDC(τ_x_, τ_y_) by multiplying the MDC responses _I,J_MDC(τ_x_,τ_y_) calculated in Section **S3.1.5** with the weighting factor _distance_W_(I,J,τx,τy,hd)_.

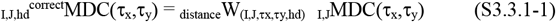 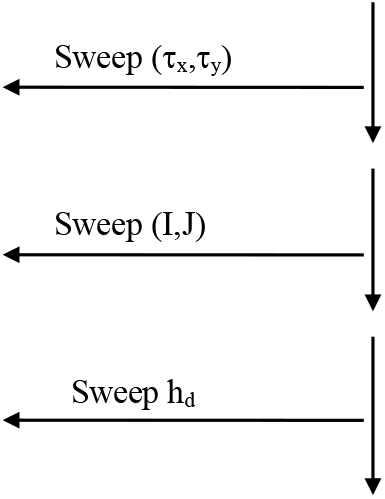
2. The corrected MDC responses _I,J,hd_^correct^MDC(τ_x_, τ_y_) have been calculated. 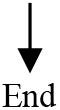

#### S3.3.2. Procedure for calculating SDC responses SDC(k,l,h_d_)

Calculate the SDC response SDC(k,l,h_d_) by applying the forward connection table _distance_T^FWD^_SDC_ [(I,J,τ_x_,τ_y_)→{(k,l,h_d_)}] created in Section **S2.4.2** to the MDC response _I,J,hd_^correct^MDC(τ_x_,τ_y_) corrected in Section **S3.3.1**. The procedure for this calculation is as follows.

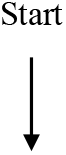

1. Reset responses SDC(k,l,h_d_) of all SDCs for (k,l,h_d_)

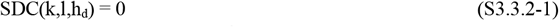 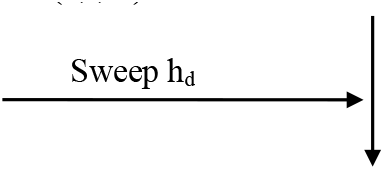 The sweep h_d_ of a SDC (k,l,h_d_) is performed by equations(S2.4.1-2a∼2b) of Section **S2.4.1**. 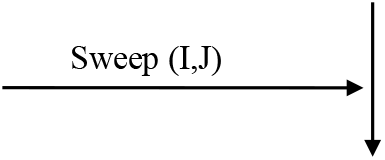 The sweep (I,J) of a RF is performed by equations(S2.1.1-1∼3d) in Section **S2.1.1**. 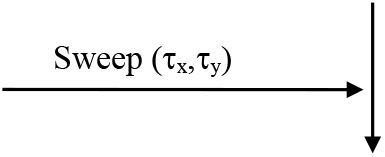 The sweep (τ_x_,τ_y_) of a MDC is performed by equations(S2.2.1-3a∼3c) of Section **S2.2.1**.
2. Using the forward connection table _dustance_T^FWD^_SDC_ [(I,J,τ_x_,τ_y_)→{(k,l,h_d_^‘^)}] created in Section **S2.4.2**, read the addresses {(k,l,h_d_^‘^)} of SDCs corresponding to the address (I,J,τ_x_,τ_y_) of each MDC: for convenience of notation, h_d_used so far has been changed to h_d_^‘^.
3. Among the above addresses {(k,l,h_d_^‘^)}, select addresses ({(k,l)},h_d_) that correspond to the cross section of the column (k,l,h_d_^‘^) at h_d_^‘^= h_d_ : these addresses ({(k,l)},h) correspond to a group of the addresses {(k,l)} shown in red dots in Figure S2-10(B).
4. Add the MDC response _I,J,hd_^correct^MDC(τ_x_, τ _y_) corrected in Section **S3.3.1** to the SDC responses {SDC(k,l,h_d_)} corresponding to the addresses ({(k,l)},h_d_) selected above. 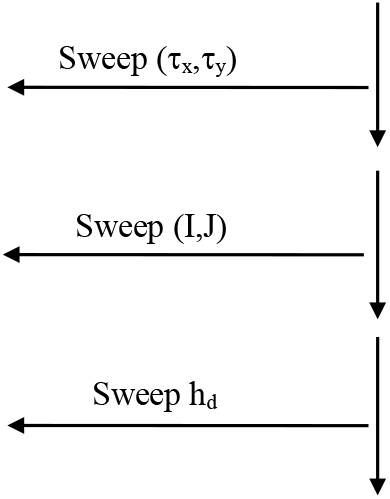
5. The SDC responses SDC(k,l,h_d_) have been calculated. 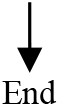

